# A Low-Cost, Modular Hardware and Software Platform for Head-Fixed Mouse Decision-Making Tasks

**DOI:** 10.64898/2026.08.03.742587

**Authors:** Maxwell B. Madden, Mahee Khatri, Arnav Mohanty, Disha Prasad, Nancy Collie-Beard, Rafiq Huda

## Abstract

Head-fixed behavior in rodents is a foundational technique in systems neuroscience which enables use of sophisticated imaging techniques in combination with animal behavior. However, accessibility of head-fixed behavior techniques is limited. Animal training consumes a large amount of experimenter labor and commercial setups, when available, are largely inflexible and financially burdensome. Here, we present a low-cost, modular, and open-source hardware and software implementation for head-fixed rodent decision-making tasks. Our design lowers experimenter labor and enables large teams of researchers to participate in animal training with minimal experimenter error using a simple touchscreen GUI and automated training progression. We demonstrate the efficacy of the platform by training a cohort of animals in a two-choice probabilistic rapid-reversal task in which mice continuously update action choices based on recent reward history. The presented design lowers the barrier to entry for laboratories seeking to conduct head-fixed rodent behavior and provides modular solutions for developing custom rigs based on experimental demands.

**Significance Statement:** Head-fixation in behaving rodents is a common neuroscience procedure that enables the use of electrical or optical techniques for neural circuit interrogation. While commercial head-fixed behavior platforms are available, their high cost and inflexibility make them unsuitable for modification or high-throughput training. This study presents a novel open-source hardware and software system for mouse head-fixed behavior. The modular design and use of low-cost components enable broad adaptability across behavioral tasks, compatibility with practical research demands, and significantly lowers the barrier to entry to head-fixed behavioral research.

## Introduction

Head-fixed behavior has become a foundational technique in systems neuroscience. Used in non-human primate research for almost 60 years (Evarts, 1966; Jasper et al., 1960), the technique has been adapted to rodents to enable the combination of highly sophisticated electrophysiological and imaging techniques with awake behavior. This expansion of head-fixed behavior into rodent has allowed the use of these neural recording techniques in combination with the genetic tools uniquely available in rodents. Additionally, head-fixation has provided access to rodent behavior modalities that are not easily accessible in their freely moving counterparts, such as musculofascial movement and pupil tracking. Both measures have been determined to be readouts for aspects of internal cognitive, affective, and arousal state (Chintalacheruvu et al., 2026; Dolensek et al., 2020; Hess & Polt, 1964; Lee & Margolis, 2016; Reimer et al., 2014; Stringer et al., 2019; Tlaie et al., 2025). These benefits come at the trade-off of increased training time, as animals must both habituate to head-fixation and learn potentially unfamiliar and nonetiological response modalities.

Great strides have been made in overcoming the behavioral limitations of head-fixed behavior, some standardization has occurred (Guo et al., 2014; Laboratory et al., 2021), and several complex navigational (Glorius et al., 2024; Stuart et al., 2024), decision-making (Jung et al., 2025; Laboratory et al., 2021), sensory discrimination (Helmchen et al., 2018), and even social (Chari et al., 2023) tasks have been developed. Despite this expansion, head-fixed behavior remains inaccessible to many labs, partly due to high financial cost of commercial head-fixation systems. Additionally, many labs develop novel tasks, or unique focused versions of existing tasks designed to isolate the specific behavior or function of interest. As a result, most head-fixed behavior apparatus is a bespoke creation of an individual lab. This has resulted in a large amount of redundant labor, as experimenters solve similar engineering problems repeatedly, thus representing an unseen burden on scientific progress. To counter this, open sharing of hardware and software solutions (Akam et al., 2022; Jung et al., 2025; Ozgur et al., 2023), especially low-cost alternatives to more costly existing systems, is vital for continued progress in the field.

Here we present a hardware and software platform for decision making tasks in mice. Our design emphasizes the use of low-cost components, thus increasing accessibility and enabling high-throughput parallelization of behavioral training. The inclusion of an optional touchscreen interface reduces the training requirements for large teams. Additionally, the system is adaptable to alternative head-fixed behavioral tasks, capable of automatic behavioral shaping, and compatible with optical and electrophysiological neural recording techniques. We demonstrate the capability of the platform using a version of an existing two-choice probabilistic rapid-reversal task (Bloem et al., 2022), in which mice must choose between two available responses to receive a water reward using only their experience of recent trials. The rewarded response is periodically switched in an uncued fashion, requiring mice to actively monitor recent reward history to make optimal choices. This hardware/software platform greatly decreases the material and labor costs associated with head-fixed behavior, increasing the accessibility and reproducibility of head-fixed behavioral experiments.

## Materials and Methods

### Animals

11 C57/BL6J (wildtype) mice were used for behavioral experiments. Mice were 7-12 weeks of age at time of stereotaxic surgery. Animals were initially housed with *ad libitum* access to food and water, with water restriction being introduced 5-10 days after surgical procedure. All mice were maintained on a reverse light/dark circadian cycle beginning at 8:00 am. All behavioral procedures were conducted in the active dark phase of the cycle, generally between zeitgeber time (ZT)14 and ZT20. All animal procedures were performed in accordance with the National Institutes of Health Guide for Care and Use of Laboratory Animals, and all procedures were approved by the Rutgers Institutional Animal Care and Use Committee (approval no. 202000004).

### Stereotaxic Surgery and Implant

Mice received surgical implantation of a custom designed headplate (eMachineShop, Figure 3B) to enable head fixation. During surgery mice were anesthetized via inhalation of isoflurane (3.5%) prior to being placed in a mouse stereotaxic frame (51500D, Stoelting). Anesthesia was subsequently maintained with inhaled isoflurane (1%) and body temperature was maintained at 37.5°C with a heating pad integrated into the base of the stereotaxic frame and a temperature controller (53800, Stoelting). Mice were given a subcutaneous injection of extended-release buprenorphine (3.25 mg/kg) prior to surgery for extended (72h) analgesia. Mice were supplemented with Meloxicam (10mg/kg) if additional analgesia was required. Scalp hair was removed using depilatory cream (Nair) and the scalp was disinfected using 3x alternating scrubs with betadine and 70% ethanol. A portion of the scalp was removed, and the skull abraded with a dental drill with a Carbon Steel 0.5mm drill bit (Fine Science Tools; No. 19007-05). The headplate was then placed over the skull and adhered in place using dental acrylic (Metabond, Parkell) mixed with mica powder. Animals were then allowed to recover in their own cage with a warm water blanket and moistened food chow. Mice were singly housed for the remainder of the experiment and were allowed at least one week of recovery before initiation of water restriction and behavioral experiments.

### Behavior Rig Construction

The behavior rig (Figure 1) was composed of three sets of hardware components; the sound attenuation chamber (Figure 1B, Figure 1-1), head restraint and behavior platform (Figure1A, Figure 1–3) and Arduino/Minicomputer controller. All components are listed in the bill of materials in extended data. Components mentioned here are labeled with an “Item Number” associated with the full component details listed in the bill of materials.

**Figure 1.**
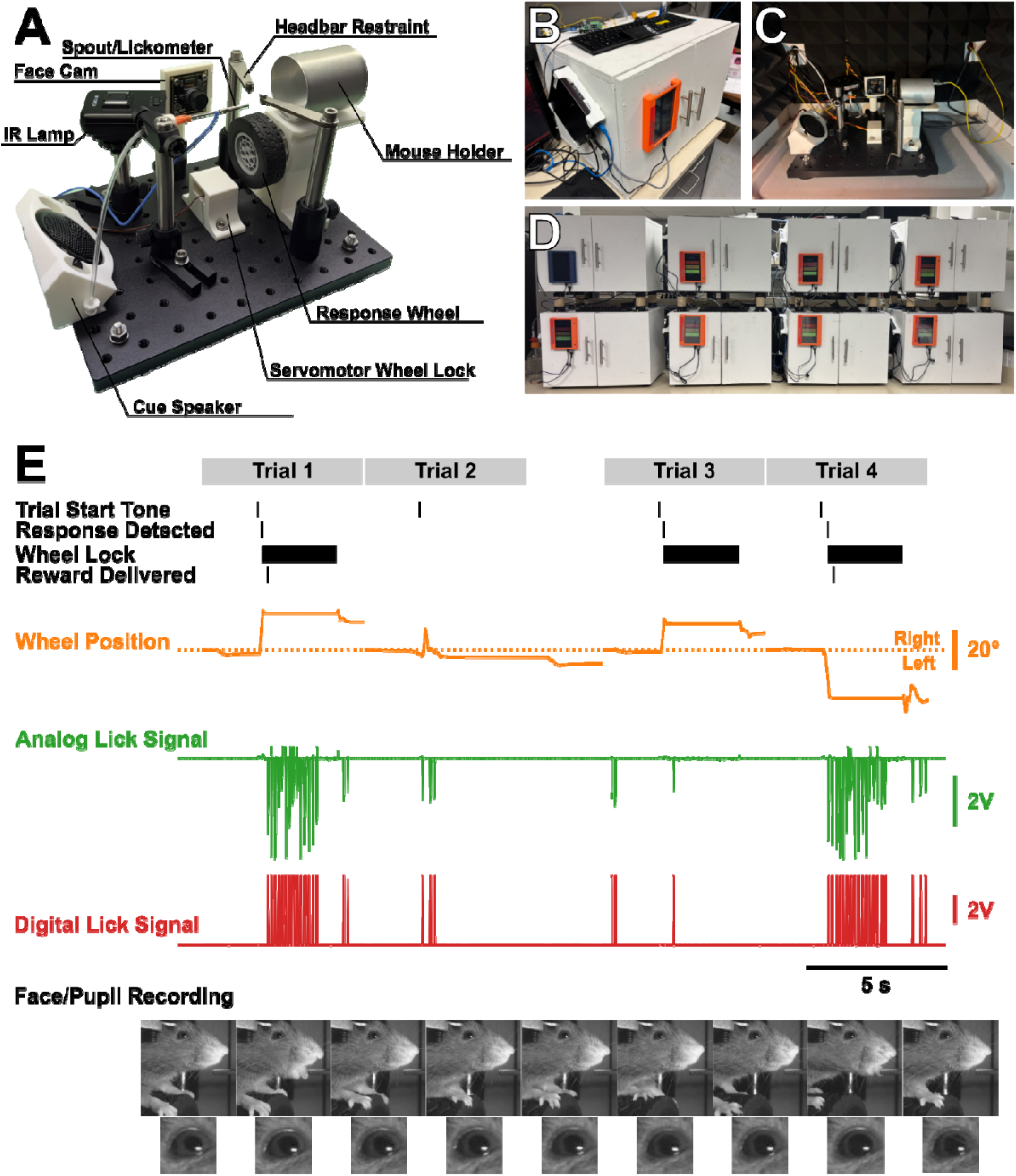
Rig Hardware Overview. **A.** Rig assembly with labeled components. **B.** Photograph of the outside of the sound attenuation chamber, including exterior mounted touchscreen interface. **C.** View of the rig assembly placed within the sound attenuation chamber. **D.** View of the stacked behavior rig systems. **E.** Representative traces of behavioral data collected by the rig including trial events (top), wheel position baselined to the 2 seconds before trail start (orange), analog (green) and digital (red) lick signals, and face/pupil recordings (bottom).

**Figure 2.**
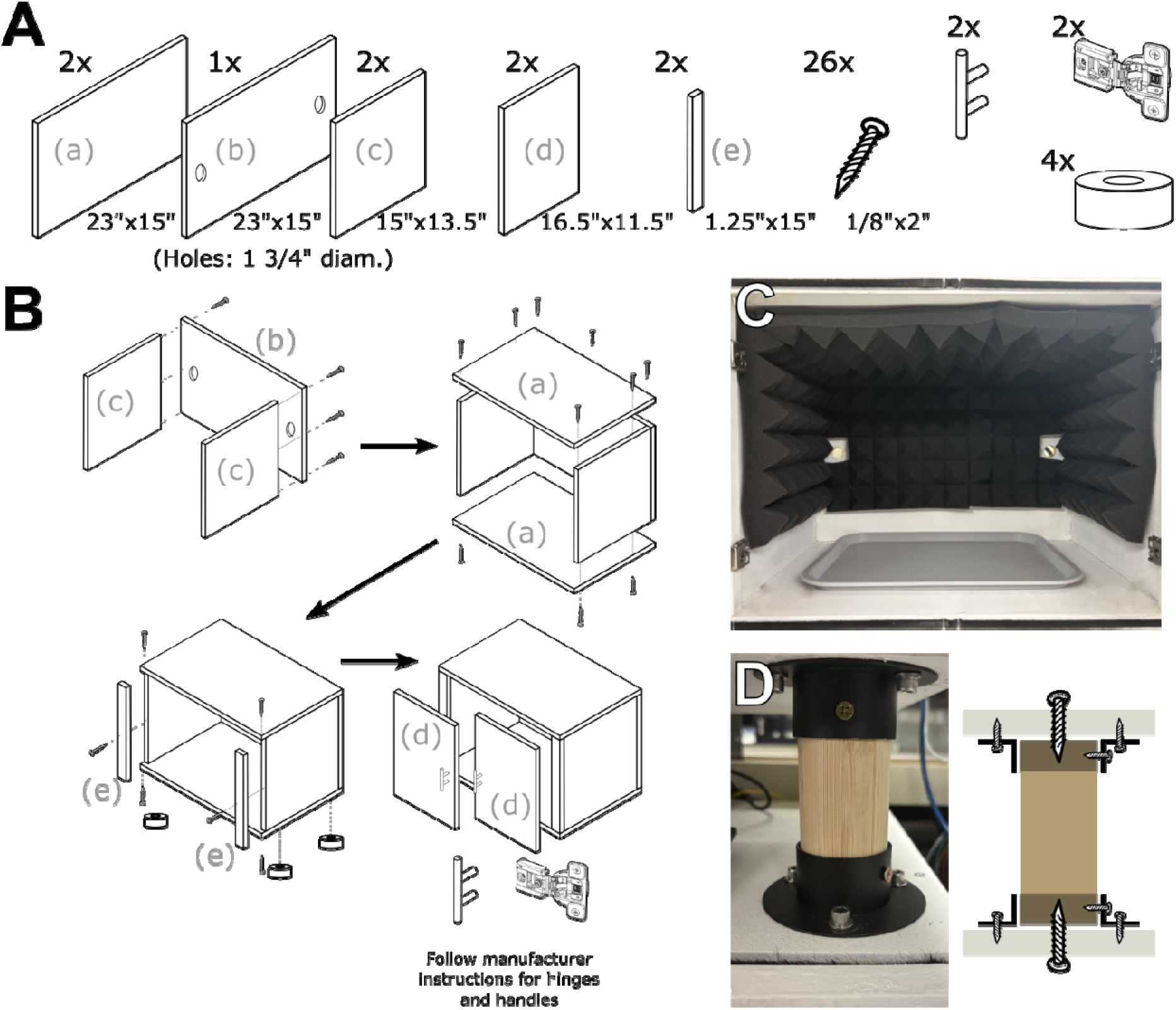
Sound Attenuation Chamber Assembly and Stacking. **A.** Cut MDF panels and materials required for assembly. **B.** Steps for assembly of sound attenuation chamber. **C.** View of sound attenuation chamber interior after installation of acoustic foam. **D.** Photograph and cartoon depiction of sound attenuation chamber stacking.

### Sound Attenuation Chamber

The sound attenuation chamber was constructed from MDF panels and components sourced from Home Depot and Amazon (Items #1-10). To construct the sound attenuation chamber, ½” thick MDF (Item #1) was cut into three 23”x15” panels, two 15”x13.5” panels, two 16”x11.5” panels, and two 1.25”x15” panels (Figure 2A). One of the 23”x15” panels had two 0.75” diameter holes drilled as cable ports, ∼2.25” inches from the edge of the panel. The sound attenuation chamber was then assembled (Figure 2B), using 1/8”x2” wood screws (Item #: 2) and wood glue (Item #3), then clamped for maximum strength of bond. After assembly, each box was painted with two coats of water-resistant primer (Item #: 4) and water-resistant paint (Item #5). Cabinet handles (Item #6) and hinges (Item #7) were installed according to manufacturer instructions, and air compressor pads (Item #8) were secured via screws as feet. A cafeteria tray (Item #9) was then placed within each sound attenuation chamber to serve as an easily removable/cleanable catch for any animal waste or leakage in water supply. Sound dampening foam (Item #10) was secured via wood staples to the inside of the sound attenuation chamber walls, ceiling and doors (Figure 2C). Sound attenuation chambers can be stacked two high to allow maximum utilization of lab space using closet rod holders (Item #11) and 2-inch diameter bamboo dowels (Item #12) cut to 4.5” of length (Figure 2D).

### Head Restraint and Behavior Platform

The head restraint and behavior platform (Figure 3) was constructed upon a Thorlabs 8”x10” aluminum breadboard (Item #13) with mounted rubber feet (Items #14,15). Thorlabs screws were used as fasteners to mount parts to the breadboard (Items #16-19). Custom components included the head-bar restraint (Item #46), mouse platform (Item #47), and rotary/wheel adapter (Item #48). The head-bar restraint (Figure 3A, left) was ordered from eMachineShop, the mouse platform was 3D printed in PLA, and the rotary encoder adapter was printed in tough resin (all files in Extended Data). It is recommended that the mouse platform be printed with the rotary encoder socket pointing upwards, to prevent splitting along layer lines. The rotary/wheel adapter should be printed with minimal supports to ensure fit.

**Figure 3.**
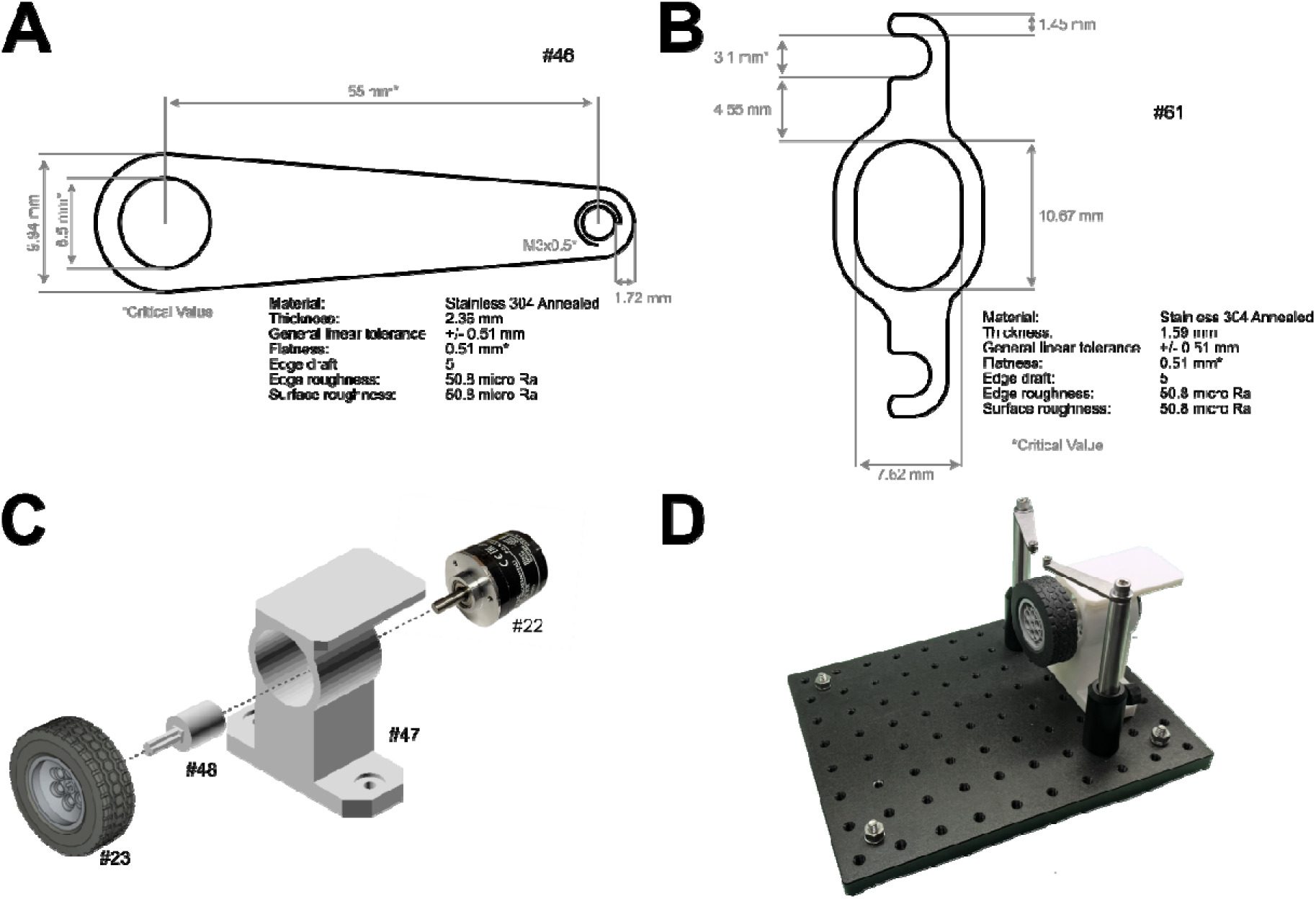
Assembly of the Head Fixation and Response Wheel Components. **A.** Specifications for the machined steel head-bars and head-plate implant. **B.** Assembly instructions for the mouse platform and response wheel. **C.** Photograph of the assembled and mounted mouse platform, response wheel, and head-bars. Item numbers are marked according to their listing in the bill of materials (Extended Data).

To construct the head restraint, the custom head-bar restraints were attached to an inverted 4” optical post (Item #20) which was then seated in a 2” post holder (Item #21). The post holder was mounted to the board using a ¾” long cap screw (Item #19), screwed through the bottom of the post holder and into the breadboard. It is vital to ensure the cap screw is fully seated to ensure positioning of the head-bar restraint at the correct height to minimize animal discomfort. The mouse platform and response wheel (Figure 3C) was constructed by seating the rotary encoder (Item #22) into the socket of the custom mouse platform (Item #47). The shaft of the rotary encoder was then joined to the response wheel (Item #23) via the rotary/wheel adapter (Item #48; Figure 3D).

Finally, an aluminum metal tube (OD: 1”, ID: 0.93”, Item #24) was attached to the top of the platform using 2” of adhesive backed hook and loop fastener (Items #25, 26). The metal tube provides an enclosure to increase animal comfort and maintain the position of the mouse’s body as well as providing an electrically conductive surface to allow lick detection.

### Water Spout

A water spout (Figure 4A), was constructed to provide water rewards to the mice, while recording consummatory licks. The water spout was constructed by first attaching a Luer Lock socket (Item #27) to a short length of 1/16” ID PVC tubing (Item #28). The tubing was then threaded through a nylon spacer (Item #29) and the Luer Lock socket and spacer were adhered together using a two-part epoxy (Item #30). A second luer lock plug (Item #31) was then inserted into the other end of the tubing to allow eventual connection to a water supply (Nalgene carboy). The water flow was controlled with a normally closed 12V solenoid valve (Item #32) joined to 1/16” tubing with a zero dead-volume PCTFE fitting (Item #33).

**Figure 4.**
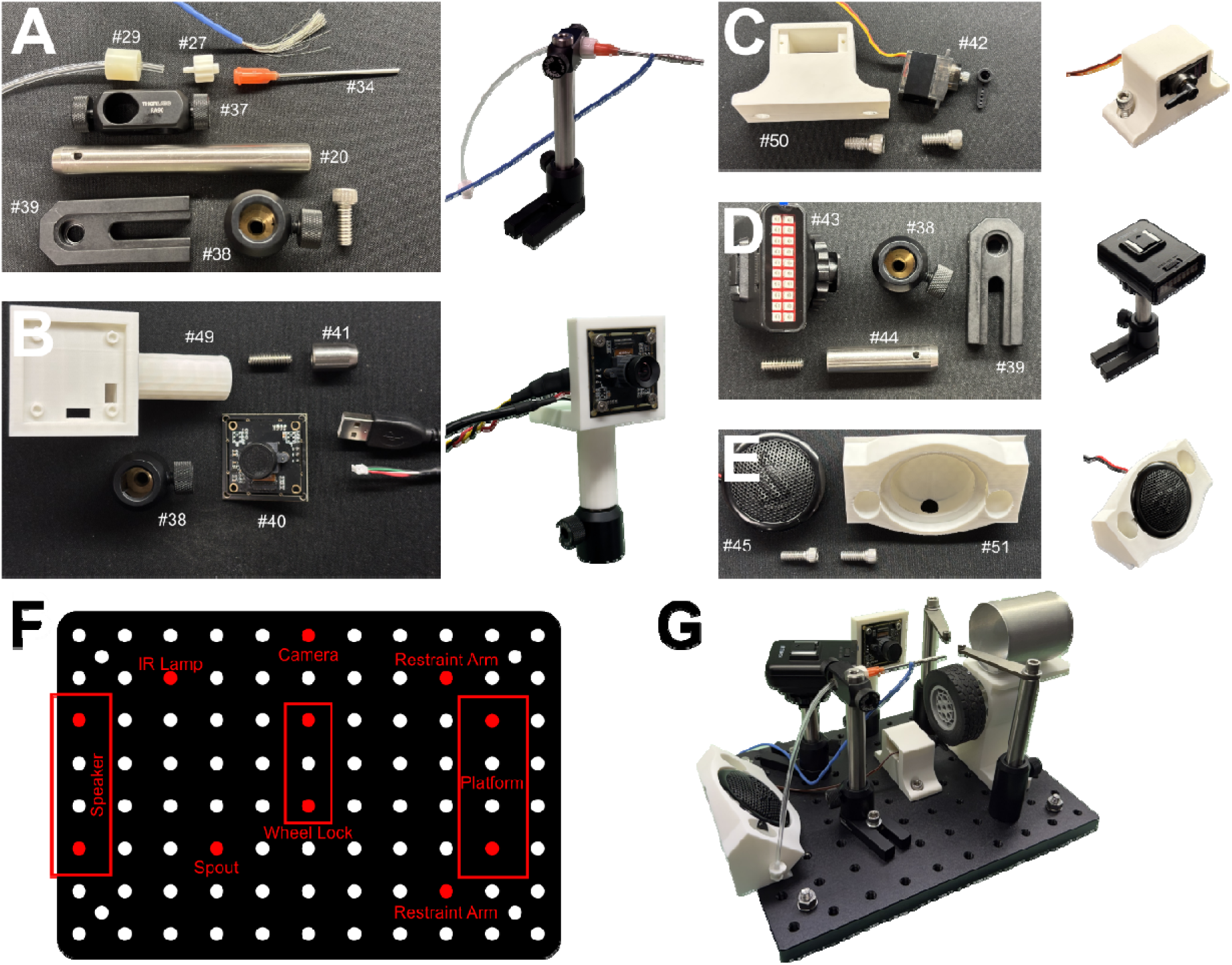
Hardware Components and Assembly. **A.** Water spout components (left) and fully assembled spout (right). **B.** Arducam camera and camera holder components (left) and fully assembled camera mount (right). **C.** Servomotor and mount for wheel lock (left) and fully assembled wheel lock (right). **D.** IR light mount components (left) and fully assembled IR light (right). **E.** Speaker and speaker mount (left) and assembled speaker assembly (right). **F.** Diagram of the mounting locations of all rig components. **G.** Photograph of all rig hardware components mounted. Item numbers are marked according to the listing in the bill of materials (Extended Data).

Next the spout, a 2” blunt stainless steel needle (Item #34), was prepared by soldering a stranded wire to the side of the needle. As soldering to stainless steel can be difficult, it is recommended to use a flux designed for use with stainless steel such as “Ruby’s Stainless Steel Soldering Flux” (Item #35). The flux is liquid, and it is therefore recommended to wrap the stranded wire tightly around the shaft of the needle prior to applying the flux with a syringe. As the solder joint is in close proximity to the mouth of the animal, we chose to use a lead-free solder (Oatey Silver Safe Flo 53061; Item #36) to minimize risk to the animal. As flux designed for stainless steel soldering is highly acidic, it is recommended to wash the joint with deionized water and to scrub with a brush or Kimwipe to prevent corrosion. The soldered needle was then connected to the spacer via the luer lock connection.

The entire spout assembly was then inserted into a right angle clamp (Item #37) which was then affixed to an inverted 4” optical post (Item #20). This was then inserted into a 0.5” optical post holder (Item #38) which was attached to a mounting base (Item #39) with a screw threaded through the bottom of the post holder. The mounting base allows for easy adjustment of the spout location to the correct position in front of the animal’s mouth (Figure 4A).

### Face/Pupil Camera

An Arducam OV9281 UVC camera (100 fps, 720p; Item #40) was used to video record the face and pupil, as we have done previously (Chintalacheruvu et al., 2026). The Arducam was mounted in a custom 3D printed camera mount (Item #49) printed in PLA. The printed camera mount accepts a setscrew (Item #17) which allows connection to a 0.75” optical post (Item #41). The optical post can then be inserted into a post holder (Item #38) mounted to the aluminum breadboard (Figure 4B). For operation, the Arducam should be connected to the controlling computer via the included USB cable. Additionally, the frame trigger (F) and ground (G) contacts will need to be connected to the Arduino microcontroller. This can be achieved either by directly soldering wires to the contacts, or by soldering header pins (Item #56). The positioning of the camera can be set via the “AMCap” software provided by Arducam, as well as any exposure or gain settings. For acquisition, the camera must be put into the “External Trigger Mode” by clicking “Options”, “Video Capture Filter”, then “Camera Control” and checking the “Low Light Compensation” setting.

### Other Hardware Components

The other hardware components include a wheel lock (Figure 4C), IR light for camera imaging (Figure 4D), and speaker mount for auditory tone presentation (Figure 4E). For the wheel lock, a high torque servomotor (Item #42) was mounted using a custom 3D printed mounting bracket (Item #50). The servomotor was secured in the bracket using the screws included in the packaging, and the small single direction arm was used to serve as the wheel brake. It is advised not to attach the arm until after the servomotor has been connected to the Arduino, as the arm should be attached at a 180° angle (parallel to the surface of the aluminum breadboard) when at rest to ensure successful braking of the wheel. The IR light (Item #43) was mounted similarly to the camera, attached via setscrew (Item #17) to an inverted optical post (Item #44) which was inserted into a post holder (Item #38) on a mounting base (Item #39). The IR light was connected to the computer via USB for power. Finally, the speaker (Item #45) was mounted to the breadboard with a custom 3D Print (Item #51) printed in PLA.

Mounting locations are shown in Figure 4F and a fully constructed setup in Figure 4G.

### Electronic/Hardware Control

An Arduino Uno (Item #54) with custom PCB shield (Item #55, Figure 5) served as the primary hardware interface. The custom PCB was ordered using JLCPCB’s printed circuit board and assembly (PCBA) service (design, gerber, BOM, and placement files are included in extended data). When ordered in batches of 20 using the “economical” PCBA service option and hand soldering the pin headers (Item #56) on the bottom of the PCB, the cost per unit (including shipping) was ∼$10.

**Figure 5.**
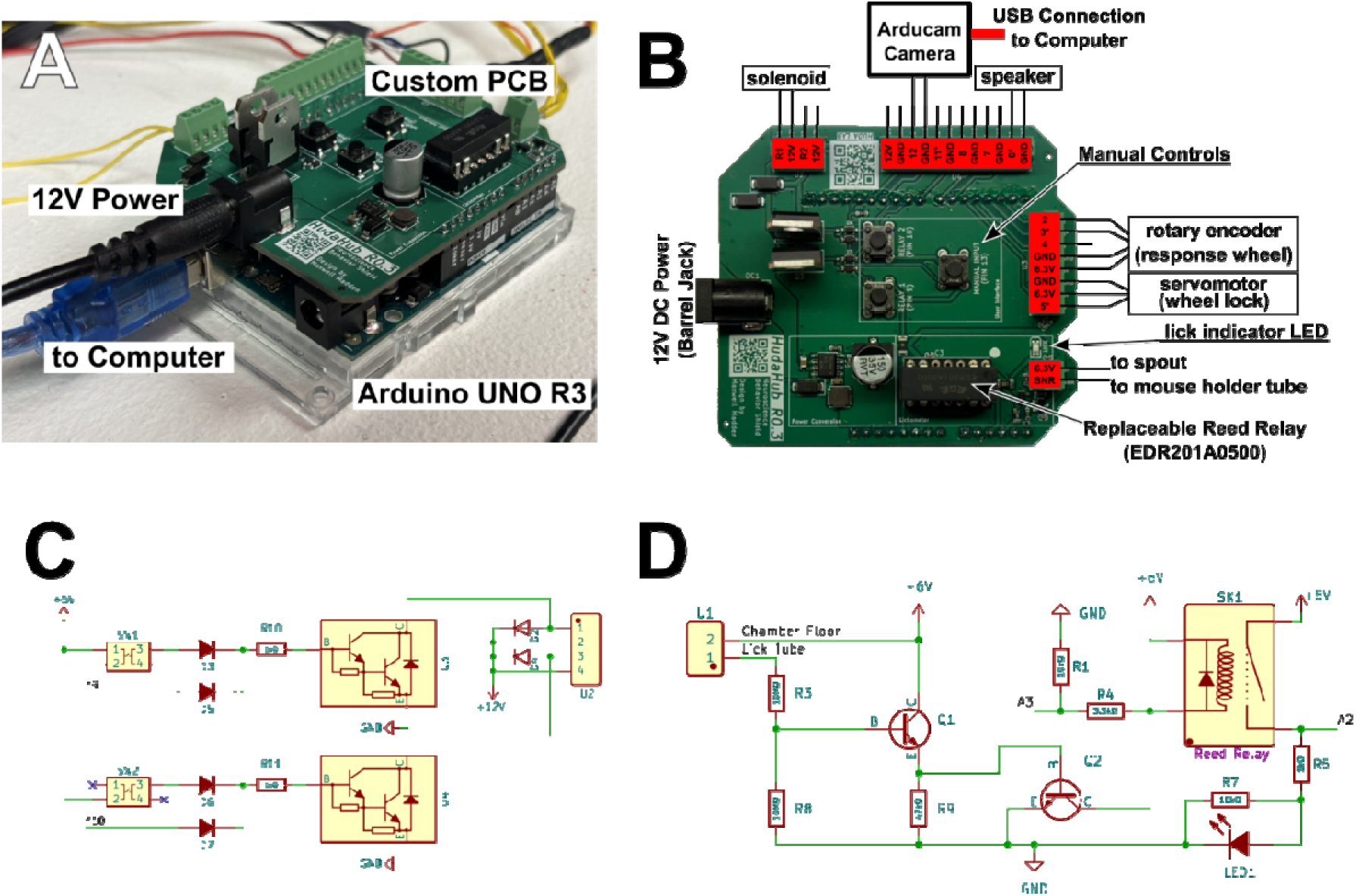
Custom PCB Diagram and Circuits. **A.** Labeled photograph of custom PCB and Arduino Uno installed on a behavior rig. **B.** top down view of custom PCB with wire connections labeled. **C.** Circuit diagram of the 12V relay circuit for solenoid control on the custom PCB. **D.** Circuit diagram of the lickometry circuit present on the custom PCB. Circuit diagrams were prepared in KiCAD. Editable KiCAD files and GERBER files for PCB ordering are available in Extended Data.

The Arduino+Shield was then connected to all active hardware according to the pin diagram in Figure 5B to allow recording and control. An onboard relay circuit (Figure 5C) enables the control of the 12V solenoid valve for reward delivery. To increase potential usefulness and expandability, an additional 12V relay circuit is included to allow the addition of an additional solenoid or other 12V device. 12V devices may be controlled manually by buttons available on the custom Arduino shield, and an additional button is available (connected to Arduino Pin 13) to allow for customizable control of any connected hardware.

Licks are measured via a simple lickometer circuit (Figure 5D) previously described and validated (Slotnick, 2009), operating at 6.3 V. One contact from the circuit (6.3V) is connected to the water spout, while the other (detector) is connected to the aluminum mouse holder tube. At the operating voltage of 6.3 V, current between the spout and detector is limited to a maximum of 0.3 μA, below the reported threshold of behavioral effects in a mouse (Weijnen, 1998). Lick signals recorded from the circuit can either be captured as an analogue signal from Arduino pin A3, or after filtering via a reed relay (EDR201A0500; Item #57) to a digital signal at Arduino pin A2 (Figure 5D). The digital output of the circuit additionally drives a live lick indicator LED, allowing the experimenter to easily ensure there are no loose or disconnected contacts, and that placement of the spout was within reach of the subject’s tongue. As the reed relay is the circuit component with the shortest expected lifetime (estimated 10-100 million operations), the PCB was designed for installation of the reed relay into a mounted socket to allow easy replacement.

The Arduino Shield was powered via a 12V input, and the Arduino was connected to the controlling computer (Item #58). The user interface was a 7” touchscreen (Item #59) mounted to the door of the sound attenuation box via a custom 3D printed mount (Item #52).

### Software Framework

The behavior rig is managed, recorded from, and controlled by a “Lightweight Automated Modular Python framework for Rodent Behavior” (LAMPyR) which was developed in Python 3.12 and is available in extended data with ongoing development and support occurring at GitHub (github.com/hudalaboratory/LAMPyR). LAMPyR was designed to be highly modular, to ease adaptation to alternative tasks, or hardware (Figure 6B). LAMPyR is operated either via a command line interface, or through the “lampyr go” GUI (Figure 6C).

**Figure 6.**
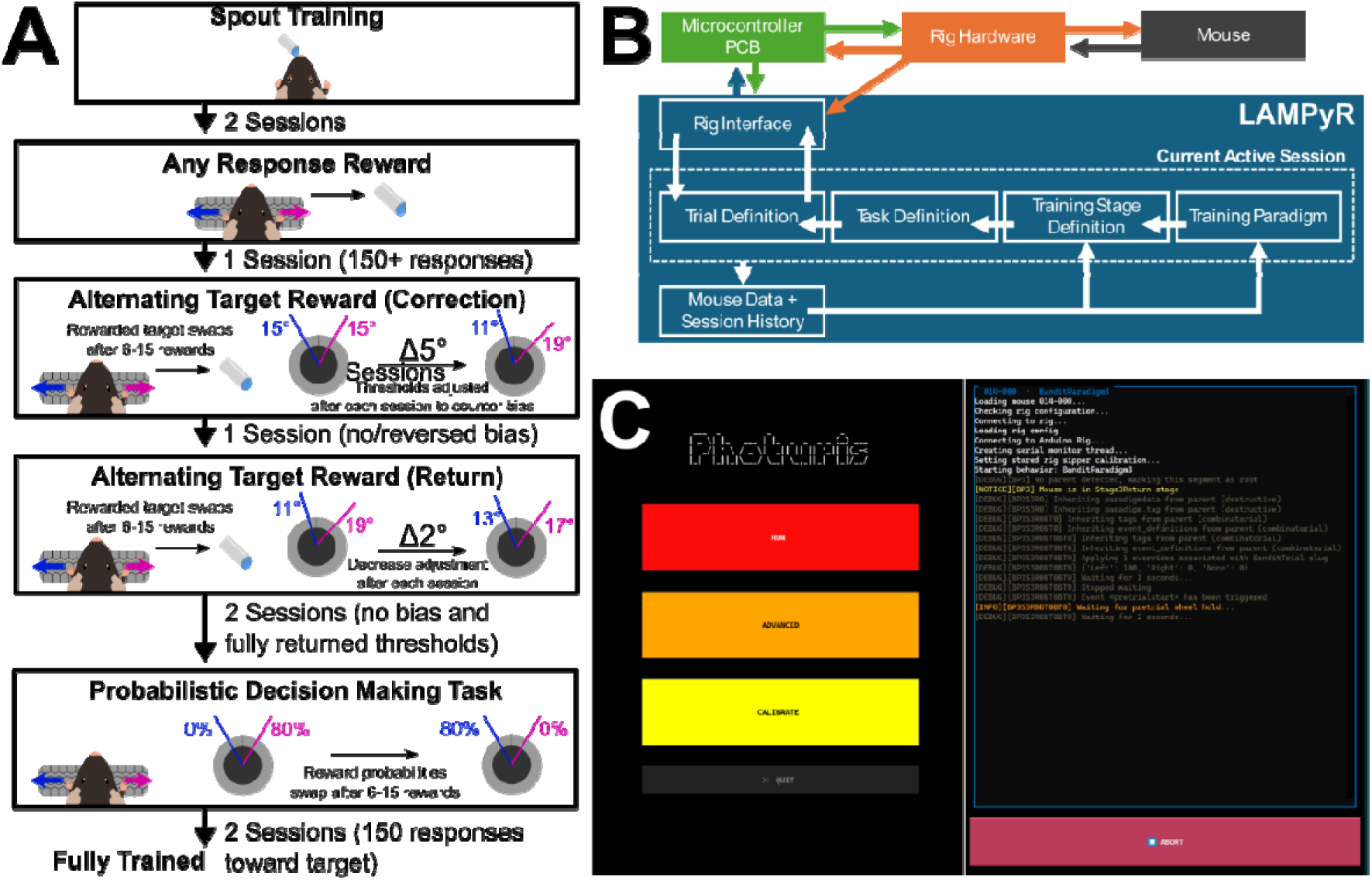
LAMPyR Software and Behavior Training. **A.** Diagram of the implemented training paradigm. **B.** Conceptual diagram of the LAMPyR framework. **C.** Screenshots of the “LAMPyR Go” GUI interface.

For LAMPyR to initiate a specific hardware action via the Arduino, a serial connection (USB 2.0) is utilized. Single letter commands are sent to initiate specific actions defined within the Arduino firmware. The Arduino in turn, reports periodic readouts from various hardware devices in the following format:

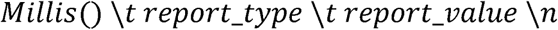

Where “millis()” is the value of the arduino internal clock in milliseconds since beginning of operation. LAMPyR then stores those values in a set of arrays with an additional value of “unix_time” which is the value of the computer clock in the Unix Epoch.

Communication and storage of the Arduino timestamp is performed to maximize accuracy of timing of events. All commands sent to the Arduino are echoed back as the “C” report_type. LAMPyR then exposes numerical data to the behavior code via a “Rig Interface Class” which defines useful methods to extract relevant information (e.g. rig.wheel.movement_since(time) and rig.licks.since(time)).

LAMPyR behavioral protocols are structured hierarchically, with each “Trial” having defined behavior that can be altered by an encompassing “Task” which controls the organization of trials within a single session. Optionally, Tasks can be placed within a Stage, which alters parameters of a Task, and subsequently within a training Paradigm, which defines the conditions for progressing between a sequence of Stages. Definition of new trials, tasks, stages, and paradigms is achieved by constructing python classes that inherit from abstract classes that implement shared functionality, or from other trials, tasks, stages, or paradigms.

Upon completion of a behavioral session, LAMPyR saves all data arrays collected from the Arduino in an .h5 file and stores a record of events and logging in a corresponding json file. These files are registered to the mouse and can be accessed via the in-built loading and analysis helper functions provided in LAMPyR or loaded externally using the h5py library and any json parser. It is recommended to assign the LAMPyR save directory to a shared network drive, to enable easy access to all animal data and easy transfer of animals between rigs when required; however, LAMPyR functions equally well when assigned to a local folder.

### Behavioral Training

Following at least a week of recovery from surgical implantation, water restriction was introduced by progressively decreasing water intake (provided via hydrogel). Final water restriction level was adjusted for each mouse to between 0.6-1.2 ml/day. During water restriction, mice were closely monitored to ensure a body weight of at least 80% initial weight and for behavioral and physiological signs of severe dehydration with logs maintained daily.

Over weekends, animals received free access to 2% Citric Acid water to maintain water restriction as previously described (Urai et al., 2021). After the establishment of water restriction, mice were habituated to the behavior rig over two consecutive days in which their entire water allotment was provided through periodic water rewards (5 μl) via the rig water spout (Spout Training, Figure 6A).

After spout training, the full task was trained via a progressive shaping paradigm. Across all stages of task training, to initiate a trial, the animal must hold the wheel still (less than 5° total movement) for two seconds. After a successful trial initiation a 100 millisecond cue (4000 Hz) is presented and the animal is given 3s to make a response with the wheel (to the left or right). If a response is registered, the wheel is locked via engagement of the wheel brake and a 5 μL water reward may or may not be presented (i.e., probabilistic reward delivery). After 2.5 seconds, if the wheel is locked, it unlocks. After an additional 1 second, the animal may now initiate a new trial (Figure 7A).

**Figure 7.**
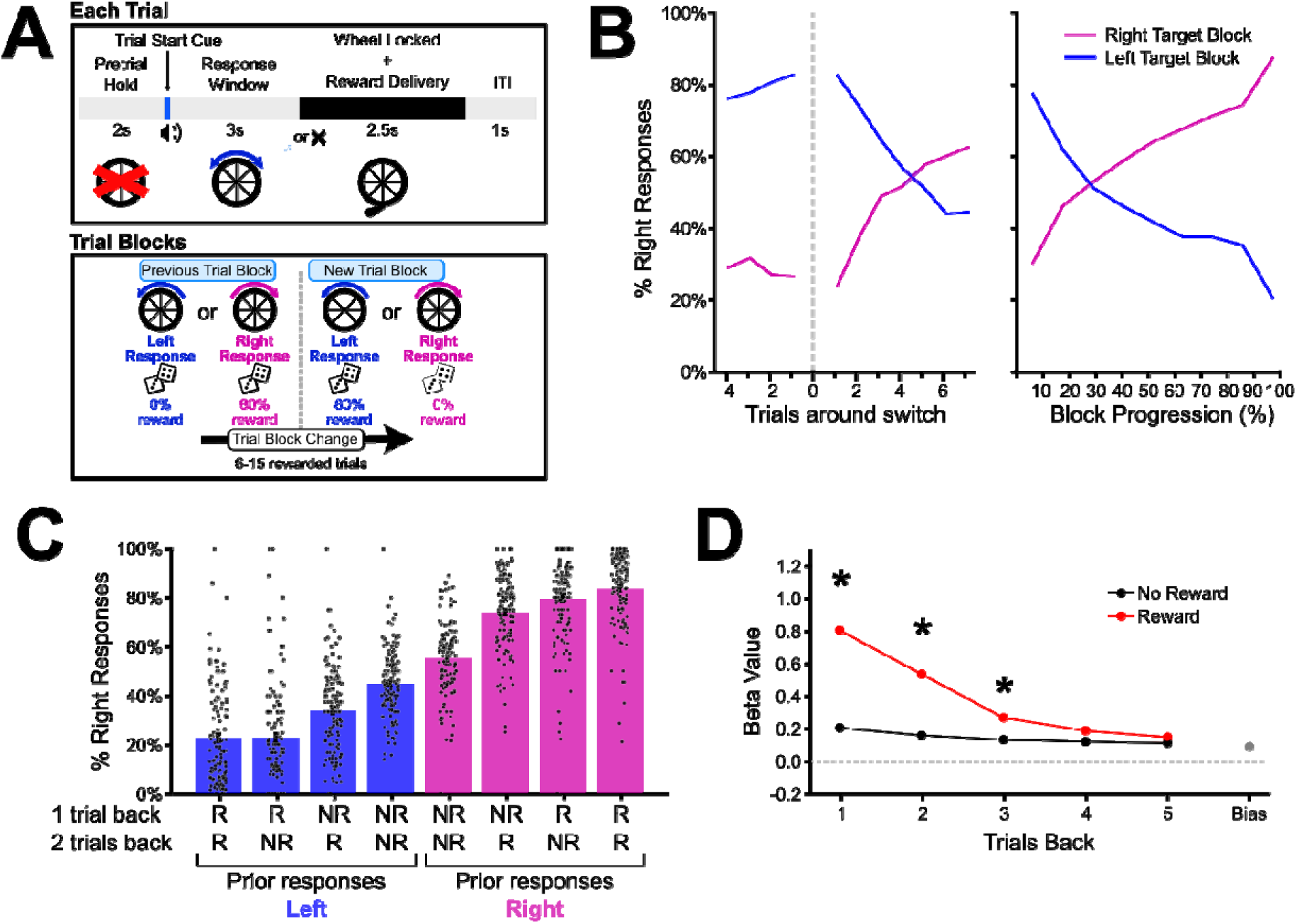
Trained Performance in the probabilistic rapid-reversal task. **A.** Diagram depicting the task paradigm. Mice were required to initiate trials by holding the wheel still for 2s. An auditory cue signaled trial start, after which the animal had three seconds to make a response (wheel movement of 15° in either direction). After response, the wheel was locked and a reward delivered based on the current probability for response reward (top). Reward probabilities for each response were switched after 6-15 rewarded trials (bottom) **B.** Choices within left and right target reward blocks centered around the block transition (left). Trials within blocks were grouped into 10 bins. Error bars show SEM across animals (N = 11 animals, 5-21 sessions per animal). **C.** Mean percent of rightward responses based on the actions and outcomes of the previous two trials. Only trials in which the previous two responses were the same were selected, for simplicity of visualization. (N = 125 sessions). **D.** Regression coefficients (mean ± SEM) from a logistic regression predicting current trial choice from the choice/outcome history of the previous 5 trials. The regression coefficients for rewarded trials were significantly different from those of unrewarded trials for the first (*p* = .002), second (p = .00008), and third (p = .016) trials back, but not the 4^th^ (p = .21) or fifth (p = .40) trials (paired t-test, N = 11 animals, Holm-Bonferroni correction for multiple comparisons).

In the first stage of training (“Any Response Reward”), animals received rewards when they turned the wheel to either the left or right by 15°. When animals have successfully completed a session with more than 150 wheel responses, they transition to the “Alternating Target Reward” stage. Within this stage, animals receive a water reward for responding to an assigned target side, which is swapped every 6-15 rewards. Initially animals exhibit a strong bias towards one side; to train responses to the other side, after each session, the degree thresholds for responses are shifted by 5° opposing the direction of bias. Once an animal has completed a session in which there was no strong bias towards one side, or bias was observed in the opposite direction, the animal transitions to the second part of the training stage, in which the thresholds are shifted by 2° after every session back to their initial 15° values, unless the animal exhibits a strong bias on that session. Finally, after the thresholds have returned to their 15° values and the animal has completed 2 consecutive sessions in which no bias was detected, they are ready to progress to the full version of the task, in which the target side is rewarded only 80% of the time. After two sessions in which the animal has successfully made responses toward the target 150 times in a session, the animal is fully trained and ready for experimental manipulations (Figure 6A).

### Logistic Regression Analysis

A logistic regression was performed using the choice and outcomes of the previous 5 trials as predictors of current trial choice (Figure 7D). The regression model can be expressed as

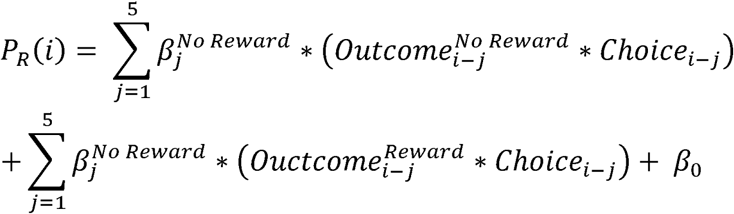

Where for each trial (j) backwards from the current trial (i), choice during that trial (i-j) was 1 if towards the right, and -1 if towards the left and each outcome (reward or no reward) is 1 if it occurred and 0 if it did not. A bias term β_0_ was also fitted.

### Data and Software Availability

All necessary Python and Arduino code are included in the Extended Data and are actively developed and maintained in our Github (github.com/hudalaboratory/LAMPyR). Associated hardware documentation is available both in Extended Data and on Github. All code and documentation on the GitHub have been shared under a permissive open source license (MIT).

## Results

The dual objective of this work was to present a versatile, cost-efficient system for head-fixed mouse behavior and to demonstrate its effectiveness in the context of a two-choice probabilistic rapid-reversal task (Figure 7A).

### Platform Adaptation and Scalability

A primary design consideration for this system was per rig cost, as for tasks requiring large amounts of training, parallelization of training across multiple rigs is paramount. The total cost of hardware components for a single rig was approximately $1,400 USD. However, the cost of per rig may be further reduced, and conversely, alternative components may be utilized at higher cost to expand the capabilities of the system.

A large portion of the per-rig cost was the Dell minicomputers (Item #58) used to control each rig, however cheaper Linux based single board computers such as the Raspberry Pi 5 (Item #60) are available and confirmed to be compatible with our provided software, bringing the total cost of the rig closer to ∼$970.

Additionally, we chose to construct our own sound attenuation chambers as this approach provided the minimum possible cost per chamber but increased the amount of labor required to construct each rig. A suggested alternative would be the purchase of fully assembled “Wall Bridge” kitchen cabinets (Home Depot, #SKU 1002751910) at the cost of some diminishment in sound dampening (due to the lower density material typically used in home cabinets). If additional sound dampening is required, the chamber can be lined with mass loaded vinyl as previously described (Ozgur et al., 2023).

We chose to utilize an Arducam camera, which is sufficient to provide detection of face movements and pupil diameter at 20Hz. Arducam has several advantages over more widely used industrial cameras; as a UVC camera, the Arducam can be accessed from Windows, Linux, and macOS computers without the installation of custom drivers or software. Additionally, the Arducam OV9281 UVC camera does not contain the IR filter present in some cameras, relieving the necessity of modifying the camera or separate purchasing of a non-IR filtering lens. For applications requiring higher fidelity or faster acquisition than is capable with an Arducam camera, we recommend the “Blackfly” camera line available from Teledyne FLIR.

### Automated training with LAMPyR

LAMPyR automatically progresses a mouse through a multistage training paradigm (Figure 6A), based on predefined performance criteria. This substantially reduces experimenter labor and enables the simultaneous training of a large numbers of mice at different stages of learning. When individualized adjustments to training are required, an experimenter can adjust mouse training stage via the command line interface or apply mouse specific “overrides” for specific task parameters by editing the values in a mouse’s associated JSON configuration file. Additional specific instructions of how to tweak mouse training performance are contained in the docs/training_management.md document on the LAMPyR Github. The modular architecture of LAMPyR (Figure 6B) allows incorporation of additional hardware, tasks, and training paradigms without modification of the core framework. The accompanying touchscreen GUI (Figure 6C) enables operation by multiple experimenters with lower onboarding requirements and supports high-throughput behavioral training (10+ rigs, 40+ animals per day) with reduced opportunities for experimenter error.

### Solutions to common hardware issues

A few issues commonly appeared while building the system that were easily addressable but required some time to identify. LAMPyR uses the serial device name from the OS to identify the Arduino controller. Thus, if Arduino drivers are not installed on the computer, or if they were installed incorrectly, LAMPyR may fail to identify rig hardware. This is easily rectified by reinstallation of Arduino drivers via the Arduino IDE. Another issue may arise if attempting to add additional PWM signal devices for simultaneous operation. The Arduino Uno has only three hardware timers, the first of which is used for the system clock. The other two timers are currently used to generate a PWM signal for control of the servomotor wheel lock and to generate audio tones. As such, if an experimenter is planning on implementing PWM control of an additional external device such as a laser or LED for ramped optical stimulation, it is recommended to either restrict PWM control signals to non-tone periods, use an additional Arduino as a stimulus generator, or to utilize a more sophisticated microcontroller board such as a Teensy or Raspberry Pi Pico. TTL based digital control of additional devices is not affected. Additional hardware and software support is available through the Github issues tracker and email.

### Behavioral adaptation in a rapid reversal learning task

A cohort of 11 animals were utilized to demonstrate the behavioral task and test rig hardware/software. Training was distributed across four experimenters, with occasional adjustments to training by the lead experimenter, demonstrating that the standardized workflow and touchscreen interface enabled consistent operation by multiple users. Once fully trained, mice performed multiple sessions in the full version of the task (Figure 7A), performing hundreds of trials (average: 366, range: 236-568) per 1hr session with tens of block switches per session (average 17.8, range: 11-24).

Animals were able to successfully adapt their responses over the course of each block, with choice probability gradually shifting toward the newly rewarded response over successive trials (Figure 7B). To characterize the history-based decision-making performance of the mice, we visualized the chance of a specific response based on outcome and choice history for the preceding two trials (Figure 7C). We additionally performed a logistic regression predicting choice in the current trial based on the choices and outcomes of the previous five trials. We observed a significant effect of reward outcome in the last three trials (Figure 7D; paired t-test, N=11 animals, Holm-Bonferroni correction for multiple comparisons), indicating that animals integrated reward history over multiple recent experiences rather than relying solely on the immediately preceding outcome.

## Discussion

Here we presented an open-source behavior rig designed for the implementation of rodent decision-making tasks and training. The system was designed to emphasize automation and low-cost implementation at scale to enable high-throughput training. We demonstrate that mice can be successfully trained in a probabilistic rapid-reversal decision-making task where animals must actively monitor recent action outcomes to maintain optimal behavior. The system is easily integrable with commercial and custom neuronal recording hardware, as both behavioral and arbitrary TTL signals can be easily output from the Arduino controller to enable precise temporal alignment. Additionally, the software framework, LAMPyR, developed in python, is easily modified to support additional hardware, tasks and training paradigms.

This design will be most useful to experimenters with some experience with Arduino and Python. We don’t believe this is a significant barrier to use, as proficiency in Python has become an increasingly common skill in the field and both Arduino and Python have extensive learning resources. The advent of large language model coding assistants has further expanded the ability of experimenters to interact with code, though sufficient human oversight to understand and test output is necessary. Nevertheless, systems such as Bonsai (Lopes et al., 2015), which utilize a visual programming interface, remain useful alternatives.

The complete per-unit cost of our behavioral rig, including computer, microcontroller, sound attenuation box, and all components, is $1,400, with potential for further reduction to <$1,000 per rig: lower than any other comparable system we are aware of. Additionally, all custom components use widely and commercially available custom manufacture technologies and services. As the behavior rig has been designed with modularity in mind, many aspects of the rig can be adapted to other tasks. The Arduino shield can be used with alternative Arduino firmware, or with other Arduino Uno format development boards. The LAMPyR software is fully adaptable to other hardware and tasks, and many portions of its code (released under a permissive open-source license) are fully separable for use in custom setups. Furthermore, the automation of training progression decreases the daily labor load on an experimenter and enables training to be run by a team with reduced onboarding and experimental errors.

Our presented approach is not unique, and several open-source or commercial behavior platforms are already available. Commercial open-source systems such as Bpod (Sanworks) and pyControl (Akam et al., 2022) offer suites of hardware and firmware to offer similar or greater functionality, though at a significantly higher price point than our system. Bpod is a modular MATLAB and Arduino based system, which offers increased performance though at significantly increased costs (a single Bpod finite state machine costs more than our entire rig). pyControl is a similar modular system based around a Micropython microcontroller which also offers tested compatibility with the pyPhotometry open-source photometry system. The pyControl breakout board remains significantly more expensive than our solution, costing more than 5x the Arduino + custom shield. Notably, the the International Brain Laboratory Consortium has created a well-documented and robust decision-making apparatus (Laboratory et al., 2021) to execute a similar, visual stimulus associated decision-making task utilizing Bpod and Bonsai. These platforms may be better suited to many experimenters and labs; however, our approach represents a significant lowering of the financial barrier to entry to head-fixed behavior, additionally, due to the modularity of our system, much of the hardware and software solutions presented here can be easily adapted for use in established commercial open-source systems or custom-built setups.

As the development of more complex and focused tasks proceeds, it is vital that labs continue to openly share detailed specifications of their behavioral setups to maximize the impact and replicability of scientific discoveries, as well as to reduce the labor and financial costs of constructing behavior rigs. It is our hope that this publication will not only provide a blueprint for those wishing to run similar decision-making tasks, but also provide useful practical details regarding rig construction that will prove useful to experimenters developing new rig hardware and tasks.

## Acknowledgements

We thank Gabe Villegas for assistance in 3D design, modeling, and printing. Financial support for this work was provided by National Institute on Drug Abuse fellowship 5T32DA055569 (Rutgers Training in Addiction Research Program) awarded to M.B.M. and National Institute on Alcohol Abuse and Alcoholism grant 5R01AA030594 awarded to R.H.

