## Extended Data for "A Low-Cost, Modular Hardware and Software Platform for Head-Fixed Mouse Decision-Making Tasks": app_architecture_and_segments.md

# LAMPyR app architecture and segment inheritance

LAMPyR is a Python framework for running behavioral rig sessions. The central idea is that an experiment is represented as a tree of **segments**. Segments inherit context from their parents, record data while they run, and dump their state into a session at the end.

This document describes the application structure and how to create new tasks by inheriting from segment objects.

## High-level architecture

Core package areas:

- `lampyr/main.py` — top-level `Lampyr` orchestrator.
- `lampyr/config.py` — JSON-backed user configuration.
- `lampyr/primatives.py` — `Mouse`, `Session`, and unique ID primitives.
- `lampyr/managers/` — data, mouse, rig, plugin, and notification managers.
- `lampyr/rigs/` — hardware rig abstractions and concrete Bandit rig components.
- `lampyr/segments/` — segment base classes for trials, tasks, stages, and paradigms.
- `lampyr/behaviors/` — concrete behavior implementations.
- `lampyr/interfaces/click_cli/` — Click CLI.
- `lampyr/interfaces/textual_tui/` — Textual GUI/TUI.
- `lampyr/analysis/` — session, mouse, colony, records, traces, and dataset helpers.

## `Lampyr`: the application object

`Lampyr` is the top-level runtime object. Constructing it creates:

- `config`: a `Config` loaded from `%LOCALAPPDATA%/lampyr/config.json`.
- `datamanager`: a `DataHandler` for mouse/session save/load and backups.
- `rigmanager`: a `RigManager` for connecting, disconnecting, and calibrating the physical rig.
- `mousemanager`: a `MouseManager` for creating, loading, saving, and retiring mice.
- `notificationmanager`: a `NotificationManager` for Pushover notifications.
- `behaviors`: a registry mapping segment class names to imported `Segment` subclasses.
- `paradigms`: an initially empty dict intended for paradigm classes/user population.

The important runtime method is:

```python
lampyr.run(segment_name: str, **kwargs)
```

It:

1. Validates that `segment_name` exists in the behavior registry.
2. Requires an active rig.
3. Creates a new `Session`, injecting the active mouse ID if available.
4. Instantiates the selected segment with `lampyr=self` and `_verbose=True`.
5. Calls `segment.run()`.
6. Prints the session summary.
7. Evaluates stop conditions.
8. Sends a completion notification.

`Lampyr.close()` disconnects the rig, saves the session, saves the active mouse, and updates local mouse backups.

## Data model

### Mouse

`Mouse` represents one experimental subject. Key fields:

- `mouseid`
- `retired`
- `mouse_behav_param_overrides`
- `paradigm`
- `properties`
- `history`

`properties` is where paradigm-specific persistent state lives. A paradigm with slug `BanditParadigm`, for example, stores current stage and shaping data under `mouse.properties['BanditParadigm']`.

`mouse_behav_param_overrides` supports per-mouse parameter overrides. Keys may be:

- `'all'`
- a segment `slug`
- one of a segment's inherited `tags`

Values are dictionaries of `{attribute_name: override_value}`. Overrides only apply to attributes that already exist on the segment.

### Session

`Session` stores all counters, limits, metadata, segment records, events, rig data, and extended data references for one run.

Important counters include:

- `merit`, `demerit`
- `trial`
- `rewards`, `reward_amount`
- `abstention`
- `participation`
- `serial_abstention`
- `duration`

Each counter has optional `_min` and `_limit` companions. `Session.evaluatestopconditions()` returns a list of stop reasons when limits are reached. `Session.lock()` finalizes the session and makes it immutable.

## Segment hierarchy

The segment classes are in `lampyr/segments/`.

```text
Segment
├── BehaviorSegment
│   ├── Task
│   └── Trial
└── ParadigmSegment
    ├── Stage
    └── Paradigm
```

### `Segment`

`Segment` is the abstract base for all executable experiment units.

Subclasses must implement:

```python
def execute(self):
    ...
```

`Segment.run()` wraps `execute()` with common behavior:

1. Prevents running the same segment twice.
2. Records `starttime`.
3. Calls `execute()`.
4. Records `endtime`.
5. Freezes the segment.
6. Calls `dump()` even if execution exits by exception.

`Segment.dump()` serializes public segment state into `session.segments`. If the segment is the root segment (`rank == 0`), it also stops rig acquisition, dumps rig data into the session, and locks the session.

### Parent inheritance

Segments form a tree. Child segments should be instantiated with `parent=self`:

```python
trial = MyTrial(parent=self)
trial.run()
```

A child automatically inherits common runtime context from its parent:

- `lampyr`
- `rig`
- `mouse`
- `session`
- `_output_func`
- `_verbose`

The child rank becomes `parent.rank + 1`. When a non-root child dumps, its `uniqueid` is appended to `parent.subdata`, giving the session a navigable tree.

If a segment has no parent but has a `lampyr` object, it inherits context directly from `lampyr`.

### `BehaviorSegment`

`BehaviorSegment` adds behavior-specific support:

- inherited/combinable `tags`
- `reports`
- `event_definitions`
- `event_records`
- registered event callbacks
- `stop_reasons`
- mouse parameter overrides
- counter logging helpers

Common methods:

```python
self.register_event(name, callback=..., description=...)
self.trigger_event(name, *args, **kwargs)
self.create_report(key, value)
self.log_merit()
self.log_demerit()
self.log_abstention()
self.log_participation()
self.log_reward(amount)
self.log_trial()
self.finish()
self.notify(message)
```

### `Task`

`Task` is for trialless behaviors or behaviors that repeatedly create trials.

Subclasses implement:

```python
def setup(self):
    ...

def loop(self):
    ...
```

`Task.execute()` calls `setup()` once and then repeatedly calls `loop()` until session stop conditions are met.

### `Trial`

`Trial` is for one discrete trial.

Subclasses implement:

```python
def setup(self):
    ...

def perform(self):
    ...
```

`Trial.execute()` calls `setup()`, then `perform()`, then logs one trial and marks itself finished.

Useful helpers:

```python
self.wait(seconds)
self.waitfor(condition, fallback_value=None, timeout=None, poll_interval=0.05)
```

`waitfor()` polls a condition until it returns a truthy value or a timeout occurs.

## Creating a new task

The usual pattern is:

1. Create a `Trial` subclass for one unit of behavior.
2. Create a `Task` subclass that runs that trial repeatedly.
3. Import the module before the `Lampyr` registry is built.
4. Run the task by class name from the CLI or GUI.

### Example: one trial plus one task

```python
# lampyr/behaviors/my_behavior.py

from dataclasses import dataclass
from lampyr.segments import Trial, Task, BehaviorSegment


def reward_callback(segment: BehaviorSegment):
    segment.rig.reward.give()
    segment.log_reward(0.005)


@dataclass
class MyChoiceTrial(Trial):
    slug: str = "MyChoiceTrial"
    response_window_s: float = 3.0
    response_threshold_deg: float = 15.0

    def setup(self):
        self.register_event(
            "reward",
            callback=reward_callback,
            description="Deliver one calibrated water reward",
        )
        self.register_event(
            "response",
            description="Wheel movement crossed response threshold",
        )

    def perform(self):
        self.rig.wheel.home()
        response = self.waitfor(
            condition=lambda: abs(self.rig.wheel.angle()) > self.response_threshold_deg,
            fallback_value=False,
            timeout=self.response_window_s,
            poll_interval=0.01,
        )

        if response:
            self.trigger_event("response")
            self.trigger_event("reward")
            self.create_report("responded", True)
            self.log_merit()
        else:
            self.create_report("responded", False)
            self.log_abstention()


@dataclass
class MyChoiceTask(Task):
    slug: str = "MyChoice"

    def setup(self):
        self.tags.append("choice")

    def loop(self):
        trial = MyChoiceTrial(parent=self)
        trial.run()
```

Run it after it is imported:

```bat
lampyr mouse run 014-003 MyChoiceTask --duration_limit 30 --reward_limit 100
```

### Prefer `field(default_factory=...)` for mutable defaults

Because segments are dataclasses, mutable defaults must use `field(default_factory=...)`.

Good:

```python
from dataclasses import dataclass, field
from lampyr.segments import Trial

@dataclass
class MyTrial(Trial):
    reward_probs: dict = field(default_factory=lambda: {"Left": 80, "Right": 20})
```

Avoid:

```python
reward_probs: dict = {"Left": 80, "Right": 20}  # do not do this
```

### Running one trial per loop with `TrialToTask`

For simple behaviors, `TrialToTask` can generate a task wrapper automatically.

```python
from dataclasses import dataclass
from lampyr.segments import Trial, TrialToTask

@TrialToTask
@dataclass
class LaserPulseTrial(Trial):
    slug: str = "LaserPulse"
    pulse_s: float = 2.0

    def setup(self):
        pass

    def perform(self):
        self.rig.laser.begin()
        self.wait(self.pulse_s)
        self.rig.laser.stop()
```

This creates a task class named `LaserPulseTrialTask` that runs `LaserPulseTrial` once per task loop.

## Inheriting from existing task/trial classes

You can create new behavior variants by subclassing an existing concrete class and overriding dataclass fields or selected methods.

Example:

```python
from dataclasses import dataclass
from lampyr.behaviors.bandit import BanditTask

@dataclass
class HighRewardLeftBandit(BanditTask):
    slug: str = "HighRewardLeft"
    target_mode: str = "Left"
    reward_prob_target: int = 90
    reward_prob_offtarget: int = 10
    taskblocks_enabled: bool = False
```

Because `HighRewardLeftBandit` inherits from `BanditTask`, it keeps the Bandit task loop but changes the parameters. It is discoverable by class name once its module is imported.

For more substantial changes, override `setup()` or `loop()` and call `super()` where appropriate:

```python
@dataclass
class MyBanditVariant(BanditTask):
    slug: str = "MyBanditVariant"

    def setup(self):
        super().setup()
        self.log_notice("Using custom setup")

    def loop(self):
        # custom pre-trial logic
        super().loop()
        # custom post-trial logic
```

## Event and report design

Use events for timestamped occurrences and reports for trial/segment outcomes.

Events:

```python
self.register_event("cue", description="Cue onset")
t_cue = self.trigger_event("cue")
```

Events are recorded in the segment and appended to `session.eventlist`.

Reports:

```python
self.create_report("response", "Left")
self.create_report("rewarded", True)
```

Reports are stored inside the dumped segment data and are useful for analysis and stage shaping.

## Session counter design

Use counter helpers so stop conditions and summaries work correctly:

- `log_trial()` increments `session.trial`.
- `log_reward(amount)` increments reward count and reward amount.
- `log_merit()` increments merit and participation.
- `log_demerit()` increments demerit and participation.
- `log_abstention()` increments abstention and serial abstention.
- `log_participation()` increments participation and resets serial abstention.

Do not update counters directly unless you have a specific reason.

## Creating a paradigm and stages

Use `Paradigm` when a mouse should progress through stages across sessions.

A `Stage` implements:

```python
def define_sessionparams(self):
    ...

def define_task(self, stage_data):
    ...

def session_summary(self):
    ...

def define_shaping(self, stage_data):
    ...
```

A `Paradigm` supplies `stagelist` and progression logic:

```python
from dataclasses import dataclass
from lampyr.segments import Stage, Paradigm

@dataclass
class StageA(Stage):
    slug: str = "StageA"

    def define_sessionparams(self):
        self.set_sessionparam("duration_limit", 30)
        self.set_sessionparam("reward_limit", 100)

    def define_task(self, stage_data):
        task = MyChoiceTask(parent=self)
        task.run()

    def session_summary(self):
        self.log_notice(str(self.session))

    def define_shaping(self, stage_data):
        stage_data["last_merit"] = self.session.merit


@dataclass
class MyParadigm(Paradigm):
    slug: str = "MyParadigm"
    stagelist: tuple = (StageA,)

    def define_progression(self, current_stage, stage_data):
        pass
```

Assign and run it:

```bat
lampyr mouse paradigm 014-003 MyParadigm --stage StageA
lampyr mouse run 014-003
```

Important paradigm details:

- `Paradigm` must run as the root segment.
- It requires a `Lampyr` instance and an active mouse.
- It stores persistent data in `mouse.properties[paradigm.slug]`.
- It selects the current stage from that persistent data.
- Stage `set_sessionparam()` respects explicit user overrides by warning instead of overwriting existing session values.

## Making new behaviors discoverable

`Lampyr.__init__()` builds its registry from already-imported subclasses:

```python
self.behaviors = {c.__name__: c for c in Segment.get_children()}
```

Therefore, defining a class is not enough; the module containing it must be imported before the `Lampyr` object is created.

Current CLI behavior imports include:

```python
from lampyr.behaviors import bandit
from lampyr.behaviors import test
```

Common options for adding a new behavior module:

1. Add the module under `lampyr/behaviors/`.
2. Import it from `lampyr/interfaces/click_cli/app.py`, or from `lampyr/behaviors/__init__.py` and ensure that package import happens before registry creation.
3. Run `lampyr list` to verify the class appears.

Example import line:

```python
from lampyr.behaviors import my_behavior
```

If the class does not appear in `lampyr list`, it has not been imported early enough.

## Hardware access from segments

Segments that inherit rig context can call rig components. For the Bandit rig, commonly used components include:

- `self.rig.wheel.home()`
- `self.rig.wheel.angle()`
- `self.rig.wheel.movement_total_since(t)`
- `self.rig.licks.since(t)`
- `self.rig.play.begintrialtone()`
- `self.rig.play.responsetone()`
- `self.rig.play.punishtone()`
- `self.rig.reward.give()`
- `self.rig.reward.setsize(size)`
- `self.rig.wheellock.lock()`
- `self.rig.wheellock.unlock()`
- `self.rig.wheellock.to_angle(angle)`
- `self.rig.laser.begin()`
- `self.rig.laser.stop()`
- `self.rig.laser.rampdown(ramp_ms)`

Always design segment code so the root segment can finish and dump cleanly. Avoid swallowing `KeyboardInterrupt` unless you re-raise it after cleanup.

## Persistence and analysis implications

Every segment dump includes:

- identifiers (`name`, `slug`, `uniqueid`)
- timing (`starttime`, `endtime`)
- tree structure (`rank`, `subdata`, `parent` representation)
- records/logs
- behavior reports/events
- inherited tags and event definitions
- `segment_type`, a list of parent segment class names

The `Session.search()` helper can later retrieve segments by type, slug, report checks, or custom filters.

Good task design makes analysis easier:

- Give classes and slugs stable names.
- Store outcome variables in `reports`.
- Use events for timestamps.
- Use counter helpers for global session metrics.
- Keep mutable task state in clearly named dataclass fields.
