## Extended Data for "A Low-Cost, Modular Hardware and Software Platform for Head-Fixed Mouse Decision-Making Tasks": cli_commands.md

# LAMPyR CLI command reference

This document describes the Click command-line interface defined in `lampyr/interfaces/click_cli/app.py`. The installed console script is `lampyr`; on Windows, commands are often shown as `LAMPyR` and are equivalent in normal use.

Most commands create a `Lampyr` application object, load configuration from `%LOCALAPPDATA%/lampyr/config.json`, and close cleanly when the command exits. Commands that run behavior connect to the configured rig, create a session, run a segment by class name, save session/mouse data, and send completion notifications when configured.

## Global form

```bat
lampyr [OPTIONS] COMMAND [ARGS]...
```

Top-level commands:

- `lampyr info`
- `lampyr list`
- `lampyr reset`
- `lampyr developer`
- `lampyr go`
- `lampyr run BEHAVIOR [SESSION OPTIONS]`
- `lampyr rig ...`
- `lampyr mouse ...`
- `lampyr user ...`

Use Click's built-in help for command-specific help:

```bat
lampyr --help
lampyr mouse run --help
lampyr user ping --help
```

## Session stop-condition options

The direct run commands (`lampyr run` and `lampyr mouse run`) accept the same set of optional session parameters. They are forwarded to `lampyr.primatives.Session` and evaluated during task execution.

| Option | Short | Type | Meaning |
|---|---:|---|---|
| `--merit_limit` | `-ml` | int | Stop when merit count is at least this value. |
| `--merit_min` | `-mm` | int | Do not stop for non-reward limits until merit count reaches this value. |
| `--demerit_limit` | `-dml` | int | Stop when demerit count is at least this value. |
| `--demerit_min` | `-dmm` | int | Do not stop for non-reward limits until demerit count reaches this value. |
| `--duration_limit` | `-dl` | int | Stop after this many minutes. |
| `--duration_min` | `-dm` | int | Do not stop for non-reward limits until this many minutes have elapsed. |
| `--trial_limit` | `-tl` | int | Stop when trial count is at least this value. |
| `--trial_min` | `-tm` | int | Do not stop for non-reward limits until trial count reaches this value. |
| `--reward_limit` | `-rl` | int | Stop when reward count is at least this value. Reward limits are privileged and are checked before minimum gates. |
| `--reward_min` | `-rm` | int | Do not stop for non-reward limits until reward count reaches this value. |
| `--abstention_limit` | `-al` | int | Stop when abstention count is at least this value. |
| `--abstention_min` | `-am` | int | Do not stop for non-reward limits until abstention count reaches this value. |
| `--participation_limit` | `-pl` | int | Stop when participation count is at least this value. |
| `--participation_min` | `-pm` | int | Do not stop for non-reward limits until participation count reaches this value. |
| `--serial_abstention_limit` | `-sal` | int | Stop when consecutive abstention count is at least this value. |
| `--serial_abstention_min` | `-sam` | int | Do not stop for non-reward limits until consecutive abstention count reaches this value. |

Notes:

- Limits with value `None` are ignored.
- Reward count and reward amount limits are intentionally privileged in the session evaluator to avoid accidental over-delivery.
- Minimums gate later limit checks: for example, `--duration_min 10 --trial_limit 20` prevents the trial limit from ending the session until 10 minutes have elapsed.

## Top-level commands

### `lampyr info`

Prints the LAMPyR version banner and the full loaded configuration.

```bat
lampyr info
```

### `lampyr list`

Lists all currently imported behavior segment subclasses grouped as paradigms, stages, tasks, and trials.

```bat
lampyr list
```

The CLI imports `lampyr.behaviors.bandit` and `lampyr.behaviors.test`, so classes from those modules are normally present. New behavior modules must be imported before the `Lampyr` object builds its subclass registry.

### `lampyr reset`

Resets configuration to defaults after an interactive confirmation prompt.

```bat
lampyr reset
```

You must type exactly `YES` when prompted. Defaults include the shared mice directory and rig defaults from `lampyr/config.py`.

### `lampyr developer`

Developer shortcut guarded by a password prompt.

```bat
lampyr developer
```

If the prompt receives the expected password, it sets:

- `rig.calibrated` to a far-future value
- `rig.name` to `Photuris`
- `rig.configured` to `True`

This is intended for development/testing, not normal rig operation.

### `lampyr go`

Launches the Textual terminal UI.

```bat
lampyr go
lampyr go --notouchoverlay
```

Options:

- `--notouchoverlay`: disables the Windows touchscreen-to-mouse overlay bridge.

On Windows, `go` toggles the terminal fullscreen with F11, disables Quick Edit mode, enables mouse input, and starts a transparent touch bridge unless disabled.

### `lampyr run BEHAVIOR`

Runs a behavior by class name without explicitly loading a mouse. The default active mouse is `UNKNOWN_MOUSE` unless code has loaded a mouse before the run.

```bat
lampyr run RewardedHabituationTask -dl 60 -rl 200
lampyr run BanditTask --trial_limit 100 --duration_limit 45
```

Before running, LAMPyR checks that:

1. `rig.configured` is truthy.
2. `rig.calibrated` is newer than five days.
3. The rig can connect.

If checks pass, `Lampyr.run()` creates a session and instantiates the named behavior from the segment subclass registry.

## Rig commands

### `lampyr rig info`

Prints the `rig` section of the current config.

```bat
lampyr rig info
```

### `lampyr rig configure`

Prompts for a rig name and marks the rig configured.

```bat
lampyr rig configure
```

Current stored values updated:

- `rig.name`
- `rig.configured = 1`

### `lampyr rig calibrate`

Runs the interactive sipper calibration routine.

```bat
lampyr rig calibrate
```

Workflow:

1. Connects to the rig if it is not already connected.
2. Tests three dispenser sizes around the current estimate.
3. Prompts for scale weights before/after reward trains.
4. Fits a line and estimates the size for ~5 µl/reward.
5. Validates the estimate.
6. Saves `rig.sipper_calib` and updates `rig.calibrated` when successful.

Entering `0` at a weight prompt restarts the calibration cycle.

## Mouse commands

### `lampyr mouse create MOUSEID`

Creates a mouse directory and mouse metadata file.

```bat
lampyr mouse create 014-003
lampyr mouse create 014-003 --paradigm BanditParadigm
lampyr mouse create 014-003 -p BanditParadigm --force
```

Arguments/options:

- `MOUSEID`: required mouse identifier.
- `--paradigm`, `-p`: optional behavior/paradigm class name to assign at creation.
- `--force`: overwrite/update even if an existing mouse is found.

Without `--force`, creation aborts if the mouse already exists. If a paradigm is provided at creation time, the command checks `lampyr.paradigms`; in the current application this dict is empty unless populated by code, so the most reliable workflow is to create the mouse first and then assign the paradigm with `lampyr mouse paradigm`.

### `lampyr mouse retire MOUSEID`

Marks a mouse as retired and saves it.

```bat
lampyr mouse retire 014-003
```

Retired mice are hidden from `mouse list` unless `-all` is used.

### `lampyr mouse list`

Prints a table of known mice.

```bat
lampyr mouse list
lampyr mouse list -recent
lampyr mouse list -all
```

Options:

- `-recent`: only show mice with a last session less than 12 hours ago.
- `-all`: include retired mice.

Columns include mouse ID, assigned paradigm, current stage, time since last session, last merit count, and last reward count.

### `lampyr mouse info MOUSEID`

Loads and prints a mouse object.

```bat
lampyr mouse info 014-003
```

### `lampyr mouse paradigm MOUSEID [PARADIGM_NAME]`

Inspect or change a mouse's assigned paradigm and current stage.

Inspect current paradigm data:

```bat
lampyr mouse paradigm 014-003
```

Set a paradigm:

```bat
lampyr mouse paradigm 014-003 BanditParadigm
```

Set a paradigm and stage:

```bat
lampyr mouse paradigm 014-003 BanditParadigm --stage AnyWheel
lampyr mouse paradigm 014-003 BanditParadigm -s AnyWheel
```

Set only the current stage for the already-assigned paradigm:

```bat
lampyr mouse paradigm 014-003 --stage AltWheel1
```

Stage names are validated against the paradigm class's `stagelist` slugs when available.

### `lampyr mouse run MOUSEID [BEHAVIOR]`

Loads a mouse, starts the rig, and runs a behavior.

```bat
lampyr mouse run 014-003
lampyr mouse run 014-003 BanditTask -dl 60 -rl 200 -sal 15
lampyr mouse run 014-003 BanditParadigm --duration_limit 60
```

Arguments/options:

- `MOUSEID`: required mouse identifier.
- `BEHAVIOR`: optional behavior/paradigm class name. If omitted, the mouse's assigned `mouse.paradigm` is used.
- Session options: all stop-condition options listed above.

Common failure cases:

- Unknown mouse ID: command prints an error and exits.
- Missing behavior when the mouse has no assigned paradigm: `Lampyr.run()` cannot find a valid segment.
- Rig unconfigured/uncalibrated: the command aborts before connecting.

## User/notification commands

Notification data is stored in the shared users config loaded by `NotificationManager`. Pushover delivery requires an app token and one or more user keys.

### `lampyr user set-token TOKEN`

Stores the shared Pushover app token.

```bat
lampyr user set-token APP_TOKEN
```

### `lampyr user create NAME`

Creates a notification user.

```bat
lampyr user create max --pushover_user_key USER_KEY
lampyr user create supervisor -super --pushover_user_key USER_KEY
```

Options:

- `--pushover_user_key`: Pushover user key.
- `-super`: mark the user as a supervisor. Supervisors are always active and cannot be deactivated.

### `lampyr user edit NAME`

Updates a notification user.

```bat
lampyr user edit max --pushover_user_key NEW_USER_KEY
lampyr user edit max --supervisor true
lampyr user edit max --supervisor false
```

Options:

- `--pushover_user_key`: replacement Pushover user key.
- `--supervisor BOOL`: set supervisor status.

### `lampyr user inspect NAME`

Prints one user's notification configuration.

```bat
lampyr user inspect max
```

### `lampyr user list`

Lists configured users and status tags.

```bat
lampyr user list
```

Tags:

- `[supervisor]`: always notified.
- `[active]`: normal active recipient.
- `[inactive]`: normal inactive recipient.

### `lampyr user activate NAME`

Marks a non-supervisor user active.

```bat
lampyr user activate max
```

### `lampyr user deactivate NAME`

Marks a non-supervisor user inactive.

```bat
lampyr user deactivate max
```

Supervisor users cannot be deactivated.

### `lampyr user ping`

Sends a test notification.

```bat
lampyr user ping
lampyr user ping --user max --message "test from rig"
lampyr user ping -u max -m "urgent test" --urgent
```

Options:

- `--user`, `-u`: send only to one user. If omitted, sends to every active/supervisor user.
- `--message`, `-m`: message body. Default is a generic test message.
- `--urgent`: send using Pushover emergency priority.

### `lampyr user remove NAME`

Deletes a notification user.

```bat
lampyr user remove max
```

## Typical CLI workflows

### First-time rig setup

```bat
lampyr info
lampyr rig configure
lampyr rig calibrate
lampyr rig info
```

### Add a mouse and assign training

```bat
lampyr mouse create 014-003 --paradigm BanditParadigm
lampyr mouse paradigm 014-003 --stage Habituation
lampyr mouse info 014-003
```

### Run a normal assigned paradigm

```bat
lampyr mouse run 014-003 -dl 60 -rl 200 -sal 15
```

### Run a specific task manually

```bat
lampyr mouse run 014-003 RewardedHabituationTask --duration_limit 30 --reward_limit 100
```

### Configure notifications

```bat
lampyr user set-token APP_TOKEN
lampyr user create max --pushover_user_key USER_KEY
lampyr user ping --user max --message "LAMPyR notification test"
```
