## Extended Data for "A Low-Cost, Modular Hardware and Software Platform for Head-Fixed Mouse Decision-Making Tasks": gui_workflow.md

# LAMPyR GUI workflow

LAMPyR's GUI is a Textual terminal user interface launched by:

```bat
lampyr go
```

The GUI is implemented in `lampyr/interfaces/textual_tui/app.py`. It is designed for touch-first operation on rig computers while still supporting mouse/keyboard interaction.

## Startup behavior

When `lampyr go` starts:

1. A `LampyrApp` Textual application is created.
2. A `Lampyr` object is created with custom input/output handlers so blocking prompts and log output can be routed through the TUI.
3. The main screen is pushed.
4. A heartbeat timer is scheduled every 20 minutes.

On Windows only, the CLI wrapper also:

- toggles fullscreen with F11;
- disables Quick Edit mode in the terminal;
- enables terminal mouse input;
- starts an invisible touch-to-mouse bridge unless `--notouchoverlay` is supplied.

```bat
lampyr go --notouchoverlay
```

Use `--notouchoverlay` if the transparent Windows touch bridge interferes with local input handling.

## Main screen

The main screen shows the configured rig name as ASCII art and four large buttons:

1. `RUN`
2. `ADVANCED`
3. `CALIBRATE`
4. `QUIT`

The rig name comes from `config['rig.name']`. If no rig name is configured, the title falls back to `Lampyr`.

## RUN workflow: normal mouse/paradigm run

Tap `RUN` for the default day-to-day workflow.

1. The GUI opens an ID-entry numpad.
2. Enter the mouse ID and confirm.
3. LAMPyR attempts to load the mouse.
4. If the mouse has an assigned `mouse.paradigm`, the GUI starts that behavior directly.
5. If the mouse does not have an assigned paradigm, the GUI opens the behavior selection screen instead.

Typical use:

```text
RUN → Enter Mouse ID → Run assigned paradigm
```

If the mouse ID cannot be loaded, the app displays an error notification and stays in the GUI.

## ADVANCED workflow: choose a task manually

Tap `ADVANCED` when you want to run a specific task instead of the mouse's assigned paradigm.

1. The GUI opens an ID-entry numpad.
2. Enter the mouse ID and confirm.
3. LAMPyR loads the mouse.
4. The behavior selection screen appears.
5. Pick a task.
6. The task-parameter screen appears.
7. Edit session limits if needed.
8. Tap `RUN`.

Typical use:

```text
ADVANCED → Enter Mouse ID → Select Task → Configure limits → RUN
```

The behavior selector lists imported subclasses of `Task` and excludes the abstract base `Task` itself. It does not list stages or paradigms in this advanced task-selection flow.

## Task parameter screen

Before an advanced task run, the GUI shows editable session parameters. Defaults are:

- `reward_limit = 200`
- `duration_limit = 60`
- `serial_abstention_limit = 15`

Editable parameters:

- Reward Limit
- Duration (min)
- Trial Limit
- Merit Limit
- Demerit Limit
- Participation Limit
- Abstention Limit
- Serial Abstention Limit

Tap a parameter to open the integer numpad. Enter a value and confirm. Entering `-` clears that parameter so it will not be passed as a session override.

When `RUN` is tapped, these values are passed as keyword arguments to `Lampyr.run()` and become `Session` stop conditions.

## Run screen

The run screen is a black log-output screen with one large action button.

During a session, the button is:

```text
ABORT
```

The run thread performs the following steps:

1. Load the mouse.
2. Check rig configuration.
3. Check calibration age.
4. Connect to the rig.
5. Start the selected behavior/paradigm.
6. Stream log output to the screen.

The rig checks mirror the CLI run checks:

- `rig.configured` must be truthy.
- `rig.calibrated` must be newer than five days.

If either check fails, the run screen reports the error and does not start a session.

### Aborting a run

Tap `ABORT` while a session is running to force a `KeyboardInterrupt` into the worker thread. The app also sets an internal `_user_aborted` flag on the `Lampyr` instance so the final stop reasons include user intervention.

After a session ends, the action button changes to a return button:

- `RETURN TO MAIN` for a clean finish.
- `RETURN (session ended with error)` for an error finish.

Returning cancels the post-session animal-left timer.

### Post-session animal-left alert

When a run finishes, the GUI starts a 30-minute timer. If the user has not returned from the run screen by then, the app sends an urgent notification warning that a mouse may have been left in the rig.

Recipients are active users and supervisors from the shared notification user configuration.

## Calibration workflow

Tap `CALIBRATE` to begin calibration.

1. A yellow confirmation screen appears.
2. Tap `BEGIN CALIBRATION` to proceed, or `CANCEL` to return.
3. The calibration screen opens a log panel.
4. LAMPyR connects to the rig if needed.
5. The calibration routine prompts for weights using the GUI numpad.
6. When calibration completes, `RETURN TO MAIN` appears.

Calibration is the same routine used by:

```bat
lampyr rig calibrate
```

It updates the stored sipper calibration and calibration timestamp on success.

## Automatic calibration reminder

The app heartbeat checks calibration age. If calibration is older than five days and no calibration/run screen is active, the GUI automatically pushes the calibration confirmation screen.

The heartbeat also touches `heartbeat.file` in the shared mice directory when no session is running. If that file operation fails, the GUI displays an error notification.

## Numpad modal

The GUI uses a touch-friendly numpad modal for:

- mouse IDs;
- integer session parameters;
- floating-point calibration weights.

Modes:

- `id`: digits plus `-`; requires a non-empty string.
- `int`: digits plus optional leading `-`; validates as an integer. A single `-` is used in the task parameter screen to clear a parameter.
- `float`: digits plus decimal point; validates as a float.

All entries use a confirmation step before submission.

## Output routing

The TUI routes normal LAMPyR text output into screen-specific RichLog widgets.

- Calibration output goes to the calibration log.
- Session output goes to the run log.
- Other screens suppress standard output unless they explicitly install an output handler.

This works through `LampyrApp.set_output()`, which updates the top-level `Lampyr` object and all managers (`rigmanager`, `mousemanager`, `datamanager`, and `notificationmanager`).

## Quitting

Tap `QUIT` on the main screen to exit the app.

Avoid closing the terminal during an active session unless necessary. The run screen's `ABORT` button gives LAMPyR a better chance to record stop reasons, dump segment data, stop rig acquisition, save the session, save the mouse, and send notifications.
