## Extended Data for "A Low-Cost, Modular Hardware and Software Platform for Head-Fixed Mouse Decision-Making Tasks": training_management.md

# LAMPyR training management

This guide covers common ways to tweak mouse training without changing task source code. It focuses on the CLI workflows in `lampyr/interfaces/click_cli/app.py`, the runtime objects in `lampyr/main.py` and `lampyr/primatives.py`, and behavior parameter overrides applied by `BehaviorSegment._checkoverrides()`.

For the full command list, see [CLI command reference](cli_commands.md). For how paradigms, stages, tasks, trials, slugs, and tags fit together, see [Application architecture and segment inheritance](app_architecture_and_segments.md).

## Normal training progression: assign paradigm and stage

A mouse stores its normal training assignment in:

- `mouse.paradigm` — the behavior/paradigm class name to run by default.
- `mouse.properties` — persistent paradigm state, including the current stage.

Use `lampyr mouse paradigm` for normal progression edits.

Inspect current paradigm data:

```bat
lampyr mouse paradigm 014-003
lampyr mouse info 014-003
```

Assign or change the paradigm:

```bat
lampyr mouse paradigm 014-003 BanditParadigm3
```

Set the current stage for the already-assigned paradigm:

```bat
lampyr mouse paradigm 014-003 --stage Stage1AnyWheel
```

Assign paradigm and stage together:

```bat
lampyr mouse paradigm 014-003 BanditParadigm3 --stage Stage2Correction
```

Stage names are stage `slug` values, not necessarily the Python class names. For Bandit examples, stage slugs include values such as `Stage0Hab`, `Stage1AnyWheel`, `Stage2Correction`, `Stage3Return`, `Stage5BanditTraining`, and `Stage6Bandit` depending on the paradigm.

After changing an assignment, verify it before running:

```bat
lampyr mouse paradigm 014-003
lampyr mouse info 014-003
```

## Run a specific task manually

The default mouse run uses the assigned paradigm:

```bat
lampyr mouse run 014-003
```

To run a specific behavior instead of the assigned paradigm, provide the behavior class name:

```bat
lampyr mouse run 014-003 RewardedHabituationTask -dl 30 -rl 50
lampyr mouse run 014-003 BanditTask --duration_limit 45 --trial_limit 100
```

This loads the mouse, so mouse-specific overrides still apply. Supplying a behavior this way does **not** change `mouse.paradigm` or the mouse's assigned stage; it is a manual one-off run.

You can see available imported behaviors with:

```bat
lampyr list
```

## Adjust session stop conditions from the CLI

`lampyr mouse run` and `lampyr run` accept session stop-condition options that are forwarded to `Session`. Common options are:

| Option | Short | Meaning |
|---|---:|---|
| `--duration_limit` | `-dl` | Stop after this many minutes. |
| `--reward_limit` | `-rl` | Stop after this many delivered rewards. |
| `--trial_limit` | `-tl` | Stop after this many trials. |
| `--serial_abstention_limit` | `-sal` | Stop after this many consecutive abstentions. |
| `--merit_limit` | `-ml` | Stop after this many merits. |
| `--demerit_limit` | `-dml` | Stop after this many demerits. |
| `--abstention_limit` | `-al` | Stop after this many abstentions. |
| `--participation_limit` | `-pl` | Stop after this many participations. |

Most limits also have corresponding minimum options, such as `--duration_min`, `--trial_min`, and `--reward_min`. Minimums gate later stop checks; for example, a trial limit with a duration minimum will not stop the session until the duration minimum has elapsed.

Examples:

```bat
REM Short, low-reward session
lampyr mouse run 014-003 -dl 30 -rl 75

REM Stop early if the mouse has 15 consecutive abstentions
lampyr mouse run 014-003 BanditTask -dl 60 -sal 15

REM Fixed-length manual task by trial count
lampyr mouse run 014-003 BanditTask --trial_limit 120 --reward_limit 200
```

If a stage sets its own session parameters internally, CLI values take precedence when they are already set on the `Session`; the stage will log a warning that a stage parameter was overridden.

## Mouse behavior parameter overrides

Per-mouse behavior tweaks live in:

```python
mouse.mouse_behav_param_overrides
```

There is not currently a dedicated CLI command known for editing this field. Use a careful Python edit or direct mouse JSON edit.

Overrides are applied when each `BehaviorSegment` is created. In the built-in hierarchy this means tasks and trials, not paradigm/stage session parameters. The override value must target an attribute that already exists on that segment; misspelled or nonexistent attributes are ignored.

### Override key matching

`BehaviorSegment._checkoverrides()` checks keys in this order:

1. `all` — applies to every behavior segment.
2. The segment's `slug`, such as `BanditTask`, `BanditTrial`, `RewHab`, or a custom slug.
3. Each inherited tag in `segment.tags`, such as `experiment` when a task defines that tag.

If multiple matching overrides set the same attribute, later applications can replace earlier values.

Example structure:

```python
{
    "BanditTrial": {
        "responsewindow_s": 5,
        "responsethresholds_deg": {"Left": 10, "Right": 10}
    },
    "BanditTask": {
        "taskblocks_enabled": False,
        "reward_prob_target": 100,
        "reward_prob_offtarget": 0
    },
    "experiment": {
        "enable_wheel_lock": True
    }
}
```

### Edit overrides with Python

This is usually safer than hand-editing JSON:

```python
from lampyr import Lampyr

app = Lampyr()
try:
    app.mousemanager.load("014-003")
    mouse = app.mouse

    overrides = mouse.mouse_behav_param_overrides
    overrides.setdefault("BanditTrial", {})["responsewindow_s"] = 5
    overrides.setdefault("BanditTask", {})["taskblocks_enabled"] = False

    app.mousemanager.save()
finally:
    app.close()
```

To remove an override when it is no longer needed:

```python
from lampyr import Lampyr

app = Lampyr()
try:
    app.mousemanager.load("014-003")
    overrides = app.mouse.mouse_behav_param_overrides

    overrides.get("BanditTrial", {}).pop("responsewindow_s", None)
    if overrides.get("BanditTrial") == {}:
        overrides.pop("BanditTrial")

    app.mousemanager.save()
finally:
    app.close()
```

### Edit overrides directly in mouse JSON

Mouse metadata is saved under the configured mice directory:

```text
<mice_directory>/<MOUSEID>/<MOUSEID>_mouse.lampyr.json
```

For example:

```text
N:/SHARED/Maxwell_LAMPyR_MouseData/014-003/014-003_mouse.lampyr.json
```

Edit only the `mouse_behav_param_overrides` field, preserve the rest of the file, keep valid JSON syntax, and make a backup first. JSON booleans are lowercase (`true`/`false`). The mouse history is stored separately in the history CSV, not in this metadata JSON. Example field value:

```json
"mouse_behav_param_overrides": {
  "BanditTrial": {
    "responsewindow_s": 5
  },
  "BanditTask": {
    "taskblocks_enabled": false,
    "reward_prob_target": 100,
    "reward_prob_offtarget": 0
  }
}
```

Do not edit raw session history files to implement training changes.

## Practical tweak examples

### Lower the session reward cap

Prefer a CLI stop condition for one-off caps:

```bat
lampyr mouse run 014-003 -rl 75
lampyr mouse run 014-003 BanditTask -rl 75 -dl 45
```

This changes only that run. It does not alter task code or mouse metadata.

### Shorten or lengthen the response window

`BanditTrial` has a `responsewindow_s` attribute. Apply a per-mouse override:

```python
mouse.mouse_behav_param_overrides.setdefault("BanditTrial", {})["responsewindow_s"] = 5
```

Remove it once the mouse should return to the task default.

### Change reward probabilities

For `BanditTask`, prefer task-level attributes when possible:

```python
mouse.mouse_behav_param_overrides.setdefault("BanditTask", {}).update({
    "reward_prob_target": 100,
    "reward_prob_offtarget": 0,
})
```

For advanced custom cases, `BanditTrial` also has `rewardprobs_perc`, but overriding trial-level probabilities bypasses the task's normal target-to-probability mapping for each created trial. Use this carefully.

### Disable or enable task blocks

`BanditTask` supports task blocks with `taskblocks_enabled`:

```python
mouse.mouse_behav_param_overrides.setdefault("BanditTask", {})["taskblocks_enabled"] = False
```

Related attributes include `taskblocks_sizerange` and `taskblocks_blockcounttype`; only use them for tasks that define those attributes.

### Adjust wheel thresholds

`BanditTrial` defines `responsethresholds_deg`:

```python
mouse.mouse_behav_param_overrides.setdefault("BanditTrial", {})[
    "responsethresholds_deg"
] = {"Left": 10, "Right": 20}
```

Be aware that some stages may also update this field through global shaping logic, so inspect the mouse after runs and document intentional manual changes.

## Warnings and best practices

- Preserve raw session history. Do not edit saved session JSON/HDF5 files to change training outcomes.
- Prefer `lampyr mouse paradigm ... --stage ...` for normal training progression.
- Prefer CLI stop-condition options for one-off shorter/longer sessions.
- Use mouse behavior overrides only for intentional per-mouse task/trial parameter changes.
- Document every override: mouse ID, date, reason, parameter, old value, new value, and planned removal criteria.
- Verify metadata before running with `lampyr mouse info MOUSEID` and `lampyr mouse paradigm MOUSEID`.
- Remove overrides when no longer needed so the mouse returns to the standard task/paradigm definition.
- Remember that overrides only apply to existing behavior segment attributes and only when future segments are constructed.
