## Extended Data for "A Low-Cost, Modular Hardware and Software Platform for Head-Fixed Mouse Decision-Making Tasks": bandit_R0_3_v0.ino

```
// This Firmware is intended for an Arduino UNO with the HudaHubR0.3 Shield

#include <Encoder.h>
#include <Servo.h>
#include <TaskScheduler.h>

// -----------------------------------------------------------------------------
// Pin Definitions
// -----------------------------------------------------------------------------

#define RIG_ID "BanditHudaHub_1"

#define SPEAKERPIN          6
#define REWARDPIN           9
#define MANUALREWARDPIN     13
#define LASERPIN            11
#define CAM_PIN

#define LICKOMETERDIGPIN    A2
#define LICKOMETERANALOGPIN A3

#define SERVOMOTORPIN       5

#define ROTARYPIN1          2
#define ROTARYPIN2          3

// -----------------------------------------------------------------------------
// Task Periods
// -----------------------------------------------------------------------------

#define LICK_PERIOD_MS      10
#define ROTARY_PERIOD_MS    20
#define CAM_PULSE_PERIOD_MS 50

// -----------------------------------------------------------------------------
// Hardware
// -----------------------------------------------------------------------------

Scheduler runner;

Encoder rotary(ROTARYPIN1, ROTARYPIN2);
Servo wheelLock;

// -----------------------------------------------------------------------------
// State
// -----------------------------------------------------------------------------

// Laser
bool rampingDown = false;
uint32_t rampStartMs = 0;
uint32_t rampDurationMs = 0;
int rampStartPWM = 255;

// Rotary
long rotarypos = 0;
long rotarypos_last = 0;
long rotarydiff = 0;

// Lick
int currentlickanalog = 0;
int currentlickdigital = 0;

// Reward
bool rewarding = false;
uint32_t rewardStart = 0;
uint32_t rewardsize = 500;      // microseconds
uint32_t rewardError = 0;

// Manual reward button
bool manualRewardState = HIGH;
bool manualRewardLastState = HIGH;

// Lock
int angle = 0;

// Loop frequency measurement
bool measureLoopRequested = false;

uint32_t loopCounter = 0;
uint32_t loopCounterStart = 0;

bool pauseAfterReport = false;
uint32_t pauseStart = 0;

// -----------------------------------------------------------------------------
// Reporting
// -----------------------------------------------------------------------------

void reportCommand(char cmd)
{
    Serial.print(millis());
    Serial.print("\tC\t");
    Serial.println(cmd);
}

void reportRotary()
{
    rotarypos = rotary.read();

    rotarydiff = rotarypos_last - rotarypos;

    Serial.print(millis());
    Serial.print("\tR\t");
    Serial.println(rotarydiff);

    rotarypos_last = rotarypos;
}

void reportLick()
{
    currentlickanalog = analogRead(LICKOMETERANALOGPIN);
    currentlickdigital = analogRead(LICKOMETERDIGPIN);

    Serial.print(millis());
    Serial.print("\tA\t");
    Serial.println(currentlickanalog);

    Serial.print(millis());
    Serial.print("\tL\t");
    Serial.println(currentlickdigital);
}

void reportRewardSize()
{
    Serial.print(millis());
    Serial.print("\tW\t");
    Serial.println(rewardsize);
}

void reportReward()
{
    Serial.print(millis());
    Serial.print("\tG\t");
    Serial.println(rewardsize);
}

void reportManualReward()
{
    Serial.print(millis());
    Serial.print("\tM\t");
    Serial.println(1);
}

void reportTime()
{
    Serial.print(millis());
    Serial.print("\tT\t");
    Serial.println(millis());
}

void reportRewardError()
{
    Serial.print(millis());
    Serial.print("\tE\t");
    Serial.println(rewardError);
}

void reportLoopHz(uint32_t hz)
{
    Serial.print(millis());
    Serial.print("\tF\t");
    Serial.println(hz);
}

// -----------------------------------------------------------------------------
// Reward Handling
// -----------------------------------------------------------------------------

void giveReward()
{
    if (rewarding)
        return;

    digitalWrite(REWARDPIN, HIGH);

    rewardStart = micros();
    rewarding = true;
}

void checkReward()
{
    if (!rewarding)
        return;

    uint32_t elapsed = micros() - rewardStart;

    if (elapsed >= rewardsize)
    {
        rewardError = elapsed - rewardsize;

        digitalWrite(REWARDPIN, LOW);

        rewarding = false;
    }
}

// -----------------------------------------------------------------------------
// Manual Reward Button
// -----------------------------------------------------------------------------

void checkManualReward()
{
    manualRewardState = digitalRead(MANUALREWARDPIN);

    if (manualRewardLastState == HIGH &&
        manualRewardState == LOW)
    {
        reportManualReward();

        giveReward();

        reportReward();
    }

    manualRewardLastState = manualRewardState;
}

// -----------------------------------------------------------------------------
// Commands
// -----------------------------------------------------------------------------

void executeCommand(char cmd)
{
    reportCommand(cmd);
    switch (cmd)
    {
        case 'p':
            tone(SPEAKERPIN, 1000, 100);
            break;

        case 'r':
            tone(SPEAKERPIN, 10000, 100);
            break;

        case 'b':
            tone(SPEAKERPIN, 4000, 100);
            break;

        case 'w':
        {
            String str = Serial.readStringUntil('\n');

            rewardsize = str.toInt();

            reportRewardSize();
            break;
        }

        case 'g':
        {
            giveReward();

            reportReward();
            break;
        }

        case 'z': // Start "zap" protocol for laser stim
        {
            rampingDown = false;
            analogWrite(LASERPIN, 255);
            break;
        }

        case 'x': // Hardcutoff to laser stim
        {
            rampingDown = false;
            analogWrite(LASERPIN, 0);
            break;
        }

        case 'c': // Slow rampdown to laser stim
        {
            String str = Serial.readStringUntil('\n');

            rampDurationMs = str.toInt() * 1000UL;
            rampStartMs = millis();
            rampStartPWM = 255;

            rampingDown = true;

            break;
        }

        case 'l': // Lock the wheel
        {
            wheelLock.write(60);
            break;
        }

        case 'u': // Unlock the wheel
        {
            wheelLock.write(90);
            break;
        }

        case 'a': // Set to specific angle
        {
            String str = Serial.readStringUntil('\n');

            angle = str.toInt();
            wheelLock.write(angle);
            break;
        }

        case 't':
        {
            reportTime();
            break;
        }

        
        case 'i':
        {
            Serial.print(millis());
            Serial.print("\tI\t");
            Serial.println(RIG_ID);
            break;
        }
        case 'e':
        {
            reportRewardError();
            break;
        }

        case 'f':
        {
            measureLoopRequested = true;

            loopCounter = 0;
            loopCounterStart = millis();

            break;
        }
    }
}

void handleSerial()
{
    while (Serial.available())
    {
        char cmd = (char)Serial.read();
        executeCommand(cmd);
    }
}

// -----------------------------------------------------------------------------
// Scheduled Tasks
// -----------------------------------------------------------------------------

Task lickTask(
    LICK_PERIOD_MS,
    TASK_FOREVER,
    &reportLick
);

Task rotaryTask(
    ROTARY_PERIOD_MS,
    TASK_FOREVER,
    &reportRotary
);

// -----------------------------------------------------------------------------
// Setup
// -----------------------------------------------------------------------------

void setup()
{
    Serial.begin(115200);

    pinMode(LED_BUILTIN, OUTPUT);

    pinMode(SPEAKERPIN, OUTPUT);

    pinMode(REWARDPIN, OUTPUT);
    digitalWrite(REWARDPIN, LOW);

    pinMode(MANUALREWARDPIN, INPUT_PULLUP);

    pinMode(LICKOMETERDIGPIN, INPUT);
    pinMode(LICKOMETERANALOGPIN, INPUT);

    wheelLock.attach(SERVOMOTORPIN);
    wheelLock.write(90);

    runner.init();

    runner.addTask(lickTask);
    runner.addTask(rotaryTask);

    lickTask.enable();
    rotaryTask.enable();
}

// -----------------------------------------------------------------------------
// Main Loop
// -----------------------------------------------------------------------------

void loop()
{
    // -------------------------------------------------------------
    // Post-report pause
    // -------------------------------------------------------------

    if (pauseAfterReport)
    {
        if ((millis() - pauseStart) >= 1000)
        {
            pauseAfterReport = false;
        }

        return;
    }

    // -------------------------------------------------------------
    // Normal operation
    // -------------------------------------------------------------

    handleSerial();

    checkManualReward();

    checkReward();

    runner.execute();

    // -------------------------------------------------------------
    // Loop frequency measurement
    // -------------------------------------------------------------

    if (measureLoopRequested)
    {
        loopCounter++;

        if ((millis() - loopCounterStart) >= 1000)
        {
            reportLoopHz(loopCounter);

            measureLoopRequested = false;

            pauseAfterReport = true;
            pauseStart = millis();
        }
    }

    // -------------------------------------------------------------
    // Laser rampdown
    // -------------------------------------------------------------
    if (rampingDown)
    {
        uint32_t elapsed = millis() - rampStartMs;

        if (elapsed >= rampDurationMs)
        {
            analogWrite(LASERPIN, 0);
            rampingDown = false;
        }
        else
        {
            int pwm =
                rampStartPWM -
                ((long)rampStartPWM * elapsed) / rampDurationMs;

            analogWrite(LASERPIN, pwm);
        }
    }
}
```
