## Extended Data for "A Low-Cost, Modular Hardware and Software Platform for Head-Fixed Mouse Decision-Making Tasks": banditv0_0_0.ino

```
#include <Encoder.h>

#define ROTARYREPORTFREQ 50 // 20Hz
#define LICKREPORTFREQ 10 // 100Hz

//pin definitions
#define SPEAKERPIN 5
#define REWARDPIN 12
#define LICKOMETERPIN A0

Encoder rotary(2,3);


long rotarypos;
long rotarypos_last;
long rotarydiff;
int currentlickval;

long rewardsize = 500;
bool rewarding;
long rewardbegin_time;

//REPORTS
long current_time;
long last_rotary_report;
long last_lick_report;

void setup() {
  // put your setup code here, to run once:
  SPCR = 0;
  digitalWrite(REWARDPIN, LOW);
  Serial.begin(115200);
  while (!Serial);
  pinMode(LED_BUILTIN, OUTPUT);
  pinMode(SPEAKERPIN, OUTPUT);
  pinMode(REWARDPIN, OUTPUT);
  digitalWrite(REWARDPIN, LOW);
  pinMode(A0, INPUT);
  delay(1000);
}

void reportRotary() {
  rotarypos = rotary.read();
  Serial.print(millis());
  Serial.print("\t");
  Serial.print("R");
  Serial.print("\t");
  rotarydiff = rotarypos_last-rotarypos;
  Serial.println(rotarydiff,DEC);
  rotarypos_last = rotarypos;
}

void reportLick() {
  currentlickval = analogRead(A0);
  Serial.print(millis());
  Serial.print("\t");
  Serial.print("L");
  Serial.print("\t");
  Serial.println(currentlickval, DEC);
}

void reportRewardSizeCalibration() {
  Serial.print(millis());
  Serial.print("\t");
  Serial.print("W");
  Serial.print("\t");
  Serial.println(rewardsize, DEC);
}

void reportReward() {
  Serial.print(millis());
  Serial.print("\t");
  Serial.print("G");
  Serial.print("\t");
  Serial.println(rewardsize, DEC);
}
\
void reportTime() {
  Serial.print(millis());
  Serial.print("\t");
  Serial.print("T");
  Serial.print("\t");
  Serial.println(current_time, DEC);
}

void executeCommand(char cmd) {
  switch (cmd) {
    case 'p': // Punishtone
      tone(SPEAKERPIN, 1000, 100);
      break;
    case 'r': // Rewardtone
      tone(SPEAKERPIN, 10000, 100);
      break;
    case 'b': // Trial begin tone
      tone(SPEAKERPIN, 4000, 100);
      break;
    case 'w': {// Set reward size 
      String str = Serial.readStringUntil('\n');
      rewardsize = str.toInt();
      reportRewardSizeCalibration();
      break;
    }
    case 'g':
      digitalWrite(REWARDPIN, HIGH);
      rewarding = true;
      rewardbegin_time = micros();
      reportReward();
      break;
    case 't':
      reportTime();
      break;
  }
}

void loop() {
  if (Serial.available()>0) {
    char cmd = (char)Serial.read();
    executeCommand(cmd);
  }
  current_time = millis();
  //HANDLE REPORTS
  if ((current_time - last_rotary_report) >= ROTARYREPORTFREQ){
    last_rotary_report = current_time;
    reportRotary();
  }
  if ((current_time - last_lick_report)>=LICKREPORTFREQ) {
    last_lick_report = current_time;
    reportLick();
  }
  if (((micros() - rewardbegin_time) >= rewardsize)) {
    digitalWrite(REWARDPIN, LOW);
    rewarding = false;
  }
}
```
