## Extended Data for "A Low-Cost, Modular Hardware and Software Platform for Head-Fixed Mouse Decision-Making Tasks": lampyr.yaml

```
name: lampyr #Environment for hardware control/behavior rigs/arduino
channels:
  - conda-forge
  - defaults
dependencies:
  - python=3.12
  - spyder-kernels=3.1
  - pytest
  - pyserial
  - numpy>=1.21
  - pandas
  - textual
  - textual-dev
  - click
  - keyboard
  - git
  - pip
  - h5py
  - requests
  - opencv>=4.12
```
