## Extended Data for "A Low-Cost, Modular Hardware and Software Platform for Head-Fixed Mouse Decision-Making Tasks": actions.py

```
# -*- coding: utf-8 -*-
"""
Created on Thu Jun 12 15:21:44 2025

@author: mm4114
"""

import click
import time
from lampyr import Lampyr

class Abort(Exception):
    """Raised by CLI action helpers to signal a non-error early exit."""
    pass

def printtitle(msg, char = '='):
    """
    Print a prominent title block surrounded by header lines.

    Parameters
    ----------
    msg : str
        Title text to display.
    char : str, optional
        Character used to build the border lines. Default is ``'='``.
    """
    printheader('',char)
    printheader(f' {msg} ', char)
    printheader('',char)

def printheader(msg, char = '-'):
    """
    Print a single centred header line padded with ``char`` to 80 characters.

    Parameters
    ----------
    msg : str
        Text to display in the centre of the line.
    char : str, optional
        Padding character. Default is ``'-'``.
    """
    prefix = char * (40 - len(msg)//2)
    if len(msg) % 2 == 1:
        suffix = prefix[:-1]
    else:
        suffix = prefix
    click.echo(prefix+msg+suffix)

def printinfo(info, indent = 0):
    """
    Recursively pretty-print a nested dict using Click.

    Parameters
    ----------
    info : dict
        Data to display.  Dict values are printed indented; list values are
        printed one item per line (with nested dicts re-entering recursion).
    indent : int, optional
        Current indentation level (tab characters). Default is 0.
    """
    prefix = '\t'*indent
    for k, v in info.items():
        if isinstance(v, dict):
            click.echo(f'{prefix}{k}')
            printinfo(v, indent+1)
        elif isinstance(v, list):
            click.echo(f'{prefix}{k}')
            for vind in v:
                if isinstance(vind, dict):
                    printinfo(vind, indent+1)
                else:
                    click.echo(f'\t{prefix}{vind}')
        else:
            click.echo(f'{prefix}{k} : {v}')

def start_rig(lampyr : Lampyr):
    """
    Verify rig configuration and calibration, then connect.

    Parameters
    ----------
    lampyr : Lampyr
        Active Lampyr instance.

    Returns
    -------
    bool
        ``True`` if the rig connected successfully.

    Raises
    ------
    Abort
        If the rig is not configured or calibration is more than 12 hours old.
    """
    if lampyr.config.get('rig.configured') < 1:
        click.echo('Command aborted because rig is not configured')
        click.echo('\nUse "lampyr rig configure" to configure rig')
        raise Abort()
    if lampyr.config.get('rig.calibrated') < time.time() - (5*24*60*60):
        click.echo('Command aborted because rig is not calibrated')
        click.echo('\nUse "lampyr rig calibrate" to calibrate rig')
        raise Abort()
    lampyr.rigmanager.connect()
    return True

def configure_rig(lampyr : Lampyr):
    """
    Interactively configure the rig name and mark it as configured.

    Prompts the user for a rig name, then stores it in
    ``config['rig.name']`` and sets ``config['rig.configured']`` to 1.

    Parameters
    ----------
    lampyr : Lampyr
        Active Lampyr instance.
    """
    name = input('Rig Name: ')
    lampyr.config.set('rig.name', name)
    lampyr.config.set('rig.configured', 1)
```
