## Extended Data for "A Low-Cost, Modular Hardware and Software Platform for Head-Fixed Mouse Decision-Making Tasks": __init__.py

```
# -*- coding: utf-8 -*-
"""
Created on Wed Dec  3 18:26:06 2025

@author: mm4114
"""

from .colony import Colony
from .dataset import MultiSessionDataset, TraceExtractionProfile, SegmentReference, TraceReference
from .traces import dynamic_trace_extraction
from .time import TimeTranslator, create_syncstream
```
