## Extended Data for "A Low-Cost, Modular Hardware and Software Platform for Head-Fixed Mouse Decision-Making Tasks": colony2.py

```
# -*- coding: utf-8 -*-
"""
Created on Mon Jun 15 16:42:30 2026

@author: mm4114
"""
from datetime import date, datetime, timedelta
from time import time

from typing import Callable, List, Tuple

from lampyr.managers.data import DataHandler


class MouseQuery:
    def __init__(self, colony, predicates: Tuple[Callable, ...] = ()):
        self.colony = colony
        self.predicates = predicates

    def _clone(self, predicate: Callable):
        return MouseQuery(self.colony, self.predicates + (predicate,))

    def where(self, predicate: Callable):
        return self._clone(predicate)

    def paradigm(self, paradigm: str):
        return self.where(lambda mouse: mouse.paradigm == paradigm)

    def stage(self, slug: str, stage: str):
        return self.where(
            lambda mouse: mouse.properties.get(slug, {}).get("stage") == stage
        )

    def retired(self, retired: bool = True):
        return self.where(lambda mouse: mouse.retired == retired)

    def active(self):
        return self.retired(False)

    def property(self, key: str, value):
        return self.where(lambda mouse: mouse.properties.get(key) == value)

    def properties(self, **properties):
        query = self
        for key, value in properties.items():
            query = query.property(key, value)
        return query

    def ids(self) -> List[str]:
        mouseids, _ = self.colony.data.mouselist()
        result = []

        for mouseid in mouseids:
            if mouseid == "UNKNOWN_MOUSE":
                continue

            mouse = self.colony.load_mouse(mouseid)
            if all(predicate(mouse) for predicate in self.predicates):
                result.append(mouseid)

        return result

    def count(self) -> int:
        return len(self.ids())

    def objects(self) -> list:
        return [self.colony.load_mouse(mouseid) for mouseid in self.ids()]

    def sessions(self, **filters):
        return self.colony.sessions(mouseids=self.ids(), **filters)


class SessionQuery:
    RANGE_KEYS = {
        "duration",
        "merit",
        "demerit",
        "trial",
        "rewards",
        "abstention",
        "participation",
        "starttime",
        "age"
    }

    def __init__(self,
                 colony,
                 mouseids: Tuple[str, ...],
                 predicates: Tuple[Callable, ...] = (),
                 clauses: Tuple[str, ...] = ()):
        self.colony = colony
        self.mouseids = mouseids
        self.predicates = predicates
        self.clauses = clauses

    def _clone(self, predicate: Callable, clause: str = None):
        clauses = self.clauses
        if clause is not None:
            clauses = clauses + (clause,)

        return SessionQuery(
            self.colony,
            self.mouseids,
            self.predicates + (predicate,),
            clauses,
        )

    def session(self, sessionid):
        for mouseid, entry in self._matched_entries():
            if self.colony._sessionid(entry) == sessionid:
                session = self.colony.load_session(
                    sessionid,
                    mouseid,
                )
                if session is not None:
                    return session
                break
    
        raise KeyError(
            f"Session '{sessionid}' not found in query or failed to load."
        )

    def where(self, predicate: Callable):
        return self._clone(predicate)

    def sessionids(self, sessionids):
        selected_ids = {
            str(sessionid)
            for sessionid in sessionids
            if sessionid not in (None, "")
        }

        return self._clone(
            lambda entry: (
                self.colony._sessionid(entry) is not None
                and str(self.colony._sessionid(entry)) in selected_ids
            ),
            f"sessionids in [{', '.join(sorted(selected_ids))}]",
        )

    def root(self, rootslug: str):
        """Filter sessions by the root segment slug saved in mouse history."""
        return self._clone(
            lambda entry: entry.get("rootslug") == rootslug,
            f"root == {rootslug}",
        )

    def range(self, key: str, minimum=None, maximum=None):
        if key not in self.RANGE_KEYS:
            raise ValueError(f"Unknown session filter: {key}")

        def predicate(entry):
            value = entry.get(key)

            if value in (None, ""):
                return False

            try:
                value = float(value)
            except (TypeError, ValueError):
                return False

            if minimum is not None and value < minimum:
                return False

            if maximum is not None and value > maximum:
                return False

            return True

        return self._clone(predicate, self._format_range_clause(key, minimum, maximum))

    def duration(self, minimum=None, maximum=None):
        return self.range("duration", minimum, maximum)

    def merit(self, minimum=None, maximum=None):
        return self.range("merit", minimum, maximum)

    def demerit(self, minimum=None, maximum=None):
        return self.range("demerit", minimum, maximum)

    def trial(self, minimum=None, maximum=None):
        return self.range("trial", minimum, maximum)

    def rewards(self, minimum=None, maximum=None):
        return self.range("rewards", minimum, maximum)

    def abstention(self, minimum=None, maximum=None):
        return self.range("abstention", minimum, maximum)

    def participation(self, minimum=None, maximum=None):
        return self.range("participation", minimum, maximum)

    def starttime(self, minimum=None, maximum=None):
        return self.range("starttime", minimum, maximum)

    @staticmethod
    def _timestamp_bound(value, *, is_end: bool):
        if value is None:
            return None

        if isinstance(value, (int, float)):
            return float(value)

        if isinstance(value, datetime):
            return value.timestamp()

        if isinstance(value, date):
            dt = datetime.combine(value, datetime.min.time())
            if is_end:
                dt += timedelta(days=1)
            return dt.timestamp()

        if isinstance(value, str):
            value = value.strip()

            if not value:
                return None

            for parser in (datetime.fromisoformat, date.fromisoformat):
                try:
                    parsed = parser(value)
                except ValueError:
                    continue

                if isinstance(parsed, datetime):
                    return parsed.timestamp()

                dt = datetime.combine(parsed, datetime.min.time())
                if is_end:
                    dt += timedelta(days=1)
                return dt.timestamp()

        raise ValueError(
            "Date bounds must be date, datetime, timestamp, or ISO-format string."
        )

    def date_range(self, start=None, end=None):
        """
        Filter sessions by calendar date or datetime range.

        `start` is inclusive. `end` is inclusive for date-only values and
        exclusive for datetime/timestamp values.
        """
        minimum = self._timestamp_bound(start, is_end=False)
        maximum = self._timestamp_bound(end, is_end=True)

        def predicate(entry):
            starttime = self.colony._starttime(entry)

            if starttime is None:
                return False

            if minimum is not None and starttime < minimum:
                return False

            if maximum is not None and starttime >= maximum:
                return False

            return True

        return self._clone(
            predicate,
            self._format_date_range_clause(start, end),
        )
    
    def age(self, minimum=None, maximum=None):
        """
        Session age in days.
        age(0, 30) => last 30 days
        age(30, None) => older than 30 days
        """
        now = time()
    
        def predicate(entry):
            starttime = self.colony._starttime(entry)
    
            if starttime is None:
                return False
    
            age_days = (now - starttime) / 86400
    
            if minimum is not None and age_days < minimum:
                return False
    
            if maximum is not None and age_days > maximum:
                return False
    
            return True
    
        return self._clone(predicate, self._format_range_clause("age_days", minimum, maximum))
    
    def younger_than(self, days: float):
        """
        Include only sessions that occurred within the last N days.
        """
        cutoff = time() - days * 24 * 60 * 60
    
        return self._clone(
            lambda entry: (
                self.colony._starttime(entry) is not None
                and self.colony._starttime(entry) >= cutoff
            ),
            f"age_days <= {days}",
        )
    
    def older_than(self, days: float):
        """
        Include only sessions older than N days.
        """
        cutoff = time() - days * 24 * 60 * 60
    
        return self._clone(
            lambda entry: (
                self.colony._starttime(entry) is not None
                and self.colony._starttime(entry) < cutoff
            ),
            f"age_days > {days}",
        )

    def filters(self, **filters):
        query = self

        for key, bounds in filters.items():
            if bounds is None:
                continue

            try:
                minimum, maximum = bounds
            except (TypeError, ValueError):
                raise ValueError(
                    f"Filter '{key}' must be a (minimum, maximum) tuple."
                ) from None

            query = query.range(key, minimum, maximum)

        return query

    def _matched_entries(self) -> list:
        result = []

        for mouseid in self.mouseids:
            mouse = self.colony.load_mouse(mouseid)

            for entry in mouse.history:
                if self.colony._sessionid(entry) is None:
                    continue

                predicate_entry = {**entry, "mouseid": mouseid}
                if all(predicate(predicate_entry) for predicate in self.predicates):
                    result.append((mouseid, entry))

        return result

    def ids(self) -> List[str]:
        return [
            self.colony._sessionid(entry)
            for _, entry in self._matched_entries()
        ]

    def matched_mouseids(self) -> List[str]:
        seen = set()
        result = []

        for mouseid, _ in self._matched_entries():
            if mouseid in seen:
                continue

            seen.add(mouseid)
            result.append(mouseid)

        return result

    def count(self) -> int:
        return len(self._matched_entries())

    def describe(self) -> str:
        matched_mouseids = self.matched_mouseids()
        lines = [
            (
                f"Resolved Mouse IDs ({len(matched_mouseids)}): "
                f"{', '.join(map(str, matched_mouseids)) if matched_mouseids else '(none)'}"
            ),
        ]

        if self.clauses:
            lines.append("Session Filters:")
            lines.extend(f"- {clause}" for clause in self.clauses)
        else:
            lines.append("Session Filters: none")

        return "\n".join(lines)

    def mice(self) -> list:
        return [
            self.colony.load_mouse(mouseid)
            for mouseid in self.matched_mouseids()
        ]

    def entries(self) -> list:
        return [
            {**entry, "mouseid": mouseid}
            for mouseid, entry in self._matched_entries()
        ]

    def objects(self) -> list:
        sessions = []

        for mouseid, entry in self._matched_entries():
            session = self.colony.load_session(self.colony._sessionid(entry), mouseid)
            if session is not None:
                sessions.append(session)

        return sessions

    def last_ids(self, n: int = 1) -> List[str]:
        sessionids = []

        for mouseid in self.mouseids:
            entries = [
                entry
                for matched_mouseid, entry in self._matched_entries()
                if (
                    matched_mouseid == mouseid
                    and self.colony._starttime(entry) is not None
                )
            ]
            entries.sort(
                key=self.colony._starttime,
                reverse=True,
            )
            sessionids.extend(
                self.colony._sessionid(entry)
                for entry in entries[:n]
            )

        return sessionids

    def last(self, n: int = 1) -> list:
        sessions = []

        for mouseid in self.mouseids:
            entries = [
                entry
                for matched_mouseid, entry in self._matched_entries()
                if (
                    matched_mouseid == mouseid
                    and self.colony._starttime(entry) is not None
                )
            ]
            entries.sort(
                key=self.colony._starttime,
                reverse=True,
            )
            for entry in entries[:n]:
                session = self.colony.load_session(
                    self.colony._sessionid(entry),
                    mouseid,
                )
                if session is not None:
                    sessions.append(session)

        return sessions

    def tail(self, n: int = 1):
        """
        Return a new SessionQuery restricted to the most recent N sessions
        for each mouse in the query.

        Unlike `last()`, this remains a query object and can still be passed
        to plots or further filtered.
        """
        selected_sessionids = []

        for mouseid in self.mouseids:
            mouse_query = SessionQuery(
                self.colony,
                (mouseid,),
                self.predicates,
                self.clauses,
            )
            selected_sessionids.extend(
                mouse_query.last_ids(n)
            )

        return self.sessionids(selected_sessionids)._with_clause(f"tail({n}) per mouse")

    def bymouse(self) -> dict:
        return {
            mouseid: SessionQuery(
                self.colony,
                (mouseid,),
                self.predicates,
                self.clauses,
            )
            for mouseid in self.matched_mouseids()
        }

    def _with_clause(self, clause: str):
        return SessionQuery(
            self.colony,
            self.mouseids,
            self.predicates,
            self.clauses + (clause,),
        )

    @staticmethod
    def _format_bound(value):
        if value is None:
            return "None"
        if isinstance(value, datetime):
            return value.isoformat(sep=" ", timespec="seconds")
        if isinstance(value, date):
            return value.isoformat()
        return str(value)

    @classmethod
    def _format_range_clause(cls, key: str, minimum=None, maximum=None):
        return (
            f"{key} in "
            f"[{cls._format_bound(minimum)}, {cls._format_bound(maximum)}]"
        )

    @classmethod
    def _format_date_range_clause(cls, start=None, end=None):
        return (
            "date_range in "
            f"[{cls._format_bound(start)}, {cls._format_bound(end)}]"
        )


class Colony:
    def __init__(self,
                 config=None,
                 keep="sessions",
                 verbose: bool = False,
                 skip_bad_sessions: bool = False):
        """
        Parameters
        ----------
        config : optional
            Configuration passed to :class:`lampyr.managers.data.DataHandler`.
        keep : {None, 'none', 'mice', 'sessions', 'all'}, optional
            Which loaded objects to cache in memory.
        verbose : bool, optional
            Print loading and skip messages.
        skip_bad_sessions : bool, optional
            If True, sessions that are missing or fail to load are skipped
            instead of raising an exception when session objects are requested.
        """
        self.data = DataHandler(config=config)
        self.keep_mice, self.keep_sessions = self._resolve_keep(keep)
        self.verbose = verbose
        self.skip_bad_sessions = skip_bad_sessions
        self.skipped_sessions = []
        self._mouse_cache = {}
        self._session_cache = {}

    def mice(self,
             paradigm=None,
             slug=None,
             stage=None,
             retired=None,
             properties=None,
             where=None) -> MouseQuery:
        query = MouseQuery(self)

        if paradigm is not None:
            query = query.paradigm(paradigm)

        if stage is not None:
            query = query.stage(slug, stage)

        if retired is not None:
            query = query.retired(retired)

        if properties is not None:
            query = query.properties(**properties)

        if where is not None:
            query = query.where(where)

        return query

    def sessions(self,
                 mouseid=None,
                 mouseids=None,
                 sessionids=None,
                 paradigm=None,
                 slug=None,
                 stage=None,
                 retired=None,
                 properties=None,
                 where_mouse=None,
                 duration=None,
                 merit=None,
                 demerit=None,
                 trial=None,
                 rewards=None,
                 abstention=None,
                 participation=None,
                 starttime=None,
                 date_range=None,
                 root=None,
                 where=None) -> SessionQuery:
        if mouseid is not None and mouseids is not None:
            raise ValueError("Pass only one of mouseid or mouseids.")

        if mouseid is not None:
            selected_mouseids = (mouseid,)
        elif mouseids is not None:
            selected_mouseids = tuple(mouseids)
        else:
            selected_mouseids = tuple(
                self.mice(
                    paradigm=paradigm,
                    slug=slug,
                    stage=stage,
                    retired=retired,
                    properties=properties,
                    where=where_mouse,
                ).ids()
            )

        query = SessionQuery(self, selected_mouseids).filters(
            duration=duration,
            merit=merit,
            demerit=demerit,
            trial=trial,
            rewards=rewards,
            abstention=abstention,
            participation=participation,
            starttime=starttime,
        )

        if where is not None:
            query = query.where(where)

        if date_range is not None:
            try:
                start, end = date_range
            except (TypeError, ValueError):
                raise ValueError(
                    "date_range must be a (start, end) tuple."
                ) from None

            query = query.date_range(start, end)

        if root is not None:
            query = query.root(root)

        if sessionids is not None:
            query = query.sessionids(sessionids)

        return query

    def load_mouse(self, mouseid: str):
        if not self.keep_mice:
            self._log(f"Loading mouse {mouseid}")
            return self.data.loadmouse(mouseid)

        if mouseid not in self._mouse_cache:
            self._log(f"Loading mouse {mouseid}")
            self._mouse_cache[mouseid] = self.data.loadmouse(mouseid)

        return self._mouse_cache[mouseid]

    def load_session(self, sessionid: str, mouseid: str):
        if not self.keep_sessions:
            self._log(f"Loading session {sessionid} for mouse {mouseid}")
            try:
                return self.data.loadsession(sessionid, mouseid)
            except Exception as exc:
                return self._handle_session_load_error(sessionid, mouseid, exc)

        key = (mouseid, sessionid)
        if key not in self._session_cache:
            self._log(f"Loading session {sessionid} for mouse {mouseid}")
            try:
                self._session_cache[key] = self.data.loadsession(sessionid, mouseid)
            except Exception as exc:
                session = self._handle_session_load_error(sessionid, mouseid, exc)
                if session is None and self.skip_bad_sessions:
                    self._session_cache[key] = None
                return session

        return self._session_cache[key]

    def _handle_session_load_error(self, sessionid: str, mouseid: str, exc: Exception):
        """Handle a failed session load, optionally skipping the bad session."""
        if self.skip_bad_sessions or isinstance(exc, FileExistsError):
            self.skipped_sessions.append({
                "mouseid": mouseid,
                "sessionid": sessionid,
                "error": exc,
            })
            self._log(f"Skipping session {sessionid} for mouse {mouseid}: {exc}")
            return None

        raise exc

    def clear_cache(self, mice: bool = True, sessions: bool = True):
        if mice:
            self._mouse_cache.clear()
        if sessions:
            self._session_cache.clear()

    @staticmethod
    def _resolve_keep(keep):
        modes = {
            None: (False, False),
            "none": (False, False),
            "mice": (True, False),
            "sessions": (False, True),
            "all": (True, True),
        }

        if keep not in modes:
            raise ValueError("keep must be one of: None, 'none', 'mice', 'sessions', 'all'")

        return modes[keep]

    def _log(self, message: str):
        if self.verbose:
            print(message)

    @staticmethod
    def _sessionid(entry: dict):
        for key in ("sessionid", "uniquesessionid", "session_id", "id"):
            if key in entry and entry[key] not in (None, ""):
                return entry[key]

        return None

    @staticmethod
    def _starttime(entry: dict):
        value = entry.get("starttime")

        if value in (None, ""):
            return None

        try:
            return float(value)
        except (TypeError, ValueError):
            return None
```
