## Extended Data for "A Low-Cost, Modular Hardware and Software Platform for Head-Fixed Mouse Decision-Making Tasks": colony.py

```
# -*- coding: utf-8 -*-
"""
Created on Fri Apr 24 16:18:48 2026

@author: mm4114
"""
from typing import Tuple, List

from lampyr.managers.abstract import AbstractManager
from lampyr.managers.data import DataHandler


class RetreiveSessions(AbstractManager):
    """
    Retrieves session data for specific animals.

    Wraps DataHandler to filter a mouse's session history by min/max bounds
    on any recorded session property.
    """
    def __init__(self, config=None, data=None):
        self._injected_data = data
        super().__init__(config=config)

    def start(self):
        self.data = self._injected_data or DataHandler(config=self.config)
        self.filters = {}
        self.mouseid = None
        self._active_mouse = None

    def retreival_settings(self,
                           duration: Tuple[int] = None,
                           merit: Tuple[float] = None,
                           demerit: Tuple[float] = None,
                           trial: Tuple[int] = None,
                           rewards: Tuple[int] = None,
                           abstention: Tuple[float] = None,
                           participation: Tuple[float] = None,
                           year: Tuple[int] = None,
                           month: Tuple[int] = None,
                           day: Tuple[int] = None):
        """
        Set min/max filters for session retrieval.

        Each parameter accepts a (min, max) tuple. Use None for either bound
        to leave it unconstrained.
        """
        self.filters = {
            'duration': duration,
            'merit': merit,
            'demerit': demerit,
            'trial': trial,
            'rewards': rewards,
            'abstention': abstention,
            'participation': participation,
            'year': year,
            'month': month,
            'day': day,
        }
        
    def load_session_from_id(self, sessionid: str, mouseid: str):
        return self.data.loadsession(sessionid, mouseid)

    @staticmethod
    def _entry_sessionid(entry: dict):
        """Return the session ID from a history entry, tolerating key variants."""
        for key in ('sessionid', 'uniquesessionid', 'session_id', 'id'):
            if key in entry and entry[key] not in (None, ''):
                return entry[key]
        return None

    def _passes_filters(self, entry: dict) -> bool:
        for key, bounds in self.filters.items():
            if bounds is None:
                continue
            val = entry.get(key)
            if val is None or val == '':
                continue
            try:
                val = float(val)
            except (TypeError, ValueError):
                continue
            lo, hi = bounds
            if lo is not None and val < lo:
                return False
            if hi is not None and val > hi:
                return False
        return True

    def set_mouse(self, mouseid: str):
        """
        Set the active mouse for subsequent retrieve calls.

        Loads the mouse from disk and caches it for repeated queries.

        Parameters
        ----------
        mouseid : str
            The mouse ID to use as the active mouse.
        """
        self.mouseid = mouseid
        self._active_mouse = self.data.loadmouse(mouseid)

    def retrieve(self, mouseid: str) -> List[str]:
        """
        Return session IDs from a mouse's history that pass the current filters.

        Parameters
        ----------
        mouseid : str
            The mouse whose session history to filter.

        Returns
        -------
        list of str
            Session IDs matching all active filter constraints.
        """
        mouse = self.data.loadmouse(mouseid)
        return [sid for entry in mouse.history
                if self._passes_filters(entry)
                and (sid := self._entry_sessionid(entry)) is not None]

    def retrieve_last(self, n: int = 1) -> list:
        """
        Return the last n loaded Session objects for the active mouse,
        ordered most-recent first.

        Applies current filters before selecting. Raises RuntimeError if no
        mouse has been set via set_mouse.

        Parameters
        ----------
        n : int, optional
            Number of sessions to return (default 1). Pass a higher value to
            retrieve earlier sessions as well.

        Returns
        -------
        list of Session
            Up to n loaded Session objects, sorted by starttime descending.
        """
        ids = self.retrieve_last_ids(n)
        return [self.data.loadsession(sid, self.mouseid) for sid in ids]

    def retrieve_last_ids(self, n: int = 1) -> List[str]:
        """
        Return the last n session IDs for the active mouse, ordered most-recent
        first.

        Same selection logic as :meth:`retrieve_last` but returns only the IDs
        without loading the Session objects from disk.

        Parameters
        ----------
        n : int, optional
            Number of session IDs to return (default 1).

        Returns
        -------
        list of str
            Up to n session IDs, sorted by starttime descending.
        """
        if self._active_mouse is None:
            raise RuntimeError('No mouse set — call set_mouse first.')
        passing = [e for e in self._active_mouse.history
                   if self._passes_filters(e)
                   and self._entry_sessionid(e) is not None]
        passing.sort(key=lambda e: float(e['starttime']), reverse=True)
        return [self._entry_sessionid(e) for e in passing[:n]]


class RetreiveMice(AbstractManager):
    """
    Retrieves and filters mice from the colony.

    Wraps DataHandler to list and filter Mouse objects by paradigm, stage,
    retirement status, or arbitrary properties.
    """
    def __init__(self, config=None, data=None):
        self._injected_data = data
        super().__init__(config=config)

    def start(self):
        self.data = self._injected_data or DataHandler(config=self.config)

    def _mouse_passes_filter(self, mouse,
                             paradigm: str,
                             slug: str,
                             stage: str,
                             retired: bool,
                             properties: dict) -> bool:
        if paradigm is not None and mouse.paradigm != paradigm:
            return False
        if stage is not None and mouse.properties.get(slug, {}).get('stage') != stage:
            return False
        if retired is not None and mouse.retired != retired:
            return False
        if properties is not None:
            for key, val in properties.items():
                if mouse.properties.get(key) != val:
                    return False
        return True

    def get_mice(self,
                 paradigm: str = None,
                 slug: str = None,
                 stage: str = None,
                 retired: bool = None,
                 properties: dict = None) -> List[str]:
        """
        Return mouse IDs from disk, optionally filtered.

        Parameters
        ----------
        paradigm : str, optional
            Keep only mice whose ``mouse.paradigm`` matches this value
            (the paradigm class name, e.g. ``'BanditParadigm'``).
        slug : str, optional
            Paradigm slug used as the key into ``mouse.properties``
            (e.g. ``'BanditParadigm2'``).  Required when filtering by
            ``stage``.
        stage : str, optional
            Keep only mice whose current stage at ``mouse.properties[slug]['stage']``
            matches this value.
        retired : bool, optional
            ``True`` — only retired mice.  ``False`` — only active mice.
            ``None`` (default) — all mice regardless of retirement status.
        properties : dict, optional
            Key/value pairs matched against top-level keys of
            ``mouse.properties``.  A mouse passes if every pair matches.

        Returns
        -------
        list of str
            Mouse IDs that satisfy all specified filters.
        """
        mouseids, _ = self.data.mouselist()
        result = []
        for mid in mouseids:
            if mid == 'UNKNOWN_MOUSE':
                continue
            mouse = self.data.loadmouse(mid)
            if self._mouse_passes_filter(mouse, paradigm, slug, stage,
                                         retired, properties):
                result.append(mid)
        return result


class Colony:
    """
    Unified handle to all acquired colony data.

    Wires up a single DataHandler and exposes both :class:`RetreiveSessions`
    and :class:`RetreiveMice` against it. Together they cover every mouse and
    every session currently on disk.

    A "focused" mouse can be set via :meth:`focus_mouse`; the loaded Mouse
    object is then available as :attr:`mouse`, and session retrieval methods
    on :attr:`sessions` operate against it.

    Attributes
    ----------
    data : DataHandler
        Shared file-level data handler.
    sessions : RetreiveSessions
        Session-history query interface.
    mice : RetreiveMice
        Mouse listing and filtering interface.
    mouse : Mouse or None
        The currently focused Mouse object (None until ``focus_mouse`` is called).
    """

    def __init__(self, config=None):
        """
        Parameters
        ----------
        config : Config, optional
            Explicit config to use.  A fresh Config is constructed if omitted.
        """
        self.data = DataHandler(config=config)
        self.sessions = RetreiveSessions(config=config, data=self.data)
        self.mice = RetreiveMice(config=config, data=self.data)

    @property
    def mouse(self):
        """The currently focused Mouse object, or None."""
        return self.sessions._active_mouse

    def focus_mouse(self, mouseid: str):
        """
        Load a mouse and set it as the focus of subsequent session retrieval.

        After calling this, ``colony.mouse`` is the loaded Mouse object and
        ``colony.sessions.retrieve_last(...)`` operates on its history.

        Parameters
        ----------
        mouseid : str
            The mouse ID to focus on.
        """
        self.sessions.set_mouse(mouseid)
```
