## Extended Data for "A Low-Cost, Modular Hardware and Software Platform for Head-Fixed Mouse Decision-Making Tasks": dataset.py

```
# -*- coding: utf-8 -*-
"""
Created on Wed Dec  3 18:26:06 2025

@author: mm4114
"""

from lampyr.primatives import Session
from lampyr import files
from typing import List, Union
from random import randint
from scipy.interpolate import interp1d
from dataclasses import asdict
from collections import namedtuple
import numpy as np
import os
import shutil


SegmentReference = namedtuple('SegmentReference', ['animal',
                                                   'session',
                                                   'segment'])
TraceReference = namedtuple('ExtractionToken', ['session',
                                                'timearray',
                                                'time_type'])
TraceExtractionProfile = namedtuple('ExtractionProfile', ['profile_name',
                                                          'mode',
                                                          'signal',
                                                          'samplerate',
                                                          'time_type',
                                                          'fill'])

def trace_extractor_factory(session, profile: TraceExtractionProfile):
    """
    Build a callable that extracts trace values from a session's rig data.

    For ``'nearest'`` and ``'linear'`` modes, returns a
    :func:`~scipy.interpolate.interp1d` object.  For windowed modes
    (``'mean'``, ``'sum'``, ``'count'``, ``'rate'``), returns a custom
    function that averages/sums/counts samples within each time window.

    Parameters
    ----------
    session : Session
        The session containing the rig data to extract from.
    profile : TraceExtractionProfile
        Specifies the signal channel, interpolation mode, sample rate, time
        type, and fill value.

    Returns
    -------
    callable
        A function ``f(time_array: np.ndarray) -> np.ndarray`` that returns
        extracted trace values at the requested times.
    """
    t_data = session.rigdata[profile.signal][profile.time_type]
    v_data = session.rigdata[profile.signal]['report_value']
    if profile.mode in ['nearest', 'linear']:
        return interp1d(t_data, v_data,
                        kind=profile.mode,
                        bounds_error=False,
                        fill_value=profile.fill)

    def windowed_extraction(time_array: np.ndarray):
        n = len(time_array)
        left_window = np.empty(n)
        right_window = np.empty(n)

        # edges
        left_window[0] = (time_array[1] - time_array[0]) / 2
        right_window[0] = left_window[0]
        left_window[-1] = (time_array[-1] - time_array[-2]) / 2
        right_window[-1] = left_window[-1]

        # interior points
        left_window[1:-1] = (time_array[1:-1] - time_array[:-2]) / 2
        right_window[1:-1] = (time_array[2:] - time_array[1:-1]) / 2

        out = np.full(n, profile.fill)
        for i, ti in enumerate(time_array):
            mask = (t_data >= ti - left_window[i]
                    ) & (t_data <= ti + right_window[i])
            if not mask.any():
                continue

            if profile.mode == "mean":
                out[i] = np.nanmean(v_data[mask])
            elif profile.mode == "sum":
                out[i] = np.nansum(v_data[mask])
            elif profile.mode == "count":
                out[i] = np.sum(~np.isnan(v_data[mask]))
            elif profile.mode == "rate":
                out[i] = np.sum(~np.isnan(v_data[mask])) / \
                    (left_window[i] + right_window[i])
            else:
                raise ValueError(f"Unknown mode: {profile.mode}")
        return out
    return windowed_extraction


class MultiSessionDataset:
    """
    A collection of sessions with disk persistence and cross-session search.

    Sessions are stored in a directory as pairs of ``.lampyr.h5`` (rig data)
    and ``.lampyr.json`` (metadata) files, indexed by
    ``msd_INDEX.lampyr.json``.

    Attributes
    ----------
    fp : str
        Path to the directory used for saving/loading session files.
    sessions : list of Session
        All sessions currently in the dataset.
    sessionsbyid : dict
        Maps ``uniquesessionid`` to Session objects.
    animals : list of str
        Sorted list of unique mouse IDs present in the dataset.
    sessionids : list of str
        List of session IDs in insertion order.
    animal_sessions : dict
        Maps mouse ID to a list of its sessions.
    """

    def __init__(self, fp: str = None, sessions: List[Session] = None,
                 destructive_overwrite=False, allow_mutation=False):
        """
        Initialise the dataset, loading existing sessions from disk.

        Parameters
        ----------
        fp : str
            Directory path for session file storage.  Created if it does not
            exist.
        sessions : list of Session, optional
            Sessions to add after loading from disk.
        destructive_overwrite : bool, optional
            If ``True``, delete all existing files in ``fp`` before adding
            new sessions.  Default is ``False``.
        """
        if sessions is None:
            sessions = []
        # Session tracking
        self._sessions_index_set = set()
        self._session_isloaded = {}
        self._session_objects = {}
        self.allow_mutation = allow_mutation

        # File attributes
        self.fp = fp
        if fp is not None:
            os.makedirs(fp, exist_ok=True)
        self._update()

        if fp is not None:
            if destructive_overwrite:
                self.clear_files()
            else:
                self._load_index()
        self.addsession(sessions)

        # Secondary attributes
        self._extractor_objects = {}
    
    def _load_session(self, sessionid):
        if sessionid not in self._sessions_index_set:
            raise ValueError('No such session is registered')
        if not self.persistent:
            raise RuntimeError('This is an in-memory only dataset, specify a filepath to load from disk')
        self._session_objects[sessionid] = files.loadsessionfile(sessionid, self.fp)
        self._session_isloaded[sessionid] = True
    
    def _save_session(self, sessionid):
        if not self.isloaded(sessionid):
            print(f'Skipping unloaded session file: {sessionid}')
            return
        if not self.persistent:
            raise RuntimeError('This is an in-memory only dataset, specify a filepath to save to disk')
        if not self.allow_mutation:
            raise RuntimeError('Mutation is not enabled')
        print(f'Saving session file: {sessionid}')
        files.savesessionfile(self._session_objects[sessionid], self.fp)
        print(f'Finished saving session file: {sessionid}')
    
    def _load_index(self):
        index_fp = os.path.join(self.fp, 'msd_INDEX.lampyr.json')
        if os.path.exists(index_fp):
            self._sessions_index_set.update(files.loadjson(index_fp))
        
    def _update(self):
        for s in self._sessions_index_set:
            if s not in self._session_isloaded:
                self._session_isloaded[s] = False
        for s in list(self._session_isloaded.keys()):
            if s not in self._sessions_index_set:
                self._session_isloaded.pop(s)
        for s in list(self._session_objects.keys()):
            if s not in self._sessions_index_set:
                self._session_objects.pop(s)
        self._extractor_objects = {}
    
    def clear(self):
        self._sessions_index_set = set()
        self._update()
    
    def save(self):
        if not self.persistent:
            raise RuntimeError('This is an in-memory only dataset, specify a filepath to save to disk')
        files.savejson(os.path.join(self.fp, 'msd_INDEX.lampyr.json'), self.sessionids)
        for sessionid in self.sessionids:
            self._save_session(sessionid)
    
    def clear_files(self):
        self.clear()
        if not self.persistent:
            return
        for item in os.listdir(self.fp):
            path = os.path.join(self.fp, item)
            if os.path.isfile(path) or os.path.islink(path):
                os.remove(path)
            elif os.path.isdir(path):
                shutil.rmtree(path)
        self.save()
        self._update()
    
    def addsession(self, session: Union[Session, List[Session]], _suppressupdate = False):
        """
        Add one or more sessions to the dataset, replacing duplicates by ID.

        Parameters
        ----------
        session : Session or list of Session
            Session(s) to add.
        _suppressupdate : bool, optional
            If ``True``, skip calling :meth:`update` (used internally for
            batch adds).  Default is ``False``.
        """
        if isinstance(session, list):
            for s in session:
                self.addsession(s, _suppressupdate=True)
            self._update()
            return
        if session.uniquesessionid in self._sessions_index_set:
            print(f'Skipped {session.uniquesessionid} because it is already in dataset')
            return
        self._sessions_index_set.add(session.uniquesessionid)
        self._session_objects[session.uniquesessionid] = session
        self._session_isloaded[session.uniquesessionid] = True
        if not _suppressupdate:
            self._update()
        
    @property
    def session_count(self):
        return len(self._sessions_index_set)
    
    @property
    def sessions(self):
        for sessionid in self.sessionids:
            if not self.isloaded(sessionid):
                self._load_session(sessionid)
            yield self._session_objects[sessionid]
    
    @property
    def animal_sessions(self):
        return {a: [s for s in self.sessions
                                    if s.mouseid == a]
                                for a in self.animals}
    
    @property
    def animals(self):
        return sorted(list(set([s.mouseid for s in self.sessions])))
    
    @property
    def sessionsbyid(self):
        return {s.uniquesessionid : s for s in self.sessions}
    
    @property
    def sessionids(self):
        return sorted(self._sessions_index_set)
        
    def isloaded(self, sessionid):
        return self._session_isloaded[sessionid]
    
    @property
    def persistent(self) -> bool:
        """True if the dataset has a backing directory on disk."""
        return self.fp is not None

    def search(self,
               *args,
               mouseid: str = None,
               return_objects : bool = False,
               **kwargs):
        """
        Search segments across all sessions.

        Parameters
        ----------
        *args
            Positional arguments forwarded to :meth:`Session.search`.
        mouseid : str, optional
            Restrict search to sessions from this mouse.
        return_objects : bool, optional
            If ``True``, return segment dicts instead of
            :class:`SegmentReference` named tuples.
        **kwargs
            Keyword arguments forwarded to :meth:`Session.search`.

        Returns
        -------
        list of SegmentReference or list of dict
            Matching segments across all sessions.
        """
        seglist = []
        for session in self.sessions:
            if mouseid is not None and mouseid != session.mouseid:
                continue
            segments = session.search(*args,
                                      return_objects=return_objects,
                                      **kwargs)
            if not return_objects:
                for seg in segments:
                    seglist.append(
                        SegmentReference(animal=session.mouseid,
                                         session=session.uniquesessionid,
                                         segment=seg)
                    )
            else:
                seglist += segments
        return seglist

    def get_segment(self, reference: SegmentReference):
        """
        Retrieve a segment dict by reference.

        Parameters
        ----------
        reference : SegmentReference
            Named tuple identifying the animal, session, and segment.

        Returns
        -------
        dict
            The serialised segment data dict from the session.
        """
        return self.sessionsbyid[reference.session].segments[reference.segment]

    def get_trace(self, reference: TraceReference,
                  profile: TraceExtractionProfile):
        """
        Extract and return a trace array for the given reference and profile.

        Caches the extractor object per ``(session, profile)`` pair.

        Parameters
        ----------
        reference : TraceReference
            Identifies the session and the time array to evaluate.
        profile : TraceExtractionProfile
            Specifies signal, mode, sample rate, time type, and fill value.

        Returns
        -------
        np.ndarray
            Extracted trace values at ``reference.timearray`` times.
        """
        if (reference.session, profile) not in self._extractor_objects:
            extractor = trace_extractor_factory(
                self.sessionsbyid[reference.session],
                profile)
            self._extractor_objects[(reference.session, profile)] = extractor
        else:
            extractor = self._extractor_objects[(reference.session, profile)]
        return extractor(reference.timearray)
```
