## Extended Data for "A Low-Cost, Modular Hardware and Software Platform for Head-Fixed Mouse Decision-Making Tasks": longtidy.py

```
# -*- coding: utf-8 -*-
"""
Created on Tue Jan  6 17:18:16 2026

@author: mm4114
"""
import inspect, hashlib, json, pickle
import pandas as pd
import numpy as np
from pathlib import Path

from lampyr.analysis.dataset import SegmentReference, TraceExtractionProfile
from lampyr.analysis.traces import dynamic_trace_extraction

from typing import Iterable, List, Callable, Dict

def _baseline_extraction_kwargs(baseline_range, n_events):
    """
    Auto-derive padding and pseudotime kwargs to extract a baseline window.

    Given a window like ``[-2, 0]`` and one or more anchor events, return the
    minimal padding and pseudotime arguments needed to extract that window
    around the first anchor (which is placed at pseudotime ``0``).
    """
    pad_start = max(0.0, -baseline_range[0])
    pad_end = max(0.0, baseline_range[1])
    event_pt = [-pad_start, 0] + [None] * (n_events - 1) + [pad_end]
    return {'padding': True,
            'pad_start': pad_start,
            'pad_end': pad_end,
            'event_pseudotimes': event_pt}


def longtidy_multidynamictraceextraction(
        data,
        segments: Iterable[SegmentReference],
        events: Iterable[str],
        extraction_profiles: Iterable[TraceExtractionProfile],
        expose_reports: List[str] = None,
        expose_properties: List[str] = None,
        expose_segmentattr: List[str] = None,
        custom_exposures: Dict[str, Callable] = None,
        perprofilekwargs: dict = None,
        baseline_range=None,
        baseline_events=None,
        baseline_kwargs: dict = None,
        **kwargs):
    """
    Extract multiple signal traces across segments and return long-form DataFrames.

    Runs :func:`~lampyr.analysis.traces.dynamic_trace_extraction` for each
    extraction profile, concatenates results into a single long-tidy trace
    DataFrame, and joins segment metadata.

    Parameters
    ----------
    data : MultiSessionDataset
        Dataset to extract from.
    segments : iterable of SegmentReference
        Segments to process.
    events : iterable of str
        Event names defining time landmarks (shared across all profiles).
    extraction_profiles : iterable of TraceExtractionProfile
        One or more extraction profiles to run.
    expose_reports : list of str, optional
        Report field names to include as columns in ``segment_info``.
    expose_properties : list of str, optional
        Segment property names to include as columns in ``segment_info``.
    expose_segmentattr : list of str, optional
        Arbitrary segment attribute names to include as columns.
    custom_exposures : dict of str -> callable, optional
        ``{column_name: func(segment_dict) -> value}`` for computed columns.
    perprofilekwargs : dict, optional
        ``{profile_name: {kwarg: value}}`` to pass additional keyword arguments
        to :func:`~lampyr.analysis.traces.dynamic_trace_extraction` for
        specific profiles only.  ``baseline_range: None`` here disables
        baselining for that profile only.
    baseline_range : tuple of (float, float), optional
        Time window over which to compute the baseline mean to subtract from
        each trace.  Interpretation depends on ``baseline_events``: if it is
        ``None``, the window is in the same alignment as ``events``; otherwise
        it is in the alignment defined by ``baseline_events`` (with the first
        baseline event at pseudotime ``0``).
    baseline_events : iterable of str, optional
        If given, perform a separate extraction aligned to these events for
        baseline computation.  Padding and pseudotimes are auto-derived from
        ``baseline_range`` so the window fits exactly.
    baseline_kwargs : dict, optional
        Override or extend the auto-derived baseline-extraction kwargs (e.g.
        force a specific ``samples`` count).  Rarely needed.
    **kwargs
        Additional keyword arguments forwarded to
        :func:`~lampyr.analysis.traces.dynamic_trace_extraction` for all
        profiles.

    Returns
    -------
    traceframe : pd.DataFrame
        Long-tidy DataFrame with columns ``'time'``, ``'value'``, ``'signal'``,
        ``'segkey'``, plus any exposed metadata columns.
    segment_info : pd.DataFrame
        One-row-per-segment metadata DataFrame with ``'segkey'``, ``'animal'``,
        ``'session'``, ``'segment'``, plus any exposed columns.
    """
    def seg2segkey(s): return f'{s.animal}_{s.session}_{s.segment}'
    exposures = (['segkey', 'animal', 'session', 'segment'] +
                 (expose_reports or []) +
                 (expose_properties or []) +
                 (expose_segmentattr or []) +
                 list(custom_exposures or {}))
    # Construct segment info dataframe
    segment_dict = {e: [] for e in exposures}
    expose_reports = expose_reports or []
    expose_properties = expose_properties or []
    expose_segmentattr = expose_segmentattr or []
    custom_exposures = custom_exposures or {}
    for segref in segments:
        segment_dict['segkey'].append(seg2segkey(segref))
        segment_dict['animal'].append(segref.animal)
        segment_dict['session'].append(segref.session)
        segment_dict['segment'].append(segref.segment)
        seg_obj = data.get_segment(segref)
        for report in expose_reports:
            segment_dict[report].append(seg_obj['reports'][report])
        for prop in expose_properties:
            segment_dict[prop].append(seg_obj['properties'][prop])
        for attr in expose_segmentattr:
            segment_dict[attr].append(seg_obj[attr])
        for expname, expcallable in custom_exposures.items():
            segment_dict[expname].append(expcallable(seg_obj))
    segment_info = pd.DataFrame(segment_dict)
    segment_info['segkey'] = segment_info['segkey'].astype('string')
    segment_info['animal'] = segment_info['animal'].astype('category')
    segment_info['session'] = segment_info['session'].astype('category')
    segment_info['segment'] = segment_info['segment'].astype('string')

    profile_dfs = []
    for profile in extraction_profiles:
        if (perprofilekwargs is not None and
                profile.profile_name in perprofilekwargs):
            profilekwargs = {**kwargs, **perprofilekwargs[profile.profile_name]}
        else:
            profilekwargs = kwargs.copy()

        # Per-profile baseline_range overrides the function-level value
        profile_baseline_range = profilekwargs.pop('baseline_range',
                                                   baseline_range)

        if baseline_events is None:
            t, sig, _ = dynamic_trace_extraction(
                data, segments=segments, events=events,
                extraction_profile=profile,
                baseline_range=profile_baseline_range,
                **profilekwargs)
        else:
            t, sig, _ = dynamic_trace_extraction(
                data, segments=segments, events=events,
                extraction_profile=profile,
                **profilekwargs)
            if profile_baseline_range is not None:
                bl_kw = _baseline_extraction_kwargs(profile_baseline_range,
                                                    len(baseline_events))
                bl_kw.update(baseline_kwargs or {})
                bl_kw.pop('baseline_range', None)
                t_bl, sig_bl, _ = dynamic_trace_extraction(
                    data, segments=segments, events=baseline_events,
                    extraction_profile=profile, **bl_kw)
                bl_mask = ((t_bl >= profile_baseline_range[0]) &
                           (t_bl <= profile_baseline_range[1]))
                baseline_means = np.nanmean(sig_bl[:, bl_mask], axis=1)
                sig = sig - baseline_means[:, np.newaxis]
        n_segs, len_t = sig.shape
        t_long = np.tile(t, n_segs)
        sig_long = sig.flatten()
        seg_keys = np.repeat(segment_dict['segkey'], len_t)
        profile_df = pd.DataFrame({'time': t_long,
                                  'value': sig_long,
                                   'signal': profile.profile_name,
                                   'segkey': seg_keys})
        profile_dfs.append(profile_df)
    traceframe = pd.concat(profile_dfs, ignore_index=True)
    traceframe = traceframe.merge(segment_info, on='segkey', how='left')
    return traceframe, segment_info

def load_parquet(path: str | Path) -> pd.DataFrame | None:
    """
    Load a Parquet file into a DataFrame, returning ``None`` if it does not exist.

    Parameters
    ----------
    path : str or Path
        Path to the ``.parquet`` file.

    Returns
    -------
    pd.DataFrame or None
        Loaded DataFrame, or ``None`` if the file is not found.
    """
    path = Path(path)
    if not path.exists():
        return None
    return pd.read_parquet(path, engine="pyarrow")


def save_parquet(path: str | Path, df: pd.DataFrame, overwrite: bool = False):
    """
    Save a DataFrame to Parquet format using Snappy compression.

    Parameters
    ----------
    path : str or Path
        Destination path.  Parent directories are created if needed.
    df : pd.DataFrame
        DataFrame to save.
    overwrite : bool, optional
        If ``False`` (default), raises ``FileExistsError`` if the file already
        exists.

    Raises
    ------
    FileExistsError
        If ``path`` exists and ``overwrite`` is ``False``.
    """
    path = Path(path)
    path.parent.mkdir(parents=True, exist_ok=True)

    if path.exists() and not overwrite:
        raise FileExistsError(path)

    df.to_parquet(path, engine="pyarrow", compression="snappy")


def ltdataset(cachefile):
    cachefile = Path(cachefile)
    cachefile.parent.mkdir(parents=True, exist_ok=True)

    def decorator(func):
        def wrapper(*args, force_reload=False, **kwargs):
            # Load metadata
            if cachefile.exists():
                with open(cachefile, "r") as f:
                    meta = json.load(f)
            else:
                meta = {}

            fid = func.__name__
            src_uid = hashlib.sha256(inspect.getsource(func).encode()).hexdigest()

            # Hash of args + kwargs
            try:
                blob = pickle.dumps((args, sorted(kwargs.items())))
            except Exception:
                blob = repr((args, sorted(kwargs.items()))).encode()
            args_uid = hashlib.sha256(blob).hexdigest()

            entry = meta.get(fid, {"src_uid": src_uid, "entries": {}})

            # Source changed → wipe ALL cached parquets for this function
            if entry.get("src_uid") != src_uid:
                for sub in entry.get("entries", {}).values():
                    for fname in (sub.get("uid_main", ""), sub.get("uid_meta", "")):
                        p = cachefile.parent / fname
                        if p.exists():
                            p.unlink()
                entry = {"src_uid": src_uid, "entries": {}}

            entries = entry["entries"]

            # Cache hit for these specific args
            if not force_reload and args_uid in entries:
                sub = entries[args_uid]
                main_path = cachefile.parent / sub["uid_main"]
                meta_path = cachefile.parent / sub["uid_meta"]
                if main_path.exists() and meta_path.exists():
                    df_main = load_parquet(main_path)
                    df_meta = load_parquet(meta_path)
                    if df_main is not None and df_meta is not None:
                        return df_main, df_meta

            # force_reload → delete the existing parquet for this args entry first
            if force_reload and args_uid in entries:
                sub = entries[args_uid]
                for fname in (sub.get("uid_main", ""), sub.get("uid_meta", "")):
                    p = cachefile.parent / fname
                    if p.exists():
                        p.unlink()

            # Run and save
            uid_main = f"{fid}_{src_uid[:16]}_{args_uid[:16]}_main.parquet"
            uid_meta = f"{fid}_{src_uid[:16]}_{args_uid[:16]}_meta.parquet"
            main_path = cachefile.parent / uid_main
            meta_path = cachefile.parent / uid_meta

            df_main, df_meta = func(*args, **kwargs)
            save_parquet(main_path, df_main, overwrite=True)
            save_parquet(meta_path, df_meta, overwrite=True)

            entries[args_uid] = {"uid_main": uid_main, "uid_meta": uid_meta}
            entry["src_uid"] = src_uid
            entry["entries"] = entries
            meta[fid] = entry

            with open(cachefile, "w") as f:
                json.dump(meta, f, indent=2)

            return df_main, df_meta
        return wrapper
    return decorator
```
