## Extended Data for "A Low-Cost, Modular Hardware and Software Platform for Head-Fixed Mouse Decision-Making Tasks": records.py

```
# -*- coding: utf-8 -*-
"""Session plotting helpers."""

from abc import ABC, abstractmethod

import matplotlib.pyplot as plt
import sqlite3


class AbstractPlot(ABC):
    extractors = ()

    def __init__(self, query, record_store, **settings):
        self.query = query
        self.record_store = record_store
        self.settings = settings

    def sessions(self):
        return self.query.objects()
    
    def preload(self):
        for extractor in self.extractors:
            self.record_store.preload(
                extractor,
                self.query,
            )

    def records(self, extractor=None):
        self.preload()

        if extractor is not None:
            extractors = (extractor,)
        elif len(self.extractors) == 1:
            extractors = (self.extractors[0],)
        else:
            extractors = tuple(self.extractors)

        grouped_records = {
            extractor_name: []
            for extractor_name in extractors
        }

        for sessionid in self.query.ids():
            sessionid = str(sessionid)
            for extractor_name in extractors:
                grouped_records[extractor_name].extend(
                    self.record_store.cache[
                        (extractor_name, sessionid)
                    ]
                )

        if len(extractors) == 1:
            return grouped_records[extractors[0]]

        return grouped_records
    
    def groupby(self, key):
        if len(self.extractors) != 1:
            raise ValueError(
                "groupby() requires a single extractor plot "
                "or an explicit records(extractor=...) call."
            )

        groups = {}

        for record in self.records():
            group_key = key(record) if callable(key) else record[key]
            groups.setdefault(group_key, []).append(record)

        return groups

    @abstractmethod
    def plot(self, ax=None):
        if ax is None:
            _, ax = plt.subplots()
        return ax


class RecordExtractor(ABC):
    name = None
    version = 1

    @abstractmethod
    def build(self, session):
        pass


class RecordStore:
    def __init__(
        self,
        db_file="reports.db",
        verbose=False,
    ):
        self.extractors = {}
        self.cache = {}
        self.verbose = verbose

        self.db_file = db_file

        self.conn = sqlite3.connect(
            self.db_file
        )
        self.conn.row_factory = sqlite3.Row

        self._init_metadata_table()
        self._init_built_sessions_table()

    def register(self, extractor):
        if extractor.name is None:
            raise ValueError(
                "Extractor must define a name."
            )

        self.extractors[extractor.name] = extractor

        current_version = self._extractor_version(
            extractor.name
        )

        if current_version != extractor.version:
            if self.verbose:
                print(
                    f"Rebuilding '{extractor.name}' "
                    f"({current_version} -> "
                    f"{extractor.version})"
                )

            self.conn.execute(
                f"DROP TABLE IF EXISTS {extractor.name}"
            )
            self._invalidate_extractor(
                extractor.name
            )

            self.conn.execute(
                """
                INSERT OR REPLACE INTO
                extractor_metadata
                (extractor, version)
                VALUES (?, ?)
                """,
                (
                    extractor.name,
                    extractor.version,
                ),
            )

            self.conn.commit()

    def get(self, extractor_name, session):
        if extractor_name not in self.extractors:
            raise KeyError(
                f"Unknown extractor: "
                f"{extractor_name}"
            )

        sessionid = self._sessionid(session)

        key = (
            extractor_name,
            sessionid,
        )

        if key not in self.cache:
            records = self._load_records(
                extractor_name,
                sessionid,
            )

            if records is None:
                if self._session_built(
                    extractor_name,
                    sessionid,
                ):
                    self.cache[key] = []
                else:
                    self._build_session(
                        extractor_name,
                        session,
                    )
            else:
                self.cache[key] = records

        return self.cache[key]

    def records(self, extractor_name):
        for (name, _), records in self.cache.items():
            if name == extractor_name:
                yield from records

    def preload(self, extractor_name, query):
        for mouseid, entry in query._matched_entries():
            sessionid = str(
                query.colony._sessionid(entry)
            )

            key = (
                extractor_name,
                sessionid,
            )

            if key in self.cache:
                continue

            records = self._load_records(
                extractor_name,
                sessionid,
            )

            if records is not None:
                self.cache[key] = records
                continue

            if self._session_built(
                extractor_name,
                sessionid,
            ):
                self.cache[key] = []
                continue

            session = query.colony.load_session(
                sessionid,
                mouseid,
            )

            if session is None:
                self.cache[key] = []
                continue

            self.get(
                extractor_name,
                session,
            )

    def clear_cache(self):
        self.cache.clear()

    def _build_session(
        self,
        extractor_name,
        session,
    ):
        extractor = self.extractors[
            extractor_name
        ]

        sessionid = self._sessionid(
            session
        )

        if self.verbose:
            print(
                f"Building '{extractor_name}' "
                f"for session {sessionid}"
            )

        built = extractor.build(session)

        if isinstance(built, dict):
            built = [built]
        else:
            built = list(built)

        for record in built:
            record["sessionid"] = sessionid
            record["mouseid"] = getattr(
                session,
                "mouseid",
                None,
            )

        self.cache[
            (
                extractor_name,
                sessionid,
            )
        ] = built

        self._mark_session_built(
            extractor_name,
            sessionid,
        )

        self._save_records(
            extractor_name,
            built,
        )

    def _init_metadata_table(self):
        self.conn.execute(
            """
            CREATE TABLE IF NOT EXISTS
            extractor_metadata (
                extractor TEXT PRIMARY KEY,
                version INTEGER NOT NULL
            )
            """
        )

        self.conn.commit()

    def _init_built_sessions_table(self):
        self.conn.execute(
            """
            CREATE TABLE IF NOT EXISTS
            extractor_sessions (
                extractor TEXT NOT NULL,
                sessionid TEXT NOT NULL,
                PRIMARY KEY (
                    extractor,
                    sessionid
                )
            )
            """
        )

        self.conn.commit()

    def _extractor_version(
        self,
        extractor_name,
    ):
        row = self.conn.execute(
            """
            SELECT version
            FROM extractor_metadata
            WHERE extractor = ?
            """,
            (extractor_name,),
        ).fetchone()

        return (
            row["version"]
            if row
            else None
        )
    
    def _invalidate_extractor(
        self,
        extractor_name,
    ):
        keys = [
            key
            for key in self.cache
            if key[0] == extractor_name
        ]
    
        for key in keys:
            del self.cache[key]

        self.conn.execute(
            """
            DELETE FROM extractor_sessions
            WHERE extractor = ?
            """,
            (extractor_name,),
        )
        self.conn.commit()

    def _session_built(
        self,
        extractor_name,
        sessionid,
    ):
        row = self.conn.execute(
            """
            SELECT 1
            FROM extractor_sessions
            WHERE extractor = ?
            AND sessionid = ?
            """,
            (
                extractor_name,
                str(sessionid),
            ),
        ).fetchone()

        return row is not None

    def _mark_session_built(
        self,
        extractor_name,
        sessionid,
    ):
        self.conn.execute(
            """
            INSERT OR REPLACE INTO
            extractor_sessions
            (extractor, sessionid)
            VALUES (?, ?)
            """,
            (
                extractor_name,
                str(sessionid),
            ),
        )
        self.conn.commit()

    @staticmethod
    def _sql_type(value):
        if value is None:
            return "REAL"
        
        if isinstance(value, bool):
            return "INTEGER"

        if isinstance(value, int):
            return "INTEGER"

        if isinstance(value, float):
            return "REAL"

        return "TEXT"

    def _ensure_table(
        self,
        extractor_name,
        records,
    ):
        exists = self.conn.execute(
            """
            SELECT name
            FROM sqlite_master
            WHERE type='table'
            AND name=?
            """,
            (extractor_name,),
        ).fetchone()

        if exists:
            return

        first = records[0]

        columns = []

        for key, value in first.items():
            columns.append(
                f"{key} "
                f"{self._sql_type(value)}"
            )

        columns.append(
            "record_index INTEGER"
        )

        sql = f"""
        CREATE TABLE
        {extractor_name}
        (
            {",".join(columns)},
            PRIMARY KEY (
                sessionid,
                record_index
            )
        )
        """

        self.conn.execute(sql)
        self.conn.commit()

    def _save_records(
        self,
        extractor_name,
        records,
    ):
        if not records:
            return

        self._ensure_table(
            extractor_name,
            records,
        )

        columns = list(
            records[0].keys()
        )

        placeholders = ",".join(
            "?"
            for _ in columns
        )

        sql = f"""
        INSERT OR REPLACE INTO
        {extractor_name}
        (
            {",".join(columns)},
            record_index
        )
        VALUES
        (
            {placeholders},
            ?
        )
        """

        for index, record in enumerate(records):
            values = [
                record[col]
                for col in columns
            ]

            self.conn.execute(
                sql,
                (*values, index),
            )

        self.conn.commit()

    def _load_records(
        self,
        extractor_name,
        sessionid,
    ):
        exists = self.conn.execute(
            """
            SELECT name
            FROM sqlite_master
            WHERE type='table'
            AND name=?
            """,
            (extractor_name,),
        ).fetchone()

        if not exists:
            return None

        rows = self.conn.execute(
            f"""
            SELECT *
            FROM {extractor_name}
            WHERE sessionid = ?
            ORDER BY record_index
            """,
            (sessionid,),
        ).fetchall()

        if not rows:
            return None

        records = []

        for row in rows:
            record = dict(row)

            record.pop(
                "record_index",
                None,
            )

            records.append(record)

        return records

    @staticmethod
    def _sessionid(session):
        for attr in (
            "sessionid",
            "uniquesessionid",
            "id",
        ):
            value = getattr(
                session,
                attr,
                None,
            )

            if value not in (
                None,
                "",
            ):
                return str(value)

        raise AttributeError(
            "Could not determine session id."
        )

    def close(self):
        if self.conn is not None:
            self.conn.close()
            self.conn = None


class BasicSessionRecord(RecordExtractor):
    name = "basic_session"
    version = 2

    def build(self, session):
        return {
            "merit": getattr(session, "merit", None),
            "demerit": getattr(session, "demerit", None),
            "duration": getattr(session, "duration", None),
            "trial": getattr(session, "trial", None),
            "reward": getattr(session, "reward", None),
            "reward_amount": getattr(session, "reward_amount", None),
            "abstention": getattr(session, "abstention", None),
            "participation": getattr(session, "participation", None),
            "starttime": getattr(session, "starttime", None),
        }


class ParticipationByMousePlot(AbstractPlot):
    extractor_name = "basic_session"

    def plot(self, ax=None):
        ax = super().plot(ax=ax)
        groups = self.groupby("mouseid")
        labels = []
        means = []

        for mouseid, records in groups.items():
            labels.append(mouseid)
            means.append(
                sum(record["participation"]
                    for record in records) / len(records)
            )

        ax.bar(labels, means)
        ax.set_ylabel("Mean participation")
        ax.set_xlabel("Mouse")
        ax.set_title("Participation by Mouse")
        ax.tick_params(axis="x", rotation=45)
        return ax
```
