## Extended Data for "A Low-Cost, Modular Hardware and Software Platform for Head-Fixed Mouse Decision-Making Tasks": segments.py

```
# -*- coding: utf-8 -*-
"""
Created on Fri Jan 23 14:26:35 2026

@author: mm4114
"""
import numpy as np

from lampyr.analysis.dataset import MultiSessionDataset, SegmentReference

from typing import Iterable, List

def extract_event_times(data: MultiSessionDataset,
                        segment: SegmentReference,
                        events: Iterable[str]) -> np.ndarray:
    """
    Extract event timestamps from a segment, aligned to a specified event order.

    Parameters
    ----------
    data : MultiSessionDataset
        Dataset containing session and segment data with event records.
    segment : SegmentReference
        Reference identifying the segment from which to extract events.
    events : Iterable[str]
        Ordered list of event names to extract. If an event appears multiple
        times in this list, multiple occurrences of that event will be filled
        in order of appearance in the segment.

    Returns
    -------
    evtimes : np.ndarray
        1D array of event times aligned to `events`. Entries are NaN when the
        corresponding event occurrence is not found in the segment.
    """
    segevents = data.get_segment(segment)['event_records']
    segevents = sorted(segevents, key=lambda x: x['time'])
    evtimes = np.full((len(events)), np.nan)
    slots = {}
    for i, ev in enumerate(events):
        slots.setdefault(ev, []).append(i)
    counters = {ev: 0 for ev in slots}
    for e in segevents:
        name = e['event']
        if name not in slots:
            continue
        c = counters[name]
        if c >= len(slots[name]):
            continue
        idx = slots[name][c]
        evtimes[idx] = e['time']
        counters[name] += 1
    return evtimes


def extract_event_times_multiple(data: MultiSessionDataset,
                                 segments: List[SegmentReference],
                                 events: List[str]) -> np.ndarray:
    """
    Extract event timestamps across multiple segments.

    Parameters
    ----------
    data : MultiSessionDataset
        Dataset containing session and segment data with event records.
    segments : List[SegmentReference]
        List of segment references from which to extract events.
    events : List[str]
        Ordered list of event names to extract for each segment.

    Returns
    -------
    event_times : np.ndarray
        2D array of shape (n_segments, n_events) containing event times.
        Each row corresponds to a segment, and each column corresponds to
        an event in `events`. Missing events are represented as NaN.
    """
    event_times = np.full((len(segments), len(events)), np.nan)
    for ind, seg in enumerate(segments):
        event_times[ind, :] = extract_event_times(data, seg, events)
    return event_times
```
