## Extended Data for "A Low-Cost, Modular Hardware and Software Platform for Head-Fixed Mouse Decision-Making Tasks": time.py

```
# -*- coding: utf-8 -*-
"""
Created on Fri Jan 23 14:26:35 2026

@author: mm4114
"""
import time

import numpy as np

from scipy.interpolate import interp1d

from lampyr.primatives import Session

from typing import Iterable, Union

def createtimereporter():
    """
    Create a closure that prints elapsed time since the last call.

    Each call to the returned ``report(prefix)`` function prints
    ``'<prefix> (<elapsed> sec)'`` and resets the internal timer.

    Returns
    -------
    callable
        A ``report(prefix: str)`` function.
    """
    start = time.time()

    def report(prefix):
        nonlocal start
        now = time.time()
        dur = now - start
        print(f'{prefix} ({dur:.1f} sec)')
        start = now
    return report

def create_dynamictimearray(points: Iterable[Union[float, int]],
                            samples: Iterable[int]):
    """
    Build a time array by linearly interpolating between anchor points.

    Creates ``samples[i]`` evenly spaced values (endpoint excluded) between
    ``points[i]`` and ``points[i+1]`` for each segment, then appends the
    final anchor point.

    Parameters
    ----------
    points : iterable of float or int
        Anchor time values.  Must have ``len(samples) + 1`` elements.
    samples : iterable of int
        Number of samples to generate in each interval.

    Returns
    -------
    np.ndarray
        1D array of length ``sum(samples) + 1``.

    Raises
    ------
    ValueError
        If ``len(points) - 1 != len(samples)``.
    """
    points = np.array(points, dtype=float)
    samples = np.array(samples, dtype=int)
    if len(points) - 1 != len(samples):
        raise ValueError("samples must have one less element than points")
    stretched = []
    for i in range(len(samples)):
        # Create linearly spaced points between consecutive points
        seg = np.linspace(points[i], points[i+1], samples[i], endpoint=False)
        stretched.append(seg)

    # Append the last point
    stretched.append([points[-1]])
    return np.concatenate(stretched)

def create_dynamictimearray_2d(points_2d: np.ndarray,
                               samples: Iterable[int]):
    """
    Apply :func:`create_dynamictimearray` to each row of a 2D array.

    Parameters
    ----------
    points_2d : np.ndarray
        2D array of shape ``(n_rows, n_points)`` where each row is a set of
        anchor times.
    samples : iterable of int
        Sample counts per interval, shared across all rows.

    Returns
    -------
    np.ndarray
        2D array of shape ``(n_rows, sum(samples) + 1)``.
    """
    newdim = sum(samples) + 1
    newarray = np.empty((points_2d.shape[0], newdim))
    for r, row  in enumerate(points_2d):
        newarray[r,:] = create_dynamictimearray(row, samples)
    return newarray

def create_syncstream(session: Session, components: list,
                      roottime: str, contribution_cutoff: int = 200):
    """
    Build a SyncStream in session.rigdata from the given rig data components.

    Samples up to ``contribution_cutoff`` points from each component channel,
    concatenates them, and sorts by ``roottime`` so the resulting arrays are
    monotonically increasing in the root time base.
    """
    timeentries = set().union(*[session.rigdata[c] for c in components])
    timeentries = [t for t in timeentries if t != 'report_value']
    session.rigdata['SyncStream'] = {k: np.empty(0, dtype=float)
                                     for k in timeentries}
    for c in components:
        arrlen = len(list(session.rigdata[c].values())[0])
        newentries = {
            k: (np.full(arrlen, np.nan) if k not in session.rigdata[c]
                else session.rigdata[c][k])
            for k in timeentries
        }
        for t, v in newentries.items():
            v_pared = v[::len(v) // contribution_cutoff] if len(v) > contribution_cutoff else v
            session.rigdata['SyncStream'][t] = np.concatenate(
                (session.rigdata['SyncStream'][t], v_pared))
    idx = np.argsort(session.rigdata['SyncStream'][roottime])
    for t in timeentries:
        session.rigdata['SyncStream'][t] = session.rigdata['SyncStream'][t][idx]


class TimeTranslator:
    """
    Interpolation-based converter between different time bases in a session.

    Uses the session's ``SyncStream`` rig data to build interpolators between
    named time axes (e.g. ``'unix_time'`` and ``'arduino_time'``).

    Attributes
    ----------
    default_source : str or None
        Default source time axis name.
    default_target : str or None
        Default target time axis name.
    """

    def __init__(self, session: Session,
                 target=None,
                 source=None,
                 kind='linear',
                 fill_value='extrapolate'):
        """
        Initialise from a session's SyncStream data.

        Parameters
        ----------
        session : Session
            Session whose ``rigdata['SyncStream']`` provides the sync data.
        target : str, optional
            Default target time axis (e.g. ``'arduino_time'``).
        source : str, optional
            Default source time axis (e.g. ``'unix_time'``).
        kind : str, optional
            Interpolation kind passed to :func:`scipy.interpolate.interp1d`.
            Default is ``'linear'``.
        fill_value : str or float, optional
            Fill behaviour for out-of-range values.  Default is
            ``'extrapolate'``.
        """
        self.default_source = source
        self.default_target = target
        self._kind = kind
        self._fill_value = fill_value
        self._syncstream = session.rigdata['SyncStream'].copy()
        self._translators = {}

    def _convert(self, arg, source, target):
        """
        Internal conversion; builds and caches the interpolator on first use.

        Parameters
        ----------
        arg : array-like
            Values in the source time base to convert.
        source : str
            Source time axis key in ``SyncStream``.
        target : str
            Target time axis key in ``SyncStream``.

        Returns
        -------
        np.ndarray
            Values in the target time base.
        """
        if (source, target) not in self._translators:
            self._translators[(source, target)] = interp1d(
                self._syncstream[source],
                self._syncstream[target],
                kind=self._kind,
                fill_value=self._fill_value)
        return self._translators[(source, target)](arg)

    def convert(self, value, source=None, target=None):
        """
        Convert ``value`` from the source time base to the target time base.

        Parameters
        ----------
        value : array-like
            Time values to convert.
        source : str, optional
            Source time axis.  Falls back to ``default_source`` if ``None``.
        target : str, optional
            Target time axis.  Falls back to ``default_target`` if ``None``.

        Returns
        -------
        np.ndarray
            Converted time values.

        Raises
        ------
        ValueError
            If neither explicit nor default source/target are set.
        """
        source = source or self.default_source
        target = target or self.default_target
        if source is None or target is None:
            raise ValueError('Source or Target not specified')
        return self._convert(value, source, target)
```
