## Extended Data for "A Low-Cost, Modular Hardware and Software Platform for Head-Fixed Mouse Decision-Making Tasks": traces.py

```
# -*- coding: utf-8 -*-
"""
Created on Fri Jan 23 14:26:35 2026

@author: mm4114
"""
import numpy as np

from scipy.interpolate import interp1d

from lampyr.primatives import Session
from lampyr.analysis.dataset import SegmentReference, TraceReference, MultiSessionDataset, TraceExtractionProfile
from lampyr.analysis.time import TimeTranslator, create_dynamictimearray, create_dynamictimearray_2d
from lampyr.analysis.segments import extract_event_times_multiple

from typing import Iterable, Literal, Union, List, Tuple


def determine_ideal_samplerate(event_times,
                               extraction_profile: TraceExtractionProfile,
                               mode: Literal['mean', 'median'] = 'mean'):
    """
    Compute the ideal number of samples per inter-event interval.

    Parameters
    ----------
    event_times : np.ndarray
        2D array of shape ``(n_segments, n_events)`` containing event timestamps.
    extraction_profile : TraceExtractionProfile
        Provides the desired sample rate (samples per second).
    mode : {'mean', 'median'}, optional
        Whether to use mean or median inter-event duration when averaging
        across segments.  Default is ``'mean'``.

    Returns
    -------
    list of int
        One integer per inter-event interval giving the ideal sample count.
        NaN intervals are replaced with 1.
    """
    diffs = np.diff(event_times, axis=1)
    if mode == 'mean':
        elapsed = diffs.mean(axis=0)
    elif mode == 'median':
        elapsed = np.median(diffs, axis=0)

    samples = extraction_profile.samplerate * elapsed
    ideal_samplerate = [int(round(s))
                        if not np.isnan(s) else 1
                        for s in samples]
    return ideal_samplerate


def timeconversion_bulk(data: MultiSessionDataset,
                        sessionids: Iterable[str],
                        event_times,
                        source_time,
                        target_time):
    """
    Convert a 2D array of event times from one time base to another across sessions.

    Parameters
    ----------
    data : MultiSessionDataset
        Dataset providing access to session rig data for time conversion.
    sessionids : iterable of str
        Session IDs corresponding to each row of ``event_times``.
    event_times : np.ndarray
        2D array of shape ``(n_segments, n_events)`` in ``source_time`` base.
    source_time : str
        Source time axis key (e.g. ``'unix_time'``).
    target_time : str
        Target time axis key (e.g. ``'arduino_time'``).

    Returns
    -------
    np.ndarray
        2D array of the same shape as ``event_times`` in ``target_time`` base.
    """
    converted_times = np.empty(event_times.shape)
    timeconverters = {}
    for ind, sessionid in enumerate(sessionids):
        if sessionid not in timeconverters:
            translator = TimeTranslator(data.sessionsbyid[sessionid],
                                        target=target_time,
                                        source=source_time,
                                        kind='linear',
                                        fill_value='extrapolate')
            timeconverters[sessionid] = translator
        nt = timeconverters[sessionid].convert(event_times[ind, :])
        converted_times[ind, :] = nt

    return converted_times


def dynamic_trace_extraction(data,
                             segments: Iterable[SegmentReference],
                             events: Iterable[str],
                             extraction_profile: TraceExtractionProfile,
                             samples: List[Union[int, None]] = None,
                             event_pseudotimes: List[Union[int, None]] = None,
                             baseline_range: Tuple[float, float] = None,
                             padding=False,
                             pad_start=2,
                             pad_end=6):
    """
    Extract aligned trace arrays across multiple segments using dynamic time warping.

    Extracts event timestamps, optionally pads them, converts to the target
    time base, builds a per-row dynamic time array, and evaluates the
    extraction profile at each time point.  Also computes a shared pseudotime
    axis based on mean inter-event intervals.

    Parameters
    ----------
    data : MultiSessionDataset
        Dataset providing session data and trace access.
    segments : iterable of SegmentReference
        Segments to extract from.
    events : iterable of str
        Ordered event names that define the time landmarks.
    extraction_profile : TraceExtractionProfile
        Specifies signal, mode, sample rate, time type, and fill value.
    samples : list of int or None, optional
        Override the ideal sample count for each interval.  ``None`` entries
        use the ideal value.  Must have length ``len(events) - 1`` (or
        ``len(events) + 1`` if ``padding=True``).
    event_pseudotimes : list of float or None, optional
        Override the pseudotime anchor for each event.  ``None`` entries use
        the ideal (mean-based) value.
    baseline_range : tuple of (float, float), optional
        If provided, subtract the mean of the trace within
        ``(baseline_range[0], baseline_range[1])`` pseudotime from each row.
    padding : bool, optional
        If ``True``, prepend ``pad_start`` seconds before the first event and
        append ``pad_end`` seconds after the last event.
    pad_start : float, optional
        Seconds of padding before the first event. Default is 2.
    pad_end : float, optional
        Seconds of padding after the last event. Default is 6.

    Returns
    -------
    pseudotime : np.ndarray
        1D pseudotime axis of length ``sum(samples) + 1``.
    result_arr : np.ndarray
        2D trace array of shape ``(n_segments, len(pseudotime))``.
    info : dict
        Additional information: ``'event_times'``, ``'samples_ideal'``,
        ``'pseudoeventtimes_ideal'``.

    Raises
    ------
    KeyError
        If any event is missing from any segment.
    ValueError
        If ``len(samples)`` is incompatible with ``len(events)``.
    """
    event_times = extract_event_times_multiple(data, segments, events)
    if np.isnan(event_times).sum() > 0:
        raise KeyError('Was not able to find all events in all segments')
    if padding:
        event_times_padded = np.empty((len(segments), len(events)+2))
        event_times_padded[:, 1:-1] = event_times
        event_times_padded[:, 0] = event_times[:, 0] - pad_start
        event_times_padded[:, -1] = event_times[:, -1] + pad_end
        event_times = event_times_padded
    samples_ideal = determine_ideal_samplerate(
        event_times, extraction_profile)
    if samples is None:
        samples = samples_ideal
    samples = [s if s is not None else i
               for s, i in zip(samples, samples_ideal)]
    if len(samples)+1 != event_times.shape[1]:
        raise ValueError('Length of samples must be length of events -1\n' +
                         f'samples is currently {len(samples)}\n' +
                         f'events is currently {event_times.shape[1]}')
    # Time conversion...if required
    if extraction_profile.time_type != 'unix_time':
        event_times = timeconversion_bulk(data,
                                          [seg.session for seg in segments],
                                          event_times,
                                          'unix_time',
                                          extraction_profile.time_type
                                          )
    tarr = create_dynamictimearray_2d(event_times, samples)
    trace_references = [TraceReference(seg.session,
                                       row,
                                       extraction_profile.time_type)
                        for seg, row in zip(segments, tarr)]
    result_arr = np.empty((tarr.shape))
    for ind, ref in enumerate(trace_references):
        result_arr[ind, :] = data.get_trace(ref, extraction_profile)

    # Take the event times and calculate average time distance between events
    diffs = np.empty((event_times.shape[1]))
    diffs[0] = 0
    diffs[1:] = np.diff(event_times, axis=1).mean(axis=0)
    pseudoeventtimes_ideal = np.cumsum(diffs)
    pseudoeventtimes_ideal = [float(e) for e in pseudoeventtimes_ideal]

    if event_pseudotimes is None:
        event_pseudotimes = pseudoeventtimes_ideal
    event_pseudotimes = [e if e is not None else i
                         for e, i in zip(event_pseudotimes,
                                         pseudoeventtimes_ideal)
                         ]

    pseudotime = create_dynamictimearray(event_pseudotimes, samples)

    if baseline_range is not None:
        bl_mask = ((pseudotime > baseline_range[0]) &
                   (pseudotime < baseline_range[1]))
        result_arr = result_arr - result_arr[:, bl_mask
                                             ].mean(axis=1)[:, np.newaxis]

    info = {'event_times': event_times,
            'samples_ideal': samples_ideal,
            'pseudoeventtimes_ideal': pseudoeventtimes_ideal
            }

    return pseudotime, result_arr, info


class SessionInterpolatedTraceExtractor:
    """
    Deprecated per-session trace extractor.

    .. deprecated::
        Use :func:`trace_extractor_factory` via :class:`MultiSessionDataset`
        instead.
    """

    def __init__(self, session: Session, time_type):
        """
        Parameters
        ----------
        session : Session
            Session to build extractors from.
        time_type : str
            Time axis key to use for interpolation.
        """
        print('SessionInterpolatedTraceExtractor is depreciated')
        self.time_type = time_type
        self.rigdata = {}
        self.rigdata_linear = {}
        self.rigdata_raw = session.rigdata
        for rdat, rdict in session.rigdata.items():
            if rdat == 'SyncStream':
                continue
            t = rdict[time_type]
            v = rdict['report_value']

            self.rigdata[rdat] = interp1d(
                t, v, kind='nearest',
                bounds_error=False, fill_value=np.nan
            )

            self.rigdata_linear[rdat] = interp1d(
                t, v, kind='linear',
                bounds_error=False, fill_value=np.nan
            )

    def extract(self, rig_data: str, time_array: np.ndarray, mode="nearest"):
        """
        Extract trace values using nearest or linear interpolation.

        Parameters
        ----------
        rig_data : str
            Report channel name.
        time_array : np.ndarray
            Time points at which to evaluate the trace.
        mode : {'nearest', 'linear'}, optional
            Interpolation mode. Default is ``'nearest'``.

        Returns
        -------
        np.ndarray
            Extracted values.
        """
        if mode == "nearest":
            return self.rigdata[rig_data](time_array)
        elif mode == "linear":
            return self.rigdata_linear[rig_data](time_array)
        else:
            raise ValueError(f"Unknown interpolation mode: {mode}")

    def extract_windowed_dynamic(self, rig_data: str, time_array: np.ndarray,
                                 mode="mean", fill=np.nan):
        """
        Extract trace values using adaptive windowed aggregation.

        Parameters
        ----------
        rig_data : str
            Report channel name.
        time_array : np.ndarray
            Time points defining window centres.
        mode : {'mean', 'sum', 'count', 'rate'}, optional
            Aggregation mode. Default is ``'mean'``.
        fill : float, optional
            Value for windows with no data. Default is ``np.nan``.

        Returns
        -------
        np.ndarray
            Aggregated values at each time point.
        """
        t_data = self.rigdata_raw[rig_data][self.time_type]
        v_data = self.rigdata_raw[rig_data]['report_value']

        n = len(time_array)
        left_window = np.empty(n)
        right_window = np.empty(n)

        # edges
        left_window[0] = (time_array[1] - time_array[0])/2
        right_window[0] = left_window[0]
        left_window[-1] = (time_array[-1] - time_array[-2]) / 2
        right_window[-1] = left_window[-1]

        # interior points
        left_window[1:-1] = (time_array[1:-1] - time_array[:-2]) / 2
        right_window[1:-1] = (time_array[2:] - time_array[1:-1]) / 2

        out = np.full(n, fill)
        for i, ti in enumerate(time_array):
            mask = (t_data >= ti - left_window[i]
                    ) & (t_data <= ti + right_window[i])
            if not mask.any():
                continue
            if mode == "mean":
                out[i] = np.nanmean(v_data[mask])
            elif mode == "sum":
                out[i] = np.nansum(v_data[mask])
            elif mode == "count":
                out[i] = np.sum(~np.isnan(v_data[mask]))
            elif mode == "rate":
                out[i] = np.sum(~np.isnan(v_data[mask])) / \
                    (left_window[i] + right_window[i])
            else:
                raise ValueError(f"Unknown mode: {mode}")
        return out
```
