## Extended Data for "A Low-Cost, Modular Hardware and Software Platform for Head-Fixed Mouse Decision-Making Tasks": bandit.py

```
# -*- coding: utf-8 -*-
"""
Created on Tue Jun 10 13:10:25 2025

@author: mm4114
"""
import time
import random
from copy import deepcopy
from typing import ClassVar, Literal, List, Tuple

from lampyr.segments import Trial, Task, BehaviorSegment
from lampyr.segments.paradigm import Stage, Paradigm
from lampyr.math import rand_int_uniform_hazard
from dataclasses import dataclass, field

# -------------- Define Event Callbacks --------------
# Simple events simply log a timepoint, complex events can also interact with the rig and log reports as needed.


def event_waterreward(self: BehaviorSegment):
    """Give a water reward via the rig and log the reward event."""
    self.log_debug('Sending give reward command to rig')
    self.rig.reward.give()
    self.log_reward(0.005)


def event_trialstart(self: BehaviorSegment):
    """Play the trial-start tone and log the event."""
    self.log_debug('Sending play trial tone command to rig')
    self.rig.play.begintrialtone()


def event_response(self: BehaviorSegment):
    """Play the response registration tone and log the event"""
    self.log_debug('Sending play response tone command to rig')
    self.rig.play.responsetone()


def event_laser_onset(self: BehaviorSegment):
    self.rig.laser.begin()
    self.log_debug('Sending laser constant illumination command to rig')


# -------------- Define Habituation Trial/Task --------------


@dataclass
class HabituationTrial(Trial):
    """
    A single habituation trial.

    1. Wait for a fixed ITI (``iti1_dur``).
    2. Deliver reward and trigger the reward event.
    3. Monitor licks for ``reward_consumption_period_s`` seconds.  If no lick is detected, extend monitoring for ``reward_consumption_nolick_delay_s`` seconds.
    4. Log whether the reward was consumed (lick detected) and count merit/abstention accordingly.
    5. Wait for a second ITI (``iti2_dur``) before finishing the trial.

    Attributes
    ----------
    iti1_dur : float
        Pre-reward inter-trial interval in seconds.
    reward_consumption_period_s : float
        Primary lick-detection window after reward delivery (seconds).
    reward_consumption_nolick_delay_s : float
        Extended wait period if no lick detected in the primary window (seconds).
    count_merits : bool
        If ``True``, log merit when the reward is consumed.
    iti2_dur : float
        Post-consumption inter-trial interval in seconds.
    """
    iti1_s: float = 1
    reward_consumption_period_s: float = 15
    reward_consumption_nolick_delay_s: float = 40
    count_merits: bool = True
    iti2_s: float = 0.5

    def setup(self):
        """Register the reward event callback and description."""
        self.register_event('reward',
                            callback=event_waterreward,
                            description='Water reward given')

    def perform(self):
        """
        Execute one habituation trial iteration: ITI → reward → consumption → ITI.
        """
        # Start behavior
        self.wait(self.iti1_s)
        self.log_info('Reward dispensed')
        rewarddelivery_time = self.trigger_event('reward')
        self.wait(self.reward_consumption_period_s)
        if not self.rig.licks.since(rewarddelivery_time):
            self.log_info(f'No licks detected  in {self.reward_consumption_period_s} seconds, waiting...')
            self.wait(self.reward_consumption_nolick_delay_s)
        lick_count = self.rig.licks.since(rewarddelivery_time)
        if lick_count > 0:
            self.create_report('reward_consumed', True)
            if self.count_merits:
                self.log_merit()
        else:
            self.create_report('reward_consumed', False)
            if self.count_merits:
                self.log_abstention()
        self.log_notice(f'Detected {lick_count} licks since reward delivery.')
        self.wait(self.iti2_s)


@dataclass
class RewardedHabituationTask(Task):
    """
    Task that repeatedly runs :class:`HabituationTrial` instances until the
    session stop conditions are met.
    """
    slug: str = 'RewHab'

    def setup(self):
        self.register_event('reward',
                            callback=event_waterreward,
                            description='Water reward given')

    def loop(self):
        """Create and run one :class:`HabituationTrial` per iteration."""
        trial = HabituationTrial(parent=self,
                                 iti1_s=1,
                                 reward_consumption_period_s=8,
                                 reward_consumption_nolick_delay_s=40,
                                 iti2_s=random.randint(4, 12),
                                 count_merits=True)
        trial.run()
        del trial

# -------------- Define Bandit Trials/Tasks --------------


@dataclass
class BanditTrial(Trial):
    iti1_s: float = 1
    pt_hold_s: float = 2
    pt_mvmt_threshold_deg: float = 5
    pt_trial_delay: float = 0
    responsewindow_s: float = 3
    responsethresholds_deg: dict = field(default_factory=lambda: {'Left': 15,
                                                                  'Right': 15})
    rewardprobs_perc: dict = field(default_factory=lambda: {'Left': 100,
                                                            'Right': 100,
                                                            'None': 0})

    reward_delay_s: float = 0
    iti2_s: float = 2.5
    enable_wheel_lock: bool = False

    # laser stuff
    pretrial_laser_enabled: bool = False
    precue_laser_enabled: bool = False
    precue_laser_offset: float = 0.5
    response_laser_enabled: bool = False
    response_laser_delay_s: float = 0.1
    
    laserstop_response_offramp_enabled: bool = False
    laserstop_response_offramp_ms: int = 200
    laserstop_trialend_offramp_enabled: bool = False
    laserstop_trialend_offramp_ms: int = 500

    def setup(self):
        """Register trial events"""
        self.register_event('reward',
                            callback=event_waterreward,
                            description='Water reward given')
        self.register_event('trialstart',
                            callback=event_trialstart,
                            description='Trial begins. Start tone played.')
        self.register_event('pretrialstart',
                            description='beginning of pretrial period')
        self.register_event('response',
                            callback=event_response,
                            description='Wheel response registered')
        self.register_event('laser_onset',
                            callback=event_laser_onset,
                            description='Laser Onset')
        self.log_debug(self.rewardprobs_perc)

        if self.response_laser_enabled and (
                self.response_laser_delay_s > self.reward_delay_s):
            self.log_warning(
                'Response laser delay cannot be larger than reward delay')
            self.response_laser_delay_s = self.reward_delay_s
            self.log_warning(f'response_laser_delay set to {self.reward_delay_s}')

    def perform(self):
        laser_on = False

        self.wait(self.iti1_s)
        if self.pretrial_laser_enabled:
            self.trigger_event('laser_onset')
            laser_on = True
        self.trigger_event('pretrialstart')
        self.log_info('Waiting for pretrial wheel hold...')
        self.wait(self.pt_hold_s)
        self.waitfor(
            condition=lambda: self.rig.wheel.movement_total_since(
                time.time()-self.pt_hold_s) < self.pt_mvmt_threshold_deg,
            timeout=None,
            poll_interval=0.01,
            while_waiting=lambda: self.log_notice(
                'Animal movement detected. Waiting for cessation.'),
            while_waiting_interval=4
        )
        if self.precue_laser_enabled:
            self.wait(max(self.pt_trial_delay - self.precue_laser_offset,0))
            self.trigger_event('laser_onset')
            laser_on = True
            self.wait(self.precue_laser_offset)
        else:
            self.wait(self.pt_trial_delay)
        self.log_info('Trial start')
        tstart_time = self.trigger_event('trialstart')
        self.rig.wheel.home()
        response = self.waitfor(
            condition=self.response_loop,
            fallback_value='None',
            timeout=self.responsewindow_s,
            poll_interval=0.005
        )
        response_time = time.time()
        if response != 'None':
            if self.enable_wheel_lock:
                self.rig.wheellock.to_angle(65)
                self.log_info('Wheel Locked')
            resp_time = self.trigger_event('response')
            self.create_report('response_delay', resp_time-tstart_time)
        self.create_report('response', response)
        highestrewardprob = max(self.rewardprobs_perc.values())
        if self.rewardprobs_perc[response] == highestrewardprob:
            self.log_merit()
            self.create_report('best_response', True)
            was_best = True
        else:
            if response == 'None':
                self.log_abstention()
            else:
                self.log_demerit()
            self.create_report('best_response', False)
            was_best = False
        self.log_info(f'Response was {response} ('
                      + ('not ' * (not was_best))
                      + 'best response)')
        probability = self.rewardprobs_perc.get(response, 0) / 100
        rand = random.random()
        self.log_debug(f'RAND:{rand},THRESH:{probability}')
        self.log_debug(f'Reward delay: {self.reward_delay_s}')
        if laser_on and self.laserstop_response_offramp_enabled:
            self.wait(self.response_laser_delay_s)
            self.rig.laser.rampdown(self.laserstop_response_offramp_ms)
            laser_on = False
            self.wait(self.reward_delay_s - self.response_laser_delay_s)
        elif self.response_laser_enabled and response != 'None':
            self.wait(self.response_laser_delay_s)
            self.trigger_event('laser_onset')
            laser_on = True
            self.wait(self.reward_delay_s - self.response_laser_delay_s)
        else:
            self.wait(self.reward_delay_s)
        anticipatory_licks = self.rig.licks.since(response_time)
        self.create_report('anticipatory_licks', anticipatory_licks)
        self.log_info(f'{anticipatory_licks} anticipatory licks logged')
        reward_time = time.time()
        if rand < probability:
            self.log_info(f'Reward given ({round(probability*100)})% chance.')
            self.trigger_event('reward')
            self.create_report('rewarded', True)
        else:
            self.log_info(
                f'No reward given ({round(probability*100)})% chance.')
            self.create_report('rewarded', False)
        self.wait(self.iti2_s)
        reward_licks = self.rig.licks.since(reward_time)
        self.create_report('reward_licks', reward_licks)
        self.log_info(f'{reward_licks} reward licks logged')
        self.rig.wheellock.unlock()
        self.log_info('Wheel Unlocked')
        if self.laserstop_trialend_offramp_enabled and laser_on:
            self.log_info('Laser ramping down')
            self.rig.laser.rampdown(self.laserstop_trialend_offramp_ms)
            laser_on = False

    def response_loop(self):
        wheel_pos = self.rig.wheel.angle()
        if wheel_pos < -self.responsethresholds_deg['Left']:
            return 'Left'
        if wheel_pos > self.responsethresholds_deg['Right']:
            return 'Right'

    def response_made(self) -> bool:
        return self.reports.get('response', 'None') != 'None'

    def satisfiesblockcondition(self, counttype: str) -> bool:
        """
        Check whether this trial satisfies a task-block counting condition.

        Parameters
        ----------
        counttype : str
            One of ``'Reward'``, ``'Merit'``, or ``'RewardedMerit'``.

        Returns
        -------
        bool
            Whether the trial meets the specified condition.
        """
        if self.reports.get('response', 'None') == 'None':
            return False
        match counttype:
            case 'Reward':
                return bool(self.reports.get('rewarded', False))
            case 'Merit':
                return bool(self.reports.get('best_response', False))
            case 'RewardedMerit':
                return bool(self.reports.get('rewarded', False)) and bool(self.reports.get('best_response', False))


@dataclass
class BanditTask(Task):
    target_mode: Literal['Random', 'Any', 'Left', 'Right'] = 'Random'
    reward_prob_target: int = 80
    reward_prob_offtarget: int = 0
    iti1_s: float = 1
    pt_trial_delay: float = 0
    reward_delay_s: float = 0

    rescue_trial_enabled: bool = False
    rescue_limit: int = 3
    rescue_cooldown: int = 20
    rescue_threshold: int = 15

    taskblocks_enabled: bool = True
    taskblocks_sizerange: tuple = (6, 15)
    taskblocks_blockcounttype: Literal['Reward',
                                       'Merit', 'RewardedMerit'] = 'Reward'

    enable_wheel_lock: bool = False
    
    #tracking for lasers
    current_response_trial_number: int = 0
    current_trial_number: int = 0

    # lasers
    enable_laser_trials: bool = False
    laser_trial_sequence: list = None
    laser_trial_sequence_type: Literal['response', 'trial'] = 'response'
    # laser param list
    laser_trial_params = (
        'pretrial_laser_enabled',
        'response_laser_enabled',
        'response_laser_delay_s',
        'laserstop_trialend_offramp_enabled',
        'laserstop_trialend_offramp_ms',
        'precue_laser_enabled',
        'precue_laser_offset',
        'laserstop_response_offramp_enabled',
        'laserstop_response_offramp_ms')
    # laser onset/offset
    pretrial_laser_enabled: bool = False
    precue_laser_enabled: bool = False
    precue_laser_offset: float = 0.5
    response_laser_enabled: bool = False
    response_laser_delay_s: float = 0.1

    laserstop_trialend_offramp_enabled: bool = False
    laserstop_trialend_offramp_ms: int = 500
    laserstop_response_offramp_enabled: bool = False
    laserstop_response_offramp_ms: int = 200

    def setup(self):
        if self.target_mode == 'Random':
            self._target = random.choice(['Left', 'Right'])
        else:
            self._target = self.target_mode
        # Trial Block Info
        self._thisblocksize = random.randint(*self.taskblocks_sizerange)
        self._trialinblockcount = 0
        # Rescue Trial Info
        self._rescue_count = 0
        self._rescue_sincelast = 10000000

        self._reward_probs = {
            'Left':  {'Left': self.reward_prob_target,   'Right': self.reward_prob_offtarget, 'None': 0},
            'Right': {'Left': self.reward_prob_offtarget, 'Right': self.reward_prob_target,   'None': 0},
            'Any':   {'Left': self.reward_prob_target,   'Right': self.reward_prob_target,    'None': 0},
        }

    def loop(self):
        # if lasers are enabled, construct the laser parameters for the current trial
        ltparams = {}
        if self.enable_laser_trials:
            if self.laser_trial_sequence is not None:
                if self.laser_trial_sequence_type == 'response':
                    idx = self.current_response_trial_number % len(
                        self.laser_trial_sequence)
                elif self.laser_trial_sequence_type == 'trial':
                    idx = self.current_trial_number % len(
                        self.laser_trial_sequence)
                else:
                    raise ValueError('laser_trial_sequence_type set to invalid key')
                if self.laser_trial_sequence[idx]:
                    ltparams = {k: getattr(self, k)
                                for k in self.laser_trial_params}

        trial = BanditTrial(parent=self,
                            iti1_s = self.iti1_s,
                            pt_trial_delay = self.pt_trial_delay,
                            reward_delay_s=self.reward_delay_s,
                            rewardprobs_perc=self._reward_probs[self._target],
                            enable_wheel_lock=self.enable_wheel_lock,
                            **ltparams)
        trial.run()
        self.current_trial_number += 1
        if trial.response_made():
            self.current_response_trial_number += 1
        if self.taskblocks_enabled:
            if trial.satisfiesblockcondition(self.taskblocks_blockcounttype):
                self._trialinblockcount += 1
            if self._trialinblockcount >= self._thisblocksize:
                self._trialinblockcount = 0
                self._thisblocksize = rand_int_uniform_hazard(
                    *self.taskblocks_sizerange)
                match self._target:
                    case 'Left':
                        self._target = 'Right'
                    case 'Right':
                        self._target = 'Left'
                    case 'Any':
                        self._target = 'Any'
                self.log_notice(f'Target has been switched to {self._target}')
                self.log_notice('Blocksize is set to '
                                + str(self._thisblocksize))
        del trial

        self._rescue_sincelast += 1
        if (self.rescue_trial_enabled
            and self.session.serial_abstention >= self.rescue_threshold
            and self._rescue_count < self.rescue_limit
                and self._rescue_sincelast >= self.rescue_cooldown):
            self.log_notice('Attempting to rescue performance...')
            rescue_trial = HabituationTrial(parent=self,
                                            slug='Rescue',
                                            count_merits=False,
                                            iti1_s=60,
                                            reward_consumption_period_s=30,
                                            reward_consumption_nolick_delay_s=60,
                                            iti2_s=0.5)
            rescue_trial.run()
            del rescue_trial
            self._rescue_count += 1
            self._rescue_sincelast = 0

        if (self.session.serial_abstention >= 20
                and self.session.serial_abstention % 10 == 0):
            self.notify(f'Animal has not responded in {self.session.serial_abstention} trials')

# -------------- Define Training Stages and Training Paradigm --------------


@dataclass
class HabituationStage(Stage):
    """
    Stage 0: free-reward habituation.

    Runs :class:`RewardedHabituationTask` and advances the mouse to the
    ``'AnyWheel'`` stage after two consecutive sessions meeting the merit
    threshold.
    """

    slug: str = 'Stage0Hab'

    def define_sessionparams(self):
        """Set duration (60 min), serial abstention (10), and reward (200) limits."""
        self.set_sessionparam('duration_limit', 90)
        self.set_sessionparam('serial_abstention_limit', 10)
        self.set_sessionparam('reward_amount_limit', 1.0)

    def define_task(self, stage_data):
        """Run the habituation task."""
        task = RewardedHabituationTask(parent=self)
        task.run()
        del task

    def session_summary(self):
        pass

    def define_shaping(self, stage_data):
        consecutive_good_sessions = stage_data.get(
            'consecutive_good', 0)
        if self.session.duration < 30:
            return
        if self.session.merit > 140:
            consecutive_good_sessions += 1
        else:
            consecutive_good_sessions = 0
        stage_data['consecutive_good'] = consecutive_good_sessions


@dataclass
class ResponseAbstractStage(Stage):
    """
    Abstract base for wheel-response stages, with shared session params and reporting.

    Provides common default session limits and the :meth:`sessionsummary`
    helper used by all response-based stages.
    """

    slug: str = 'ResponseAbstractStage'

    def define_sessionparams(self):
        """Set shared defaults: 60 min max, 20 min min, serial abstention 30, reward 200."""
        self.set_sessionparam('duration_limit', 60)
        self.set_sessionparam('duration_min', 60)
        self.set_sessionparam('serial_abstention_limit', 30)
        self.set_sessionparam('reward_amount_limit', 1.0)

    def _compute_sb_metric(self):
        """Returns (Right - Left) / (Right + Left), or None if no directional responses."""
        trials = self.searchsubsegments(slug='BanditTrial', type='Trial')
        responses = self.summarize_reportsinsegments('response', trials)
        resp_r = responses.get('Right', 0)
        resp_l = responses.get('Left', 0)
        if resp_r + resp_l == 0:
            return None
        return (resp_r - resp_l) / (resp_l + resp_r)

    def define_globalshaping(self, global_data):
        adjustment = global_data.get('adjustmentvalue', 0)
        def allowedrange(deg): return min(max(5, deg), 40)
        responsethresholds_deg = {'Left': allowedrange(15-adjustment),
                                  'Right': allowedrange(15+adjustment)}
        self.mouse.mouse_behav_param_overrides[
            'BanditTrial']['responsethresholds_deg'] = responsethresholds_deg

    def session_summary(self):
        """Log response counts and side bias. Returns sb_metric (float) or None."""
        trials = self.searchsubsegments(slug='BanditTrial', type='Trial')
        responsecounts = self.summarize_reportsinsegments('response', trials)
        for resp, num in responsecounts.items():
            self.log_info(f'Detected {num} {resp} responses.')
        sb_metric = self._compute_sb_metric()
        if sb_metric is None:
            self.log_info(
                'Sidebias could not be calculated due to no responses.')
        else:
            sb_perc = (sb_metric * 50) + 50
            self.log_info(
                f'Sidebias was: {round(sb_metric, 2)} ({round(sb_perc)}% Right)'
            )


@dataclass
class AnyWheelStage(ResponseAbstractStage):
    """
    Stage 1: any-direction wheel training with 100% reward probability.

    Advances to ``'AltWheel'`` after the required number of consecutive
    sessions meeting the participation threshold.
    """
    slug: str = 'Stage1AnyWheel'

    def define_task(self, stage_data):
        """Run an any-direction BanditTask with 100% reward and rescue trials enabled."""
        task = BanditTask(parent=self,
                          target_mode='Any',
                          reward_prob_target=100,
                          reward_prob_offtarget=0,
                          reward_delay_s=0,
                          rescue_trial_enabled=True,
                          taskblocks_enabled=False)
        task.run()
        del task

    def define_shaping(self, stage_data):
        """
        Advance to AltWheel after the required consecutive sessions above participation threshold.
        """
        consecutive_good_sessions = stage_data.get(
            'consecutive_good', 0)
        if self.session.duration < 30:
            return
        if self.session.participation >= 150:
            consecutive_good_sessions += 1
            self.log_info('This session was a good session (>=150 responses)')
        else:
            consecutive_good_sessions = 0

        self.log_info(
            f'Number of consecutive good sessions is {consecutive_good_sessions}')
        stage_data['consecutive_good'] = consecutive_good_sessions


@dataclass
class AltWheelStage1(ResponseAbstractStage):
    """
    Stage 2: alternating-target bandit with adaptive threshold correction.

    Runs a standard bandit task and applies a three-phase threshold adjustment
    algorithm (correction → return → complete) to correct side bias while
    keeping wheel thresholds within bounds.
    """
    slug: str = 'Stage2Correction'

    def define_task(self, stage_data):
        """Run the alternating bandit task with 100/0 reward probabilities."""
        task = BanditTask(parent=self,
                          reward_prob_target=100,
                          reward_prob_offtarget=0,
                          rescue_trial_enabled=True,
                          reward_delay_s=0)
        task.run()
        del task

    def define_shaping(self, stage_data):
        consecutive_good_sessions = stage_data.get(
            'consecutive_good', 0)
        global_paradigm_data = self.get_globalparadigmdata()
        adj_val = global_paradigm_data.get('adjustmentvalue', 0)
        side_bias = self._compute_sb_metric()

        if side_bias is None or self.session.duration < 30:
            return

        def sign(x): return 1 if x > 0 else -1 if x < 0 else 0
        def signdiff(x, y): return (sign(x) * sign(y)) == -1

        if side_bias < 0:
            bias = 'Leftward'
        elif side_bias > 0:
            bias = 'Rightward'
        else:
            bias = 'Neutral'

        if adj_val == 0:  # No adjustment value set
            self.log_info('Assigning adjustment value for first time')
            if side_bias == 0:
                consecutive_good_sessions += 1
            else:
                adj_val += 5*sign(side_bias)
                consecutive_good_sessions = 0
        elif abs(side_bias) > 0.2:
            if signdiff(adj_val, side_bias):
                self.log_info('Adjustment value has overshot bias')
                self.log_info('This was a good session (contrary side bias)')
                consecutive_good_sessions += 1
            else:
                self.log_info(
                    f'{bias} bias detected. Adjustment value updated.')
                adj_val += 5*sign(side_bias)
                consecutive_good_sessions = 0
        else:
            self.log_info('This was a good session (no side bias detected)')
            consecutive_good_sessions += 1

        self.log_info('Adjustment Value is ' + str(adj_val))
        self.log_info('Consecutive good sessions is ' +
                      str(consecutive_good_sessions))
        adj_val = max(-40, min(40, adj_val))
        global_paradigm_data['adjustmentvalue'] = adj_val
        stage_data['consecutive_good'] = consecutive_good_sessions


@dataclass
class AltWheelStage2(ResponseAbstractStage):
    """
    Stage 2: alternating-target bandit with adaptive threshold correction.

    Runs a standard bandit task and applies a three-phase threshold adjustment
    algorithm (correction → return → complete) to correct side bias while
    keeping wheel thresholds within bounds.
    """
    slug: str = 'Stage3Return'

    def define_task(self, stage_data):
        """Run the alternating bandit task with 100/0 reward probabilities."""
        task = BanditTask(parent=self,
                          reward_prob_target=100,
                          reward_prob_offtarget=0,
                          rescue_trial_enabled=True,
                          reward_delay_s=0)
        task.run()
        del task

    def define_shaping(self, stage_data):
        consecutive_good_sessions = stage_data.get(
            'consecutive_good', 0)
        global_paradigm_data = self.get_globalparadigmdata()
        adj_val = global_paradigm_data.get('adjustmentvalue', 0)
        side_bias = self._compute_sb_metric()

        if side_bias is None or self.session.duration < 30:
            return

        # If side bias is against adjustment value, return by 2
        def sign(x): return 1 if x > 0 else -1 if x < 0 else 0
        def signdiff(x, y): return (sign(x) * sign(y)) == -1

        if side_bias < 0:
            bias = 'Leftward'
        elif side_bias > 0:
            bias = 'Rightward'
        else:
            bias = 'Neutral'

        if (adj_val == 0 and (-0.15 < side_bias < 0.15)
                and self.session.merit >= 150):
            self.log_info(
                'No/low bias and sufficient performance for good session')
            consecutive_good_sessions += 1
        else:
            self.log_info('Bias or low performance detected')
            consecutive_good_sessions = 0

        if signdiff(side_bias, adj_val) or abs(side_bias) < 0.1:
            self.log_info(f'No/opposite bias at adjustment score {adj_val}')
            adj_val = adj_val - (sign(adj_val) * min(abs(adj_val), 2))

        self.log_info('Adjustment Value is ' + str(adj_val))
        self.log_info('Consecutive good sessions is ' +
                      str(consecutive_good_sessions))
        adj_val = max(-40, min(40, adj_val))
        global_paradigm_data['adjustmentvalue'] = adj_val
        stage_data['consecutive_good'] = consecutive_good_sessions


@dataclass
class AltWheelDelayStage(ResponseAbstractStage):
    """
    Intermediate stage that progressively introduces a reward delay.

    Steps through ``BanditParadigm.DELAY_STEPS`` based on merit performance,
    advancing to ``'BanditTraining'`` once the longest delay step is reached.
    """
    slug: str = 'Stage4AltWheelDelay'
    delay_progression = (0.1, 0.25, 0.5, 1)

    def define_task(self, stage_data):
        """Run the alternating bandit task with 100/0 reward probabilities."""
        current_delay = stage_data.get('current_delay', 0)
        rwdelay = self.delay_progression[current_delay]

        task = BanditTask(parent=self,
                          reward_prob_target=100,
                          reward_prob_offtarget=0,
                          rescue_trial_enabled=True,
                          reward_delay_s=rwdelay)
        task.run()
        del task

    def define_shaping(self, stage_data):
        consecutive_good_sessions = stage_data.get(
            'consecutive_good', 0)
        current_delay = stage_data.get('current_delay', 0)

        if self.session.duration < 30:
            return

        if current_delay < (len(self.delay_progression)-1):
            if self.session.merit >= 150:
                self.log_info('This was a good session (>=150 merit)')
                self.log_notice('Increasing reward delay...')
                current_delay += 1
        else:
            if self.session.merit >= 150:
                consecutive_good_sessions += 1
            else:
                consecutive_good_sessions = 0

        self.log_info('Current delay is ' +
                      str(self.delay_progression[current_delay]))
        self.log_info('Consecutive good sessions is ' +
                      str(consecutive_good_sessions))
        stage_data['consecutive_good'] = consecutive_good_sessions
        stage_data['current_delay'] = current_delay


@dataclass
class BanditTrainingStage(ResponseAbstractStage):
    """
    Full bandit training stage with the default reward probability schedule.
    """
    slug: str = 'Stage5BanditTraining'
    _task: object = None

    def define_task(self, stage_data):
        """Run the bandit task with rescue trials and retain a reference for analysis."""
        task = BanditTask(parent=self,
                          rescue_trial_enabled=True,
                          reward_delay_s=0)
        task.run()
        del task

    def define_shaping(self, stage_data):
        consecutive_good_sessions = stage_data.get(
            'consecutive_good', 0)

        if self.session.duration < 30:
            return

        if self.session.merit >= 150:
            self.log_info('This was a good session (merit >= 150)')
            consecutive_good_sessions += 1
        else:
            consecutive_good_sessions = 0

        self.log_notice('Consecutive good sessions is ' +
                        str(consecutive_good_sessions))
        stage_data['consecutive_good'] = consecutive_good_sessions


@dataclass
class BanditEndStage(ResponseAbstractStage):
    """
    Final bandit stage run without rescue trials.
    """
    slug: str = 'Stage6Bandit'

    def define_task(self, stage_data):
        """Run the bandit task without rescue trials."""
        task = BanditTask(parent=self,
                          rescue_trial_enabled=False)
        task.run()
        del task

    def define_shaping(self, stage_data):
        pass


@dataclass
class BanditParadigm(Paradigm):
    slug: str = 'BanditParadigm2'
    stagelist: tuple = (HabituationStage,
                        AnyWheelStage,
                        AltWheelStage1,
                        AltWheelStage2,
                        AltWheelDelayStage,
                        BanditTrainingStage,
                        BanditEndStage)

    def define_progression(self, current_stage, stage_data):
        if current_stage is HabituationStage:
            if stage_data.get('consecutive_good', 0) >= 2:
                self.progress()
        elif current_stage is AnyWheelStage:
            if stage_data.get('consecutive_good', 0) >= 2:
                self.progress()
        elif current_stage is AltWheelStage1:
            if stage_data.get('consecutive_good', 0) >= 2:
                self.progress()
        elif current_stage is AltWheelStage2:
            if stage_data.get('consecutive_good', 0) >= 2:
                self.progress()
        elif current_stage is AltWheelDelayStage:
            if stage_data.get('consecutive_good', 0) >= 2:
                self.progress()
        elif current_stage is BanditTrainingStage:
            if stage_data.get('consecutive_good', 0) >= 2:
                self.progress()
        elif current_stage is BanditEndStage:
            pass


# BANDIT PARADIGM 3 - Created 6.12.26 in response to issues with animal training
# Animal response curves not sufficient
@dataclass
class AnyWheelStageB3(ResponseAbstractStage):
    """
    Stage 1: any-direction wheel training with 100% reward probability.

    Advances to ``'AltWheel'`` after the required number of consecutive
    sessions meeting the participation threshold.
    """
    slug: str = 'Stage1AnyWheel'

    def define_task(self, stage_data):
        """Run an any-direction BanditTask with 100% reward and rescue trials enabled."""
        task = BanditTask(parent=self,
                          target_mode='Any',
                          reward_prob_target=100,
                          reward_prob_offtarget=0,
                          reward_delay_s=0,
                          rescue_trial_enabled=True,
                          taskblocks_enabled=False,
                          enable_wheel_lock=True)
        task.run()
        del task

    def define_shaping(self, stage_data):
        """
        Advance to AltWheel after the required consecutive sessions above participation threshold.
        """
        consecutive_good_sessions = stage_data.get(
            'consecutive_good', 0)
        if self.session.duration < 30:
            return
        if self.session.participation >= 100:
            consecutive_good_sessions += 1
            self.log_info('This session was a good session (>=150 responses)')
        else:
            consecutive_good_sessions = 0

        self.log_info(
            f'Number of consecutive good sessions is {consecutive_good_sessions}')
        stage_data['consecutive_good'] = consecutive_good_sessions


@dataclass
class AltWheelStage1B3(ResponseAbstractStage):
    """
    Stage 2: alternating-target bandit with adaptive threshold correction.

    Runs a standard bandit task and applies a three-phase threshold adjustment
    algorithm (correction → return → complete) to correct side bias while
    keeping wheel thresholds within bounds.
    """
    slug: str = 'Stage2Correction'

    def define_task(self, stage_data):
        """Run the alternating bandit task with 100/0 reward probabilities."""
        task = BanditTask(parent=self,
                          reward_prob_target=100,
                          reward_prob_offtarget=0,
                          rescue_trial_enabled=True,
                          taskblocks_enabled=True,
                          reward_delay_s=0,
                          enable_wheel_lock=True
                          )
        task.run()
        del task

    def define_shaping(self, stage_data):
        consecutive_good_sessions = stage_data.get(
            'consecutive_good', 0)
        global_paradigm_data = self.get_globalparadigmdata()
        adj_val = global_paradigm_data.get('adjustmentvalue', 0)
        side_bias = self._compute_sb_metric()

        if side_bias is None or self.session.duration < 30:
            return

        def sign(x): return 1 if x > 0 else -1 if x < 0 else 0
        def signdiff(x, y): return (sign(x) * sign(y)) == -1

        if side_bias < 0:
            bias = 'Leftward'
        elif side_bias > 0:
            bias = 'Rightward'
        else:
            bias = 'Neutral'

        if adj_val == 0:  # No adjustment value set
            self.log_info('Assigning adjustment value for first time')
            if side_bias == 0:
                consecutive_good_sessions += 1
            else:
                adj_val += 5*sign(side_bias)
                consecutive_good_sessions = 0
        elif abs(side_bias) > 0.2:
            if signdiff(adj_val, side_bias):
                self.log_info('Adjustment value has overshot bias')
                self.log_info('This was a good session (contrary side bias)')
                consecutive_good_sessions += 1
            else:
                self.log_info(
                    f'{bias} bias detected. Adjustment value updated.')
                adj_val += 5*sign(side_bias)
                consecutive_good_sessions = 0
        else:
            self.log_info('This was a good session (no side bias detected)')
            consecutive_good_sessions += 1

        self.log_info('Adjustment Value is ' + str(adj_val))
        self.log_info('Consecutive good sessions is ' +
                      str(consecutive_good_sessions))
        adj_val = max(-40, min(40, adj_val))
        global_paradigm_data['adjustmentvalue'] = adj_val
        stage_data['consecutive_good'] = consecutive_good_sessions


@dataclass
class AltWheelStage2B3(ResponseAbstractStage):
    """
    Stage 2: alternating-target bandit with adaptive threshold correction.

    Runs a standard bandit task and applies a three-phase threshold adjustment
    algorithm (correction → return → complete) to correct side bias while
    keeping wheel thresholds within bounds.
    """
    slug: str = 'Stage3Return'

    def define_task(self, stage_data):
        """Run the alternating bandit task with 100/0 reward probabilities."""
        task = BanditTask(parent=self,
                          reward_prob_target=100,
                          reward_prob_offtarget=0,
                          rescue_trial_enabled=True,
                          taskblocks_enabled=True,
                          reward_delay_s=0,
                          enable_wheel_lock=True
                          )
        task.run()
        del task

    def define_shaping(self, stage_data):
        consecutive_good_sessions = stage_data.get(
            'consecutive_good', 0)
        global_paradigm_data = self.get_globalparadigmdata()
        adj_val = global_paradigm_data.get('adjustmentvalue', 0)
        side_bias = self._compute_sb_metric()

        if side_bias is None or self.session.duration < 30:
            return

        # If side bias is against adjustment value, return by 2
        def sign(x): return 1 if x > 0 else -1 if x < 0 else 0
        def signdiff(x, y): return (sign(x) * sign(y)) == -1

        if side_bias < 0:
            bias = 'Leftward'
        elif side_bias > 0:
            bias = 'Rightward'
        else:
            bias = 'Neutral'

        if (adj_val == 0 and (-0.15 < side_bias < 0.15)
                and self.session.merit >= 150):
            self.log_info(
                'No/low bias and sufficient performance for good session')
            consecutive_good_sessions += 1
        else:
            self.log_info('Bias or low performance detected')
            consecutive_good_sessions = 0

        if signdiff(side_bias, adj_val) or abs(side_bias) < 0.1:
            self.log_info(f'No/opposite bias at adjustment score {adj_val}')
            adj_val = adj_val - (sign(adj_val) * min(abs(adj_val), 2))

        self.log_info('Adjustment Value is ' + str(adj_val))
        self.log_info('Consecutive good sessions is ' +
                      str(consecutive_good_sessions))
        adj_val = max(-40, min(40, adj_val))
        global_paradigm_data['adjustmentvalue'] = adj_val
        stage_data['consecutive_good'] = consecutive_good_sessions


@dataclass
class BanditTrainingStageB3(ResponseAbstractStage):
    """
    Full bandit training stage with the default reward probability schedule.
    """
    slug: str = 'Stage5BanditTraining'
    _task: object = None

    def define_task(self, stage_data):
        """Run the bandit task with rescue trials and retain a reference for analysis."""
        task = BanditTask(parent=self,
                          rescue_trial_enabled=True,
                          reward_delay_s=0,
                          enable_wheel_lock=True)
        task.run()
        del task

    def define_shaping(self, stage_data):
        consecutive_good_sessions = stage_data.get(
            'consecutive_good', 0)

        if self.session.duration < 30:
            return

        if self.session.merit >= 150:
            self.log_info('This was a good session (merit >= 150)')
            consecutive_good_sessions += 1
        else:
            consecutive_good_sessions = 0

        self.log_notice('Consecutive good sessions is ' +
                        str(consecutive_good_sessions))
        stage_data['consecutive_good'] = consecutive_good_sessions


@dataclass
class BanditEndStageB3(ResponseAbstractStage):
    """
    Final bandit stage run without rescue trials.
    """
    slug: str = 'Stage6Bandit'

    def define_task(self, stage_data):
        """Run the bandit task without rescue trials."""
        task = BanditTask(parent=self,
                          rescue_trial_enabled=False,
                          enable_wheel_lock=True)
        task.run()
        del task

    def define_shaping(self, stage_data):
        pass

@dataclass
class EXPeriment_LaserInhibitionRandom20(BanditTask):
    slug : str = 'EXPeriment_LaserInhibitionRandom20'
    tags : list = field(default_factory= lambda : ['experiment'])
    rescue_trial_enabled : bool = False
    enable_wheel_lock : bool = True
    
    reward_delay_s : float = 0.2
    
    enable_laser_trials: bool = True
    laser_trial_sequence_type: Literal['response', 'trial'] = 'response'
    response_laser_enabled: bool = True
    laser_trial_sequence: list = None
    response_laser_delay_s: float = 0.1
    laserstop_trialend_offramp_enabled: bool = True
    laserstop_trialend_offramp_ms: int = 500
    
    percentage_trials : int = 20
    def setup(self):
        super().setup()
        self.laser_trial_sequence = self.nonconsecutive_trial_sequence(
            self.percentage_trials,
            20,50
            )
        self.log_notice('Attempting to initialize rodent face-cam')
        self.rig.initialize_mousecam()
        self.log_notice('Allowing 6 seconds for autoexposure calibration...')
        time.sleep(6)
    
    @staticmethod
    def nonconsecutive_trial_sequence(percentage = 25,
                                      blocks = 20,
                                      blocksize = 50):
        perc = percentage/100
        num_true = round(blocksize * perc)
        bsize = blocksize - 1
        true_positions = set()
        for block in range(blocks):
            offset = block * blocksize
            positions = random.sample(range(bsize-num_true + 1), num_true)
            for i, pos in enumerate(sorted(positions)):
                true_positions.add(
                    pos+i+offset
                    )
        return [i in true_positions for i in range(blocks*blocksize)]

@dataclass
class EXPeriment_LaserInhibitionFullTrial_Random20(EXPeriment_LaserInhibitionRandom20):
    slug : str = 'EXPeriment_LaserInhibitionFullTrial_Random20'
    tags : list = field(default_factory= lambda : ['experiment'])
    
    response_laser_enabled: bool = False
    laser_trial_sequence_type: Literal['response', 'trial'] = 'trial'
    pretrial_laser_enabled: bool = True
    laserstop_trialend_offramp_enabled:bool = True
    laserstop_trialend_offramp_ms: int = 500
    percentage_trials : int = 20

@dataclass
class EXPeriment_LaserInhibitionCueResponse_Random25(EXPeriment_LaserInhibitionRandom20):
     slug : str = 'EXPeriment_LaserInhibitionCueResponse_Random25'
     tags : list = field(default_factory= lambda : ['experiment'])
     
     response_laser_enabled: bool = False
     laserstop_trialend_offramp_enabled:bool = False
     laser_trial_sequence_type: Literal['response', 'trial'] = 'trial'
     
     precue_laser_enabled: bool = True
     precue_laser_offset: float = 0.3
     pt_trial_delay:float = 0.3
     iti1_s: float = 0.7
     response_laser_delay_s : float = 0.1
     laserstop_response_offramp_enabled: bool = True
     laserstop_response_offramp_ms: int = 100
     
     percentage_trials : int = 33
     
@dataclass
class EXPeriment_LASERCUE_ZERO(EXPeriment_LaserInhibitionRandom20):
     slug : str = 'EXPeriment_LASERCUE_ZERO'
     tags : list = field(default_factory= lambda : ['experiment'])
     
     response_laser_enabled: bool = False
     laserstop_trialend_offramp_enabled:bool = False
     laser_trial_sequence_type: Literal['response', 'trial'] = 'trial'
     
     precue_laser_enabled: bool = True
     precue_laser_offset: float = 0
     pt_trial_delay:float = 0
     iti1_s: float = 1
     response_laser_delay_s : float = 0.1
     laserstop_response_offramp_enabled: bool = True
     laserstop_response_offramp_ms: int = 100
     
     percentage_trials : int = 33

@dataclass
class EXPeriment_LaserControlSession(BanditTask):
    slug : str = 'EXPeriment_LaserControlSession'
    tags : list = field(default_factory= lambda : ['experiment'])
    rescue_trial_enabled : bool = False
    enable_wheel_lock : bool = True
    
    reward_delay_s : float = 0.2
    
    enable_laser_trials: bool = False
    def setup(self):
        super().setup()
        self.log_notice('Attempting to initialize rodent face-cam')
        self.rig.initialize_mousecam()
        self.log_notice('Allowing 6 seconds for autoexposure calibration...')
        time.sleep(6)

@dataclass
class EXPeriment_LaserControlSessionNOCAMERA(BanditTask):
    slug : str = 'EXPeriment_LaserControlSessionNOCAMERA'
    tags : list = field(default_factory= lambda : ['experiment'])
    rescue_trial_enabled : bool = False
    enable_wheel_lock : bool = True
    
    reward_delay_s : float = 0.2
    
    enable_laser_trials: bool = False
    def setup(self):
        super().setup()
        self.log_notice('Skipping rodent face-cam')
        time.sleep(6)
        

@dataclass
class EXPeriment_BanditForPhotom(BanditTask):
    slug : str = 'EXPeriment_BanditForPhotom'
    tags : list = field(default_factory= lambda : ['experiment'])
    rescue_trial_enabled : bool = False
    enable_wheel_lock : bool = True
    
    reward_delay_s : float = 0.2
    
    enable_laser_trials: bool = False
    
    def setup(self):
        super().setup()
        self.log_notice('Attempting to initialize rodent face-cam')
        self.rig.initialize_mousecam()
        self.log_notice('Allowing 6 seconds for autoexposure calibration...')
        time.sleep(6)

@dataclass
class EXPeriment_AltChoiceForPhotom(BanditTask):
    slug : str = 'EXPeriment_BanditForPhotom'
    tags : list = field(default_factory= lambda : ['experiment'])
    
    reward_prob_target : int=100,
    reward_prob_offtarget : int=0,
    rescue_trial_enabled : bool=False,
    taskblocks_enabled: bool=True,
    reward_delay_s: float=0.2,
    enable_wheel_lock: bool=True
    
    reward_delay_s : float = 0.2
    
    enable_laser_trials: bool = False
    
    def setup(self):
        super().setup()
        self.log_notice('Attempting to initialize rodent face-cam')
        self.rig.initialize_mousecam()
        self.log_notice('Allowing 6 seconds for autoexposure calibration...')
        time.sleep(6)
    

@dataclass
class BanditParadigm3(Paradigm):
    slug: str = 'BanditParadigm3'
    stagelist: tuple = (HabituationStage,
                        AnyWheelStageB3,
                        AltWheelStage1B3,
                        AltWheelStage2B3,
                        BanditTrainingStageB3,
                        BanditEndStageB3)

    def define_progression(self, current_stage, stage_data):
        if current_stage is HabituationStage:
            if stage_data.get('consecutive_good', 0) >= 2:
                self.progress()
        elif current_stage is AnyWheelStageB3:
            if stage_data.get('consecutive_good', 0) >= 1:
                self.progress()
        elif current_stage is AltWheelStage1B3:
            if stage_data.get('consecutive_good', 0) >= 1:
                self.progress()
        elif current_stage is AltWheelStage2B3:
            if stage_data.get('consecutive_good', 0) >= 1:
                self.progress()
        elif current_stage is BanditTrainingStageB3:
            if stage_data.get('consecutive_good', 0) >= 2:
                self.progress()
        elif current_stage is BanditEndStageB3:
            pass
```
