## Extended Data for "A Low-Cost, Modular Hardware and Software Platform for Head-Fixed Mouse Decision-Making Tasks": test.py

```
# -*- coding: utf-8 -*-
"""
Created on Mon Jun  8 12:28:43 2026

@author: mm4114
"""

import time
import random
from copy import deepcopy
from typing import ClassVar, Literal, List, Tuple

from lampyr.segments import Trial, Task, BehaviorSegment, TrialToTask
from lampyr.segments.paradigm import Stage, Paradigm
from dataclasses import dataclass, field

# -------------- Define Event Callbacks --------------
# Simple events simply log a timepoint, complex events can also interact with the rig and log reports as needed.


def event_response(self : BehaviorSegment):
    """Play the response registration tone and log the event"""
    self.log_debug('Sending play response tone command to rig')
    self.rig.play.responsetone()

# -------------- Define Habituation Trial/Task --------------


@dataclass
class RotaryTest(Trial):
    """
    A single habituation trial.

    1. Wait for a fixed ITI (``iti1_dur``).
    2. Deliver reward and trigger the reward event.
    3. Monitor licks for ``reward_consumption_period_s`` seconds.  If no lick is detected, extend monitoring for ``reward_consumption_nolick_delay_s`` seconds.
    4. Log whether the reward was consumed (lick detected) and count merit/abstention accordingly.
    5. Wait for a second ITI (``iti2_dur``) before finishing the trial.

    Attributes
    ----------
    iti1_dur : float
        Pre-reward inter-trial interval in seconds.
    reward_consumption_period_s : float
        Primary lick-detection window after reward delivery (seconds).
    reward_consumption_nolick_delay_s : float
        Extended wait period if no lick detected in the primary window (seconds).
    count_merits : bool
        If ``True``, log merit when the reward is consumed.
    iti2_dur : float
        Post-consumption inter-trial interval in seconds.
    """

    def setup(self):
        """Register the reward event callback and description."""
        self.register_event('response',
                            callback=event_response,
                            description='Water reward given')

    def perform(self):
        """
        Execute one habituation trial iteration: ITI → reward → consumption → ITI.
        """
        # Start behavior
        stime = time.time()
        self.waitfor(
            condition=lambda: self.rig.wheel.movement_total_since(stime) > 359,
            timeout=None,
            poll_interval=0.01,
            while_waiting=lambda: self.log_notice(
                'No Response yet detected'),
            while_waiting_interval=4
        )
        self.notify('Detected Full Revolution')
        self.trigger_event('response')
        self.wait(5)


@dataclass
class RotaryTestTask(Task):
    """
    Task that repeatedly runs :class:`RotaryTest` instances until the
    session stop conditions are met.
    """
    slug: str = 'RotaryTest'

    def setup(self):
        pass

    def loop(self):
        """Create and run one :class:`HabituationTrial` per iteration."""
        trial = RotaryTest(parent=self)
        trial.run()
        del trial

@TrialToTask
@dataclass
class LaserTest(Trial):
    slug : str = 'LaserTest'
    def setup(self):
        pass
    
    def perform(self):
        self.log_notice('LASER ON')
        self.rig.laser.begin()
        self.wait(20)
        self.log_notice('LASER OFF')
        self.rig.laser.stop()
        self.wait(2)
```
