## Extended Data for "A Low-Cost, Modular Hardware and Software Platform for Head-Fixed Mouse Decision-Making Tasks": config.py

```
# -*- coding: utf-8 -*-
"""
Created on Mon Aug 25 19:47:06 2025

@author: mm4114
"""
from lampyr.version import __version__

import os, json, time
from copy import deepcopy

class ConfigFile:
    """
    Persistent JSON-backed configuration store with dot-path access.

    Merges a supplied default configuration with any values found on disk,
    so that new keys are always available while user-set values are preserved.
    """

    def __init__(self, default_config : dict, fp : str):
        """
        Load configuration from disk, filling missing keys from ``default_config``.

        Parameters
        ----------
        default_config : dict
            Nested dict of default values.  Any key absent from the on-disk
            file is supplied from here.
        fp : str
            Path to the JSON file to load from and save to.  Created on the
            first :meth:`save` call if it does not yet exist.
        """
        self._default = deepcopy(default_config)
        self._syncfp = fp
        self._load()

    def _load(self):
        loaded_config = {}
        if os.path.exists(self._syncfp):
            try:
                with open(self._syncfp, 'r') as f:
                    loaded_config = json.load(f)
            except Exception:
                pass
        self._config = self._merge_configs(deepcopy(self._default), loaded_config)

    def reload(self):
        """Re-read config from disk, picking up out-of-process changes."""
        self._load()

    def _merge_configs(self, default, loaded):
        """
        Recursively merge ``default`` into ``loaded``, adding missing keys.

        Parameters
        ----------
        default : dict
            Source of default values.
        loaded : dict
            User-supplied values that take precedence over defaults.

        Returns
        -------
        dict
            ``loaded`` dict with missing keys populated from ``default``.
        """
        for key, value in default.items():
            if key in loaded and isinstance(value, dict) and isinstance(loaded[key], dict):
                loaded[key] = self._merge_configs(value, loaded[key])
            elif key not in loaded:
                loaded[key] = value
        return loaded

    def get(self, key_path):
        """
        Retrieve a configuration value by dot-separated key path.

        Parameters
        ----------
        key_path : str
            Dot-separated path into the config dict
            (e.g. ``'lampyr.mice_directory'``).

        Returns
        -------
        object
            A deep copy of the value at ``key_path``.

        Raises
        ------
        KeyError
            If any segment of ``key_path`` does not exist in the config.
        """
        keys = key_path.split('.')
        current = self._config
        for key in keys:
            if isinstance(current, dict) and key in current:
                current = current[key]
            else:
                raise KeyError(f"Key not found: {key_path}")
        return deepcopy(current)

    def set(self, key_path, value):
        """
        Set a configuration value by dot-separated key path and persist to disk.

        Parameters
        ----------
        key_path : str
            Dot-separated path into the config dict
            (e.g. ``'rig.calibrated'``).
        value : object
            Value to store at ``key_path``.

        Raises
        ------
        KeyError
            If any intermediate segment of ``key_path`` does not exist.
        """
        keys = key_path.split('.')
        current = self._config
        for key in keys[:-1]:
            if isinstance(current, dict) and key in current:
                current = current[key]
            else:
                raise KeyError(f"Key path not found: {key_path}")
        current[keys[-1]] = value
        self.save()
    
    def save(self):
        """
        Persist the current configuration to the backing JSON file.
        """
        with open(self._syncfp, 'w') as file:
            json.dump(self.to_dict(), file, indent = 2)
    
    def to_dict(self):
        """
        Return a deep copy of the full configuration as a plain dict.

        Returns
        -------
        dict
            Deep copy of the internal config dictionary.
        """
        return deepcopy(self._config)


class Config(ConfigFile):
    """
    Application-level configuration for Lampyr, backed by the user's AppData.

    Extends :class:`ConfigFile` with a fixed path
    (``%LOCALAPPDATA%/lampyr/config.json``), a well-known default schema,
    and helpers for loading secondary config files from both the local AppData
    directory and the shared mice directory.
    """

    _APP_DATA_DIR = os.path.join(os.getenv('LOCALAPPDATA'), 'lampyr')
    _CONFIG_FILE_PATH = os.path.join(_APP_DATA_DIR, 'config.json')

    DEFAULT_CONFIG = {
        'lampyr': {
            'configured': False,
            'mice_directory': 'N:/SHARED/Maxwell_Lampyr_MouseData',
            'plugin_folder': None,
            'enable_saveload_failsafe': True,
            'enable_local_mouse_backups' : True
        },
        'rig': {
            'name': None,
            'rig_type' : 'BanditRig',
            'rig_type_last_load' : None, #not implemented but will eventually force configuration
            'configuration' : {}, #implicit area for persistant rig data
            'calibrated': 0, #To be removed
            'configured': False, #To be removed
            'sipper_calib': 10000 #To be removed
        },
        'notifications': {}
    }

    def __init__(self):
        """
        Initialise the application config, creating AppData directory if needed.

        Also stamps the current Lampyr version into the config on every
        startup so that ``lampyr.version`` always reflects the installed
        package.
        """
        os.makedirs(self._APP_DATA_DIR, exist_ok=True)
        super().__init__(self.DEFAULT_CONFIG, self._CONFIG_FILE_PATH)
        self.set('lampyr.version', __version__)
    
    def load_extended_config(self, key, default = {}):
        """
        Load (or create) a secondary config file from the local AppData directory.

        Parameters
        ----------
        key : str
            Filename stem; the file will be
            ``%LOCALAPPDATA%/lampyr/<key>.json``.
        default : dict, optional
            Default config dict used if the file does not exist.

        Returns
        -------
        ConfigFile
            A :class:`ConfigFile` instance backed by the requested file.
        """
        fp = os.path.join(self._APP_DATA_DIR, f'{key}.json')
        return ConfigFile(default, fp)

    def load_shared_extended_config(self, key, default = {}):
        """
        Load (or create) a secondary config file from the shared mice directory.

        Parameters
        ----------
        key : str
            Filename stem; the file will be
            ``<lampyr.mice_directory>/<key>.json``.
        default : dict, optional
            Default config dict used if the file does not exist.

        Returns
        -------
        ConfigFile
            A :class:`ConfigFile` instance backed by the requested file in the
            shared directory.
        """
        shared_dir = self.get('lampyr.mice_directory')
        fp = os.path.join(shared_dir, f'{key}.json')
        return ConfigFile(default, fp)
```
