## Extended Data for "A Low-Cost, Modular Hardware and Software Platform for Head-Fixed Mouse Decision-Making Tasks": files.py

```
# -*- coding: utf-8 -*-
"""
Created on Thu Sep 18 11:48:06 2025

@author: mm4114
"""


import h5py
import os
import numpy as np
import pickle
import json
import glob
from dataclasses import is_dataclass, asdict, fields
from lampyr.primatives import Session, Mouse
from lampyr.config import Config
import shutil
import csv
import time
import hashlib

from typing import Union, List


def savejson(fp, data, saveasis=True):
    """
    Serialise data to a JSON file.

    Parameters
    ----------
    fp : str or os.PathLike
        Destination file path.
    data : object
        Data to serialise.  Dataclasses are converted with ``asdict`` if
        ``saveasis=False``.
    saveasis : bool, optional
        If ``True`` (default), write ``data`` directly.  If ``False`` and
        ``data`` is a dataclass, convert it first.

    Returns
    -------
    str or os.PathLike
        The path ``fp`` that was written to.
    """
    if not saveasis:
        if is_dataclass(data):
            data = asdict(data)
    with open(fp, 'w') as f:
        json.dump(data, f, indent=2)
    return fp


def loadjson(fp):
    """
    Load a JSON file and return the parsed object.

    Parameters
    ----------
    fp : str or os.PathLike
        Path to the JSON file.

    Returns
    -------
    object
        Parsed JSON content.
    """
    try:
        with open(fp, 'r') as f:
            return json.load(f)
    except Exception:
        print(fp)
        raise


def savecsv(fp, data: List[dict]):
    """
    Write a list of dicts to a CSV file.

    All keys found across all dicts are used as column headers.  Missing
    values are left blank.

    Parameters
    ----------
    fp : str or os.PathLike
        Destination file path.
    data : list of dict
        Rows to write; each dict maps column name to value.
    """
    all_keys = []
    for entry in data:
        for key in entry.keys():
            if key not in all_keys:
                all_keys.append(key)

    with open(fp, 'w', newline='') as f:
        writer = csv.DictWriter(f, fieldnames=all_keys)
        writer.writeheader()
        writer.writerows(data)
        
def loadcsv(fp):
    """
    Load a CSV file and return a list of row dicts.

    Parameters
    ----------
    fp : str or os.PathLike
        Path to the CSV file.

    Returns
    -------
    list of dict
        One dict per row, keyed by column header.
    """
    data = []
    with open(fp, 'r') as f:
        reader = csv.DictReader(f)
        for row in reader:
            data.append(row)
    return data


def savepickle(fp, data):
    """
    Pickle and save ``data`` to a binary file.

    Parameters
    ----------
    fp : str or os.PathLike
        Destination file path.
    data : object
        Object to pickle.

    Returns
    -------
    str or os.PathLike
        The path ``fp`` that was written to.
    """
    with open(fp, 'wb') as f:
        pickle.dump(data, f)
    return fp


def loadpickle(filepath):
    """
    Load and return a pickled object from a binary file.

    Parameters
    ----------
    filepath : str or os.PathLike
        Path to the pickle file.

    Returns
    -------
    object
        The unpickled object.
    """
    with open(filepath, 'rb') as f:
        data = pickle.load(f)
    return data


def loadh5(fp):
    """
    Load an HDF5 file written by :func:`saveh5` and reconstruct its Python object.

    Recursively traverses the HDF5 group structure, using the ``'_type'``
    attribute stored by :func:`saveh5` to reconstruct dicts, lists, ndarrays,
    and scalars.  Falls back to heuristic type detection for files written
    without ``'_type'`` attributes.

    Parameters
    ----------
    fp : str or os.PathLike
        Path to the ``.h5`` file.

    Returns
    -------
    object or None
        The reconstructed Python object rooted at the ``'root'`` HDF5 group,
        or ``None`` if no ``'root'`` group exists.
    """
    def recursive_load(h5group):
        # Get the type hint saved by saveh5
        node_type = h5group.attrs.get('_type')
        if node_type == 'ndarray':
            return h5group['array'][()]
        elif node_type == 'scalar' or node_type == 'unknown_fallback_str':
            val = h5group.attrs['value']
            original_py_type = h5group.attrs.get('_original_py_type')

            # Attempt to convert back to original Python type
            if original_py_type == 'int':
                return int(val)
            elif original_py_type == 'float':
                return float(val)
            elif original_py_type == 'bool':
                # Handle boolean string conversion carefully
                if isinstance(val, str):
                    return val.lower() == 'true'
                return bool(val)  # For actual boolean values saved directly
            # Add more types if needed (e.g., complex, bytes)
            # elif original_py_type == 'str_fallback': # This means it was an unhandled type saved as string
            #     # You might want to log a warning here
            #     pass
            return val  # Default for string or if type conversion not handled

        elif node_type == 'dict':
            keys = list(h5group.keys())
            return {k: recursive_load(h5group[k]) for k in keys}
        elif node_type == 'list':  # List or tuple, using the explicit flag
            keys = list(h5group.keys())
            indices = sorted(int(k) for k in keys)
            # Ensure it's a contiguous list of indices for true list reconstruction
            if len(indices) == len(keys) and all(idx == i for i, idx in enumerate(indices)):
                return [recursive_load(h5group[str(i)]) for i in indices]
            else:
                # Fallback to dict if keys are not contiguous for a list
                print(f"Warning: List expected for group '{
                      h5group.name}', but keys are not contiguous. Loading as dictionary.")
                return {k: recursive_load(h5group[k]) for k in keys}
        else:
            # Fallback for old files or unexpected structures
            # If no '_type' attribute, try to guess based on existing children/attributes
            if 'array' in h5group:
                return h5group['array'][()]
            elif 'value' in h5group.attrs:
                return h5group.attrs['value']
            # Assume it's a group that might be a dict or list (try dict first for safety)
            else:
                keys = list(h5group.keys())
                # Try to guess if it's a list by checking if all keys are integer strings
                try:
                    parsed_keys = sorted(int(k) for k in keys)
                    if len(parsed_keys) > 0 and all(pk == i for i, pk in enumerate(parsed_keys)):
                        return [recursive_load(h5group[str(i)]) for i in parsed_keys]
                    else:
                        return {k: recursive_load(h5group[k]) for k in keys}
                except ValueError:  # Not all keys are integers, so definitely a dict
                    return {k: recursive_load(h5group[k]) for k in keys}

    with h5py.File(fp, mode='r') as file:
        if 'root' in file:
            return recursive_load(file['root'])
        else:
            print(f"Warning: 'root' group not found in HDF5 file: {
                  fp}. Returning None.")
            return None  # Or raise an error


def saveh5(fp, data):
    """
    Recursively save a Python object to an HDF5 file.

    Supports dicts, lists/tuples, numpy arrays, and scalars (int, float, str,
    bool, numpy number types).  Unsupported types are stored as their
    ``str()`` representation with a ``'unknown_fallback_str'`` type tag.
    Dataclasses are converted with ``asdict`` before saving.

    Parameters
    ----------
    fp : str or os.PathLike
        Destination ``.h5`` file path.
    data : object
        Python object to save.

    Returns
    -------
    str or os.PathLike
        The path ``fp`` that was written to.
    """
    def recursive_save(obj, h5group):
        if isinstance(obj, dict):
            h5group.attrs['_type'] = 'dict'  # Indicate it's a dictionary
            for k, v in obj.items():
                grp = h5group.create_group(str(k))
                recursive_save(v, grp)
        elif isinstance(obj, (list, tuple)):
            h5group.attrs['_type'] = 'list'  # Indicate it's a list/tuple
            for i, v in enumerate(obj):
                grp = h5group.create_group(str(i))
                recursive_save(v, grp)
        elif isinstance(obj, np.ndarray):
            h5group.attrs['_type'] = 'ndarray'  # Indicate it's an ndarray
            h5group.create_dataset('array', data=obj)
        # Added bool for explicit type saving
        elif isinstance(obj, (int, float, str, bool, np.number)):
            h5group.attrs['_type'] = 'scalar'
            h5group.attrs['value'] = obj
            # Store original Python type name for better reconstruction
            h5group.attrs['_original_py_type'] = type(obj).__name__
        else:
            # Fallback for unhandled types (e.g., set, complex, custom objects)
            # Save as string, mark as fallback
            h5group.attrs['_type'] = 'unknown_fallback_str'
            h5group.attrs['value'] = str(obj)
            h5group.attrs['_original_py_type'] = 'str_fallback'

    if is_dataclass(data):
        data = asdict(data)

    with h5py.File(fp, mode='w') as file:
        # Changed to pass data first
        recursive_save(data, file.create_group("root"))

    return fp


def savesessionfile(session: Session,
                    dir_fp: Union[str, os.PathLike] = os.getcwd()
                    ) -> List[os.PathLike]:
    """
    Save a Session object in Lampyr format within a specific directory.

    This function saves two files:
    - A `.lampyr.h5` file containing rig data in HDF5 format.
    - A `.lampyr.json` file containing the rest of the session metadata.

    Parameters
    ----------
    session : Session
        Lampyr Session object to save.
    dir_fp : Union[str, os.PathLike], optional
        Directory in which to save session files. Defaults to current working directory.

    Returns
    -------
    fps : List[os.PathLike]
        List of file paths to the saved files (HDF5 and JSON).
    """
    os.makedirs(dir_fp, exist_ok=True)
    rigdata = session.rigdata
    np_rigdata = {}
    fps = []
    if rigdata is not None:
        for reporttype, reports in rigdata.items():
            np_rigdata[reporttype] = {}
            for datalabel, data in reports.items():
                np_rigdata[reporttype][datalabel] = np.asarray(data)
        fp = saveh5(os.path.join(dir_fp, f'{session.uniquesessionid}.lampyr.h5'),
                    np_rigdata)
        if fp is not None:
            fps.append(fp)
    data = asdict(session)
    data['rigdata'] = None
    fp = savejson(os.path.join(dir_fp, f'{session.uniquesessionid}.lampyr.json'),
                  data)
    if fp is not None:
        fps.append(fp)
    return fps


def loadsessionfile(sessionid: str,
                    dir_fp: Union[str, os.PathLike] = os.getcwd()
                    ) -> Session:
    """
    Load a Lampyr session object from saved HDF5 and JSON files.

    This function reconstructs a Session object using the `.lampyr.h5` and
    `.lampyr.json` files stored in the specified directory. The rig data is
    loaded from the HDF5 file, and all other attributes are restored from
    the JSON file.

    Parameters
    ----------
    sessionid : str
        The unique session ID corresponding to the saved session files.
    dir_fp : Union[str, os.PathLike], optional
        Directory where the session files are located. Defaults to current working directory.

    Raises
    ------
    FileExistsError
        If the directory or required session files do not exist.

    Returns
    -------
    Session
        A reconstructed Lampyr Session object.
    """
    # Check if dir_fp exists
    if not os.path.exists(dir_fp):
        raise FileExistsError(f'{dir_fp} does not exist')

    # Determine the h5 and json filepaths
    h5file = os.path.join(dir_fp,
                          f'{sessionid}.lampyr.h5')
    jsonfile = os.path.join(dir_fp,
                            f'{sessionid}.lampyr.json')

    # Check if the h5 and json files exist
    if not os.path.exists(h5file) or not os.path.exists(jsonfile):
        raise FileExistsError(f'Session is invalid or does not exist. {
                              h5file} or {jsonfile} not found.')

    # Load all data and input into a session object, saves the endtime last
    # to avoid premature locking of session data
    session = Session(rigdata=loadh5(h5file))
    jsonobj = loadjson(jsonfile)
    for index, value in jsonobj.items():
        if index not in ['endtime', 'rigdata']:
            session.__setattr__(index, value)
    session.endtime = jsonobj['endtime']
    return session


def savemousefile(mouse: Mouse,
                  dir_fp: Union[str, os.PathLike] = os.getcwd()
                  ) -> List[os.PathLike]:
    """
    Save a Mouse object in Lampyr format within a specific directory.

    This function saves two files:
    - A `.lampyr.json` file containing mouse metadata.
    - A `.lampyr.csv` file containing mouse session history (if available).

    Parameters
    ----------
    mouse : Mouse
        A Lampyr Mouse object to be saved. Contains metadata and session history.
    dir_fp : Union[str, os.PathLike], optional
        The directory in which to save the mouse files. Defaults to the current working directory.

    Returns
    -------
    List[os.PathLike]
        A list of file paths to the saved files (JSON and optionally CSV).
    """
    os.makedirs(dir_fp, exist_ok=True)

    mouse_json_fp = os.path.join(dir_fp, f'{mouse.mouseid}_mouse.lampyr.json')
    mouse_csv_fp = os.path.join(dir_fp, f'{mouse.mouseid}_history.lampyr.csv')

    mouse_metadata = asdict(mouse)
    mouse_history = mouse_metadata.pop('history', None)

    savejson(mouse_json_fp, mouse_metadata)

    if mouse.mouseid == 'UNKNOWN_MOUSE' or mouse_history is None:
        return [mouse_json_fp]

    savecsv(mouse_csv_fp, mouse_history)

    return [mouse_json_fp, mouse_csv_fp]


def loadmousefile(mouseid: str,
                  dir_fp: Union[str, os.PathLike] = os.getcwd()
                  ) -> Mouse:
    """
    Load a Lampyr Mouse object from saved JSON and CSV files.

    This function reconstructs a Mouse object using:
    - A `.lampyr.json` file containing mouse metadata.
    - A `.lampyr.csv` file containing mouse session history (if available).

    Parameters
    ----------
    mouseid : str
        The unique mouse ID corresponding to the saved files.
    dir_fp : Union[str, os.PathLike], optional
        Directory where the mouse files are stored. Defaults to the current working directory.

    Raises
    ------
    FileExistsError
        If the required JSON file does not exist in the specified directory.

    Returns
    -------
    Mouse
        A reconstructed Mouse object with metadata and optional session history.
    """
    if mouseid == 'UNKNOWN_MOUSE':
        return Mouse(mouseid=mouseid)
    
    mouse_json_fp = os.path.join(dir_fp, f'{mouseid}_mouse.lampyr.json')
    mouse_csv_fp = os.path.join(dir_fp, f'{mouseid}_history.lampyr.csv')
    
    if not os.path.exists(mouse_json_fp):
        raise FileExistsError(f'{mouse_json_fp} does not exist')
    
    try:
        mouse_data = loadjson(mouse_json_fp)
    except json.JSONDecodeError as e:
        e.add_note(f'Mouse {mouseid} has malformed json')
        raise e
    
    if os.path.exists(mouse_csv_fp):
        mouse_data['history'] = loadcsv(mouse_csv_fp)
        
    allowed = {f.name for f in fields(Mouse)}
    mouse_data = {k: v for k, v in mouse_data.items() if k in allowed}
    
    return Mouse(**mouse_data)
```
