## Extended Data for "A Low-Cost, Modular Hardware and Software Platform for Head-Fixed Mouse Decision-Making Tasks": app.py

```
# -*- coding: utf-8 -*-
"""
Created on Wed May 14 15:02:02 2025

@author: mm4114
"""

import click
import time
import os
import numpy as np
from lampyr import config, Lampyr
from lampyr.version import __version__
from lampyr.segments import Task, Trial, Stage, Paradigm
from lampyr.behaviors import bandit
from lampyr.behaviors import test
from lampyr.rigs.rigcontrol import ArduinoBanditRig_0, SerialMonitor
from lampyr import actions

@click.group()
@click.pass_context
def cli(ctx):
    ctx.obj = Lampyr()
    if (ctx.obj.config.get('lampyr.configured') is False
        and ctx.invoked_subcommand != 'configure'):
        raise click.UsageError('Lampyr is not configured. Run "lampyr configure"')
    @ctx.call_on_close
    def cleanup():
        ctx.obj.close()

@cli.command()
@click.pass_obj
def configure(lampyr : Lampyr):
    actions.printtitle('CONFIGURE LAMPYR')
    mouse_fp = click.prompt('Session/Mouse Data Path:')
    sl = click.prompt('Enable save/load failsafe(y/n)?\n') == 'y'
    mback = click.prompt('Enable local mouse file backup (y/n)?\n') == 'y'
    if click.prompt('Would you like to enable plugins (y/n)?\n') == 'y':
        plugin_fp = click.prompt('Plugin Folder Path:')
    else:
        plugin_fp = None
    lampyr.config.set('lampyr.mice_directory', mouse_fp)
    lampyr.config.set('lampyr.plugin_folder', plugin_fp)
    lampyr.config.set('lampyr.enable_saveload_failsafe', mouse_fp)
    lampyr.config.set('lampyr.enable_local_mouse_backups', mouse_fp)
    lampyr.config.set('lampyr.configured', True)
    click.echo('You have successfully configured your lampyr software')

def _origin(cls):
    """Return a short label for where a behaviour class was defined."""
    mod = cls.__module__
    if mod.startswith('plugin_'):
        return f'plugin:{mod.removeprefix("plugin_")}'
    return 'builtin'


@cli.command()
@click.pass_obj
def list(lampyr : Lampyr):
    actions.printtitle('LOADED BEHAVIORS')
    actions.printheader('PARADIGMS')
    for behav in Paradigm.get_children():
        click.echo(f'{behav.__name__:50}  [{_origin(behav)}]')
    actions.printheader('STAGES')
    for behav in Stage.get_children():
        click.echo(f'{behav.__name__:50}  [{_origin(behav)}]')
    actions.printheader('TASKS')
    for behav in Task.get_children():
        click.echo(f'{behav.__name__:50}  [{_origin(behav)}]')
    actions.printheader('TRIALS')
    for behav in Trial.get_children():
        click.echo(f'{behav.__name__:50}  [{_origin(behav)}]')

@cli.command()
@click.pass_obj
def reset(lampyr):
    passw = click.prompt("Type YES to confirm")
    if passw != 'YES':
        click.echo('Denied')
    else:
        lampyr.config.reset_to_default()
        click.echo("Configuration reset to default.")

@cli.command()
@click.pass_obj
def developer(lampyr):
    passw = click.prompt("Input Password")
    if passw != 'photuris':
        click.echo('Denied')
    else:
        lampyr.config.set('rig.calibrated', 100000000000)
        lampyr.config.set('rig.name', 'Photuris')
        lampyr.config.set('rig.configured', True)
        click.echo("Configuration set to developer mode.")
    
@cli.command()
@click.pass_obj
def info(lampyr):
    actions.printtitle(f'LAMPYR BEHAVIOR SOFTWARE v{__version__}')
    actions.printheader('CONFIG')
    info = lampyr.config._config
    actions.printinfo(info)

@cli.group()
@click.pass_obj
def rig(ctx):
    pass

@rig.command(name = 'info')
@click.pass_obj
def rig_info(lampyr):
    info = lampyr.config.get('rig')
    actions.printinfo(info)

@rig.command()
@click.pass_obj
def configure(lampyr):
    actions.configure_rig(lampyr)
        
@rig.command()
@click.pass_obj
def calibrate(ctx):
    ctx.rigmanager.calibrate()

@cli.group()
def mouse():
    pass

@mouse.command(name = 'create')
@click.argument('mouseid')
@click.option('--paradigm','-p', type = str, required = False)
@click.option('--force', is_flag = True, help='overwrite existing mouse')
@click.pass_obj
def mouse_create(lampyr : Lampyr, mouseid, force, **kwargs):
    if not force:
        if lampyr.mousemanager.exists(mouseid):
            click.echo(f'\nFailed to create mouse: Mouse name {mouseid} already exists.')
            return
        mousedir = os.path.join(lampyr.config.get('lampyr.mice_directory'),mouseid)
        if not os.path.exists(mousedir):
            os.makedirs(mousedir)
            click.echo(f'Created folder: {mousedir}')
    if kwargs['paradigm'] is not None:
        if kwargs['paradigm'] not in lampyr.paradigms:
            click.echo(f'\nFailed to create mouse: {kwargs["paradigm"]} Paradigm does not exist.')
            return
    lampyr.mousemanager.create(mouseid, **kwargs)
    
@mouse.command(name = 'retire')
@click.argument('mouseid')
@click.pass_obj
def mouse_retire(lampyr : Lampyr, mouseid):
    if lampyr.mousemanager.exists(mouseid):
        lampyr.mousemanager.load(mouseid)
        lampyr.mousemanager.retire()
    else:
        click.echo('Mouse does not exist')
        

@mouse.command(name = 'list')
@click.option('-recent', 'recent_only', is_flag=True, default=False, help='Only show recently run mice')
@click.option('-all', 'all', is_flag=True, default=False, help='Show all mice, including retired mice')
@click.pass_obj
def mouse_list(lampyr, recent_only, all):
    def _time_ago(ts):
        delta = int(time.time() - ts)
        if delta < 60:
            return f"{delta}s ago"
        elif delta < 3600:
            return f"{delta // 60}m ago"
        elif delta < 86400:
            h = delta // 3600
            m = (delta % 3600) // 60
            return f"{h}h {m}m ago"
        else:
            d = delta // 86400
            h = (delta % 86400) // 3600
            return f"{d}d {h}h ago"

    mids, _ = lampyr.datamanager.mouselist()
    rows = []
    for mid in mids:
        mouse = lampyr.datamanager.loadmouse(mid)
        if mouse.retired and not all:
            continue
        paradigm = mouse.paradigm or 'none'
        if mouse.paradigm is not None:
            paradigmslug = lampyr.behaviors.get(paradigm).slug
            paradigm_data = mouse.properties.get(paradigmslug, {})
            stage = paradigm_data.get('stage', 'pending...')
        else:
            stage = '-'
        time_since_last = 9999999999999
        if mouse.history:
            last_entry = mouse.history[-1]
            last = _time_ago(float(last_entry['starttime']))
            time_since_last = int(time.time() - float(last_entry['starttime']))
            merit = str(last_entry.get('merit', ''))
            rewards = str(last_entry.get('rewards', ''))
        else:
            last, merit, rewards = 'never', '', ''
        if recent_only and time_since_last > 43200:
            continue
        rows.append((mid, paradigm, stage, last, merit, rewards))
    rows = sorted(rows, key = lambda x : x[0])
    headers = ('NAME', 'PARADIGM', 'STAGE', 'LAST SESSION', 'MERIT', 'REWARDS')
    widths = [max(len(h), max((len(r[i]) for r in rows), default=0))
              for i, h in enumerate(headers)]

    actions.printheader('MICE')
    click.echo('  '.join(h.ljust(widths[i]) for i, h in enumerate(headers)))
    click.echo('  '.join('-' * w for w in widths))
    for row in rows:
        click.echo('  '.join(row[i].ljust(widths[i]) for i in range(len(headers))))
    click.echo('\n*Note: Paradigm and stage may not correspond to last session')

@mouse.command(name = 'info')
@click.argument('mouseid')
@click.pass_obj
def mouse_info(lampyr, mouseid):
    lampyr.mousemanager.load(mouseid)
    info = lampyr.mouse
    click.echo(info)

@mouse.command(name = 'run')
@click.argument('mouseid')
@click.argument('behavior', required = False)
@click.option('--merit_limit', '-ml', type=int, required=False)
@click.option('--merit_min', '-mm', type=int, required=False)
@click.option('--demerit_limit', '-dml', type=int, required=False)
@click.option('--demerit_min', '-dmm', type=int, required=False)
@click.option('--duration_limit', '-dl', type=int, required=False)
@click.option('--duration_min', '-dm', type=int, required=False)
@click.option('--trial_limit', '-tl', type=int, required=False)
@click.option('--trial_min', '-tm', type=int, required=False)
@click.option('--reward_limit', '-rl', type=int, required=False)
@click.option('--reward_min', '-rm', type=int, required=False)
@click.option('--abstention_limit', '-al', type=int, required=False)
@click.option('--abstention_min', '-am', type=int, required=False)
@click.option('--participation_limit', '-pl', type=int, required=False)
@click.option('--participation_min', '-pm', type=int, required=False)
@click.option('--serial_abstention_limit', '-sal', type=int, required=False)
@click.option('--serial_abstention_min', '-sam', type=int, required=False)
@click.pass_obj
def mouse_run(lampyr, mouseid, behavior, **kwargs):
    try:
        lampyr.mousemanager.load(mouseid)
    except KeyError as e:
        click.echo(f'Error loading mouse: {e}')
        return
    if behavior is None:
        behavior = lampyr.mouse.paradigm
    try:
        actions.start_rig(lampyr)
    except actions.Abort as e:
        click.echo(f'Error starting rig: {e}')
        return
    lampyr.run(behavior, **kwargs)

@mouse.command(name='paradigm')
@click.argument('mouseid')
@click.argument('paradigm_name', required=False)
@click.option('--stage', '-s', type=str, default=None, help='Set the current stage for this paradigm')
@click.pass_obj
def mouse_paradigm(lampyr, mouseid, paradigm_name, stage):
    try:
        lampyr.mousemanager.load(mouseid)
    except Exception as e:
        click.echo(f'Failed to load mouse: {e}')
        return

    if paradigm_name is None:
        if stage is not None:
            current_paradigm = lampyr.mouse.paradigm
            if not current_paradigm:
                click.echo('No paradigm set. Provide a paradigm name or set one first.')
                return
            paradigm_cls = lampyr.behaviors.get(current_paradigm)
            valid_stages = getattr(paradigm_cls, 'stagelist', ())
            valid_stages = [s.slug for s in valid_stages]
            if valid_stages and stage not in valid_stages:
                click.echo(f'Invalid stage "{stage}". Valid stages: {", ".join(valid_stages)}')
                return
            lampyr.mouse.properties[paradigm_cls.slug]['stage'] = stage
            click.echo(f'Set {mouseid} stage → {stage} (paradigm: {current_paradigm})')
            lampyr.mousemanager.save()
            return
        # Dump current paradigm info
        m = lampyr.mouse
        current_paradigm = m.paradigm
        paradigm_cls = lampyr.behaviors.get(current_paradigm)
        pdata = m.properties.get(paradigm_cls.slug,{})
        if pdata:
            click.echo(f'Paradigm data for {current_paradigm}:')
            import json
            click.echo(json.dumps(pdata, indent=2))
        else:
            click.echo(f'Paradigm data for {current_paradigm}:')
            click.echo('None')
        return

    # Set paradigm
    if paradigm_name not in lampyr.behaviors:
        click.echo(f'Failed: {paradigm_name} is not a valid behavior.')
        return
    lampyr.mouse.paradigm = paradigm_name
    click.echo(f'Set {mouseid} paradigm → {paradigm_name}')

    if stage is not None:
        paradigm_cls = lampyr.behaviors.get(paradigm_name)
        valid_stages_cls = getattr(paradigm_cls, 'stagelist', ())
        valid_stages = [v.slug for v in valid_stages_cls]
        if valid_stages and stage not in valid_stages:
            click.echo(f'Invalid stage "{stage}". Valid stages: {", ".join(valid_stages)}')
            return
        lampyr.mouse.properties[paradigm_name]['stage'] = stage
        click.echo(f'Set {mouseid} stage → {stage} (paradigm: {paradigm_name})')

    lampyr.mousemanager.save()


@cli.group()
def user():
    pass

@user.command(name='create')
@click.argument('name')
@click.option('-super', 'is_super', is_flag=True, default=False, help='Make user a supervisor (always active)')
@click.option('--pushover_user_key', type=str, default=None, help='Pushover user key')
@click.pass_obj
def user_create(lampyr, name, is_super, pushover_user_key):
    nm = lampyr.notificationmanager
    if name in nm.userdata._config:
        click.echo(f"User '{name}' already exists. Use 'user edit' to modify.")
        return
    nm.add_user(name, pushover_user_key or '', supervisor=is_super)
    click.echo(f"User '{name}' created.")

@user.command(name='edit')
@click.argument('name')
@click.option('--pushover_user_key', type=str, default=None, help='Pushover user key')
@click.option('--supervisor', type=click.BOOL, default=None, help='Set supervisor status (true/false)')
@click.pass_obj
def user_edit(lampyr, name, pushover_user_key, supervisor):
    nm = lampyr.notificationmanager
    if name not in nm.userdata._config:
        click.echo(f"User '{name}' not found.")
        return
    existing = nm.userdata._config[name]
    puk = pushover_user_key if pushover_user_key is not None else existing.get('pushover_user_key', '')
    sup = supervisor if supervisor is not None else existing.get('supervisor', False)
    nm.add_user(name, puk, supervisor=sup, active=existing.get('active', True))
    click.echo(f"User '{name}' updated.")

@user.command(name='set-token')
@click.argument('token')
@click.pass_obj
def user_set_token(lampyr, token):
    lampyr.notificationmanager.set_app_token(token)
    click.echo("Pushover app token updated.")

@user.command(name='inspect')
@click.argument('name')
@click.pass_obj
def user_inspect(lampyr, name):
    users = lampyr.notificationmanager.userdata._config
    if name not in users:
        click.echo(f"User '{name}' not found.")
        return
    actions.printheader(name.upper())
    actions.printinfo(users[name])

@user.command(name='list')
@click.pass_obj
def user_list(lampyr):
    users = {n: d for n, d in lampyr.notificationmanager.userdata._config.items() if isinstance(d, dict)}
    if not users:
        click.echo('No users configured.')
        return
    actions.printheader('USERS')
    for name, data in users.items():
        if data.get('supervisor'):
            tag = ' [supervisor]'
        elif data.get('active', True):
            tag = ' [active]'
        else:
            tag = ' [inactive]'
        click.echo(f'{name}{tag}')

@user.command(name='activate')
@click.argument('name')
@click.pass_obj
def user_activate(lampyr, name):
    nm = lampyr.notificationmanager
    try:
        nm.set_user_active(name, True)
        click.echo(f"User '{name}' activated.")
    except KeyError as e:
        click.echo(f"Error: {e}")

@user.command(name='deactivate')
@click.argument('name')
@click.pass_obj
def user_deactivate(lampyr, name):
    nm = lampyr.notificationmanager
    try:
        nm.set_user_active(name, False)
        click.echo(f"User '{name}' deactivated.")
    except (KeyError, ValueError) as e:
        click.echo(f"Error: {e}")

@user.command(name='ping')
@click.option('--user', '-u', 'target', type=str, default=None, help='User to ping (defaults to all active users)')
@click.option('--message', '-m', type=str, default='This is a test notification from Lampyr.', help='Message to send')
@click.option('--urgent', is_flag=True, default=False, help='Send as emergency priority (bypasses Do Not Disturb)')
@click.pass_obj
def user_ping(lampyr, target, message, urgent):
    nm = lampyr.notificationmanager
    users = {n: d for n, d in nm.userdata._config.items() if isinstance(d, dict)}
    if not users:
        click.echo('No users configured.')
        return
    title = 'Lampyr Emergency' if urgent else 'Lampyr Ping'
    if target is not None:
        if target not in users:
            click.echo(f"User '{target}' not found.")
            return
        nm._send_to_user(target, message, title, urgent=urgent)
    else:
        for name, data in users.items():
            if data.get('active', True):
                nm._send_to_user(name, message, title, urgent=urgent)

@user.command(name='remove')
@click.argument('name')
@click.pass_obj
def user_remove(lampyr, name):
    nm = lampyr.notificationmanager
    try:
        nm.delete_user(name)
        click.echo(f"User '{name}' removed.")
    except KeyError as e:
        click.echo(f"Error: {e}")

@cli.command()
@click.argument('behavior')
@click.option('--merit_limit', '-ml', type=int, required=False)
@click.option('--merit_min', '-mm', type=int, required=False)
@click.option('--demerit_limit', '-dml', type=int, required=False)
@click.option('--demerit_min', '-dmm', type=int, required=False)
@click.option('--duration_limit', '-dl', type=int, required=False)
@click.option('--duration_min', '-dm', type=int, required=False)
@click.option('--trial_limit', '-tl', type=int, required=False)
@click.option('--trial_min', '-tm', type=int, required=False)
@click.option('--reward_limit', '-rl', type=int, required=False)
@click.option('--reward_min', '-rm', type=int, required=False)
@click.option('--abstention_limit', '-al', type=int, required=False)
@click.option('--abstention_min', '-am', type=int, required=False)
@click.option('--participation_limit', '-pl', type=int, required=False)
@click.option('--participation_min', '-pm', type=int, required=False)
@click.option('--serial_abstention_limit', '-sal', type=int, required=False)
@click.option('--serial_abstention_min', '-sam', type=int, required=False)
@click.pass_obj
def run(lampyr : Lampyr, behavior, **kwargs):
    try:
        actions.start_rig(lampyr)
    except actions.Abort:
        return
    lampyr.run(behavior, **kwargs)

def _start_touch_mouse_bridge():
    """Invisible fullscreen Win32 overlay that converts WM_TOUCH/WM_POINTER touch
    events to mouse clicks via SendInput.  Windows Terminal suppresses touch-to-mouse
    promotion for its own window, so Textual never sees taps without this shim."""
    import ctypes
    import ctypes.wintypes
    import threading
    import time

    user32   = ctypes.windll.user32
    kernel32 = ctypes.windll.kernel32

    user32.DefWindowProcW.restype  = ctypes.c_ssize_t   # LRESULT (pointer-sized)
    user32.DefWindowProcW.argtypes = [
        ctypes.c_void_p,  # HWND
        ctypes.c_uint,    # UINT Msg
        ctypes.c_size_t,  # WPARAM
        ctypes.c_size_t,  # LPARAM
    ]

    WS_POPUP          = 0x80000000
    WS_EX_LAYERED     = 0x00080000
    WS_EX_TRANSPARENT = 0x00000020
    WS_EX_TOPMOST     = 0x00000008
    WS_EX_NOACTIVATE  = 0x08000000
    WS_EX_TOOLWINDOW  = 0x00000080
    LWA_ALPHA         = 0x00000002
    GWL_EXSTYLE       = -20
    WM_TOUCH          = 0x0240
    WM_POINTERDOWN    = 0x0246
    WM_POINTERUP      = 0x0247
    WM_DESTROY        = 0x0002
    TOUCHEVENTF_DOWN  = 0x0002
    TOUCHEVENTF_UP    = 0x0004
    WM_LBUTTONDOWN  = 0x0201
    WM_LBUTTONUP    = 0x0202
    WM_RBUTTONDOWN  = 0x0204
    WM_RBUTTONUP    = 0x0205
    WM_MOUSEWHEEL   = 0x020A
    INPUT_MOUSE           = 0
    MOUSEEVENTF_MOVE      = 0x0001
    MOUSEEVENTF_LEFTDOWN  = 0x0002
    MOUSEEVENTF_LEFTUP    = 0x0004
    MOUSEEVENTF_RIGHTDOWN = 0x0008
    MOUSEEVENTF_RIGHTUP   = 0x0010
    MOUSEEVENTF_WHEEL     = 0x0800
    MOUSEEVENTF_ABSOLUTE  = 0x8000
    MOUSEEVENTF_VIRTUALDESK = 0x4000

    # Virtual desktop bounds (handles multi-monitor)
    sw = user32.GetSystemMetrics(78) or user32.GetSystemMetrics(0)
    sh = user32.GetSystemMetrics(79) or user32.GetSystemMetrics(1)
    sx = user32.GetSystemMetrics(76)
    sy = user32.GetSystemMetrics(77)

    class TOUCHINPUT(ctypes.Structure):
        _fields_ = [
            ('x', ctypes.c_long), ('y', ctypes.c_long),
            ('hSource', ctypes.c_void_p), ('dwID', ctypes.c_ulong),
            ('dwFlags', ctypes.c_ulong), ('dwMask', ctypes.c_ulong),
            ('dwTime', ctypes.c_ulong), ('dwExtraInfo', ctypes.POINTER(ctypes.c_ulong)),
            ('cxContact', ctypes.c_ulong), ('cyContact', ctypes.c_ulong),
        ]

    class MOUSEINPUT(ctypes.Structure):
        _fields_ = [
            ('dx', ctypes.c_long), ('dy', ctypes.c_long),
            ('mouseData', ctypes.c_ulong), ('dwFlags', ctypes.c_ulong),
            ('time', ctypes.c_ulong), ('dwExtraInfo', ctypes.POINTER(ctypes.c_ulong)),
        ]

    class INPUT(ctypes.Structure):
        class _U(ctypes.Union):
            _fields_ = [('mi', MOUSEINPUT)]
        _anonymous_ = ('_u',)
        _fields_ = [('type', ctypes.c_ulong), ('_u', _U)]

    overlay = [None]   # mutable hwnd ref shared with closures
    injecting = [False]

    def inject(x, y, flags):
        """Inject a mouse event at screen coords (x, y), briefly making the overlay
        WS_EX_TRANSPARENT so the event reaches the terminal below."""
        inp = INPUT()
        inp.type = INPUT_MOUSE
        inp.mi.dx = int((x - sx) * 65535 // sw)
        inp.mi.dy = int((y - sy) * 65535 // sh)
        inp.mi.dwFlags = flags | MOUSEEVENTF_ABSOLUTE | MOUSEEVENTF_VIRTUALDESK
        hwnd = overlay[0]
        injecting[0] = True
        if hwnd:
            old = user32.GetWindowLongW(hwnd, GWL_EXSTYLE)
            user32.SetWindowLongW(hwnd, GWL_EXSTYLE, old | WS_EX_TRANSPARENT)
            user32.SendInput(1, ctypes.byref(inp), ctypes.sizeof(inp))
            # Brief pause ensures SendInput is dispatched before we remove transparency
            time.sleep(0.01)
            user32.SetWindowLongW(hwnd, GWL_EXSTYLE, old)
        else:
            user32.SendInput(1, ctypes.byref(inp), ctypes.sizeof(inp))
        injecting[0] = False

    def inject_wheel(delta):
        """Inject a mouse-wheel event, briefly making the overlay transparent."""
        inp = INPUT()
        inp.type = INPUT_MOUSE
        inp.mi.mouseData = ctypes.c_ulong(delta).value
        inp.mi.dwFlags = MOUSEEVENTF_WHEEL
        hwnd = overlay[0]
        injecting[0] = True
        if hwnd:
            old = user32.GetWindowLongW(hwnd, GWL_EXSTYLE)
            user32.SetWindowLongW(hwnd, GWL_EXSTYLE, old | WS_EX_TRANSPARENT)
            user32.SendInput(1, ctypes.byref(inp), ctypes.sizeof(inp))
            time.sleep(0.01)
            user32.SetWindowLongW(hwnd, GWL_EXSTYLE, old)
        else:
            user32.SendInput(1, ctypes.byref(inp), ctypes.sizeof(inp))
        injecting[0] = False

    def get_xy(lp):
        """Extract signed screen coords from a WM_POINTER lParam."""
        return (ctypes.c_short(lp & 0xFFFF).value,
                ctypes.c_short((lp >> 16) & 0xFFFF).value)

    WNDPROC = ctypes.WINFUNCTYPE(
        ctypes.c_long, ctypes.c_void_p, ctypes.c_uint,
        ctypes.c_size_t, ctypes.c_size_t,
    )

    def wndproc(hwnd, msg, wp, lp):
        # ── WM_TOUCH (reliable path when RegisterTouchWindow is used) ────────
        if msg == WM_TOUCH:
            count  = wp & 0xFFFF
            handle = ctypes.c_void_p(lp)
            touches = (TOUCHINPUT * count)()
            if user32.GetTouchInputInfo(handle, count,
                                        ctypes.byref(touches),
                                        ctypes.sizeof(TOUCHINPUT)):
                for t in touches:
                    # WM_TOUCH coords are in hundredths of pixels
                    x, y = t.x // 100, t.y // 100
                    if t.dwFlags & TOUCHEVENTF_DOWN:
                        inject(x, y, MOUSEEVENTF_MOVE)
                        inject(x, y, MOUSEEVENTF_LEFTDOWN)
                    elif t.dwFlags & TOUCHEVENTF_UP:
                        inject(x, y, MOUSEEVENTF_LEFTUP)
            user32.CloseTouchInputHandle(handle)
            return 0

        # ── Real mouse clicks (overlay intercepts these; forward them through) ──
        if not injecting[0]:
            pt = ctypes.wintypes.POINT()
            user32.GetCursorPos(ctypes.byref(pt))
            x, y = pt.x, pt.y
            if msg == WM_LBUTTONDOWN:
                inject(x, y, MOUSEEVENTF_MOVE)
                inject(x, y, MOUSEEVENTF_LEFTDOWN)
                return 0
            if msg == WM_LBUTTONUP:
                inject(x, y, MOUSEEVENTF_LEFTUP)
                return 0
            if msg == WM_RBUTTONDOWN:
                inject(x, y, MOUSEEVENTF_MOVE)
                inject(x, y, MOUSEEVENTF_RIGHTDOWN)
                return 0
            if msg == WM_RBUTTONUP:
                inject(x, y, MOUSEEVENTF_RIGHTUP)
                return 0
            if msg == WM_MOUSEWHEEL:
                delta = ctypes.c_short(wp >> 16).value
                inject_wheel(delta)
                return 0

        # ── WM_POINTER (fallback) ─────────────────────────────────────────────
        if msg == WM_POINTERDOWN and not injecting[0]:
            x, y = get_xy(lp)
            inject(x, y, MOUSEEVENTF_MOVE)
            inject(x, y, MOUSEEVENTF_LEFTDOWN)
            return 0
        if msg == WM_POINTERUP and not injecting[0]:
            x, y = get_xy(lp)
            inject(x, y, MOUSEEVENTF_LEFTUP)
            return 0

        if msg == WM_DESTROY:
            user32.PostQuitMessage(0)
            return 0
        return user32.DefWindowProcW(hwnd, msg, wp, lp)

    cb = WNDPROC(wndproc)
    _start_touch_mouse_bridge._cb_ref = cb  # prevent GC

    def run():
        class WNDCLASSW(ctypes.Structure):
            _fields_ = [
                ('style', ctypes.c_uint), ('lpfnWndProc', WNDPROC),
                ('cbClsExtra', ctypes.c_int), ('cbWndExtra', ctypes.c_int),
                ('hInstance', ctypes.c_void_p), ('hIcon', ctypes.c_void_p),
                ('hCursor', ctypes.c_void_p), ('hbrBackground', ctypes.c_void_p),
                ('lpszMenuName', ctypes.c_wchar_p), ('lpszClassName', ctypes.c_wchar_p),
            ]

        hmod = kernel32.GetModuleHandleW(None)
        wc = WNDCLASSW()
        wc.lpfnWndProc   = cb
        wc.hInstance     = hmod
        wc.lpszClassName = 'LampyrTouchBridge'
        user32.RegisterClassW(ctypes.byref(wc))

        hwnd = user32.CreateWindowExW(
            WS_EX_LAYERED | WS_EX_TOPMOST | WS_EX_NOACTIVATE | WS_EX_TOOLWINDOW,
            'LampyrTouchBridge', None, WS_POPUP,
            sx, sy, sw, sh, None, None, hmod, None,
        )
        if not hwnd:
            return

        overlay[0] = hwnd
        # Alpha=1 (not 0): alpha=0 causes Windows to exclude the window from
        # hit-testing entirely, so it never receives touch events.
        user32.SetLayeredWindowAttributes(hwnd, 0, 1, LWA_ALPHA)
        user32.RegisterTouchWindow(hwnd, 0x00000002)  # TWF_WANTPALM
        user32.ShowWindow(hwnd, 4)   # SW_SHOWNOACTIVATE
        user32.UpdateWindow(hwnd)

        class MSG(ctypes.Structure):
            _fields_ = [
                ('hwnd', ctypes.c_void_p), ('message', ctypes.c_uint),
                ('wParam', ctypes.c_size_t), ('lParam', ctypes.c_size_t),
                ('time', ctypes.c_ulong), ('pt', ctypes.wintypes.POINT),
            ]

        msg = MSG()
        while user32.GetMessageW(ctypes.byref(msg), None, 0, 0) != 0:
            user32.TranslateMessage(ctypes.byref(msg))
            user32.DispatchMessageW(ctypes.byref(msg))

    threading.Thread(target=run, daemon=True).start()
    time.sleep(0.15)  # allow the overlay window to finish initialising


@cli.command()
@click.option(
    '--notouchoverlay',
    is_flag=True,
    default=False,
    help='Disable the Windows touchscreen click overlay bridge.',
)
def go(notouchoverlay):
    """Launch the Lampyr TUI."""
    import sys
    if sys.platform == "win32":
        import ctypes
        # Fullscreen
        KEYEVENTF_KEYUP = 0x0002
        VK_F11 = 0x7A
        ctypes.windll.user32.keybd_event(VK_F11, 0, 0, 0)
        ctypes.windll.user32.keybd_event(VK_F11, 0, KEYEVENTF_KEYUP, 0)
        # Disable Quick Edit Mode
        STD_INPUT_HANDLE  = -10
        ENABLE_MOUSE_INPUT = 0x0010
        ENABLE_QUICK_EDIT  = 0x0040
        ENABLE_EXTENDED    = 0x0080
        handle = ctypes.windll.kernel32.GetStdHandle(STD_INPUT_HANDLE)
        mode = ctypes.c_ulong()
        ctypes.windll.kernel32.GetConsoleMode(handle, ctypes.byref(mode))
        ctypes.windll.kernel32.SetConsoleMode(
            handle,
            (mode.value | ENABLE_MOUSE_INPUT | ENABLE_EXTENDED) & ~ENABLE_QUICK_EDIT,
        )
        # Touch-to-mouse bridge
        if not notouchoverlay:
            _start_touch_mouse_bridge()
    from lampyr.interfaces.textual_tui.app import LampyrApp
    LampyrApp().run()
    sys.exit(0)
```
