## Extended Data for "A Low-Cost, Modular Hardware and Software Platform for Head-Fixed Mouse Decision-Making Tasks": app.py

```
# -*- coding: utf-8 -*-
"""
Lampyr Textual TUI
Launched via: lampyr go
"""

import ctypes
import re as _re
import threading
import time
from importlib import resources as impres
from typing import Callable

import art
from textual import on
from textual.app import App, ComposeResult
from textual.containers import Container, Vertical, VerticalScroll
from textual.message import Message
from textual.screen import ModalScreen, Screen
from rich.text import Text
from textual.widgets import Button, Footer, Label, RichLog

from lampyr import Lampyr
from lampyr import actions
from lampyr.segments.behavior import Task


# ---------------------------------------------------------------------------
# TUI display helper — shorten [segment_name] slug to uppercase letters + digits
# ---------------------------------------------------------------------------

def _shorten_segment_slug(text: str) -> str:
    """TUI display only: compress the [segment_name] bracket to uppercase+digits."""
    count = [0]
    def _replace(m):
        count[0] += 1
        if count[0] == 2:
            return '[' + _re.sub(r'[^A-Z0-9]', '', m.group(1)) + ']'
        return m.group(0)
    return _re.sub(r'\[([^\]]*)\]', _replace, text)


# ---------------------------------------------------------------------------
# Abort helper — raises an exception in a running thread
# ---------------------------------------------------------------------------

def _async_raise(tid, exctype):
    res = ctypes.pythonapi.PyThreadState_SetAsyncExc(
        ctypes.c_ulong(tid), ctypes.py_object(exctype)
    )
    if res == 0:
        raise ValueError("Invalid thread id")


# ---------------------------------------------------------------------------
# Threading bridge: routes blocking _input_func calls to the TUI numpad
# ---------------------------------------------------------------------------

class TUIInputBridge:
    """Callable that stands in for input().

    When calibration (or any blocking code) calls self._input_func(prompt),
    this bridge signals the TUI to show a NumpadModal, then blocks the worker
    thread until the user submits a value.
    """

    def __init__(self):
        self._event = threading.Event()
        self._value = None
        self.show_numpad_cb: Callable[[str], None] = lambda p: None

    def __call__(self, prompt: str) -> str:
        self._event.clear()
        self._value = None
        self.show_numpad_cb(prompt)
        self._event.wait()
        return self._value

    def submit(self, value: str) -> None:
        self._value = value
        self._event.set()


# ---------------------------------------------------------------------------
# NumpadModal — touch-friendly numeric / ID entry
# ---------------------------------------------------------------------------

class NumpadModal(ModalScreen):
    """Touch-friendly numpad modal.

    mode='float'  →  decimal point key; validates as float on OK
    mode='int'    →  no decimal key; validates as int on OK
    mode='id'     →  dash key; validates as non-empty string on OK
    """

    def __init__(self, prompt: str, mode: str = "float", calibration: bool = False):
        super().__init__()
        self._prompt = prompt
        self._mode = mode
        self._calibration = calibration
        self._current = ""
        self._pending = ""   # value awaiting confirmation

    def compose(self) -> ComposeResult:
        extra_label = "." if self._mode == "float" else "-"
        classes = "numpad-calibration" if self._calibration else ""
        with Container(id="numpad-modal", classes=classes):
            yield Label(self._prompt, id="numpad-prompt")
            yield Label("0" if self._mode == "float" else "", id="numpad-display")
            # ── Entry widgets ──────────────────────────────
            with Container(id="numpad-grid"):
                yield Button("7", id="n7",    classes="numpad-digit")
                yield Button("8", id="n8",    classes="numpad-digit")
                yield Button("9", id="n9",    classes="numpad-digit")
                yield Button("4", id="n4",    classes="numpad-digit")
                yield Button("5", id="n5",    classes="numpad-digit")
                yield Button("6", id="n6",    classes="numpad-digit")
                yield Button("1", id="n1",    classes="numpad-digit")
                yield Button("2", id="n2",    classes="numpad-digit")
                yield Button("3", id="n3",    classes="numpad-digit")
                yield Button(extra_label, id="nextra", classes="numpad-digit")
                yield Button("0", id="n0",    classes="numpad-digit")
                yield Button("⌫", id="nback", classes="numpad-digit")
            yield Button("OK", id="numpad-ok")
            # ── Confirmation widgets (hidden until OK pressed) ──
            yield Button("✓  CONFIRM",  id="numpad-confirm")
            yield Button("✗  RE-ENTER", id="numpad-reenter")

    def on_mount(self) -> None:
        self.query_one("#numpad-confirm").display = False
        self.query_one("#numpad-reenter").display = False

    def _update_display(self) -> None:
        display = self._current or ("0" if self._mode == "float" else "")
        self.query_one("#numpad-display", Label).update(display)

    def _show_confirm(self, val: str) -> None:
        """Switch to confirmation state."""
        self._pending = val
        self.query_one("#numpad-display", Label).update(f"Confirm:  {val}")
        self.query_one("#numpad-grid").display   = False
        self.query_one("#numpad-ok").display     = False
        self.query_one("#numpad-confirm").display  = True
        self.query_one("#numpad-reenter").display  = True

    def _show_entry(self) -> None:
        """Switch back to entry state."""
        self._pending = ""
        self.query_one("#numpad-grid").display    = True
        self.query_one("#numpad-ok").display      = True
        self.query_one("#numpad-confirm").display   = False
        self.query_one("#numpad-reenter").display   = False
        self._update_display()

    @on(Button.Pressed)
    def on_button(self, event: Button.Pressed) -> None:
        btn_id = event.button.id

        # ── Confirmation state buttons ──
        if btn_id == "numpad-confirm":
            self.dismiss(self._pending)
            return
        if btn_id == "numpad-reenter":
            self._show_entry()
            return

        # ── Entry state buttons ──
        if btn_id == "numpad-ok":
            val = self._current
            if self._mode == "float":
                try:
                    float(val)
                    self._show_confirm(val)
                except (ValueError, TypeError):
                    self.query_one("#numpad-display", Label).update("invalid!")
            elif self._mode == "int":
                if val == "-":
                    self._show_confirm(val)
                else:
                    try:
                        int(val)
                        self._show_confirm(val)
                    except (ValueError, TypeError):
                        self.query_one("#numpad-display", Label).update("invalid!")
            else:
                if val:
                    self._show_confirm(val)
                else:
                    self.query_one("#numpad-display", Label).update("enter ID!")
            return

        if btn_id == "nback":
            self._current = self._current[:-1]
        elif btn_id == "nextra":
            if self._mode == "float" and "." not in self._current:
                self._current += "."
            elif self._mode == "id":
                self._current += "-"
            elif self._mode == "int" and self._current == "":
                self._current = "-"
        else:
            self._current += str(event.button.label)

        self._update_display()


# ---------------------------------------------------------------------------
# CalibrationConfirmScreen — yellow gate shown before calibration starts
# ---------------------------------------------------------------------------

class CalibrationConfirmScreen(Screen):
    """Shown before starting calibration so an unattended rig check never
    silently opens the serial port."""

    def compose(self) -> ComposeResult:
        yield Label("⚠  CALIBRATION", id="calib-header")
        yield Label(
            "This will connect to the Arduino rig and begin the calibration "
            "procedure.\n\nEnsure the rig is available and press CONFIRM to proceed.",
            id="calib-subheader",
        )
        yield Button("✓  BEGIN CALIBRATION", id="confirm-yes")
        yield Button("✗  CANCEL",            id="confirm-no")

    @on(Button.Pressed, "#confirm-yes")
    def on_yes(self) -> None:
        self.app.switch_screen(CalibrationScreen())

    @on(Button.Pressed, "#confirm-no")
    def on_no(self) -> None:
        self.app.pop_screen()


# ---------------------------------------------------------------------------
# CalibrationScreen — walks user through calibration cycle
# ---------------------------------------------------------------------------

class CalibrationScreen(Screen):

    class CalibrationDone(Message):
        pass

    class LogOutput(Message):
        def __init__(self, text: str):
            super().__init__()
            self.text = text

    def compose(self) -> ComposeResult:
        yield Label("⚠  CALIBRATION REQUIRED", id="calib-header")
        yield Label(
            "Follow the prompts below. Enter weights using the numpad.",
            id="calib-subheader",
        )
        yield RichLog(id="calib-log", markup=True, highlight=True)
        yield Button("◀  RETURN TO MAIN", id="calib-return")

    def on_mount(self) -> None:
        self.query_one("#calib-return").display = False
        log = self.query_one("#calib-log", RichLog)
        log.border_title = "Calibration Output"
        # post_message is explicitly thread-safe; use it instead of call_from_thread
        self.app.set_output(
            lambda msg: self.post_message(CalibrationScreen.LogOutput(str(msg)))
        )
        threading.Thread(target=self._run_calibration, daemon=True).start()

    @on(LogOutput)
    def on_log(self, event: LogOutput) -> None:
        self.query_one("#calib-log", RichLog).write(Text.from_ansi(event.text))

    def _run_calibration(self) -> None:
        try:
            self.app.lampyr.rigmanager.calibrate()
            self.post_message(CalibrationScreen.CalibrationDone())
        except Exception as e:
            self.post_message(
                CalibrationScreen.LogOutput(f"\x1b[1;31mCalibration error:\x1b[0m {e}")
            )
            self.post_message(CalibrationScreen.CalibrationDone())

    @on(CalibrationDone)
    def on_done(self) -> None:
        self.query_one("#calib-return", Button).display = True

    @on(Button.Pressed, "#calib-return")
    def on_return(self) -> None:
        self.app.pop_screen()


# ---------------------------------------------------------------------------
# BehaviorSelectScreen — pick a behavior (ADVANCED flow)
# ---------------------------------------------------------------------------

class BehaviorSelectScreen(Screen):

    def __init__(self, mouseid: str):
        super().__init__()
        self._mouseid = mouseid

    def compose(self) -> ComposeResult:
        yield Label(
            f"Select Behavior  ·  Mouse: {self._mouseid}",
            id="behsel-header",
        )
        user_behaviors = [
            name for name, cls in self.app.lampyr.behaviors.items()
            if issubclass(cls, Task) and cls is not Task
        ]
        with VerticalScroll(id="behavior-list"):
            for name in user_behaviors:
                yield Button(name, classes="behavior-btn")

    @on(Button.Pressed, ".behavior-btn")
    def on_behavior(self, event: Button.Pressed) -> None:
        behavior_name = str(event.button.label)
        self.app.push_screen(TaskParamScreen(self._mouseid, behavior_name))


# ---------------------------------------------------------------------------
# TaskParamScreen — configure session limits before running a task
# ---------------------------------------------------------------------------

_SESSION_PARAMS = [
    ("reward_limit",          "Reward Limit"),
    ("duration_limit",        "Duration (min)"),
    ("trial_limit",           "Trial Limit"),
    ("merit_limit",           "Merit Limit"),
    ("demerit_limit",         "Demerit Limit"),
    ("participation_limit",   "Participation Limit"),
    ("abstention_limit",      "Abstention Limit"),
    ("serial_abstention_limit", "Serial Abstention Limit"),
]


class TaskParamScreen(Screen):

    def __init__(self, mouseid: str, behavior: str):
        super().__init__()
        self._mouseid = mouseid
        self._behavior = behavior
        self._params: dict = {
            "reward_limit": 200,
            "duration_limit": 60,
            "serial_abstention_limit": 15,
        }

    def compose(self) -> ComposeResult:
        yield Label(
            f"Configure  ·  {self._behavior}  ·  Mouse: {self._mouseid}",
            id="taskparam-header",
        )
        with VerticalScroll(id="taskparam-list"):
            for param, label in _SESSION_PARAMS:
                yield Button(
                    self._param_btn_label(param, label),
                    id=f"param-{param}",
                    classes="taskparam-btn",
                )
        with Container(id="taskparam-actions"):
            yield Button("▶  RUN",  id="taskparam-run",  variant="success")
            yield Button("◀  BACK", id="taskparam-back")

    def _param_btn_label(self, param: str, label: str) -> str:
        val = self._params.get(param)
        val_str = f"[ {val} ]" if val is not None else "[  —  ]"
        return f"{label}  {val_str}"

    @on(Button.Pressed, ".taskparam-btn")
    def on_param_btn(self, event: Button.Pressed) -> None:
        param = event.button.id.removeprefix("param-")
        label = next(lbl for p, lbl in _SESSION_PARAMS if p == param)
        self.app.push_screen(
            NumpadModal(f"Set {label}:", mode="int"),
            lambda val, p=param, lbl=label: self._on_param_set(p, lbl, val),
        )

    def _on_param_set(self, param: str, label: str, val: str | None) -> None:
        if val is not None:
            if val == "-":
                self._params.pop(param, None)
            else:
                self._params[param] = int(val)
            self.query_one(f"#param-{param}", Button).label = self._param_btn_label(param, label)

    @on(Button.Pressed, "#taskparam-run")
    def on_run(self) -> None:
        self.app.switch_screen(RunScreen(self._mouseid, self._behavior, session_params=self._params))

    @on(Button.Pressed, "#taskparam-back")
    def on_back(self) -> None:
        self.app.pop_screen()


# ---------------------------------------------------------------------------
# RunScreen — black session output screen with ABORT
# ---------------------------------------------------------------------------

class RunScreen(Screen):

    class RunDone(Message):
        def __init__(self, error: bool = False):
            super().__init__()
            self.error = error

    class LogOutput(Message):
        def __init__(self, text: str):
            super().__init__()
            self.text = text

    def __init__(self, mouseid: str, behavior: str, session_params: dict = None):
        super().__init__()
        self._mouseid = mouseid
        self._behavior = behavior
        self._session_params = session_params or {}
        self._thread: threading.Thread | None = None
        self._animal_timer: threading.Timer | None = None

    def compose(self) -> ComposeResult:
        yield RichLog(id="run-output", markup=True, highlight=True, wrap=True)
        yield Button("⏹  ABORT", id="action-btn", variant="error")

    def on_mount(self) -> None:
        log = self.query_one("#run-output", RichLog)
        log.border_title = f"{self._mouseid}  ·  {self._behavior}"
        # post_message is explicitly thread-safe; use it instead of call_from_thread
        self.app.set_output(
            lambda msg: self.post_message(RunScreen.LogOutput(str(msg)))
        )
        self._thread = threading.Thread(target=self._run, daemon=True)
        self._thread.start()

    @on(LogOutput)
    def on_log(self, event: LogOutput) -> None:
        self.query_one("#run-output", RichLog).write(Text.from_ansi(_shorten_segment_slug(event.text)))

    def _run(self) -> None:
        import traceback as _tb

        # Always use post_message directly — it's guaranteed thread-safe.
        def out(msg: str) -> None:
            self.post_message(RunScreen.LogOutput(str(msg)))

        error = False
        try:
            out(f"Loading mouse {self._mouseid}...")
            self.app.lampyr.mousemanager.load(self._mouseid)

            # Describe rig start failures explicitly — actions.Abort has no message
            out("Checking rig configuration...")
            configured = self.app.lampyr.config.get("rig.configured")
            calibrated = self.app.lampyr.config.get("rig.calibrated")
            if not configured or configured < 1:
                out("\x1b[1;31mERROR: Rig is not configured.\x1b[0m")
                out("\x1b[33mRun 'lampyr rig configure' or use developer mode.\x1b[0m")
                error = True
                return
            if calibrated < time.time() - (5 * 24 * 60 * 60):
                out("\x1b[1;31mERROR: Rig calibration has expired.\x1b[0m")
                out("\x1b[33mTap CALIBRATE on the main screen.\x1b[0m")
                error = True
                return

            out("Connecting to rig...")
            self.app.lampyr.rigmanager.connect()

            out(f"Starting behavior: {self._behavior}")
            self.app.lampyr.run(self._behavior, **self._session_params)

        except actions.Abort:
            out("\x1b[1;31mERROR: Rig start aborted (unconfigured or uncalibrated).\x1b[0m")
            error = True
        except Exception as e:
            out(f"\x1b[1;31mERROR: {e}\x1b[0m")
            out(_tb.format_exc())
            error = True
        finally:
            self.post_message(RunScreen.RunDone(error=error))

    @on(RunDone)
    def on_done(self, event: RunDone) -> None:
        """Session finished — clean up and flip button to RETURN rather than auto-popping."""
        try:
            self.app.lampyr.close()
        except Exception:
            pass
        btn = self.query_one("#action-btn", Button)
        if event.error:
            btn.label = "◀  RETURN  (session ended with error)"
            btn.variant = "error"
            self.add_class("run-done-error")
        else:
            btn.label = "◀  RETURN TO MAIN"
            btn.variant = "success"
            self.add_class("run-done-success")
        self._animal_timer = threading.Timer(1800, self._animal_left_alert)
        self._animal_timer.daemon = True
        self._animal_timer.start()
        btn.disabled = False

    def _animal_left_alert(self) -> None:
        rig_name = self.app.lampyr.config.get("rig.name") or "UNKNOWN"
        msg = f"Rig {rig_name} may have a mouse in it that has been left for 30 minutes after session end. Please check on this rig/mouse."
        nm = self.app.lampyr.notificationmanager
        nm.userdata.reload()
        for name, data in nm.userdata.to_dict().items():
            if not isinstance(data, dict):
                continue
            if data.get('active', True) or data.get('supervisor'):
                nm._send_to_user(name, msg, "URGENT NOTICE", urgent=True)

    @on(Button.Pressed, "#action-btn")
    def on_action_btn(self, event: Button.Pressed) -> None:
        if self._thread and self._thread.is_alive():
            # Still running → ABORT
            event.button.disabled = True
            event.button.label = "Aborting…"
            # Set flag directly on lampyr so main.py's finally block
            # reliably records 'user intervention' regardless of where
            # the KeyboardInterrupt lands in the segment hierarchy.
            self.app.lampyr._user_aborted = True
            try:
                _async_raise(self._thread.ident, KeyboardInterrupt)
            except ValueError:
                pass
        else:
            # Session finished → return to main, not the prior task-selection screen.
            if self._animal_timer is not None:
                self._animal_timer.cancel()
                self._animal_timer = None
            for screen in reversed(self.app.screen_stack):
                if isinstance(screen, MainScreen):
                    screen.pop_until_active()
                    break
            else:
                self.app.pop_screen()


# ---------------------------------------------------------------------------
# MainScreen — three large touch buttons
# ---------------------------------------------------------------------------

class MainScreen(Screen):

    def compose(self) -> ComposeResult:
        rig_name = self.app.lampyr.config.get("rig.name") or "Lampyr"
        with Vertical(id="main-content"):
            yield Label(art.text2art(rig_name, font="small"), id="rig-title")
            yield Button("RUN",       id="btn-run",       classes="main-btn")
            yield Button("ADVANCED",  id="btn-advanced",  classes="main-btn")
            yield Button("CALIBRATE", id="btn-calibrate", classes="main-btn")
            yield Button("✕  QUIT",   id="btn-quit",      classes="main-btn")

    # ── RUN ──────────────────────────────────────────────────────────────

    @on(Button.Pressed, "#btn-run")
    def on_run(self) -> None:
        self.app.push_screen(
            NumpadModal("Enter Mouse ID:", mode="id"),
            self._start_run,
        )

    def _start_run(self, mouseid: str | None) -> None:
        if not mouseid:
            return
        try:
            self.app.lampyr.mousemanager.load(mouseid)
        except Exception as e:
            self.app.notify(f"Mouse not found: {mouseid}", severity="error", timeout=5)
            return
        paradigm = self.app.lampyr.mouse.paradigm
        if paradigm:
            self.app.push_screen(RunScreen(mouseid, paradigm))
        else:
            self.app.push_screen(BehaviorSelectScreen(mouseid))

    # ── ADVANCED ─────────────────────────────────────────────────────────

    @on(Button.Pressed, "#btn-advanced")
    def on_advanced(self) -> None:
        self.app.push_screen(
            NumpadModal("Enter Mouse ID:", mode="id"),
            self._start_advanced,
        )

    def _start_advanced(self, mouseid: str | None) -> None:
        if not mouseid:
            return
        try:
            self.app.lampyr.mousemanager.load(mouseid)
        except Exception:
            self.app.notify(f"Mouse not found: {mouseid}", severity="error", timeout=5)
            return
        self.app.push_screen(BehaviorSelectScreen(mouseid))

    # ── CALIBRATE ────────────────────────────────────────────────────────

    @on(Button.Pressed, "#btn-calibrate")
    def on_calibrate(self) -> None:
        self.app.push_screen(CalibrationConfirmScreen())

    # ── QUIT ─────────────────────────────────────────────────────────────

    @on(Button.Pressed, "#btn-quit")
    def on_quit(self) -> None:
        self.app.exit()


# ---------------------------------------------------------------------------
# LampyrApp — root application
# ---------------------------------------------------------------------------

class LampyrApp(App):
    """Lampyr TUI — launched via `lampyr go`."""

    CSS_PATH = impres.files("lampyr").joinpath(
        "interfaces/textual_tui/app.tcss"
    )

    def __init__(self):
        self.bridge = TUIInputBridge()
        self.lampyr = Lampyr(
            _input_func=self.bridge,
            _output_func=lambda msg: None,  # overridden per-screen via set_output()
        )
        self.bridge.show_numpad_cb = lambda prompt: self.call_from_thread(
            self._show_numpad, prompt
        )
        super().__init__()

    def set_output(self, func: Callable) -> None:
        """Update _output_func on Lampyr and all sub-managers at once.

        AbstractManager copies _output_func at construction time, so we must
        push updates to each manager directly whenever the active screen changes.
        """
        self.lampyr._output_func = func
        for mgr in (
            self.lampyr.rigmanager,
            self.lampyr.mousemanager,
            self.lampyr.datamanager,
            self.lampyr.notificationmanager,
        ):
            mgr._output_func = func

    def _show_numpad(self, prompt: str) -> None:
        """Push the numpad modal and route its result back to the bridge."""
        calibration = isinstance(self.screen, CalibrationScreen)
        if calibration:
            # Echo the prompt + entered value to the calibration log
            calib_screen = self.screen
            def on_calib_submit(value: str) -> None:
                calib_screen.post_message(
                    CalibrationScreen.LogOutput(f"{prompt} {value}")
                )
                self.bridge.submit(value)
            self.push_screen(NumpadModal(prompt, calibration=True), on_calib_submit)
        else:
            self.push_screen(NumpadModal(prompt), self.bridge.submit)

    def on_mount(self) -> None:
        self.theme = "textual-dark"
        self.push_screen(MainScreen())
        self._heartbeat()
        self.set_interval(1200, self._heartbeat) #every 20 minutes

    def _heartbeat(self) -> None:
        """Push CalibrationConfirmScreen whenever calibration has expired.

        Scans the full screen stack so that a NumpadModal sitting on top of a
        CalibrationScreen, or an already-open confirm screen, does not trigger
        a second push.
        """
        expired = self.lampyr.config.get("rig.calibrated") < time.time() - (5*24*60*60)
        already_active = any(
            isinstance(s, (CalibrationScreen, CalibrationConfirmScreen))
            for s in self.screen_stack
        )
        session_running = any(isinstance(s, RunScreen) for s in self.screen_stack)
        if expired and not already_active and not session_running:
            self.push_screen(CalibrationConfirmScreen())
        if not session_running:
            try:
                self.lampyr.datamanager.heartbeat_filetouch()
            except Exception as e:
                self.notify(
                    f"Heartbeat file touch failed.\n{type(e).__name__}: {e}",
                    severity="error", timeout=20*60)


if __name__ == "__main__":
    LampyrApp().run()
```
