## Extended Data for "A Low-Cost, Modular Hardware and Software Platform for Head-Fixed Mouse Decision-Making Tasks": main.py

```
# -*- coding: utf-8 -*-
"""
Created on Thu May 22 15:48:30 2025

@author: mxwll
"""

import os
import traceback
import json
import time

from lampyr.config import Config
from lampyr.primatives import Session

from lampyr.segments.abstract import Segment
from lampyr.managers import DataHandler, MouseManager, RigManager
from lampyr.managers.notification import NotificationManager
from lampyr.managers.plugins import PluginManager

class Lampyr:
    """
    Top-level orchestrator for a Lampyr behavioural experiment session.

    Initialises all managers (data, rig, mouse, notifications) and provides
    the :meth:`run` method to execute a named behaviour segment against the
    currently loaded mouse.

    Attributes
    ----------
    config : Config
        Application configuration.
    session : Session or None
        The session created by the most recent :meth:`run` call, or ``None``
        if no session has been started yet.
    datamanager : DataHandler
        Handles saving/loading of sessions and mice.
    rigmanager : RigManager
        Manages connection to the physical rig.
    mousemanager : MouseManager
        Manages the active mouse.
    notificationmanager : NotificationManager
        Sends push notifications on session end.
    behaviors : dict
        Maps segment class name strings to their classes, populated from all
        known :class:`~lampyr.segments.abstract.Segment` subclasses.
    paradigms : dict
        Maps paradigm name strings to paradigm classes (user-populated).
    """

    def __init__(self, _input_func = input, _output_func=print):
        """
        Initialise Lampyr and all sub-managers.

        Parameters
        ----------
        _input_func : callable, optional
            Function used to prompt for user input.  Defaults to built-in
            ``input``.  Override in tests or non-interactive contexts.
        _output_func : callable, optional
            Function used for text output.  Defaults to built-in ``print``.
        """
        self._output_func = _output_func
        self._input_func = _input_func
        self.config = Config()
        self.session = None
        if self.config.get('lampyr.configured') is False:
            return
        
        self.pluginmanager = PluginManager(self)
        self.datamanager = DataHandler(self)
        
        self.rigmanager = RigManager(self)
        self.mousemanager = MouseManager(self)
        self.notificationmanager = NotificationManager(self)

        self.behaviors = {c.__name__: c for c in Segment.get_children()}
        self.paradigms = {}

    @property
    def rig(self):
        """The connected rig object, or ``None`` if not yet connected."""
        if hasattr(self, 'rigmanager'):
            return self.rigmanager.rig
        return None

    @property
    def mouse(self):
        """The currently active :class:`~lampyr.primatives.Mouse` object."""
        if hasattr(self, 'mousemanager'):
            return self.mousemanager.mouse
        return None

    def run(self, segment_name: str, **kwargs):
        """
        Create a session and run the named behaviour segment.

        Backs up mouse data before starting, creates a new
        :class:`~lampyr.primatives.Session`, instantiates the segment, runs
        it, and sends a push notification on completion regardless of how the
        session ends.

        Parameters
        ----------
        segment_name : str
            Name of a registered behaviour class in :attr:`behaviors`.
        **kwargs
            Additional keyword arguments forwarded to
            :class:`~lampyr.primatives.Session`.

        Raises
        ------
        KeyError
            If ``segment_name`` is not in :attr:`behaviors`.
        RuntimeError
            If a paradigm segment is requested without an active mouse, or if
            no rig is connected.
        """
        if segment_name not in self.behaviors:
            raise KeyError('Not a valid behavior')
        if segment_name in self.paradigms and self.mouse is None:
            raise RuntimeError(
                'Paradigms cannot be run without an active mouse')
        if self.rig is None:
            raise RuntimeError(
                'Segments cannot be run without an active rig')
            
        self.datamanager._backupmice()
        self._createsession(**kwargs)
        behav = self.behaviors[segment_name](lampyr=self,
                                             _verbose=True
                                             )
        stopcodes = []
        try:
            behav.run()
        except KeyboardInterrupt:
            self._output_func('\nAborted by user!\n')
            stopcodes.append('user intervention')
        except Exception as error:
            self._output_func(traceback.format_exc())
        finally:
            self._output_func(behav.session)
            stopcodes += behav.session.evaluatestopconditions()
            if getattr(self, '_user_aborted', False):
                stopcodes.append('user intervention')
                self._user_aborted = False
            if not stopcodes:
                stopcodes.append('finished task')
            rname = self.config.get('rig.name')
            mid = self.mouse.mouseid
            self.notificationmanager.send_notification(
                f'{mid} has ended behavior due to {" and ".join(stopcodes)}',
                title=f'Lampyr - {rname}'
                )
    
    def notify(self, message):
        rigname = self.config.get('rig.name')
        self.notificationmanager.send_notification(
            message,
            title=f'Lampyr - {rigname}')
    
    def _createsession(self, **kwargs):
        """
        Create a new Session, injecting the active mouse ID if available.

        Parameters
        ----------
        **kwargs
            Forwarded to the :class:`~lampyr.primatives.Session` constructor.
        """
        if self.mouse is not None:
            kwargs['mouseid'] = self.mouse.mouseid
        self.session = Session(**kwargs)

    def close(self):
        """
        Gracefully shut down Lampyr: disconnect rig, save session and mouse.

        Safe to call even if the rig was never connected or no session was
        started.  Mouse data is backed up after saving to ensure the shared
        directory is up to date.
        """
        if self.rig is not None:
            self.rigmanager.disconnect()
        if self.session is not None:
            self.datamanager.savesession()
        if self.mouse is not None:
            self.mousemanager.save() # Important that mouse is saved after session
            self.datamanager._backupmice()


if __name__ == '__main__':
    try:
        lamp = Lampyr()
        lamp.mousemanager.load('014-003')
    finally:
        lamp.close()
```
