## Extended Data for "A Low-Cost, Modular Hardware and Software Platform for Head-Fixed Mouse Decision-Making Tasks": abstract.py

```
# -*- coding: utf-8 -*-
"""
Created on Mon Aug 25 18:40:48 2025

@author: mm4114
"""
from lampyr.config import Config

class AbstractManager():
    """
    Base class for all Lampyr manager objects.

    Provides consistent initialisation whether the manager is instantiated
    standalone (for scripted use) or as part of a :class:`~lampyr.main.Lampyr`
    instance.  Subclasses override :meth:`start` to perform manager-specific
    setup.

    Attributes
    ----------
    config : Config
        Application configuration (from lampyr or freshly constructed).
    _input_func : callable
        Function used to prompt for user input.
    _output_func : callable
        Function used for text output.
    lampyr : Lampyr or None
        Parent Lampyr instance, or ``None`` if running standalone.
    """

    def __init__(self, lampyr = None, config = None):
        """
        Initialise the manager, wiring up config and I/O functions.

        Parameters
        ----------
        lampyr : Lampyr, optional
            Parent Lampyr instance.  If provided, config and I/O functions
            are taken from it.
        config : Config, optional
            Explicit config object to use when running standalone.  If both
            ``lampyr`` and ``config`` are ``None``, a fresh :class:`Config`
            is created.
        """
        if lampyr is not None:
            self.config = lampyr.config
            self._input_func = lampyr._input_func
            self._output_func = lampyr._output_func
            self.lampyr = lampyr
        if lampyr is None:
            if config is not None:
                self.config = config
            else:
                self.config = Config()
            self._input_func = input
            self._output_func = print
            self.lampyr = None
            
        self.start()
    
    def start(self):
        """
        Manager-specific startup hook called at the end of ``__init__``.

        Base implementation is a no-op.  Subclasses override to perform
        initialisation that requires config or lampyr to be set.
        """
        pass
```
