## Extended Data for "A Low-Cost, Modular Hardware and Software Platform for Head-Fixed Mouse Decision-Making Tasks": data.py

```
# -*- coding: utf-8 -*-
"""
Created on Mon Aug 25 18:40:05 2025

@author: mm4114
"""

from lampyr.managers.abstract import AbstractManager
from lampyr.files import savemousefile, loadmousefile, loadsessionfile, savesessionfile, loadjson, savejson
import os
from pathlib import Path
import json
import glob
from lampyr.primatives import Session, Mouse
import shutil
import csv
import hashlib
from datetime import datetime

from typing import List


def hash_file(file_path, algo='sha256', block_size=65536):
    """
    Compute the hash digest of a file.

    Reads the file in fixed-size chunks to avoid loading large files into
    memory all at once.

    Parameters
    ----------
    file_path : str or os.PathLike
        Path to the file to hash.
    algo : str, optional
        Hash algorithm name accepted by :func:`hashlib.new`. Default is
        ``'sha256'``.
    block_size : int, optional
        Number of bytes to read per chunk. Default is 65536 (64 KiB).

    Returns
    -------
    str
        Hex-encoded digest string of the file contents.
    """
    hasher = hashlib.new(algo)
    with open(file_path, 'rb') as f:
        while chunk := f.read(block_size):
            hasher.update(chunk)

    return hasher.hexdigest()


def hashcheck_copyoverwrite(sourcefile, targetfile):
    """
    Copy a source file to a target path only if the contents differ.

    If the target already exists and its hash matches the source, no copy is
    performed. This avoids unnecessary writes when the files are identical.

    Parameters
    ----------
    sourcefile : str or os.PathLike
        Path to the file to copy from. Must exist.
    targetfile : str or os.PathLike
        Path to copy the file to. Will be created or overwritten if the
        contents differ from the source.

    Returns
    -------
    bool
        ``True`` if the file was copied, ``False`` if it was skipped because
        the target already contained identical content.
    """
    if os.path.exists(targetfile):
        if hash_file(sourcefile) == hash_file(targetfile):
            return False
    shutil.copy(sourcefile, targetfile)
    return True


class DataHandler(AbstractManager):
    CONFIG_FAILSAFE_DEFAULT = {'sessions': [],
                               'files': [], }

    def start(self):
        """
        DataHandler startup. If this object is instantiated with a config file,
        determine if backups and failsafes are enabled, then execute those methods.


        Returns
        -------
        None.

        """
        configured = self.config is not None
        haslampyr = self.lampyr is not None
        if not configured:
            return
        self.enable_failsafe = self.config.get(
            'lampyr.enable_saveload_failsafe') and haslampyr
        self.enable_localbackup = self.config.get(
            'lampyr.enable_local_mouse_backups') and haslampyr

        # Create required information for backups and cleanup
        self.local_save_dir = self.config._APP_DATA_DIR
        self.config_failsafe_data = self.config.load_extended_config('data_failsafe',
                                                                     default=self.CONFIG_FAILSAFE_DEFAULT)
        # Check if
        m_dir_present = os.path.exists(
            self.config.get('lampyr.mice_directory'))
        if m_dir_present and self.enable_localbackup:
            self._backupmice()
        if m_dir_present and self.enable_failsafe:
            self._runfailsafecleanup()

    def _backupmice(self):
        """
        Copy all mouse data files from the shared directory to local app data.

        Iterates over every mouse returned by :meth:`mouselist` and uses
        :func:`hashcheck_copyoverwrite` to copy the metadata JSON and, where
        present, the history CSV to the local AppData backup directory. Files
        that are already up to date (matching hash) are skipped silently.

        Mice that have no history CSV on disk (e.g. ``UNKNOWN_MOUSE``) are
        handled gracefully — only the JSON is backed up.

        Per-mouse errors are caught and reported via ``_output_func`` without
        interrupting the backup of remaining mice.

        Returns
        -------
        None
        """
        if not self.enable_localbackup:
            return
        miceids, _ = self.mouselist()
        apdir = self.config._APP_DATA_DIR
        data_dir = self.config.get('lampyr.mice_directory')
        for mid in miceids:
            try:
                mfile_bname = os.path.join(data_dir,
                                           mid,
                                           mid)
                bup_mfile_bname = os.path.join(apdir,
                                               mid)
                mfmove = hashcheck_copyoverwrite(
                    f'{mfile_bname}_mouse.lampyr.json',
                    f'{bup_mfile_bname}_mouse.lampyr.json')
                src_csv = f'{mfile_bname}_history.lampyr.csv'
                hfmove = (hashcheck_copyoverwrite(src_csv,
                          f'{bup_mfile_bname}_history.lampyr.csv')
                          if os.path.exists(src_csv) else False)
                if mfmove or hfmove:
                    self._output_func(f'Updated local backup of {mid}.')
            except FileNotFoundError:
                self._output_func(
                    f'FAILED TO BACK UP {mid} due to FILE NOT FOUND')
            except PermissionError:
                self._output_func(
                    f'FAILED TO BACK UP {mid} due to PERMISSION DENIED')
            except Exception as e:
                self._output_func(
                    f'FAILED TO BACK UP {mid} due to UNEXPECTED ERROR')
                self._output_func(str(e))

    def _runfailsafecleanup(self):
        """
        Replay any session file copies that failed during a previous run.

        Reads the ``data_failsafe`` extended config, which records sessions
        whose files could not be moved to their target directory. For each
        valid entry, the source files are copied to the intended target using
        :func:`shutil.copy`.

        Entries are skipped (with a warning via ``_output_func``) if they are
        not a ``dict`` or are missing the required ``'fps'`` and ``'target'``
        keys. In that case the user is instructed to inspect the remaining
        files manually.

        Returns
        -------
        None
        """
        if not self.enable_failsafe:
            return
        failsafe = self.config.load_extended_config('data_failsafe',
                                                    default=self.CONFIG_FAILSAFE_DEFAULT)
        failed_sessions = failsafe.get('sessions')
        for session in failed_sessions:
            if not isinstance(session, dict):
                self._output_func(
                    'DataHandler found an invalid session failsafe entry!!!')
                continue
            if not {'fps', 'target'} <= session:
                self._output_func(
                    'DataHandler found an invalid session failsafe entry!!!')
                self._output_func(
                    'You must manually inspect and register any remaining failed files.')
                continue
            for fp in session['fps']:
                shutil.copy(fp, session['target'])

    def _logfailure(self, failure_type: str, fps: list, target: str):
        """
        Append a failed file-operation entry to the failsafe config.

        Records the source file paths and intended target directory so that
        :meth:`_runfailsafecleanup` can retry the copy on the next startup.

        Parameters
        ----------
        failure_type : str
            Key in the failsafe config under which to log the failure
            (e.g. ``'sessions'``).
        fps : list of str
            List of source file paths that could not be moved.
        target : str or os.PathLike
            The destination directory or path the files were intended for.

        Returns
        -------
        None
        """
        failures = self.config_failsafe_data.get(failure_type, [])
        failures.append({'fps': fps,
                         'target': target})
        self.config_failsafe_data.set(failure_type, failures)

    def savesession(self, session: Session = None, register=True):
        """
        Save a session to the mouse's session history directory.

        Writes the session files under
        ``<mice_directory>/<mouseid>/lampyr_sessionhistory/`` via
        :func:`~lampyr.files.savesessionfile`. Optionally registers the
        session summary in the mouse's in-memory history via
        :meth:`register_session_to_mouse`.

        Parameters
        ----------
        session : Session, optional
            The session to save. If ``None``, uses the active session from the
            attached lampyr instance. Raises ``KeyError`` if no lampyr instance
            is available.
        register : bool, optional
            If ``True`` (default) and a lampyr instance is attached, the
            session is also registered to the current mouse's history.

        Raises
        ------
        KeyError
            If ``session`` is ``None`` and no lampyr instance is attached.

        Returns
        -------
        None
        """
        if session is None:
            if self.lampyr is None:
                raise KeyError(
                    'Mouse must be specified if outside lampyr instance')
            session = self.lampyr.session
        mouseid = session.mouseid
        data_dir = self.config.get('lampyr.mice_directory')
        dir_fp = os.path.join(data_dir,
                              mouseid,
                              'lampyr_sessionhistory')
        savesessionfile(session, dir_fp)
        self.collect_extended_data(session)
        if register is True and self.lampyr is not None:
            self.register_session_to_mouse(self.lampyr.mouse,
                                           session)

    def loadsession(self, sessionid: str, mouseid: str = None):
        """
        Load a session from a mouse's session history directory.

        Looks up the session files under
        ``<mice_directory>/<mouseid>/lampyr_sessionhistory/`` via
        :func:`~lampyr.files.loadsessionfile`.

        Parameters
        ----------
        sessionid : str
            The unique session ID to load.
        mouseid : str, optional
            The mouse whose session history to search. If ``None``, uses the
            active mouse from the attached lampyr instance. Raises ``KeyError``
            if no lampyr instance is available.

        Raises
        ------
        KeyError
            If ``mouseid`` is ``None`` and no lampyr instance is attached.

        Returns
        -------
        Session
            The reconstructed Lampyr Session object.
        """
        data_dir = self.config.get('lampyr.mice_directory')
        if mouseid is None:
            if self.lampyr is not None:
                mouseid = self.lampyr.mouse.mouseid
            else:
                raise KeyError(
                    'Mouseid must be specified if outside lampyr instance')
        data_dir = os.path.join(data_dir,
                                mouseid,
                                'lampyr_sessionhistory')
        return loadsessionfile(sessionid, data_dir)

    def mouseexists(self, mouseid):
        """
        Check whether a mouse has a saved metadata file on disk.

        Parameters
        ----------
        mouseid : str
            The mouse ID to look up.

        Returns
        -------
        bool
            ``True`` if the mouse's ``_mouse.lampyr.json`` file exists in the
            configured mice directory, ``False`` otherwise.
        """
        return os.path.exists(os.path.join(self.config.get('lampyr.mice_directory'),
                                           mouseid,
                                           f'{mouseid}_mouse.lampyr.json'))

    def mouselist(self) -> List[str]:
        """
        Return all mouse IDs and their metadata file paths found on disk.

        Scans the configured mice directory recursively for
        ``*.lampyr.json`` files and retains only those whose filename stem
        matches the pattern ``<parentdir>_mouse``, identifying them as mouse
        metadata files rather than session files.

        Returns
        -------
        mouseidlist : list of str
            List of mouse ID strings derived from the parent directory names.
        mousepaths : list of str
            Corresponding list of absolute paths to each mouse's
            ``_mouse.lampyr.json`` file.
        """
        data_dir = self.config.get('lampyr.mice_directory')
        candidate_mice = glob.glob(os.path.join(data_dir,
                                                '**',
                                                '*.lampyr.json'))
        mousepaths = [mpath for mpath in candidate_mice
                      if os.path.basename(mpath).split('.')[0] ==
                      os.path.basename(os.path.dirname(mpath)) + '_mouse']
        mouseidlist = [os.path.basename(os.path.dirname(m))
                       for m in mousepaths]
        return mouseidlist, mousepaths

    def savemouse(self, mouse: Mouse = None):
        """
        Save a Mouse object to the configured mice directory.

        Writes the mouse metadata JSON and, if the mouse has session history,
        the history CSV under ``<mice_directory>/<mouseid>/`` via
        :func:`~lampyr.files.savemousefile`.

        Parameters
        ----------
        mouse : Mouse, optional
            The Mouse object to save. If ``None``, uses the active mouse from
            the attached lampyr instance. Raises ``KeyError`` if no lampyr
            instance is available.

        Raises
        ------
        KeyError
            If ``mouse`` is ``None`` and no lampyr instance is attached.

        Returns
        -------
        None
        """
        if mouse is None:
            if self.lampyr is None:
                raise KeyError(
                    'Mouse must be specified if outside lampyr instance')
            mouse = self.lampyr.mouse
        data_dir = self.config.get('lampyr.mice_directory')
        data_dir = os.path.join(data_dir,
                                mouse.mouseid)
        savemousefile(mouse, data_dir)

    def loadmouse(self, mouseid: str):
        """
        Load a Mouse object from the configured mice directory.

        Reconstructs the Mouse from the ``_mouse.lampyr.json`` and, if
        present, the ``_history.lampyr.csv`` files stored under
        ``<mice_directory>/<mouseid>/`` via
        :func:`~lampyr.files.loadmousefile`.

        Parameters
        ----------
        mouseid : str
            The ID of the mouse to load.

        Returns
        -------
        Mouse
            The reconstructed Lampyr Mouse object.
        """
        data_dir = os.path.join(self.config.get('lampyr.mice_directory'),
                                mouseid)
        return loadmousefile(mouseid, data_dir)

    def register_session_to_mouse(self, mouse: Mouse, session: Session):
        """
        Append a session summary entry to a mouse's in-memory history.

        Extracts key performance metrics from the session and appends them as
        a dict to ``mouse.history``. The timestamp is also decomposed into
        year, month, and day fields for convenient downstream filtering.

        Parameters
        ----------
        mouse : Mouse
            The Mouse object whose history will be updated.
        session : Session
            The completed session from which to extract the summary entry.

        Returns
        -------
        None
        """
        sessionentry = {}
        dt = datetime.fromtimestamp(session.starttime).astimezone()
        sessionentry['sessionid'] = session.uniquesessionid
        sessionentry['starttime'] = session.starttime
        sessionentry['year'] = dt.year
        sessionentry['month'] = dt.month
        sessionentry['day'] = dt.day
        sessionentry['rootslug'] = session.segments.get(session.root, {}
                                                        ).get('slug', 'NA')
        for entry in ["merit", "demerit", "duration", "trial", "rewards",
                      "abstention", "participation"]:
            sessionentry[entry] = getattr(session, entry)
        mouse.history.append(sessionentry)

    def collect_extended_data(self, session: Session):
        if session._extendeddata is None:
            return
        mouseid = session.mouseid
        data_dir = self.config.get('lampyr.mice_directory')
        for e in session._extendeddata:
            fname = os.path.basename(e['fp'])
            target_fp = os.path.join(data_dir,
                                     mouseid,
                                     'lampyr_extendeddata',
                                     session.uniquesessionid,
                                     e['type'],
                                     fname)
            os.makedirs(os.path.dirname(target_fp), exist_ok=True)
            shutil.move(e['fp'], target_fp)
            self._output_func('Collected {fname}')

    def _mouse_session_list_from_files(self, mouseid):
        data_dir = self.config.get('lampyr.mice_directory')
        dir_fp = os.path.join(data_dir,
                              mouseid,
                              'lampyr_sessionhistory',
                              '*.lampyr.json')
        all_sessions = glob.glob(dir_fp)
        all_sessions = [Path(p).name.removesuffix('.lampyr.json')
                        for p in all_sessions]
        return all_sessions

    def reconstruct_all_mouse_history_from_files(self):
        mouselist, _ = self.mouselist()
        for mouse in mouselist:
            print(f'Reconstructing {mouse} history')
            mouse_obj = self.loadmouse(mouse)
            mouse_obj.history = []
            if mouse == 'UNKNOWN_MOUSE':
                continue
            sessionlist = self._mouse_session_list_from_files(mouse)
            print(f'Processing {len(sessionlist)} sessions')
            for session in sessionlist:
                try:
                    s = self.loadsession(session, mouse)
                    self.register_session_to_mouse(mouse_obj, s)
                except:
                    print(f'Failed to parse {mouse} - {session}')
            print('Successfully registered ' +
                  f'{len(mouse_obj.history)}/{len(sessionlist)} sessions')
            self.savemouse(mouse_obj)

    def heartbeat_filetouch(self):
        datadir = self.config.get('lampyr.mice_directory')
        hbeatfile = os.path.join(datadir, 'heartbeat.file')
        if not os.path.exists(hbeatfile):
            savejson(hbeatfile, {})
        loadjson(hbeatfile)


class MouseManager(AbstractManager):
    def start(self):
        """
        MouseManager startup.

        Ensures ``UNKNOWN_MOUSE`` exists on disk (creating it if necessary),
        then loads it as the initially active mouse. Raises ``RuntimeError``
        if no lampyr instance is attached, as this manager cannot function
        without one.

        Raises
        ------
        RuntimeError
            If no lampyr instance is attached to this manager.

        Returns
        -------
        None
        """
        if self.lampyr is None:
            raise RuntimeError(
                'MouseManager cannot operate without a lampyr instance')
        self.mouse = None
        if not self.exists('UNKNOWN_MOUSE'):
            self.create('UNKNOWN_MOUSE')
        self.load('UNKNOWN_MOUSE')

    def create(self, mouseid, **kwargs):
        """
        Create a new Mouse, set it as active, and save it to disk.

        Parameters
        ----------
        mouseid : str
            ID to assign to the new mouse.
        **kwargs
            Additional keyword arguments forwarded to the :class:`Mouse`
            constructor.

        Returns
        -------
        None
        """
        mouse = Mouse(mouseid=mouseid, **kwargs)
        self.mouse = mouse
        self.save()

    def retire(self):
        """
        Marks mouse as retired

        Parameters
        ----------
        mouseid : str
            DESCRIPTION.

        Returns
        -------
        None.

        """
        self.mouse.retired = True
        self.save()

    def deretire(self):
        self.mouse.retired = False
        self.save()

    def list(self):
        """
        Return a list of all mouse IDs found in the mice directory.

        Returns
        -------
        list of str
            Mouse IDs discovered on disk.
        """
        mlist, _ = self.lampyr.datamanager.mouselist()
        return mlist

    def exists(self, mouseid):
        """
        Check whether a mouse exists on disk.

        Parameters
        ----------
        mouseid : str
            The mouse ID to look up.

        Returns
        -------
        bool
            ``True`` if the mouse's metadata file is present, ``False``
            otherwise.
        """
        return self.lampyr.datamanager.mouseexists(mouseid)

    def load(self, mouseid):
        """
        Load a mouse from disk and set it as the active mouse.

        Parameters
        ----------
        mouseid : str
            The ID of the mouse to load.

        Returns
        -------
        None
        """
        self.mouse = self.lampyr.datamanager.loadmouse(mouseid)

    def save(self):
        """
        Save the currently active mouse to disk.

        Returns
        -------
        None
        """
        self.lampyr.datamanager.savemouse(self.mouse)


if __name__ == '__main__':
    dhandler = DataHandler()
    print(dhandler.loadmouse('014-000'))
```
