## Extended Data for "A Low-Cost, Modular Hardware and Software Platform for Head-Fixed Mouse Decision-Making Tasks": hardware.py

```
# -*- coding: utf-8 -*-
"""
Created on Mon Aug 25 18:40:27 2025

@author: mm4114
"""

from lampyr.managers.abstract import AbstractManager
from lampyr.rigs.rigcontrol import ArduinoBanditRig_0
from lampyr.rigs.abstract import all_rig_definitions, AbstractHardwareRig
from lampyr.rigs.banditrig import BanditRig

import numpy as np
import time


class RigManager(AbstractManager):
    """
    Manages connection to and calibration of the physical rig.

    Attributes
    ----------
    rig : ArduinoBanditRig_0 or None
        The connected rig object; ``None`` until :meth:`connect` is called.
    connected : bool
        Whether a rig is currently connected.
    """

    def start(self):
        """Initialise rig and connection state to disconnected."""
        all_rigs = all_rig_definitions()
        rig_type = self.config.get('rig.rig_type')
        self.rig_cls = all_rigs.get(rig_type, BanditRig)
        self.rig = None
        self.connected = False

    def connect(self):
        """
        Connect to the Arduino rig and apply the stored sipper calibration.

        Instantiates :class:`~lampyr.rigs.rigcontrol.ArduinoBanditRig_0`,
        starts the serial listener thread, and sets the dispenser size from
        ``config['rig.sipper_calib']``.
        """
        self._output_func('Loading rig config')
        self._output_func('Connecting to Arduino Rig...')
        self.rig = self.rig_cls(config = self.config)
        self._output_func('Creating serial monitor thread...')
        self.rig.start()
        self._output_func('Setting stored rig sipper calibration...')
        self.rig.reward.setsize(self.config.get('rig.sipper_calib')) #to be removed
        self.connected = True

    def disconnect(self):
        """
        Stop the serial monitor thread and close the rig serial port.
        """
        if self.rig is None:
            return
        if not self.connected:
            self._output_func('Rig is already shut down...')
            return
        self._output_func('Closing rig monitoring threads...')
        self.rig.stop()
        self.rig.disconnect()
        self.connected = False

    def calibrate(self):
        """
        Interactively calibrate the sipper dispenser to deliver ~5 µl per reward.

        Performs a multi-point gravimetric calibration:

        1. Tests three dispense sizes around the current estimate.
        2. Fits a linear regression to the measured volumes.
        3. Solves for the size that produces 5 µl.
        4. Validates with a final test dispense.
        5. Repeats until ``|measured_volume - 0.005 ml| < 0.0005 ml``.

        Stores the calibrated size in ``config['rig.sipper_calib']`` and
        updates ``config['rig.calibrated']`` with the current timestamp.
        Connects to the rig first if not already connected, and disconnects
        afterwards if it was not connected before.
        """
        was_connected = self.connected
        if not self.connected:
            self.connect()

        def linreg(x, y):
            x = np.asarray(x)
            y = np.asarray(y)
            a, b = np.polyfit(x, y, 1)
            y_pred = a * x + b
            r2 = 1 - np.sum((y - y_pred)**2) / np.sum((y - np.mean(y))**2)

            return a, b, r2

        def inputfloat(prompt):
            while True:
                val = self._input_func(prompt)
                try:
                    number = float(val)
                    break
                except ValueError:
                    self._output_func("Please enter a valid number.")
            return number

        def calib_disp(disp_size, initial_value=None):
            disp_size = int(disp_size)
            self.rig.reward.setsize(disp_size)
            if initial_value is None:
                initial_value = inputfloat('INPUT WEIGHT (g): ')
                if initial_value == 0.0:
                    self._output_func("Zero entered — restarting calibration cycle.")
                    return None
            time.sleep(0.1)
            for i in range(200):
                self.rig.reward.give()
                time.sleep(0.3)
            new_value = inputfloat('INPUT NEW WEIGHT (g):')
            if new_value == 0.0:
                self._output_func("Zero entered — restarting calibration cycle.")
                return None
            dvol = (new_value - initial_value) / 200
            self._output_func(f'Reward Size: {disp_size} produces {str(dvol)[:10]} ml reward')
            return dvol, new_value

        try:
            est_sipp = None
            while True:
                self._output_func('\nBEGINING CALIBRATION')
                if est_sipp is None:
                    try:
                        est_sipp = int(self.config.get('rig.sipper_calib'))
                    except:
                        est_sipp = 10000
                if est_sipp <= 6000:
                    est_sipp = 10000
                dsizes = [int(est_sipp*(2/3)), est_sipp, int(est_sipp*1.5)]
                dvols = []
                next_initial = None
                restart = False
                for disp_size in dsizes:
                    result = calib_disp(disp_size, initial_value=next_initial)
                    if result is None:
                        restart = True
                        break
                    dvol, next_initial = result
                    dvols.append(dvol)
                if restart:
                    continue
                slope, coeff, r2 = linreg(dsizes, dvols)
                if r2 < 0.9:
                    self._output_func('Failed to produce linear regression. Repeating Calibration.')
                    continue
                est_sipp = int((0.005 - coeff) / slope)
                self._output_func(f'Estimated correct reward size is {est_sipp}')
                self._output_func('Beginning dispenser test...')
                result = calib_disp(est_sipp, initial_value=next_initial)
                if result is None:
                    continue
                dvol, _ = result
                if abs(0.005 - dvol) < 0.0005:
                    break
                else:
                    self._output_func('Calibration failed. Repeating calibration.')
            self._output_func('Calibration success')
            self._output_func(f'Rig reward size is set to {est_sipp}')
            self.config.set('rig.sipper_calib', est_sipp)
            self.config.set('rig.calibrated', round(time.time()))
        finally:
            if not was_connected:
                self.disconnect()
```
