## Extended Data for "A Low-Cost, Modular Hardware and Software Platform for Head-Fixed Mouse Decision-Making Tasks": notification.py

```
# -*- coding: utf-8 -*-
"""
Created on Mon Aug 25 18:44:53 2025

@author: mm4114
"""

from lampyr.managers.abstract import AbstractManager
from lampyr.files import loadjson, savejson
import requests

class NotificationManager(AbstractManager):
    """
    Manages push notifications via the Pushover API.

    Notifications are sent to every active user (supervisors are always active)
    registered in the shared ``users.json`` config file.

    Attributes
    ----------
    userdata : ConfigFile
        Shared user database containing Pushover credentials.
    """

    def start(self):
        """Load the shared user database."""
        self.userdata = self.config.load_shared_extended_config('users')

    def add_user(self, name, pushover_user_key, supervisor=False, active=True):
        """
        Register a new user in the shared user database.

        Parameters
        ----------
        name : str
            Unique user name.
        pushover_user_key : str
            Pushover user key for the account.
        supervisor : bool, optional
            If ``True``, this user always receives notifications and cannot be
            deactivated.  Default is ``False``.
        active : bool, optional
            If ``False``, the user is excluded from notifications.  Ignored for
            supervisors, who are always active.  Default is ``True``.
        """
        self.userdata._config[name] = {
            'pushover_user_key': pushover_user_key,
            'supervisor': supervisor,
            'active': True if supervisor else active,
        }
        self.userdata.save()

    def set_app_token(self, token):
        """Store the shared Pushover application token in the users config."""
        self.userdata._config['_app_token'] = token
        self.userdata.save()

    def set_user_active(self, name, active):
        """
        Set the active/inactive state of a user.

        Parameters
        ----------
        name : str
            User name to update.
        active : bool
            ``True`` to enable notifications for this user, ``False`` to disable.

        Raises
        ------
        KeyError
            If ``name`` is not in the user database.
        """
        if name not in self.userdata._config:
            raise KeyError(f"User {name!r} not found.")
        if self.userdata._config[name].get('supervisor'):
            raise ValueError(f"User {name!r} is a supervisor and cannot be deactivated.")
        self.userdata._config[name]['active'] = active
        self.userdata.save()

    def delete_user(self, name):
        """
        Remove a user from the shared user database.

        Parameters
        ----------
        name : str
            User name to delete.

        Raises
        ------
        KeyError
            If ``name`` is not in the user database.
        """
        if name not in self.userdata._config:
            raise KeyError(f"User {name!r} not found.")
        del self.userdata._config[name]
        self.userdata.save()

    def _get_targets(self):
        """
        Build the list of user names that should receive the next notification.

        Returns every active user (supervisors are always active).

        Returns
        -------
        list of str
            Names to notify.
        """
        self.userdata.reload()
        all_users = {n: d for n, d in self.userdata.to_dict().items() if isinstance(d, dict)}
        return [n for n, d in all_users.items() if d.get('active', True) or d.get('supervisor')]

    def _send_to_user(self, name, message, title, urgent=False):
        """
        Send a Pushover notification to a single user.

        Silently skips users whose credentials are not configured.

        Parameters
        ----------
        name : str
            User name to look up in the user database.
        message : str
            Notification body text.
        title : str
            Notification title.
        urgent : bool, optional
            If ``True``, sends at priority 2 (Emergency), which bypasses Do Not
            Disturb and retries every 30 s for up to 1 hour until acknowledged.

        Raises
        ------
        RuntimeError
            If the Pushover API returns a non-200 status code.
        """
        self.userdata.reload()
        try:
            app_token = self.userdata.get('_app_token')
        except KeyError:
            print('Pushover app token not set — run: lampyr user set-token <token>')
            return
        try:
            user_key = self.userdata.get(f'{name}.pushover_user_key')
        except KeyError:
            print(f'Pushover user key not configured for {name!r} — skipping.')
            return
        payload = {"token": app_token, "user": user_key, "title": title, "message": message}
        if urgent:
            payload.update({"priority": 2, "retry": 30, "expire": 10800})
        response = requests.post("https://api.pushover.net/1/messages.json", data=payload)
        if response.status_code != 200:
            raise RuntimeError(f"Pushover notification failed for {name!r}: {response.text}")
        print(f"Notification sent to {name!r}.")

    def send_notification(self, message, title="Notification"):
        """
        Send a notification to all target users.

        Parameters
        ----------
        message : str
            Notification body.
        title : str, optional
            Notification title. Default is ``'Notification'``.
        """
        targets = self._get_targets()
        if not targets:
            print('Notifications not configured — skipping push notification.')
            return
        for name in targets:
            self._send_to_user(name, message, title)
        

if __name__ == '__main__':
    NotificationManager().send_notification('test','Lampyr')
```
