## Extended Data for "A Low-Cost, Modular Hardware and Software Platform for Head-Fixed Mouse Decision-Making Tasks": plugins.py

```
# -*- coding: utf-8 -*-
"""
Created on Mon Aug 25 19:44:11 2025

@author: mm4114
"""
from lampyr.managers.abstract import AbstractManager

import importlib.util
from pathlib import Path

class PluginManager(AbstractManager):
      def start(self):
          if self.config.get('lampyr.plugin_folder') is None:
              return
          self._load_directory()
          

      def _load_directory(self):
          fp = self.config.get('lampyr.plugin_folder')
          for path in Path(fp).glob("*.py"):
            if path.stem.startswith("_"):
                continue
            spec = importlib.util.spec_from_file_location(f"plugin_{path.stem}", path)
            mod = importlib.util.module_from_spec(spec)
            spec.loader.exec_module(mod)
```
