## Extended Data for "A Low-Cost, Modular Hardware and Software Platform for Head-Fixed Mouse Decision-Making Tasks": math.py

```
# -*- coding: utf-8 -*-
"""
Created on Mon Jun 15 14:58:17 2026

@author: mm4114
"""
import random, math

def rand_int_uniform_hazard(min_val=1, max_val=10):
    """
    Samples from a truncated geometric distribution using the
    ideal hazard for the specified range.
    """
    h = _ideal_hazard(min_val, max_val)

    u = random.random()

    k = math.floor(
        math.log(1 - u) /
        math.log(1 - h)
    )

    return min(min_val + k, max_val)

def _ideal_hazard(min_val=1, max_val=10):
    N = max_val - min_val + 1

    if N == 1:
        return 1.0

    return 1 - N**(-1/(N-1))
```
