## Extended Data for "A Low-Cost, Modular Hardware and Software Platform for Head-Fixed Mouse Decision-Making Tasks": primatives.py

```
# -*- coding: utf-8 -*-
"""
Created on Tue Jun 24 11:33:55 2025

@author: mm4114
"""

from dataclasses import dataclass, field
import time
from copy import deepcopy
from typing import List
import random


def uniqueid(type, name=None):
    """
    Generate a unique identifier string with an optional name component.

    Parameters
    ----------
    type : str
        Category prefix for the identifier (e.g. ``'session'``, ``'segment'``).
    name : str, optional
        Additional label to embed in the identifier. Omitted if ``None``.

    Returns
    -------
    str
        Unique identifier of the form
        ``'<type>_[<name>_]created<unix_timestamp>_<random6digits>'``.
    """
    uid = f'{type}_'
    if name is not None:
        uid += f'{name}_'
    uid += f'created{round(time.time())}_{random.randint(0, 999999):06d}'
    return uid


@dataclass
class Mouse:
    """
    Represents a single experimental subject.

    Attributes
    ----------
    mouseid : str
        Unique identifier for the mouse (e.g. ``'014-000'``).
    mouse_behav_param_overrides : dict
        Per-slug or per-tag behavioral parameter overrides applied at segment
        initialisation. Keys are slug/tag strings; values are dicts of
        ``{param: value}`` mappings.
    paradigm : str or None
        Name of the paradigm currently assigned to this mouse.
    paradigm_stage : dict
        Maps paradigm tags to the mouse's current stage string within that
        paradigm (e.g. ``{'BanditParadigm': 'AnyWheel'}``).
    properties : dict
        Paradigm-specific persistent data keyed by paradigm slug.
    history : list of dict
        Chronological list of session summary entries appended after each
        completed session.
    """
    mouseid: str = '014-000'
    retired: bool = False
    mouse_behav_param_overrides: dict = field(default_factory=dict)
    paradigm: str = None
    properties: dict = field(default_factory=dict)
    history: List = field(default_factory=list)


@dataclass
class Session:
    """
    Tracks all counters, limits, metadata, and segment records for a single
    experimental session.

    Each scalar counter (``merit``, ``trial``, ``rewards``, etc.) has an
    optional ``_min`` and ``_limit`` companion.  ``evaluatestopconditions``
    uses these to decide when to end a session.

    The session becomes immutable once :meth:`lock` is called (i.e. after
    ``endtime`` is set).  Any subsequent ``__setattr__`` call raises
    ``RuntimeError``.

    Attributes
    ----------
    merit_limit, merit_min, merit : int or None
        Upper limit, lower minimum, and running count of merit events.
    demerit_limit, demerit_min, demerit : int or None
        Upper limit, lower minimum, and running count of demerit events.
    duration_limit, duration_min, duration : int or None
        Upper limit (minutes), lower minimum (minutes), and elapsed duration.
    trial_limit, trial_min, trial : int or None
        Upper limit, lower minimum, and running trial count.
    reward_limit, reward_min, rewards : int or None
        Upper limit, lower minimum, and running reward count.
    abstention_limit, abstention_min, abstention : int or None
        Upper limit, lower minimum, and running abstention count.
    participation_limit, participation_min, participation : int or None
        Upper limit, lower minimum, and running participation count.
    serial_abstention_limit, serial_abstention_min, serial_abstention : int or None
        Upper limit, lower minimum, and consecutive abstention count (resets
        on each participation event).
    starttime : float
        Unix timestamp when the session was created.
    endtime : float or None
        Unix timestamp set by :meth:`lock`; ``None`` while session is active.
    uniquesessionid : str
        Unique identifier generated at creation.
    mouseid : str or None
        ID of the mouse that ran this session.
    mouseid_unique : str or None
        Secondary unique mouse identifier.
    root : str or None
        ``uniqueid`` of the top-level segment that ran this session.
    segmentlist : list of str
        Ordered list of segment ``uniqueid`` strings added during the session.
    segments : dict
        Maps segment ``uniqueid`` to the serialised segment data dict.
    eventlist : list of dict
        Chronological list of event records written by :meth:`Trial.trigger_event`.
    rigdata : dict or None
        Raw rig data snippet extracted at session end.
    """
    # Session limits and minimums
    merit_limit: int = None
    merit_min: int = None
    merit: int = 0
    demerit_limit: int = None
    demerit_min: int = None
    demerit: int = 0
    duration_limit: int = None
    duration_min: int = None
    duration: float = None
    trial_limit: int = None
    trial_min: int = None
    trial: int = 0
    reward_limit: int = None
    reward_min: int = None
    rewards: int = 0
    reward_amount_limit: float = None
    reward_amount_min: float = None
    reward_amount: float = 0.0
    abstention_limit: int = None
    abstention_min: int = None
    abstention: int = 0
    participation_limit: int = None
    participation_min: int = None
    participation: int = 0
    serial_abstention_limit: int = None
    serial_abstention_min: int = None
    serial_abstention: int = 0

    # General information
    starttime: float = field(default_factory=time.time)
    endtime: float = None
    uniquesessionid: str = field(default_factory=lambda: uniqueid('session'))

    # Mouseid
    mouseid: str = None
    mouseid_unique: str = None

    # Segment records
    root: str = None
    segmentlist: list = field(default_factory=list)
    segments: dict = field(default_factory=dict)

    # Session Events and rig data
    eventlist: List = field(default_factory=list)
    rigdata: dict = None
    rigproperties: dict = None
    extendeddata: list = None  # populated on shutdown
    _extendeddata: list = None

    def __setattr__(self, key, value):
        """
        Prevent any attribute modification after the session has been locked.

        Parameters
        ----------
        key : str
            Attribute name to set.
        value : object
            Value to assign.

        Raises
        ------
        RuntimeError
            If the session has already been finalised (``endtime`` is not ``None``).
        """
        if hasattr(self, "endtime") and self.endtime is not None:
            raise RuntimeError(f"Cannot modify '{key}'; session is finalized.")
        super().__setattr__(key, value)

    def lock(self):
        """
        Finalise the session by recording the end time.

        Sets ``endtime`` to the current Unix timestamp, making the session
        immutable.  Any subsequent attribute modification will raise
        ``RuntimeError``.
        """
        self.duration = (time.time() - self.starttime) / 60
        self.endtime = time.time()

    def __repr__(self):
        """
        Return a human-readable summary of session counters and limits.

        Returns
        -------
        str
            Multi-line string with one row per counter showing current value,
            minimum, and maximum, followed by timing and identity fields.
        """
        fields = [
            ("merit", self.merit_min, self.merit_limit, self.merit),
            ("demerit", self.demerit_min, self.demerit_limit, self.demerit),
            ("duration", self.duration_min, self.duration_limit,
             0 if self.duration is None else round(self.duration, 2)),
            ("trial", self.trial_min, self.trial_limit, self.trial),
            ("reward", self.reward_min, self.reward_limit, self.rewards),
            ("reward_amount", self.reward_amount_min,
             self.reward_amount_limit, round(self.reward_amount, 2)),
            ("abstention", self.abstention_min,
             self.abstention_limit, self.abstention),
            ("participation", self.participation_min,
             self.participation_limit, self.participation),
            ("serial_abstention", self.serial_abstention_min,
             self.serial_abstention_limit, self.serial_abstention),
        ]
        msg = ''
        for name, min_val, max_val, val in fields:
            msg += f"{name:22}: {str(val):4} (Min: {str(min_val)                                                    :6} Max: {str(max_val):6})\n"
        msg += f"{'starttime':22}: {self.starttime}\n"
        msg += f"{'endtime':22}: {self.endtime}\n"
        msg += f"{'uniquesessionid':22}: {self.uniquesessionid}\n"
        msg += f"{'mouseid':22}: {self.mouseid}\n"
        msg += f"{'mouseid_unique':22}: {self.mouseid_unique}\n"
        return msg

    def search(self, root=None,
               type=None, types=None,
               slug=None, slugs=None,
               custom_checks_permissive=None,
               custom_checks_strict=None,
               sort=None, return_objects=False,
               filt=None,
               _seglist=None):
        """
        Recursively search the segment tree for segments matching given criteria.

        Traverses the tree starting from ``root``, collecting segment IDs (or
        objects) that satisfy all supplied filters.

        Parameters
        ----------
        root : str, optional
            ``uniqueid`` of the segment to start searching from.  Defaults to
            the session root.
        type : str, optional
            Single segment type string that must appear in
            ``segment['segment_type']``.
        types : list of str, optional
            Any one of these type strings must appear in
            ``segment['segment_type']`` (permissive OR match).
        slug : str, optional
            Single slug that must match ``segment['slug']``.
        slugs : list of str, optional
            Any one of these slugs must match (permissive OR match).
        custom_checks_permissive : dict, optional
            ``{field: [values]}`` — any value in the list may match the
            segment field (permissive OR logic per field).
        custom_checks_strict : dict, optional
            ``{field: [values]}`` — all values in the list must be present in
            the segment field (strict AND logic per field).
        sort : str, optional
            If given, sort matching segment IDs by this field from the segment
            dict.
        return_objects : bool, optional
            If ``True``, return segment dicts instead of ID strings.
        filt : callable, optional
            Additional filter ``filt(segment_dict) -> bool`` applied after all
            other checks.
        _seglist : list, optional
            Internal accumulator; do not pass explicitly.

        Returns
        -------
        list of str or list of dict
            Matching segment IDs, or segment dicts when ``return_objects=True``.
        """
        if _seglist is None:
            _seglist = []
        if root is None:
            root = self.root
        segment = self.segments[root]

        custom_checks_permissive = custom_checks_permissive or {}
        custom_checks_strict = custom_checks_strict or {}

        def search_filter(singleton, multioption, target, strict=False):
            if singleton is None and multioption is None:
                return True
            if not isinstance(target, (list, tuple, set, dict)):
                target = [target]
            if singleton is not None:
                if singleton not in target:
                    return False
            if multioption is not None:
                for option in multioption:
                    if strict:
                        if option not in target:
                            return False
                    else:
                        if option in target:
                            return True
                if strict:
                    return True
                else:
                    return False
            return True

        valid = search_filter(type, types, segment['segment_type'])
        valid = valid and search_filter(slug, slugs, segment['slug'])
        for check, vals in custom_checks_permissive.items():
            if check not in segment:
                valid = False
                break
            valid = valid and search_filter(None, vals, segment[check],
                                            strict=False)
        for check, vals in custom_checks_strict.items():
            if check not in segment:
                valid = False
                break
            valid = valid and search_filter(None, vals, segment[check],
                                            strict=True)

        if valid:
            _seglist.append(root)
        for subseg in segment['subdata']:
            self.search(root=subseg,
                        type=type,
                        types=types,
                        slug=slug,
                        slugs=slugs,
                        custom_checks_permissive=custom_checks_permissive,
                        custom_checks_strict=custom_checks_strict,
                        _seglist=_seglist)
        if sort is not None:
            _seglist = sorted(_seglist, key=lambda x: self.segments[x][sort])
        if filt is not None:
            _seglist = [seg for seg in _seglist
                        if filt(self.segments[seg])]
        if return_objects:
            return [self.segments[seg] for seg in _seglist]
        return _seglist

    @property
    def segment_types(self):
        """
        Return all unique segment type strings present in this session.

        Returns
        -------
        list of str
            Deduplicated list of type strings drawn from every segment's
            ``'segment_type'`` field.
        """
        typs = set()
        for seg in self.segments.values():
            for typ in seg['segment_type']:
                typs.add(typ)
        return list(typs)

    def evaluatestopconditions(self):
        """
        Evaluate all session stop conditions and return any that are triggered.

        Reward limits are checked first regardless of minimums, to protect
        against accidental over-delivery.  All ``_min`` requirements must be
        satisfied before any ``_limit`` triggers are tested.

        Returns
        -------
        list of str
            Names of the stop conditions currently met.  An empty list means
            the session should continue.  Possible values include
            ``'reward'``, ``'trial'``, ``'duration'``, ``'merit'``,
            ``'demerit'``, ``'participation'``, ``'abstention'``,
            ``'serialabstention'``.
        """
        stops = []
        if self.endtime is None:
            self.duration = (time.time() - self.starttime) / 60
        duration = self.duration

        # Reward limits are privileged to avoid senarios where water restriction is broken
        if self.reward_limit is not None:
            if self.rewards >= self.reward_limit:
                stops.append('reward')
        if self.reward_amount_limit is not None:
            if self.reward_amount >= self.reward_amount_limit:
                stops.append('reward')

        # All minimum requirements must be met before limits are tested
        if self.trial_min is not None:
            if self.trial < self.trial_min:
                return stops
        if self.duration_min is not None:
            if duration < self.duration_min:
                return stops
        if self.participation_min is not None:
            if self.participation < self.participation_min:
                return stops
        if self.merit_min is not None:
            if self.merit < self.merit_min:
                return stops
        if self.demerit_min is not None:
            if self.demerit < self.demerit_min:
                return stops
        if self.abstention_min is not None:
            if self.abstention < self.abstention_min:
                return stops
        if self.serial_abstention_min is not None:
            if self.serial_abstention < self.serial_abstention_min:
                return stops
        if self.reward_amount_min is not None:
            if self.reward_amount < self.reward_amount_min:
                return stops
        if self.reward_min is not None:
            if self.rewards < self.reward_min:
                return stops

        # Test all limits and report reasons for stop
        if self.trial_limit is not None:
            if self.trial >= self.trial_limit:
                stops.append('trial')
        if self.duration_limit is not None:
            if duration >= self.duration_limit:
                stops.append('duration')
        if self.merit_limit is not None:
            if self.merit >= self.merit_limit:
                stops.append('merit')
        if self.demerit_limit is not None:
            if self.demerit >= self.demerit_limit:
                stops.append('demerit')
        if self.participation_limit is not None:
            if self.participation >= self.participation_limit:
                stops.append('participation')
        if self.abstention_limit is not None:
            if self.abstention >= self.abstention_limit:
                stops.append('abstention')
        if self.serial_abstention_limit is not None:
            if self.serial_abstention >= self.serial_abstention_limit:
                stops.append('serialabstention')
        return stops
```
