## Extended Data for "A Low-Cost, Modular Hardware and Software Platform for Head-Fixed Mouse Decision-Making Tasks": abstract.py

```
# -*- coding: utf-8 -*-
"""
Created on Tue May 19 15:53:37 2026

@author: mm4114

NOT YET IMPLEMENTED

"""
from abc import ABC, abstractmethod
from threading import Lock
from collections import namedtuple
import os, time

from typing import List

def all_rig_definitions():
    def subclasses(cls):
        for sub in cls.__subclasses__():
            yield sub
            yield from subclasses(sub)
    rigs = {sub.__name__ : sub for sub in subclasses(AbstractHardwareRig)}
    return rigs

class AbstractHardwareRig(ABC):
    def __init__(self, *args, config = None, **kwargs):
        self.interfaces = {}
        self.components = {}
        self.config = config or {} # this is expected to be the lampyr config object
        self.setup(*args, **kwargs)
    
    @abstractmethod
    def setup(self, *args, **kwargs):
        pass
    
    @abstractmethod
    def is_calibrated(self):
        pass
    
    @abstractmethod
    def is_configured(self):
        pass
    
    @abstractmethod
    def calibrate(self):
        pass
    
    @abstractmethod
    def configure(self):
        pass
    
    def register_component(self, name, component_obj):
        if hasattr(self, name):
            raise ValueError('That name is already reserved')
        setattr(self, name, component_obj)
        self.components[name] = component_obj
    
    def register_interface(self, name, interface_obj):
        if name in self.interfaces:
            raise ValueError(f'Attempted to register {name} twice')
        self.interfaces[name] = interface_obj
    
    def start(self):
        for interface in self.interfaces.values():
            interface.start()
    
    def stop(self):
        for component in self.components.values():
            component.stop()
        time.sleep(2)
        for interface in self.interfaces.values():
            interface.stop()
    
    def disconnect(self):
        for interface in self.interfaces.values():
            interface.disconnect()
    
    def dump(self):
        accumulated_json_data = {}
        accumulated_array_data = {}
        extended_data_files = []
        accumulated_json_data['RIG_TYPE'] = self.__class__.__name__
        for interfacename, interface in self.interfaces.items():
            accumulated_json_data[interfacename] = interface.dump()
            accumulated_json_data[interfacename]['INTERFACE_TYPE'] = \
                interface.__class__.__name__
            accumulated_json_data[interfacename]['REPORTS'] = []
            extendeddata = interface.extendeddata
            
            accumulated_json_data[interfacename]['EXTENDED_DATA'] = extendeddata.copy()
            extended_data_files.extend(extendeddata.copy())
            
            for reporttype in interface.data.reports:
                if reporttype not in accumulated_array_data:
                    accumulated_array_data[reporttype] = interface.data.reports[reporttype]
                    accumulated_json_data[interfacename]['REPORTS'].append(reporttype)
                else:
                    rtype = f'{interfacename}_{reporttype}'
                    accumulated_array_data[rtype] = interface.data.reports[reporttype]
                    accumulated_json_data[interfacename]['REPORTS'].append(rtype)
        return accumulated_json_data, accumulated_array_data, extended_data_files


class InterfaceData:
    def __init__(self, report_values : List[str], report_types : List[str] = None):
        report_types = report_types or []
        self.reports = {name : {v : [] for v in report_values}
                        for name in report_types}
        self.report_values = frozenset(report_values)
        self.errlog = []
        self.lock = Lock()
    
    def log_report(self, reporttype, **kwargs):
        with self.lock:
            if reporttype not in self.reports:
                self.reports[reporttype] = {v : [] for v in self.report_values}
            if set(kwargs.keys()) != self.report_values:
                self.errlog.append({reporttype : kwargs})
                return
            for k, v in kwargs.items():
                self.reports[reporttype][k].append(v)
    
    def _scanback(self, reporttype, report_condition):
        report = self.reports[reporttype]
        if not report:
            return None
        n = len(next(iter(report.values())))
        start_idx = n
        for i in reversed(range(n)):
            row = {
                key: values[i]
                for key, values in report.items()
            }

            if report_condition(row):
                start_idx = i
            else:
                break
        return start_idx
    
    def _scanrange(self, reporttype, report_condition):
        report = self.reports[reporttype]
        n = len(next(iter(report.values())))
        start_idx = 0
        end_idx = 0
        found = False
        for i in reversed(range(n)):
            row = {
                key: values[i]
                for key, values in report.items()
            }
            if report_condition(row):
                if not found:
                    end_idx = i + 1
                    found = True
                start_idx = i
            elif found:
                break
        return start_idx, end_idx
    
    def _scan(self, reporttype, report_condition):
        report = self.reports[reporttype]
        n = len(next(iter(report.values())))
    
        return [
            i
            for i in range(n)
            if report_condition({
                key: values[i]
                for key, values in report.items()
            })
        ]
    
    def scanback_values(self, reporttype, report_value, report_condition):
        with self.lock:
            s = self._scanback(reporttype, report_condition)
            values = self.reports[reporttype][report_value][s:]
        return values
    
    def scanback_report(self, reporttype, report_condition):
        with self.lock:
            s = self._scanback(reporttype, report_condition)
            values = {k : v[s:]
                      for k, v in self.reports[reporttype].items()}
        return values
    
    def scanrange_values(self, reporttype, report_value, report_condition):
        with self.lock:
            s, e = self._scanrange(reporttype, report_condition)
            values = self.reports[reporttype][report_value][s:e]
        return values
    
    def scanrange_report(self, reporttype, report_condition):
        with self.lock:
            s, e = self._scanrange(reporttype, report_condition)
            values = {k : v[s:e]
                      for k, v in self.reports[reporttype].items()}
        return values
    
    def scan_values(self, reporttype, report_value, report_condition):
        with self.lock:
            idxs = self._scan(reporttype, report_condition)
            report_values = self.reports[reporttype][report_value]
            values = [report_values[i] for i in idxs]
        return values
    
    def scan_report(self, reporttype, report_condition):
        with self.lock:
            idxs = self._scan(reporttype, report_condition)
            values = {
                k: [v[i] for i in idxs]
                for k, v in self.reports[reporttype].items()
            }
        return values


class AbstractInterface(ABC):
    def __init__(self, *args, **kwargs):
        self.data = None
        self.extendeddata = []
        self.setup(*args, **kwargs)
    
    @abstractmethod
    def setup(self, *args, **kwargs):
        pass
        
    @abstractmethod
    def start(self):
        pass
    
    @abstractmethod
    def stop(self):
        pass
    
    @abstractmethod
    def disconnect(self):
        pass
    
    @abstractmethod
    def dump(self):
        pass
    
    def register_extendeddatafile(self, filepath, data_type):
        self.extendeddata.append({'fp' : filepath,
                                  'type' : data_type})

class Component(ABC):
    def __init__(self, *args, **kwargs):
        self.setup(*args, **kwargs)
    
    @abstractmethod
    def setup(self, *args, **kwargs):
        pass
    
    def stop(self):
        pass
```
