## Extended Data for "A Low-Cost, Modular Hardware and Software Platform for Head-Fixed Mouse Decision-Making Tasks": banditrig.py

```
# -*- coding: utf-8 -*-
"""
Created on Wed May 14 15:02:37 2025

@author: mm4114

NOT YET IMPLEMENTED

"""

import threading
from serial.tools import list_ports
import serial
from dataclasses import dataclass, field
from collections import defaultdict
import time
from lampyr.rigs.abstract import AbstractHardwareRig, Component
from lampyr.rigs.interfaces import SerialInterface_ArduinoTRV, CameraInterface_Arducam


class Wheel(Component):
    def setup(self, serialinterface):
        self.serial = serialinterface
        self.home()

    def movement_total_since(self, time):
        components = self.movement_components_since(time)
        return components[1] - components[0]

    def movement_since(self, time):
        def condition(report): return report['unix_time'] > time
        values = self.serial.data.scanback_values(
            'R', 'report_value', condition)
        return sum(values)/4096*360

    def movement_components_since(self, time):
        def condition(report): return report['unix_time'] > time
        values = self.serial.data.scanback_values(
            'R', 'report_value', condition)
        sum_positive = sum(x for x in values if x > 0)/4096*360
        sum_negative = sum(x for x in values if x < 0)/4096*360
        return sum_negative, sum_positive

    def angle(self):
        return self.movement_since(self.home_t)

    def home(self):
        self.home_t = time.time()


class Lick(Component):
    def setup(self, serialinterface):
        self.serial = serialinterface

    def since(self, time):
        """
        Count the number of lick onset events since ``time``.

        Detects lick onsets as low-to-high threshold crossings (threshold
        512) in the raw lick signal.

        Parameters
        ----------
        time : float
            Unix timestamp; only events after this time are counted.

        Returns
        -------
        int
            Number of detected lick onsets.
        """
        def condition(report): return report['unix_time'] > time
        lickdat = self.serial.data.scanback_values(
            'L', 'report_value', condition)
        lcount = 0
        for l1, l2 in zip(lickdat, lickdat[1:]):
            if l1 < 512 and l2 > 512:
                lcount += 1
        return lcount


class Speaker(Component):
    """Speaker interface for playing tones via the rig Arduino."""

    def setup(self, serialinterface):
        self.serial = serialinterface

    def begintrialtone(self):
        """Play the trial-start tone (serial command ``'b'``)."""
        self.serial.send_command('b')

    def responsetone(self):
        """Play the response tone (serial command ``'r'``)."""
        self.serial.send_command('r')

    def punishtone(self):
        """Play the punishment tone (serial command ``'p'``)."""
        self.serial.send_command('p')


class Sipper(Component):
    """Water sipper interface for delivering liquid rewards."""

    def setup(self, serialinterface):
        self.serial = serialinterface

    def give(self):
        """Dispense one reward (serial command ``'g'``)."""
        self.serial.send_command('g')

    def setsize(self, size: int):
        """
        Set the sipper reward size on the Arduino.

        Parameters
        ----------
        size : int
            Dispense size value (arbitrary Arduino units, calibrated to
            ~5 µl per reward).
        """
        self.serial.send_command(f'w{size}')


class WheelLock(Component):
    def setup(self, serialinterface):
        self.serial = serialinterface

    def lock(self):
        self.serial.send_command('l')

    def unlock(self):
        self.serial.send_command('u')

    def to_angle(self, angle):
        self.serial.send_command(f'a{angle}')
        
    def stop(self):
        self.unlock()


class LaserControl(Component):
    def setup(self, serialinterface):
        self.serial = serialinterface

    def begin(self):
        self.serial.send_command('z')

    def stop(self):
        self.serial.send_command('x')

    def rampdown(self, ramp_ms=500):
        self.serial.send_command(f'c{ramp_ms}')


class Camera(Component):
    def setup(self, serialinterface, camerainterface):
        self.serial = serialinterface
        self.cam = camerainterface

    def begin(self):
        self.serial.send_command('o')

    def stop(self):
        self.serial.send_command('k')

    def get_framecount_inlast(self, seconds):
        cutoff = time.time() - seconds
        return len(
            self.cam.data._scanback('cam_frame_times',
                                    lambda r: r['unix_time'] > cutoff)
        )
    
    def get_total_frames(self):
        idx = 0
        with self.cam.lock:
            idx = self.cam.idx
        return idx

class BanditRig(AbstractHardwareRig):
    def setup(self):
        serialinterface = SerialInterface_ArduinoTRV(baud=115200, timeout=1)
        self.register_interface('HudaHub', serialinterface)
        self.register_component('wheel', Wheel(serialinterface))
        self.register_component('licks', Lick(serialinterface))
        self.register_component('play', Speaker(serialinterface))
        self.register_component('reward', Sipper(serialinterface))
        self.register_component('wheellock', WheelLock(serialinterface))
        self.register_component('laser', LaserControl(serialinterface))

    def initialize_mousecam(self):
        try:
            apdir = self.config._APP_DATA_DIR
        except:
            apdir = None
    
        camerainterface = CameraInterface_Arducam()
        self.register_interface('Arducam', camerainterface)
        # This will require manually starting the camera interface, as the
        # Rig wide start command has already occurred
        camerainterface.start()
        camerainterface.ready.wait(timeout = 5)
        # This is not advisable as general practice but this hack allows
        # The same rig to run camera and non-camera sessions wihout reconfig
        time.sleep(1)
        serialinterface = self.interfaces['HudaHub']
        cam = Camera(serialinterface, camerainterface)
        self.register_component('camera', cam)
        self.camera.begin()

    def is_calibrated(self):
        pass

    def is_configured(self):
        pass

    def calibrate(self):
        pass

    def configure(self):
        pass
```
