## Extended Data for "A Low-Cost, Modular Hardware and Software Platform for Head-Fixed Mouse Decision-Making Tasks": interfaces.py

```
# -*- coding: utf-8 -*-
"""
Created on Tue Jun 23 12:52:15 2026

@author: mm4114
"""

from lampyr.rigs.abstract import AbstractInterface, InterfaceData
import numpy as np
import threading
import time


class SerialInterface_ArduinoTRV(AbstractInterface):
    def setup(self, baud=115200, timeout=1):
        self.baud = baud
        self.timeout = timeout
        self._find_device()
        self.data = InterfaceData(
            ['arduino_time', 'unix_time', 'report_value'])
        self.abort_flag = False
        self.stop_flag = False
        self.thread = None

    def _find_device(self):
        from serial.tools import list_ports
        import serial
        """
        Scan COM ports for an Arduino and open the first one found.

        Raises
        ------
        RuntimeError
            If no Arduino is found in the list of available serial ports.
        """
        ports = list_ports.comports()
        ports = [p for p in ports if 'Arduino' in p.description]
        if not ports:
            raise RuntimeError(
                'No Arduino device found. Check USB connection.')
        port = ports[0]
        self.ser = serial.Serial(port.device, self.baud, timeout=self.timeout)
        time.sleep(2)
        self.ser.reset_input_buffer()
        self.ser.flush()

    def start(self):
        if self.thread is not None:
            self.stop_flag = False
            return
        self.thread = threading.Thread(target=self.listen, daemon=True)
        self.thread.start()
        time.sleep(2)
            

    def listen(self):
        """
        Background thread target: continuously read from the serial port.

        Resets the input buffer, then loops calling :meth:`_readserial` until
        the abort flag is set.  Resets the abort flag on exit.
        """
        time.sleep(1)
        self.ser.reset_input_buffer()
        self.ser.flush()
        while not self.abort_flag:
            while not self.stop_flag:
                try:
                    self._readserial()
                except Exception as error:
                    print(f'WARNING! Unknown serial read error occurred. {error}')
                time.sleep(0.001)
            time.sleep(0.001)

        self.abort_flag = False

    def _readserial(self):
        """
        Drain the serial buffer and store all valid incoming lines.

        Parses each line as ``arduino_timestamp<TAB>event_type<TAB>value``.
        Malformed lines and unicode errors are printed as warnings.  All
        successfully parsed entries are logged into the interface data object.

        Returns
        -------
        list of tuple
            Newly parsed ``(unix_time, [arduino_time, event_type, value])``
            entries from this read cycle.
        """
        responses = []
        while self.ser.in_waiting > 0:
            try:
                response = self.ser.readline().decode().strip()
            except UnicodeDecodeError as error:
                print(error)
                print(
                    'WARNING: UnicodeDecodeError detected. If this reoccurs, restart your script!!!!')
                break
            response = response.split('\t')
            # print(response)
            if len(response) == 3:
                try:
                    timestamp = int(response[0])
                    event_type = str(response[1])
                    value = int(response[2])
                    responses.append(
                        (time.time(), [timestamp, event_type, value]))
                except ValueError as e:
                    print(f"ValueError processing data: {response} - {e}")
            else:
                print(f"Received unexpected data format: {response}")
        for unix_time, (arduino_time, report_type, report_value) in responses:
            self.data.log_report(report_type, arduino_time=arduino_time,
                                 unix_time=unix_time,
                                 report_value=report_value
                                 )
        return responses

    def send_command(self, cmd):
        cmdent = f'{cmd}\n'
        self.ser.write(cmdent.encode())
    
    def stop(self):
        self.stop_flag = True

    def disconnect(self):
        self.abort_flag = True
        self.stop_flag = True
        if self.thread is not None:
            self.thread.join(timeout=2)
        self.ser.close()

    def dump(self):
        return {'COM PORT': self.ser.port,
                'BAUD': self.ser.baudrate,
                'BYTESIZE': self.ser.bytesize}

class CameraInterface_Arducam(AbstractInterface):
    def setup(self, 
              camera_index = 0,
              tempfiletarget = None,
              playback_framerate = 20,
              check_cooldown_ms = 4,
              height = 480,
              width = 640,
              data_type = 'face_cam',
              threshold_RMS = 0.001
              ):
        #settings
        self.camera_index = camera_index
        if tempfiletarget is None:
            from lampyr.primatives import uniqueid
            import tempfile
            import os
            temp_dir = tempfile.gettempdir()
            uniqueid = uniqueid('ArducamVideo', data_type)
            self.tempfiletarget = os.path.join(temp_dir, f"{uniqueid}.avi")
        else:
            self.tempfiletarget = tempfiletarget
        self.fps = playback_framerate
        self.height = height
        self.width = width
        self.data_type = data_type
        self.threshold_RMS = threshold_RMS
        
        self.data = InterfaceData(report_values = ['unix_time','frame_num'],
                                  report_types = ['cam_frame_times'])
        
        # Threading/Runtime
        self.abort_flag = False
        self.thread = None
        self.idx = 0
        self.check_cooldown_ms = check_cooldown_ms
        self.ready = threading.Event()
        
        # set during startup
        self.cap = None
        self.out = None
    
    def start(self):
        import os
        os.environ["OPENCV_LOG_LEVEL"] = "ERROR"
        import cv2
        self.cap = cv2.VideoCapture(self.camera_index, cv2.CAP_MSMF)
        if not self.cap.isOpened():
            self.cap.release()
            raise RuntimeError("Could not open Camera.")
        self.cap.set(cv2.CAP_PROP_FRAME_WIDTH, self.width)
        self.cap.set(cv2.CAP_PROP_FRAME_HEIGHT, self.height)
        
        fourcc = cv2.VideoWriter_fourcc(*"XVID")
        self.out = cv2.VideoWriter(self.tempfiletarget,
                                   fourcc, self.fps,
                                   (self.width, self.height),
                                   isColor=False)
        if not self.out.isOpened():
            self.cap.release()
            raise RuntimeError("Could not open VideoWriter.")
        
        self.register_extendeddatafile(self.tempfiletarget,
                                       self.data_type)
        
        self.thread = threading.Thread(target=self._listen,
                                       daemon=True)
        self.thread.start()
    
    def _listen(self):
        import cv2
        self.ready.set()
        last_accepted_frame = None
        while not self.abort_flag:
            readtime = time.time()
            ret, frame = self.cap.read()
            if not ret or frame is None:
                time.sleep(self.check_cooldown_ms/1000)
                continue
            
            #Force Greyscale
            if len(frame.shape) == 3:
                frame = cv2.cvtColor(frame, cv2.COLOR_BGR2GRAY)
            # Force correct size for VideoWriter if driver returns a different shape.
            if frame.shape[1] != self.width or frame.shape[0] != self.height:
                frame = cv2.resize(frame, (self.width, self.height))
            
            # Filter frame out if not reached RMS threshold
            if last_accepted_frame is not None:
                diff = frame.astype(np.float32) - last_accepted_frame.astype(np.float32)
                rms = np.sqrt(np.mean(diff * diff))
    
                if rms < self.threshold_RMS:
                    continue
            # ABOVE IS IMPORTANT WITH MSMF BACKEND!!!!
            # MSMF will continue to supply duplicate frames even when a new
            # frame has not been grabbed.
            
            last_accepted_frame = frame.copy()
            self.out.write(frame)
            self.data.log_report('cam_frame_times', 
                                 unix_time = readtime,
                                 frame_num = self.idx)
            self.idx+=1
            time.sleep(self.check_cooldown_ms/1000)
    
    def stop(self):
        self.abort_flag = True
        if self.thread is not None:
            self.thread.join(timeout=2)
        if self.out is not None:
            self.out.release()
        if self.cap is not None:
            self.cap.release()
        self.thread = None
        self.out = None
        self.cap = None
        self.ready.clear()
    
    def disconnect(self):
        pass
    
    def dump(self):
        return {'DIMENSIONS' : (self.width, self.height),
                'FRAMES_CAPTURED' : self.idx,
                'tempfiletarget' : self.tempfiletarget,
                'playback_framerate' : self.fps}


if __name__ == '__main__':
    interface = CameraInterface_Arducam()
    interface.start()
    time.sleep(2)
    print('gogogo')
    time.sleep(10)
    interface.stop()
    interface.disconnect()
    print(interface.data.reports)
```
