## Extended Data for "A Low-Cost, Modular Hardware and Software Platform for Head-Fixed Mouse Decision-Making Tasks": rigcontrol2.py

```
# -*- coding: utf-8 -*-
"""
Created on Wed May 14 15:02:37 2025

@author: mm4114
"""

import threading
from serial.tools import list_ports
import serial
from dataclasses import dataclass, field
from collections import defaultdict
import time

class ArduinoBanditRig_0:
    """
    High-level interface to the Arduino-based bandit rig (hardware version 0).

    Wraps a :class:`SerialMonitor` and exposes named sub-objects for each
    hardware component: wheel encoder, lick sensor, speaker, and sipper.

    Attributes
    ----------
    serial : SerialMonitor
        Serial communication handler.
    data : SerialData
        Shared data store populated by the serial monitor thread.
    wheel : Wheel
        Wheel encoder interface.
    licks : Lick
        Lick sensor interface.
    play : Speaker
        Speaker/tone interface.
    reward : Sipper
        Water sipper interface.
    """

    class _Measure():
        """Base class for rig components that read from ``SerialData``."""

        def __init__(self, parent):
            self.data = parent.data

    class _Control():
        """Base class for rig components that write to the serial port."""

        def __init__(self, parent):
            self.serial = parent.serial

    class Wheel(_Measure):
        """
        Rotary wheel encoder interface.

        Reads raw encoder ticks from report type ``'R'`` and converts to
        degrees (4096 ticks per revolution).
        """

        def __init__(self, *args, **kwargs):
            super().__init__(*args, **kwargs)
            self.home()

        def movement_total_since(self, time):
            """
            Return the net signed displacement (degrees) since ``time``.

            Equivalent to ``positive_component - negative_component``.

            Parameters
            ----------
            time : float
                Unix timestamp; only events after this time are counted.

            Returns
            -------
            float
                Net displacement in degrees.
            """
            components = self.movement_components_since(time)
            return components[1] - components[0]

        def movement_since(self, time):
            """
            Return the sum of all encoder ticks (signed) since ``time``, in degrees.

            Parameters
            ----------
            time : float
                Unix timestamp; only events after this time are counted.

            Returns
            -------
            float
                Signed total displacement in degrees.
            """
            return sum(self.data.get_reportvals_since('R', time))/4096*360

        def movement_components_since(self, time):
            """
            Return negative and positive movement components separately.

            Parameters
            ----------
            time : float
                Unix timestamp; only events after this time are counted.

            Returns
            -------
            tuple of (float, float)
                ``(negative_degrees, positive_degrees)`` — both values are
                expressed as non-negative magnitudes for the respective
                directions.
            """
            sum_positive = sum(x for x in self.data.get_reportvals_since('R', time)
                               if x > 0)/4096*360
            sum_negative = sum(x for x in self.data.get_reportvals_since('R', time)
                               if x < 0)/4096*360
            return sum_negative, sum_positive

        def angle(self):
            """
            Return the current wheel angle relative to the last :meth:`home` call.

            Returns
            -------
            float
                Signed displacement in degrees from the home position.
            """
            return self.movement_since(self.home_t)

        def home(self):
            """
            Reset the wheel home position to the current time.

            Subsequent calls to :meth:`angle` will measure displacement from
            this moment.
            """
            self.home_t = time.time()

    class Lick(_Measure):
        """Lick detector interface reading report type ``'L'``."""

        def since(self, time):
            """
            Count the number of lick onset events since ``time``.

            Detects lick onsets as low-to-high threshold crossings (threshold
            512) in the raw lick signal.

            Parameters
            ----------
            time : float
                Unix timestamp; only events after this time are counted.

            Returns
            -------
            int
                Number of detected lick onsets.
            """
            lickdat = self.data.get_reportvals_since('L', time)
            lcount = 0
            for l1, l2 in zip(lickdat,lickdat[1:]):
                if l1 < 512 and l2 > 512:
                    lcount += 1
            return lcount

    class Speaker(_Control):
        """Speaker interface for playing tones via the rig Arduino."""

        def begintrialtone(self):
            """Play the trial-start tone (serial command ``'b'``)."""
            self.serial._writeserial('b')

        def responsetone(self):
            """Play the response tone (serial command ``'r'``)."""
            self.serial._writeserial('r')

        def punishtone(self):
            """Play the punishment tone (serial command ``'p'``)."""
            self.serial._writeserial('p')

    class Sipper(_Control):
        """Water sipper interface for delivering liquid rewards."""

        def give(self):
            """Dispense one reward (serial command ``'g'``)."""
            self.serial._writeserial('g')

        def setsize(self, size : int):
            """
            Set the sipper reward size on the Arduino.

            Parameters
            ----------
            size : int
                Dispense size value (arbitrary Arduino units, calibrated to
                ~5 µl per reward).
            """
            self.serial._writeserial('w')
            self.serial._writeserial(f'{size}\n')

    def __init__(self, customserialmonitor=None):
        """
        Initialise the rig, creating a :class:`SerialMonitor` if none is provided.

        Parameters
        ----------
        customserialmonitor : SerialMonitor, optional
            Use a pre-constructed serial monitor (useful for testing).  If
            ``None``, a new :class:`SerialMonitor` at 115200 baud is created.
        """
        if customserialmonitor is None:
            self.serial = SerialMonitor(115200)
        else:
            self.serial =  customserialmonitor
        self.data = self.serial.data
        self.wheel = self.Wheel(self)
        self.licks = self.Lick(self)
        self.play = self.Speaker(self)
        self.reward = self.Sipper(self)

    def listen(self):
        """
        Start the serial monitor thread and purge any buffered data.

        Blocks for 1 second to allow the Arduino to reset before purging.
        """
        self.serial.listen()
        time.sleep(1)
        self.serial.purge()

    def abort(self):
        """Signal the serial monitor thread to stop."""
        self.serial.abort()

    def close(self):
        """Close the underlying serial port."""
        self.serial.close()

# Helper function to create the default dictionary for reports.
def _create_report_dict():
    """Returns the default dictionary structure for a new report type."""
    return {'unix_time': [], 'arduino_time': [], 'report_value': []}

@dataclass
class SerialData:
    """
    Thread-safe store for all data received from the Arduino.

    Attributes
    ----------
    lock : threading.Lock
        Protects concurrent access from the reader thread and user code.
    log : list of tuple
        Raw log entries: ``(unix_time, [arduino_time, event_type, value])``.
    reports : defaultdict of dict
        Per-report-type time series.  Each value is a dict with keys
        ``'unix_time'``, ``'arduino_time'``, and ``'report_value'`` (lists).
    """
    lock: threading.Lock = field(default_factory=lambda: threading.Lock())
    log: list = field(default_factory=lambda: [])
    reports: dict = field(default_factory=lambda: defaultdict(_create_report_dict))

    def get_reportvals_since(self, reporttype, unixtime):
        """
        Return all report values for ``reporttype`` recorded at or after ``unixtime``.

        Parameters
        ----------
        reporttype : str
            Report channel identifier (e.g. ``'R'`` for wheel, ``'L'`` for lick).
        unixtime : float
            Cutoff Unix timestamp (inclusive).

        Returns
        -------
        list
            Slice of the ``'report_value'`` list starting at the first entry
            whose ``unix_time`` >= ``unixtime``.  Returns an empty list if no
            matching entries exist.
        """
        with self.lock:
            reports = self.reports[reporttype]
            times = reports['unix_time']
            for i, t in enumerate(times):
                if t >= unixtime:
                    return reports['report_value'][i:]
        return []

    def get_report_snippet(self, start_unixtime, end_unixtime):
        """
        Extract a time-bounded snippet of all report data.

        Parameters
        ----------
        start_unixtime : float
            Start of the time window (inclusive).
        end_unixtime : float
            End of the time window (inclusive).

        Returns
        -------
        dict
            Nested dict ``{reporttype: {'unix_time': [...], 'arduino_time': [...], 'report_value': [...]}}``
            containing only entries within ``[start_unixtime, end_unixtime]``.
        """
        result = {}
        for reporttype, data in self.reports.items():
            result[reporttype] = {'unix_time': [],
                                  'arduino_time': [], 'report_value': []}
            for tu, ta, v in zip(data['unix_time'], data['arduino_time'], data['report_value']):
                if start_unixtime <= tu <= end_unixtime:
                    result[reporttype]['unix_time'].append(tu)
                    result[reporttype]['arduino_time'].append(ta)
                    result[reporttype]['report_value'].append(v)
        return result


class SerialMonitor:
    """
    Manages serial communication with the Arduino in a background thread.

    Finds the first connected Arduino, opens a serial port, and continuously
    reads incoming tab-separated ``timestamp\tevent_type\tvalue`` lines,
    storing parsed data in a :class:`SerialData` instance.
    """

    def __init__(self, baud, timeout=1):
        """
        Locate and open the Arduino serial port.

        Parameters
        ----------
        baud : int
            Baud rate for the serial connection.
        timeout : int, optional
            Serial read timeout in seconds. Default is 1.

        Raises
        ------
        RuntimeError
            If no Arduino device is detected on any available COM port.
        """
        self.baud = baud
        self.timeout = timeout
        self.find_device()
        self.data = SerialData()
        self.abort_flag = False
        self.threadlock = threading.Lock()

    def abort(self):
        """Set the abort flag, causing the background listener thread to exit."""
        self.abort_flag = True

    def close(self):
        """Close the serial port."""
        self.ser.close()

    def find_device(self):
        """
        Scan COM ports for an Arduino and open the first one found.

        Raises
        ------
        RuntimeError
            If no Arduino is found in the list of available serial ports.
        """
        ports = list_ports.comports()
        ports = [p for p in ports if 'Arduino' in p.description]
        if not ports:
            raise RuntimeError('No Arduino device found. Check USB connection.')
        port = ports[0]
        self.ser = serial.Serial(port.device, self.baud, timeout=self.timeout)
        time.sleep(2)
        self.ser.reset_input_buffer()
        self.ser.flush()

    def listen(self):
        """
        Start the background serial reader thread and wait 2 seconds for it to stabilise.
        """
        thread = threading.Thread(target=self._listen, daemon=True)
        thread.start()
        time.sleep(2)

    def purge(self):
        """
        Clear all accumulated data from the :class:`SerialData` store.

        Resets ``data.log`` and ``data.reports`` under the data lock so the
        experiment starts with a clean buffer.
        """
        with self.data.lock:
            self.data.log = []
            self.data.reports = defaultdict(_create_report_dict)

    def _listen(self):
        """
        Background thread target: continuously read from the serial port.

        Resets the input buffer, then loops calling :meth:`_readserial` until
        the abort flag is set.  Resets the abort flag on exit.
        """
        time.sleep(1)
        self.ser.reset_input_buffer()
        self.ser.flush()
        while not self.abort_flag:
            try:
                self._readserial()
            except Exception as error:
                print(f'WARNING! Unknown serial read error occurred. {error}')
            time.sleep(0.001)

        self.abort_flag = False

    def _readserial(self):
        """
        Drain the serial buffer and store all valid incoming lines.

        Parses each line as ``arduino_timestamp<TAB>event_type<TAB>value``.
        Malformed lines and unicode errors are printed as warnings.  All
        successfully parsed entries are appended to ``data.log`` and
        ``data.reports``.

        Returns
        -------
        list of tuple
            Newly parsed ``(unix_time, [arduino_time, event_type, value])``
            entries from this read cycle.
        """
        responses = []
        while self.ser.in_waiting > 0:
            try:
                response = self.ser.readline().decode().strip()
            except UnicodeDecodeError as error:
                print(error)
                print('WARNING: UnicodeDecodeError detected. If this reoccurs, restart your script!!!!')
                break
            response = response.split('\t')
            # print(response)
            if len(response) == 3:
                try:
                    timestamp = int(response[0])
                    event_type = str(response[1])
                    value = int(response[2])
                    responses.append(
                        (time.time(), [timestamp, event_type, value]))
                except ValueError as e:
                    print(f"ValueError processing data: {response} - {e}")
            else:
                print(f"Received unexpected data format: {response}")
        self.data.log += responses
        for unix_time, (arduino_time, report_type, report_value) in responses:
            self.data.reports[report_type]['unix_time'].append(unix_time)
            self.data.reports[report_type]['arduino_time'].append(arduino_time)
            self.data.reports[report_type]['report_value'].append(report_value)
        return responses

    def _writeserial(self, cmd):
        """
        Encode and send a command string to the Arduino.

        Parameters
        ----------
        cmd : str
            Command character or string to send.
        """
        self.ser.write(cmd.encode())
```
