## Extended Data for "A Low-Cost, Modular Hardware and Software Platform for Head-Fixed Mouse Decision-Making Tasks": abstract.py

```
# -*- coding: utf-8 -*-
"""
Created on Thu Jun 26 17:21:05 2025

@author: mm4114
"""

from dataclasses import dataclass, field
import time
from copy import deepcopy
from typing import Callable, List, Literal
from abc import ABC, abstractmethod
import os

from lampyr.primatives import Mouse, Session, uniqueid


@dataclass
class Segment(ABC):
    """
    Abstract base class for all Lampyr segment types.

    A segment represents a discrete, timed unit of experimental behaviour
    (trial, task, stage, paradigm, etc.).  Subclasses implement
    :meth:`execute` to define what happens when the segment runs.

    Segments form a tree: each segment knows its ``parent`` and accumulates
    ``subdata`` references.  Property inheritance, logging, and session I/O
    are all handled here.

    Attributes
    ----------
    name : str or None
        Human-readable name, auto-generated from parent and slug if not set.
    slug : str or None
        Short class-level label; defaults to the class name.
    uniqueid : str or None
        Globally unique identifier generated at initialisation.
    starttime : float or None
        Unix timestamp recorded at the start of :meth:`run`.
    endtime : float or None
        Unix timestamp recorded after :meth:`execute` returns.
    rank : int
        Depth in the segment tree; root segments have rank 0.
    subdata : list of str
        ``uniqueid`` strings of child segments appended during execution.
    lampyr : object or None
        Parent Lampyr instance; ``None`` when running standalone.
    parent : Segment or None
        Immediate parent segment; ``None`` for root segments.
    records : list of tuple
        General-purpose log: ``(unix_time, prefix, message)`` entries.
    rig : object or None
        Hardware rig object, inherited from parent or lampyr.
    mouse : Mouse or None
        Active mouse, inherited from parent or lampyr.
    session : Session or None
        Active session, inherited from parent or lampyr.
    _output_func : callable or None
        Output function used by the logging methods.
    _verbose : bool
        If ``True``, DEBUG messages are also written to output.
    _frozen : bool
        Set to ``True`` after :meth:`run` completes; prevents re-running.
    """

    # General identification
    name: str = None
    slug: str = None
    uniqueid: str = None
    starttime: float = None
    endtime: float = None

    # References to other segments and lampyr
    rank: int = 0
    subdata: list = field(default_factory=list)
    lampyr: object = None
    parent: object = None

    # Data and logs
    records: List = field(default_factory=list)  # General purpose log

    # Mouse, rig, session data
    rig: object = None
    mouse: Mouse = None
    session: Session = None

    # Input/Output functions
    _output_func: Callable = None  # inherited from lampyr; falls back to print if standalone
    _verbose: bool = False

    # Utilities
    _parent_inheritproperties: list = field(default_factory=list)
    _lampyr_inheritproperties: list = field(default_factory=list)
    _parent_inheritproperties_combine: list = field(default_factory=list)
    _dump_reducetorepresentations: list = field(default_factory=list)
    _dump_exclusions: list = field(default_factory=list)
    _frozen: bool = False
    
    # Logging nonsense
    _last_log_notice_t: float = field(default=0, init=False)
    _last_log_notice: str = field(default='', init=False)
    _last_log_info_t: float = field(default=0, init=False)
    _last_log_info: str = field(default='', init=False)

    def __post_init__(self):
        """
        Finalise segment initialisation after dataclass ``__init__``.

        Sets default ``slug``, ``name``, and ``uniqueid``; inherits properties
        from the parent segment and lampyr instance; creates a fallback
        :class:`~lampyr.primatives.Session` if none was found; and marks root
        segments (rank 0).
        """
        self._configure()
        if self.slug is None:
            self.slug = self.__class__.__name__
        if self.name is None:
            if self.parent is not None:
                self.name = f'{self.parent.name}_{self.slug}{len(self.parent.subdata)}'
            else:
                self.name = self.slug
        if self.uniqueid is None:
            self.uniqueid = uniqueid(
                'segment', self.__class__.__name__)
        self._inherit()
        self._inherit_from_lampyr()
        if self.session is None:
            self.log_warning('Not able to find a session. Creating new session (destructive).')
            self.session = Session()
        if self.parent is None:
            self.rank = 0
            self.log_debug('No parent detected, marking this segment as root')
            self.session.root = self.uniqueid

    @classmethod
    def get_children(cls, recursive=True):
        """
        Return all subclasses of this segment class.

        Parameters
        ----------
        recursive : bool, optional
            If ``True`` (default), include indirect subclasses at all depths.

        Returns
        -------
        set of type
            Set of subclass objects.
        """
        children = set()
        for subclass in cls.__subclasses__():
            children.add(subclass)
            if recursive:
                children.update(subclass.get_children())
        return children
    
    @classmethod
    def get_parents(cls, recursive=True):
        """
        Return all :class:`Segment` ancestor classes of this class.

        Parameters
        ----------
        recursive : bool, optional
            If ``True`` (default), walk the full MRO and include grandparents.

        Returns
        -------
        set of type
            Set of ancestor Segment subclasses (excludes non-Segment bases).
        """
        parents = set()
        for parent in cls.__bases__:
            if not issubclass(parent, Segment):
                continue
            parents.add(parent)
            if recursive and hasattr(parent, 'get_parents'):
                parents.update(parent.get_parents())
        return parents
    
    def run(self):
        """
        Execute the segment, recording start/end times and serialising data.

        Calls :meth:`execute`, then :meth:`dump` in a ``finally`` block so
        data is always persisted even if execution raises.

        Raises
        ------
        RuntimeError
            If called on a segment that has already been run (``_frozen``).
        KeyboardInterrupt
            Re-raised after logging, so the caller can handle forced quits.
        """
        if self._frozen:
            raise RuntimeError('Running a segment twice is forbidden')
        self.starttime = time.time()
        try:
            self.execute()
        except KeyboardInterrupt as error:
            self.log_error(f'User has initiated a force-quit. {self.name} closing...')
            raise
        finally:
            self.endtime = time.time()
            self._frozen = True
            self.dump()    
    
    @abstractmethod
    def execute(self):
        """
        Define the behaviour for this segment.

        Called by :meth:`run`.  Subclasses must implement this method with
        whatever logic the segment type requires.
        """
        pass

    def _log(self, prefix, message: str, output=True, style='\x1b[33m'):
        """
        Internal log dispatcher: write to output and append to records.

        Parameters
        ----------
        prefix : str
            Log level label (e.g. ``'INFO'``, ``'ERROR'``).
        message : str
            Text to log.
        output : bool, optional
            If ``True``, pass the formatted string to ``_output_func``.
        style : str, optional
            ANSI escape code prefix for terminal colouring.

        Returns
        -------
        float
            Unix timestamp of the log event.
        """
        if output:
            out = self._output_func if self._output_func is not None else print
            out(f'{style}[{prefix}][{self.name}] {message}\033[0m')
        self.records.append((time.time(), prefix, message))
        return time.time()

    def log_info(self, message: str, delay = None):
        """
        Log an informational message (cyan).

        Parameters
        ----------
        message : str
            Message to log.
        delay : float, optional
            Minimum seconds between consecutive emissions of the same message.
            If the same message was emitted within ``delay`` seconds, it is
            suppressed.

        Returns
        -------
        float
            Unix timestamp of the log event.
        """
        if delay is None:
            return self._log('INFO', message, output=True)
        if message != self._last_log_info or self._last_log_info_t + delay < time.time():
            self._last_log_info = message
            self._last_log_info_t = time.time()
            return self._log('INFO', message, output=True)
        return time.time()

    def log_debug(self, message: str):
        """
        Log a debug message (dark grey). Only shown when ``_verbose`` is True.

        Parameters
        ----------
        message : str
            Message to log.

        Returns
        -------
        float
            Unix timestamp of the log event.
        """
        return self._log('DEBUG', message, output=self._verbose, style='\x1b[90m')
    
    def log_notice(self, message: str):
        """
        Log a notice message (yellow).

        Parameters
        ----------
        message : str
            Message to log.
        delay : float, optional
            Minimum seconds between consecutive emissions of the same message.

        Returns
        -------
        float
            Unix timestamp of the log event.
        """
        return self._log('NOTICE', message, output=True, style='\033[93m')
    
    def log_warning(self, message: str):
        """
        Log a warning message (orange).

        Parameters
        ----------
        message : str
            Message to log.

        Returns
        -------
        float
            Unix timestamp of the log event.
        """
        return self._log('WARNING', message, output=True, style='\033[38;5;208m')

    def log_error(self, message: str):
        """
        Log an error message (red).

        Parameters
        ----------
        message : str
            Message to log.

        Returns
        -------
        float
            Unix timestamp of the log event.
        """
        return self._log('ERROR', message, output=True, style='\033[38;5;196m')
    
    def notify(self, message):
        """
        Notify relevant users via lampyr.notificationmanager

        Parameters
        ----------
        message : str
            Message to send to user.

        Returns
        -------
        None.

        """
        self.log_notice(message)
        if self.lampyr is None:
            self.log_warning('User push notification is not possible without lampyr instance')
            return
        self.lampyr.notify(message)

    def dump(self):
        """
        Serialise segment state into the session and, if root, finalise it.

        Collects all public attributes (excluding private, excluded, and
        reduced fields), replaces complex references with their ``uniqueid``
        or ``repr``, and stores the result in ``session.segments``.  Root
        segments (rank 0) additionally extract rig data and lock the session.

        Returns
        -------
        dict
            The serialised segment data dict stored in the session.
        """
        all_data = {k: v for k, v in self.__dict__.items()
                    if k not in self._dump_reducetorepresentations
                    and k not in self._dump_exclusions
                    and not k.startswith('_')}
        for k in self._dump_reducetorepresentations:
            if k not in self.__dict__:
                continue
            v = self.__dict__[k]
            if hasattr(v, 'uniqueid'):
                all_data[k] = v.uniqueid
            elif v is None:
                all_data[k] = None
            else:
                all_data[k] = v.__repr__()
        all_data['segment_type'] = [cls.__name__ for cls in self.get_parents()]
        try:
            all_data = deepcopy(all_data)
        except:
            print([[k, type(v)] for k, v in all_data.items()])
        self.log_debug('Storing data in session...')
        self.session.segmentlist.append(self.uniqueid)
        self.session.segments[self.uniqueid] = all_data
        if self.rank == 0:
            self.log_notice('Detected that self is highest ranked segment')
            self.log_notice('Stopping data aquisition')
            self.rig.stop()
            self.log_debug('Allowing grace period for rig data to finish accumulating')
            time.sleep(0.5)
            if self.lampyr is None:
                self.log_warning("Detected no lampyr instance, attempting to fully close rig")
                self.rig.disconnect()
            self.log_notice('Extracting rig data and saving to session')
            rig_json_representation, rig_data_numerical, rig_extended_data = self.rig.dump()
            self.session.rigproperties = rig_json_representation
            self.session.rigdata = rig_data_numerical
            self.session._extendeddata = rig_extended_data
            self.session.extendeddata = \
                [os.path.join(e['type'],
                              os.path.basename(e['fp'])
                              )
                 for e in rig_extended_data]
            
            self.log_notice('Session data is now LOCKED')
            self.session.lock()
            
            
        else:
            self.log_debug('Storing reference to self in parent segment')
            self.parent.subdata.append(self.uniqueid)
        return all_data

    def _configure(self):
        """
        Populate default inheritance and dump control lists.

        Called during ``__post_init__`` before any inheritance takes place.
        Subclasses should call ``super()._configure()`` and then extend the
        relevant lists.
        """
        self._parent_inheritproperties += ['lampyr',
                                           'rig', 'mouse', 'session', '_output_func',
                                           '_verbose']
        self._lampyr_inheritproperties += ['rig', 'mouse', 'session', '_output_func']
        self._dump_reducetorepresentations += ['lampyr', 'parent']
        self._dump_exclusions += ['rig', 'session', 'mouse']

    def _inherit(self):
        """
        Pull listed properties from the parent segment.

        Sets ``rank`` to one greater than the parent's rank, then copies
        each property in ``_parent_inheritproperties`` (destructive replace)
        and combines those in ``_parent_inheritproperties_combine``.
        """
        if self.parent is None:
            return
        self.rank = self.parent.rank + 1
        for prop in self._parent_inheritproperties:
            self.log_debug(f'Inheriting {prop} from parent (destructive)')
            self._inheritproperty(self.parent, prop, 'replace')
        for prop in self._parent_inheritproperties_combine:
            self.log_debug(f'Inheriting {prop} from parent (combinatorial)')
            self._inheritproperty(self.parent, prop, 'combine')
    
    def _inherit_from_lampyr(self):
        """
        Fill any ``None`` properties by pulling them from the lampyr instance.

        Only copies a property from lampyr if the segment's own value is
        currently ``None``, preserving any value already set by parent
        inheritance.
        """
        if self.lampyr is None:
            self.log_warning('You are running segments without a lampyr instance.')
            return
        for prop in self._lampyr_inheritproperties:
            if getattr(self, prop) is None:
                self.log_debug(f'Inheriting {prop} from parent (destructive)')
                self._inheritproperty(self.lampyr, prop, 'replace')
        

    def _inheritproperty(self, source, name: str, mode: Literal['replace', 'combine']):
        """
        Copy or merge a single property from ``source`` into this segment.

        Parameters
        ----------
        source : object
            The object to copy from (a parent segment or lampyr instance).
        name : str
            Attribute name to inherit.
        mode : {'replace', 'combine'}
            ``'replace'`` overwrites the local value; ``'combine'`` merges
            lists (concatenation) or dicts (update) with the local value.

        Raises
        ------
        TypeError
            If ``mode='combine'`` but the source and local types are
            incompatible (e.g. list vs dict).
        """
        if not hasattr(source, name):
            return
        attr = getattr(source, name)
        if attr is None:
            return
        if mode == 'replace':
            setattr(self, name, attr)
            return
        myattr = getattr(self, name)
        hint = getattr(type(self), '__annotations__', {}).get(name, None)
        if myattr is None:
            setattr(self, name, attr)
            return
        if isinstance(attr, list) and isinstance(myattr, list):
            combined = attr + myattr
        elif isinstance(attr, dict) and isinstance(myattr, dict):
            combined = {**attr, **myattr}
        else:
            raise TypeError(f'{self.name} has attempted to inherit {name} from parent and has experienced a type mismatch.' +
                            f'Parent type was {type(attr)}, object type was {type(myattr)} and object annotation was for {hint}')
        setattr(self, name, combined)
```
