## Extended Data for "A Low-Cost, Modular Hardware and Software Platform for Head-Fixed Mouse Decision-Making Tasks": behavior.py

```
# -*- coding: utf-8 -*-
"""
Created on Thu Jun 26 17:21:59 2025

@author: mm4114
"""
from dataclasses import dataclass, field
import time
from copy import deepcopy
from typing import Callable, List, Literal
from abc import ABC, abstractmethod

from lampyr.primatives import Mouse, Session, uniqueid
from lampyr.segments.abstract import Segment


"""
REFACTOR NEEDED

The plan is for BehaviorSegment to implement all methods and event registration, with Task and Trial
as subclasses that provide structure for different types of behavior.  Task is for "trialless" behaviors or those
that include multiple trials, while trial is for discrete trial-based behaviors.
"""

@dataclass
class BehaviorSegment(Segment):
    # Tags allow for tag based mouse overrides and are inherited and combined across the segment hierarchy.
    tags: List[str] = field(default_factory=list)  # Inherited from parents
    # Describe the results of the segment
    reports: dict = field(default_factory=dict)
    # describe discrete events and timepoints of interest during the segment
    event_definitions: dict = field(default_factory=dict)
    event_records: list = field(default_factory=list)
    _events: dict = field(default_factory=dict)

    # Reasons for stopping the session
    stop_reasons: list = field(default_factory=list)

    def __post_init__(self):
        """
        Extend base ``__post_init__`` to apply any per-mouse parameter overrides.
        """
        super().__post_init__()
        self._checkoverrides()
        

    def _configure(self):
        """
        Extend base ``_configure`` to set up inheritance and dumping for behavior-specific attributes.
        """
        super()._configure()
        self._parent_inheritproperties_combine += ['tags', 'event_definitions']
        self._dump_reducetorepresentations += ['_events']

    def _applyoverrides(self, overrides: dict):
        """
        Apply a dict of ``{param: value}`` overrides to this segment's attributes.

        Parameters
        ----------
        overrides : dict
            Mapping of attribute names to override values.  Only attributes
            that already exist on the segment are updated.
        """
        for param, val in overrides.items():
            if hasattr(self, param):
                setattr(self, param, val)

    def _checkoverrides(self):
        """
        Look up and apply mouse-specific parameter overrides for this segment.

        Checks the mouse's ``mouse_behav_param_overrides`` dict for entries
        keyed by ``'all'``, the segment's ``slug``, and each of its ``tags``,
        applying any matches via :meth:`_applyoverrides`.
        """
        if self.mouse is None:
            return
        if 'all' in self.mouse.mouse_behav_param_overrides:
            overrides = self.mouse.mouse_behav_param_overrides['all']
            self.log_debug(f'Applying {len(overrides)} overrides')
            self._applyoverrides(overrides)
        if self.slug in self.mouse.mouse_behav_param_overrides:
            overrides = self.mouse.mouse_behav_param_overrides[self.slug]
            self.log_debug(f'Applying {len(overrides)} overrides associated with ' +
                          f'{self.slug} slug')
            self._applyoverrides(overrides)
        for tag in self.tags:
            if tag in self.mouse.mouse_behav_param_overrides:
                overrides = self.mouse.mouse_behav_param_overrides[tag]
                self.log_debug(f'Applying {len(overrides)} overrides associated with ' +
                              f'{tag} tag')
                self._applyoverrides(overrides)

    def finish(self):
        """
        Signal that this segment should stop on the next loop evaluation.

        Appends ``'finished'`` to ``stop_reasons``, which causes
        :meth:`execute` to exit the loop on its next iteration.
        """
        self.stop_reasons.append('finished')
    
    def log_merit(self, increment=1):
        """
        Increment the session merit counter and log participation.

        Parameters
        ----------
        increment : int, optional
            Amount to add to ``session.merit``. Default is 1.
        """
        if self.session:
            self.session.merit += increment
            self.log_participation(increment)
            self.log_debug(f'Merit +{increment} → {self.session.merit}.')

    def log_demerit(self, increment=1):
        """
        Increment the session demerit counter and log participation.

        Parameters
        ----------
        increment : int, optional
            Amount to add to ``session.demerit``. Default is 1.
        """
        if self.session:
            self.session.demerit += increment
            self.log_participation(increment)
            self.log_debug(f'Demerit +{increment} → {self.session.demerit}.')

    def log_abstention(self, increment=1):
        """
        Increment the abstention and serial abstention counters.

        Parameters
        ----------
        increment : int, optional
            Amount to add. Default is 1.
        """
        if self.session:
            self.session.abstention += increment
            self.session.serial_abstention += increment
            self.log_debug(
                f'Abstention +{increment} → {self.session.abstention}.')
            self.log_debug(
                f'SerialAbstention +{increment} → {self.session.serial_abstention}.')

    def log_participation(self, increment=1):
        """
        Increment the participation counter and reset serial abstention to 0.

        Parameters
        ----------
        increment : int, optional
            Amount to add to ``session.participation``. Default is 1.
        """
        if self.session:
            self.session.participation += increment
            self.session.serial_abstention = 0
            self.log_debug(
                f'Participation +{increment} → {self.session.participation}. SerialAbstention reset.')

    def log_reward(self, amount):
        """
        Increment the session reward counter.

        Parameters
        ----------
        increment : int, optional
            Amount to add to ``session.rewards``. Default is 1.
        """
        if self.session:
            self.session.rewards += 1
            self.session.reward_amount += amount
            self.log_debug(f'Reward +1 → {self.session.rewards}.')
            self.log_debug(f'Reward +{amount} → {self.session.reward_amount}.')

    def log_trial(self, increment=1):
        """
        Increment the session trial counter.

        Parameters
        ----------
        increment : int, optional
            Amount to add to ``session.trial``. Default is 1.
        """
        if self.session:
            self.session.trial += increment
            self.log_debug(f'Trial +{increment} → {self.session.trial}.')

    def create_report(self, key, value):
        """
        Store a key/value result in this segment's ``reports`` dict.

        Parameters
        ----------
        key : str
            Report field name.
        value : object
            Value to store.  Overwriting an existing key emits a warning.
        """
        if key in self.reports:
            self.log_warning(f'Overwrote {key} as {value}. This is not recommended behavior.')
        self.reports[key] = value
    
    def register_event(self, name: str,
                       callback: Callable = lambda *args, **kwargs: None,
                       description: str = None):
        """
        Register a named event with an optional callback.

        Parameters
        ----------
        name : str
            Unique event name.  Logs debug statement if already registered.
        callback : callable, optional
            Function called when the event is triggered.  Receives the trial
            segment as the first argument, followed by any ``*args``/``**kwargs``
            passed to :meth:`trigger_event`.  Defaults to a no-op.
        description : str, optional
            Human-readable description stored in ``event_definitions``.

        Raises
        ------
        RuntimeError
            If ``name`` is already registered on this trial.
        """
        if name in self._events:
            self.log_debug('Did not create event {name} because it already exists.')
        self._events[name] = callback
        self.event_definitions[name] = {'description': description,
                                        'callback': callback.__repr__()}

    def trigger_event(self, event, *args, **kwargs):
        """
        Fire a registered event, invoking its callback and recording the occurrence.

        Appends an entry to both ``event_records`` and ``session.eventlist``.

        Parameters
        ----------
        event : str
            Name of the event to trigger; must have been registered via
            :meth:`register_event`.
        *args
            Positional arguments forwarded to the event callback.
        **kwargs
            Keyword arguments forwarded to the event callback.

        Returns
        -------
        float or None
            Unix timestamp of the trigger, or ``None`` if the event was not
            found.
        """
        if event not in self._events:
            self.log_error(f'Attempted to trigger <{
                           event}> which is not a registered event.')
            return
        t = self.log_debug(f'Event <{event}> has been triggered')
        try:
            self._events[event](self, *args, **kwargs)
        except TypeError:
            self.log_error(f'Failed to trigger <{event}> due to incorrect arguments. Event callbacks should accept segment as first argument then *args **kwargs.')
        self.event_records.append({'time': t,
                                   'event': event,
                                   'args': args,
                                   'kwargs': kwargs})
        self.session.eventlist.append({'segment': self.uniqueid,
                                       'time': t,
                                       'event': event,
                                       'args': args,
                                       'kwargs': kwargs})
        return t
    

                
@dataclass
class Task(BehaviorSegment):
    """
    Implements methods for trawling for trial data and extracting aggregate information
    """
    _LOOP_DELAY : float = 0.01
    def execute(self):
        """
        Run the setup-loop cycle, polling stop conditions each iteration.

        Calls :meth:`setup` once, then repeatedly calls :meth:`loop` until
        the session reports one or more stop conditions.
        """
        self.setup()
        while True:
            self.stop_reasons += self.session.evaluatestopconditions()
            if self.stop_reasons:
                break
            self.loop()
            time.sleep(self._LOOP_DELAY)
    
    @abstractmethod
    def setup(self):
        """Perform one-time initialisation before the main loop."""
        pass

    @abstractmethod
    def loop(self):
        """Execute a single iteration of the segment's main loop."""
        pass


@dataclass
class Trial(BehaviorSegment):
    def execute(self):
        """
        Run the trial, calling :meth:`setup` once and :meth:`perform` until completion.
        """
        self.setup()
        self.perform()
        self.log_trial()
        self.finish()
    
    @abstractmethod
    def setup(self):
        """Perform one-time initialisation before the main loop."""
        pass

    @abstractmethod
    def perform(self):
        """Trial Logic goes here."""
        pass
    
    def waitfor(
        self,
        condition,
        fallback_value=None,
        timeout=None,
        poll_interval=0.05,
        while_waiting=None,
        while_waiting_interval=0.5,
    ):
        if poll_interval <= 0:
            raise ValueError("poll_interval must be > 0")
        if while_waiting is not None and while_waiting_interval <= 0:
            raise ValueError("while_waiting_interval must be > 0")
    
        now = time.monotonic
        deadline = None if timeout is None else now() + timeout
        next_callback = None if while_waiting is None else now() + while_waiting_interval
    
        while True:
            result = condition()
            if result:
                return result
    
            current = now()
            if deadline is not None and current >= deadline:
                return fallback_value
    
            if next_callback is not None and current >= next_callback:
                while_waiting()
                next_callback = current + while_waiting_interval
                continue
    
            sleep_for = poll_interval
            if deadline is not None:
                sleep_for = min(sleep_for, deadline - current)
            if next_callback is not None:
                sleep_for = min(sleep_for, next_callback - current)
    
            time.sleep(max(0, sleep_for))

    
    def wait(self, duration: float):
        """
        Block for a specified duration.

        Parameters
        ----------
        duration : float
            Time to wait in seconds.
        """
        self.log_debug(f'Waiting for {duration} seconds...')
        time.sleep(duration)
        self.log_debug('Stopped waiting')
        
def TrialToTask(trial_cls):
    """Return a Task class that runs ``trial_cls`` once per loop."""
    if not issubclass(trial_cls, Trial):
        raise TypeError('TrialToTask expects a Trial subclass')

    @dataclass
    class GeneratedTask(Task):
        trial_kwargs: dict = field(default_factory=dict)

        def setup(self):
            pass

        def loop(self):
            trial = trial_cls(parent=self, **self.trial_kwargs)
            trial.run()

    GeneratedTask.__name__ = f'{trial_cls.__name__}Task'
    GeneratedTask.__qualname__ = GeneratedTask.__name__
    GeneratedTask.__module__ = trial_cls.__module__
    return GeneratedTask
```
