## Extended Data for "A Low-Cost, Modular Hardware and Software Platform for Head-Fixed Mouse Decision-Making Tasks": paradigm.py

```
# -*- coding: utf-8 -*-
"""
Created on Thu Jun 26 17:31:16 2025

@author: mm4114
"""
from dataclasses import dataclass
from lampyr.segments.abstract import Segment
from abc import abstractmethod

@dataclass
class ParadigmSegment(Segment):
    """
    Base class for paradigm-level segments (Paradigm, Stage).

    Adds a ``paradigm_tag`` label and a ``_paradigmdata`` reference that is
    shared across the paradigm hierarchy.

    Attributes
    ----------
    paradigm_tag : str or None
        Label used to key into the mouse's ``paradigm_stage`` and
        ``properties`` dicts.  Defaults to ``slug`` if not set.
    paradigmdata : dict or None
        Reference to the paradigm's persistent data dict from the mouse,
        populated by the root :class:`Paradigm`.
    """
    paradigm_tag : str = None
    paradigmdata : dict = None
    
    def __post_init__(self):
        """
        Set ``paradigm_tag`` to ``slug`` if not explicitly provided.
        """
        super().__post_init__()
        if self.paradigm_tag is None:
            self.paradigm_tag = self.slug

    def _configure(self):
        """
        Add ``paradigmdata`` and ``paradigm_tag`` to the parent-inherit list.
        """
        super()._configure()
        self._parent_inheritproperties += ['paradigmdata',
                                           'paradigm_tag']

class Stage(ParadigmSegment):
    """
    Represents one stage within a Paradigm.

    A Stage runs a single task session and then applies shaping logic to
    determine whether the mouse should advance to the next stage.  Subclasses
    implement :meth:`define_sessionparams`, :meth:`define_task`, and
    :meth:`define_shaping`.
    """

    def execute(self):
        """
        Run the stage: set session parameters, run the task, then apply shaping.

        ``KeyboardInterrupt`` during :meth:`define_task` is caught and logged
        so that shaping logic in :meth:`define_shaping` still runs on forced
        exits.
        """
        self.define_sessionparams()
        global_paradigm_data = self.get_globalparadigmdata()
        self.define_globalshaping(global_paradigm_data)
        try:
            stage_paradigm_data = self.paradigmdata.get(self.slug, {})
            self.define_task(stage_paradigm_data)
        except KeyboardInterrupt:
            self.log_error('Detected user initiated force quit.')
            self._post_execute()
            raise
        self._post_execute()
            
    def _post_execute(self):
        self.log_notice('SESSION SUMMARY:')
        self.session_summary()
        if self.paradigmdata is not None:
            stage_paradigm_data = self.paradigmdata.get(self.slug, None)
            if stage_paradigm_data is None:
                stage_paradigm_data = {}
                self.paradigmdata[self.slug] = stage_paradigm_data
            self.log_notice('APPLYING SHAPING PROTOCOL:')
            self.define_shaping(stage_paradigm_data)
            global_paradigm_data = self.get_globalparadigmdata()
            self.define_globalshaping(global_paradigm_data)
        else:
            self.log_error('Paradigm Data does not exist. Shaping cannot occur.')
    
    @abstractmethod
    def define_sessionparams(self):
        """Set session limits and minimums via :meth:`set_sessionparam`."""
        pass

    @abstractmethod
    def define_task(self, stage_data):
        """Instantiate and run the task(s) for this stage."""
        pass
    
    @abstractmethod
    def session_summary(self):
        pass

    @abstractmethod
    def define_shaping(self, stage_data):
        """Evaluate session outcome and update shaping state or advance stage."""
        pass
    
    def define_globalshaping(self, global_data):
        pass
    
    def get_globalparadigmdata(self):
        if 'global' not in self.paradigmdata:
            self.paradigmdata['global'] = {}
        return self.paradigmdata['global']
    
    def set_sessionparam(self, param, value):
        """
        Set a session parameter only if it has not already been overridden.

        If the session attribute is currently ``None``, it is set to
        ``value``.  If it already has a value, a warning is logged instead
        so the user is aware of the override.

        Parameters
        ----------
        param : str
            Session attribute name (e.g. ``'duration_limit'``).
        value : object
            Value to assign.
        """
        current_val = getattr(self.session, param)
        if current_val is not None:
            self.log_warning(f'Stage parameter {param} was overridden by user to be {current_val}. If this is not intentional, please quit and remove the override.')
        else:
            setattr(self.session, param, value)
    
    def searchsubsegments(self, *args, **kwargs):
        segments = []
        for child_id in self.subdata:
            segments.extend(self.session.search(root=child_id, *args, **kwargs))
        return segments
    
    def summarize_reportsinsegments(self, report, segmentlist):
        reportcounts = {}
        for seg_id in segmentlist:
            seg = self.session.segments[seg_id]
            val = seg['reports'].get(report, 'NO REPORT FOUND')
            if val not in reportcounts:
                reportcounts[val] = 0
            reportcounts[val] += 1
        return reportcounts


@dataclass
class Paradigm(ParadigmSegment):
    """
    Root-level segment that orchestrates stage selection and session history.

    A Paradigm must always be the highest-ranked segment (rank 0) and must
    be run within a Lampyr instance.  It initialises per-mouse paradigm data
    from ``mouse.properties``, then delegates to the appropriate
    :class:`Stage` based on the mouse's current progress.

    Class Variables
    ---------------
    DEFAULT_PROPERTIES : dict
        Defines the initial structure of the per-mouse paradigm data dict.
        Subclasses must override this.
    STAGES : list of str
        Ordered list of stage name strings for this paradigm.

    Attributes
    ----------
    sessionhistory_sessionlimit : int
        Maximum number of past sessions to load when reviewing history.
    sessionhistory_dayslimit : int
        Maximum age (days) of past sessions to consider.
    _paradigmdata : dict or None
        Reference to ``mouse.properties[slug]``, populated during
        :meth:`execute`.
    """
    paradigmdata : dict = None
    stagelist : tuple = None
    
    def __post_init__(self):
        """
        Validate that a lampyr instance and DEFAULT_PROPERTIES are present.

        Raises
        ------
        RuntimeError
            If the paradigm is not at rank 0, if no lampyr instance is
            attached, or if ``DEFAULT_PROPERTIES`` has not been overridden.
        """
        super().__post_init__()
        if self.rank != 0:
            raise RuntimeError(f'Attempted to create paradigm {self.name} at a low rank. Paradigms must always be the highest ranked segment.')
        if self.lampyr is None:
            raise RuntimeError('Attempted to run paradigm outside of lampyr.')
        if self.stagelist is None:
            raise RuntimeError('stagelist is undefined')
    
    def execute(self):
        if self.slug not in self.mouse.properties:
            self.mouse.properties[self.slug] = {'stage' : None}
        self.paradigmdata = self.mouse.properties.get(self.slug,{})
        self._createstagemap()
        stageid = self.paradigmdata.get('stage', self._defaultstage)
        if stageid is None:
            stageid = self._defaultstage
            self.paradigmdata['stage'] = self._defaultstage
        if stageid not in self.paradigmdata:
            self.paradigmdata[stageid] = {}
        self.log_notice(f'Mouse is in {stageid} stage')
        StageClass = self._stagemap[stageid]
        stage = StageClass(parent=self)
        try:
            stage.run()
        except KeyboardInterrupt:
            self.log_notice(f'CHECKING FOR PROGRESSION CRITERIA ({stageid}):')
            self.define_progression(StageClass, self.paradigmdata[stageid])
            newstage = self.paradigmdata.get('stage', self._defaultstage)
            if newstage != stageid:
                self.log_notice('Next session will be {newstage}')
            raise
        self.log_notice(f'CHECKING FOR PROGRESSION CRITERIA ({stageid}):')
        self.define_progression(StageClass,  self.paradigmdata[stageid])
        newstage = self.paradigmdata.get('stage', self._defaultstage)
        if newstage != stageid:
            self.log_notice(f'Next session will be {newstage}')
    
    def _createstagemap(self):
        self._stagemap = {}
        self._defaultstage = self.stagelist[0].slug
        for stage in self.stagelist:
            if stage.slug in self._stagemap:
                raise RuntimeError(f'Detected duplicate {stage.slug} stages')
            self._stagemap[stage.slug] = stage
    
    @abstractmethod
    def define_progression(self, current_stage, stage_data):
        pass
    
    def progress(self):
        stageid = self.paradigmdata.get('stage', self._defaultstage)
        for ind, stageclass in enumerate(self.stagelist):
            if stageid == stageclass.slug:
                break
        newstage = self.stagelist[ind+1].slug
        self.paradigmdata['stage'] = newstage
        self.log_debug(f'Progressed Paradigm Stage from {stageid} to {newstage}')
    
    def setstagebyclass(self, stageclass):
        self.setstagebyslug(stageclass.slug)
    
    def setstagebyslug(self, slug):
        if slug not in self._stagemap:
            raise RuntimeError(f'{slug} not in stagemap')
    
    def _configure(self):
        super()._configure()
        self._dump_reducetorepresentations += ['stagelist']
```
