## Extended Data for "A Low-Cost, Modular Hardware and Software Platform for Head-Fixed Mouse Decision-Making Tasks": lampyr_setup.bat

```
taskkill /F /IM python.exe
taskkill /F /IM pythonw.exe
rmdir /s /q "%LOCALAPPDATA%\miniforge3\envs"
mkdir "%LOCALAPPDATA%\miniforge3\envs"
call conda activate
call conda env create -f https://raw.githubusercontent.com/mxwllmadden/Lampyr/main/mx_hardware.yaml -y
call conda activate lampyr
call conda install -y git
pip install "git+https://github.com/mxwllmadden/Lampyr.git@main"
```
