## Extended Data for "A Low-Cost, Modular Hardware and Software Platform for Head-Fixed Mouse Decision-Making Tasks": lampyr_update.bat

```
@echo off

call conda activate
call conda activate lampyr
call conda env update --name mx_hardware --f https://raw.githubusercontent.com/mxwllmadden/Lampyr/main/mx_hardware.yaml --prune
pip install art
pip uninstall lampyr -y
pip install --no-deps "git+https://github.com/mxwllmadden/Lampyr.git@main"
lampyr go
cmd /k
```
