## Extended Data for "A Low-Cost, Modular Hardware and Software Platform for Head-Fixed Mouse Decision-Making Tasks": pyproject.toml

```
[project]
name = "Lampyr"
version = "0.0.1"
description = "NA"
authors = [
    { name = "Maxwell Madden", email = "" }
]
dependencies = [
    "pyserial",
    "click",
    "textual",
    "art",
    "numpy>=1.21",
    "pandas",
    "h5py",
    "requests",
]
readme = ""
requires-python = ">=3.10"

[project.optional-dependencies]
video = [
    "opencv-python",
]

[project.urls]
Homepage = "https://github.com/mxwllmadden"

[project.scripts]
lampyr = "lampyr.interfaces.click_cli.app:cli"

[tool.setuptools.packages.find]
where = ["."]
include = ["lampyr", "lampyr.*"]

[tool.setuptools.package-data]
"lampyr" = ["**/*.tcss", "**/*.css"]
```
