## Extended Data for "A Low-Cost, Modular Hardware and Software Platform for Head-Fixed Mouse Decision-Making Tasks": README.md

# LAMPyR

LAMPyR is a Python framework for running and managing behavioral experiment rigs. It provides a command-line interface and a Textual terminal UI for configuring rig hardware/interfaces, tracking mouse performance, running automated behavioral training paradigms, and has a notification manager to alert experimenters to session completion.

## Features

- Click-based `LAMPyR` command-line interface
- Textual TUI launcher (`LAMPyR go`)
- Rig configuration and calibration helpers
- Mouse creation, retirement, paradigm assignment, and run history
- Session and mouse data persistence using JSON/HDF5-backed utilities
- Optional Pushover notifications
- Hardware build files in `hardware/`
- Arduino firmware sketches for Bandit rig variants in `firmware/`

## Documentation

- [CLI command reference](docs/cli_commands.md)
- [GUI workflow](docs/gui_workflow.md)
- [Application architecture and segment inheritance](docs/app_architecture_and_segments.md)
- [Training management](docs/training_management.md)

## Requirements

- Windows is recommended for hardware/TUI operation; some rig and touch-overlay features are Windows-specific.
- Linux is supported for CLI operation with a Linux compatible TUI planned soon.
- Conda or Miniforge/Miniconda
- Python 3.10 or newer; the provided Conda environment uses Python 3.12
- Git, if installing directly from GitHub
- Arduino drivers/firmware as required by your rig hardware

## Installation

### Recommended: Conda environment from this repository

From the repository root:

```bat
conda env create -f mx_hardware.yaml
conda activate LAMPyR
pip install .
```

Verify the install:

```bat
LAMPyR --help
LAMPyR info
```

### Developer install

Use an editable install if you plan to modify the source code:

```bat
conda env create -f mx_hardware.yaml
conda activate LAMPyR
pip install -e .
```

### Install directly from GitHub

```bat
conda create -n LAMPyR python=3.12 -y
conda activate LAMPyR
conda install -c conda-forge git -y
pip install "git+https://github.com/mxwllmadden/LAMPyR.git@main"
```

### Optional video support

If you install without the provided Conda environment and need OpenCV/video support:

```bat
pip install "LAMPyR[video]"
```

For a local checkout, use:

```bat
pip install ".[video]"
```

## Windows setup scripts

This repository includes batch files for convenience:

- `LAMPyR_setup.bat` - installs/refreshes a LAMPyR Conda environment from the GitHub `main` branch.
- `LAMPyR_update.bat` - updates dependencies and reinstalls LAMPyR from GitHub.
- `install.bat` - local environment/install helper.

Review these scripts before running them, especially on machines with existing Conda environments.

## Configuration

LAMPyR stores user configuration under:

```text
%LOCALAPPDATA%\LAMPyR\config.json
```

Default settings include a shared mouse data directory:

```text
N:/SHARED/Maxwell_LAMPyR_MouseData
```

Use the CLI to inspect and configure the installation:

```bat
LAMPyR info
LAMPyR rig info
LAMPyR rig configure
LAMPyR rig calibrate
```

To reset configuration to defaults:

```bat
LAMPyR reset
```

## Basic usage

Launch the terminal UI:

```bat
LAMPyR go
```

List available behaviors:

```bat
LAMPyR list
```

Create and inspect a mouse:

```bat
LAMPyR mouse create MOUSE_ID --paradigm Bandit
LAMPyR mouse info MOUSE_ID
LAMPyR mouse list
```

Set or inspect a mouse paradigm/stage:

```bat
LAMPyR mouse paradigm MOUSE_ID
LAMPyR mouse paradigm MOUSE_ID PARADIGM_NAME --stage STAGE_NAME
```

Run a behavior for a mouse:

```bat
LAMPyR mouse run MOUSE_ID
LAMPyR mouse run MOUSE_ID BEHAVIOR_NAME
```

Run a behavior directly:

```bat
LAMPyR run BEHAVIOR_NAME
```

Common stop-condition options include:

```text
--duration_limit / -dl
--trial_limit / -tl
--reward_limit / -rl
--merit_limit / -ml
--demerit_limit / -dml
```

Use command help for the full option list:

```bat
LAMPyR mouse run --help
LAMPyR run --help
```

## Notifications

LAMPyR supports Pushover notifications. Configure an app token and users with:

```bat
LAMPyR user set-token TOKEN
LAMPyR user create USERNAME --pushover_user_key USER_KEY
LAMPyR user list
LAMPyR user ping --user USERNAME --message "LAMPyR notification test"
```

## Hardware build files

Hardware files for building LAMPyR rigs are in `hardware/`:

- `hardware/BOM_Hardware.xlsx` - hardware bill of materials.
- `hardware/3d models/` - printable 3D model files.
- `hardware/Custom PCB shield/` - PCB shield fabrication files, BOM, designators, and pick-and-place positions.
- `hardware/eMachineShop design files/` - eMachineShop design files for machined components.

## Firmware

Arduino sketches for supported rig variants are in `firmware/`. Flash the sketch matching your rig hardware before attempting to run sessions.

## Project layout

```text
LAMPyR/                 Python package
LAMPyR/interfaces/      CLI and Textual TUI entry points
LAMPyR/managers/        Data, rig, mouse, plugin, and notification managers
LAMPyR/segments/        Segment, task, trial, stage, and paradigm abstractions
LAMPyR/behaviors/       Behavior implementations
LAMPyR/analysis/        Data analysis helpers
hardware/               Hardware BOM, 3D models, PCB shield files, and machining files
firmware/               Arduino firmware sketches
mx_hardware.yaml        Conda environment definition
```

## LLM Assisted Coding Disclosure

Some code in this repository was generated or written with the assistance of an LLM coding agent. ALL code has been tested, read, and verified by Maxwell Madden.

Specifically, LLMs were used to generate the LAMPyR.analysis.colony query based session retreival code and the 'LAMPyR go' GUI. All other code was manually written.

## License

This project is distributed under the MIT License.
